# Management recommendations for Alpine protection forests: the importance of regeneration quality and initial stand composition

**DOI:** 10.1101/2024.04.25.591179

**Authors:** Ueli Schmid, Monika Frehner, Harald Bugmann

## Abstract

The protection of infrastructure against gravitational natural hazards is one of the most important ecosystem services (ES) of mountain forests in Alpine countries. For a continuous provision of this ES, forests need to have a high protective effect, e.g., high canopy cover and/or stem numbers, while being resistant to and resilient after disturbances by being well-structured and stable, having a species composition adapted to the local site conditions and sufficient regeneration, all on a relatively small spatial scale.

While “natural” forests may fulfill these prerequisites without human intervention, management history and high levels of ungulate browsing have produced unsustainable stand structures in many protection forests that need to be improved by management. The general principles of protection forest management are well established, but there are no quantitative, science-based recommendations for management regimes, i.e., specific sequences of interventions, that ensure a continuous protective quality. Our goal was to derive such recommendations for different stand types across three elevational zones, from mixed forests of the upper montane to spruce forests of the subalpine zone.

We used an updated version of the model ProForM to simulate stand development under different levels of ungulate browsing, testing a large number of management regimes that vary in the spatial aggregation of tree removal, the intensity and interval of the interventions. We investigated the influence of browsing pressure and management on the protective quality using Boosted Regression Trees and Beta regression.

High levels of ungulate browsing had such a strong negative effect on the protective quality that it could not be improved through forest management. This underlines the need for maintaining ungulate densities in Alpine forests at levels that allow for the successful regeneration of all key tree species. In stands that are influenced less by browsing, the protective quality can be improved through management in many cases, with specific management recommendations differing mostly depending on the initial stand conditions and, to a lesser extent, on the elevational zone. Well-structured stands provide a high protective quality without management interventions during at least a century across all elevational zones. In young and in mature stands, we generally recommend management regimes with relatively long return intervals of 30 to 40 years and low intervention intensities of 10 to 20% basal area removal.

## 1. Introduction

Mountain forests are an important habitat for a multitude of species and provide crucial ecosystem services (ES) such as timber production, carbon storage, scenic beauty, and protection form natural hazards (EEA, 2010). Many forests provide multiple ES simultaneously, a feature known as multifunctionality. In mountainous regions, protection against natural hazards is one of the most important forest ES (Wehrli et al., 2007a). These so-called protection forests reduce the risks to human lives and infrastructure resulting from gravitational natural hazards such as avalanches, rockfall, landslides, erosion or debris flows by lowering the probability of mass movements being initialized, or by reducing their energy during transit (Brang, 2001; Dorren et al., 2004). The degree to which a forest mitigates these risks depends on the intensity of the hazard process, the length of the forested slope between the source and the infrastructure, and the state of the forest itself.

The most important stand characteristics providing a high protective effect are a large basal area and/or stem numbers – especially in rockfall protection forests where trees act as obstacles to falling rocks – and a high canopy cover, as gaps (1) are potential release zones for avalanches due to a more homogeneous snow pack and (2) feature a reduced or missing root reinforcement of the soil, which enhances landslide risk. As the protective function should be continuous on a relatively small spatial scale, stands in protection forests need to be resistant and resilient to biotic and abiotic disturbances (Brang, 2001; Dorren et al., 2004; Frehner et al., 2005). Resistance is associated with stand stability, either through individual tree stability or collective stability, a diverse diameter distribution, and a close-to-natural species composition. Resilience is mainly determined by the abundance and composition of regeneration, ensuring the potential for a rapid recovery of the protective function after a major disturbance.

The relevant characteristics for the protective function of a forest can be influenced by management, but they are also influenced by natural factors. One of the main sylvicultural challenges in determining appropriate management regimes are the trade-offs between the current protective quality and the sustainability of stand structure: while the current protective effect is usually highest in dense stands, sufficient light and therefore canopy gaps need to be available to ensure continuous regeneration. The ideal structure of a protection forest is therefore often multi-aged with continuous canopy cover (O’Hara, 2006), and managers strive to maintain forest structure within limits that may be narrower than those resulting from natural dynamics (Brang, 2001; Brang et al., 2006).

A key challenge for the management of protection forests in the European Alps are the high ungulate densities that locally influence the abundance and composition of regeneration negatively (Ammer, 1996; Brang, 2017; Kupferschmid et al., 2020; Schulze et al., 2014). Browsing is highly selective regarding tree species (Motta, 1998), often resulting not only in a reduced density of regeneration, but also lower species richness among the regeneration (Ramirez et al., 2018; Schulze et al., 2014). In the long term, even the total absence of heavily browsed tree species in the upper canopy can result (Kupferschmid et al., 2019). Some pivotal tree species of Alpine forests in general and protection forests in particular, such as *Abies alba, Acer pseudoplatanus*, or *Sorbus aucuparia*, are especially prone to browsing (Kupferschmid et al., 2015; Motta, 1998). High browsing pressure is thus expected to have a negative effect on the protective quality of forests. Only few studies have addressed the impact of browsing on the protective function (Didion et al., 2009; Thrippleton et al., 2020; Wehrli et al., 2007b; Woltjer et al., 2008), all of them confirming the negative effect. However, none of them have investigated the implications of protection forest management under different browsing pressures.

Forests, and particularly mountain forests, develop slowly and natural hazard events are usually infrequent. Therefore, empirical research on the impact of forest structure and management on the protective quality is scarce (but see, e.g., Bebi et al., 2015; Ringenbach et al., 2022). In some studies, natural hazard processes in forests were modeled using measured stand data (e.g., Caduff et al., 2022; Schönenberger et al., 2005; Vacchiano et al., 2015), but most studies relied on simulations of forest dynamics (e.g., Elkin et al., 2013; Maroschek et al., 2015; Mina et al., 2017; Moos et al., 2021). To quantify the protective effect of simulated stands, two main approaches are used: (1) coupling forest simulation models with models simulating the natural hazard process itself (e.g., Dupire et al., 2016; Fuhr et al., 2015; Moos et al., 2021; Scheidl et al., 2020) and (2) deriving the protective quality from simulated stand structures using linker functions or indices (Blattert et al., 2017; Cordonnier et al., 2014).

Many of these indices are based on the Swiss framework for assessing the quality of protection forests and deriving suitable management measures (abbreviated as “NaiS”; Frehner et al., 2005). The assessment process in NaiS is based on a comparison of the current forest state to target states, so-called profiles, that describe forest states which are expected to provide a good protective quality, and to be resistant to and resilient after possible disturbances. The profiles differ by site type and the prevailing natural hazard, and they encompass threshold values for a range of stand characteristics such as species mixture, structure, and regeneration. Most linker functions commonly used to assess the protective quality of simulated stands focus on the current protective effect (i.e., stand density, canopy cover and gaps, and/or diameter distribution) while neglecting crucial aspects of resistance and resilience, such as species composition, stand stability, and particularly the abundance of regeneration (Elkin et al., 2013; Hillebrand et al., 2023; Irauschek et al., 2017a; Maroschek et al., 2015; Mina et al., 2017; Pardos et al., 2017; Thrippleton et al., 2021). In contrast, the recently developed model ProForM (Schmid et al., 2023) assesses the protective quality by closely emulating the system NaiS, determining the discrepancies between the stand’s state and the target profiles in six categories: species mixture, vertical and horizontal arrangement, stand stability, and seedling and sapling abundance and composition.

The necessity and type of management as well as its intensity (e.g., volume to be removed, return interval) that are required to maintain the protective function of mountain forests have been addressed on a general level (Brang, 2001; Brang et al., 2006; Dorren et al., 2004). There, it was often concluded that stands that exhibit a close-to-natural composition and structure are usually able to provide a good and continuous protective effect and have a high resistance to and resilience after natural disturbances, and therefore do not necessarily have to be managed. However, many of our current protective forests are heavily influenced by past management that often has led to high stand densities, uniform structures and/or a low or even missing regeneration layer. While such stands may currently provide a high protective effect, it is unsustainable in the long term. In these cases, management interventions mimicking small-scale natural disturbances should be applied while working in line with natural forest dynamics as much as possible.

On the stand and landscape scales, the effects of different management options on the protective effect have been investigated with simulation studies. There, initial forest conditions were usually based on the current situation in the respective study area and can therefore be expected to not represent ideal protection forest structures. Nevertheless, many studies have concluded that, under current climate, not intervening was the best management strategy over the course of roughly one century (Irauschek et al., 2017a, 2017b; Maroschek et al., 2015; Mina et al., 2017; Pardos et al., 2017; Scheidl et al., 2020). Only a few studies found management schemes specifically designed for protection forests to enhance the protective effect compared to a no-intervention scenario (Rammer et al., 2015; Woltjer et al., 2008). However, these studies did neither include measures of resistance and resilience to assess the protective quality, as mentioned above, nor explicitly include disturbance events. In one of the few studies that did include measures of resilience, Thrippleton et al. (2020) found previously managed forests to be more resilient after large disturbances because managed forests feature abundant advance regeneration, i.e. regeneration that is present prior to the disturbance.

In most studies, the range of management options considered was relatively narrow. Usually, they included a no-management and a business-as-usual scenario (Irauschek et al., 2017a; Maroschek et al., 2015; Scheidl et al., 2020; Thrippleton et al., 2020). In some cases, one or two additional alternative management regimes were investigated (Mina et al., 2017; Pardos et al., 2017; Rammer et al., 2015; Woltjer et al., 2008). Only in very few studies, management parameters were varied systematically, e.g., intervention intensity (Thrippleton et al., 2023, 2021) or the spatial pattern of tree removal (Irauschek et al., 2017b). However, the return period and management intensity that is required to warrant the protective effect is important from an economic perspective if harvesting is not profitable, which is often the case in steep mountain terrain and particularly in high-wage countries such as Switzerland. Under such conditions, governmental subsidies are often paid for management geared towards maintaining the protective effect of forests, usually per intervention, with their amount depending on either the volume harvested or the area treated.

Based on these considerations, the overall goal of our study was (a) to investigate the influence of high browsing pressure and forest management on the protective quality of forest stands, and (b) to derive management recommendations regarding the minimum level of intervention to warrant the protective effect of forest stands with respect to two key hazards, i.e., avalanches and landslides/erosion/debris flows. The stands that we investigated in a simulation modeling study span a large environmental gradient, encompassing (1) mixed species forests of the upper montane zone with Norway spruce (*Picea abies* (L.) H. Karst), silver fir (*Abies alba* Mill.), European Beech (*Fagus sylvatica* L.), and sycamore maple (*Acer pseudo-platanus* L.), spruce-fir forests of the high montane zone, and subalpine spruce forests, (2) two site qualities and (3) three initial stand types – young, well-structured, and mature stands. The management regimes vary systematically with regard to the spatial aggregation of tree removal, interval, and intensity. To that end, the forest simulation model ProForM (Schmid et al., 2023) was further developed by adding a range of management options that reflect intervention types frequently used in mountain forest management (single tree selection, group selection, slit cuts, and cable yarding) and by improving the process of assessing the protective quality based on the Swiss system NaiS (Frehner et al., 2005) to render more realistic results.

In particular, we address the following research questions:

1. How do suitable management regimes (i.e., intervention type, interval, and intensity) for protection forests differ between sites of good and low regeneration quality?
2. What are “optimal” management regimes and how do they vary along an elevational gradient, i.e., for different forest types?
3. How do these “optimal” management regimes differ based on the initial stand conditions, i.e., for young, well-structured, and mature stands, and in the short vs. the long term (90 vs. 1’000 years)?

## 2. Material and methods

### 2.1. Simulation model

To simulate forest dynamics and assess the protective quality of stands, we used the dynamic simulation model ProForM (Schmid et al., 2023). ProForM was designed to simulate the dynamics of Swiss mountain forest stands while allowing for different management options. The protective quality of the simulated stands can be assessed based on the Swiss guidelines for protection forest management (NaiS; Frehner et al., 2005). ProForM includes four tree species – Norway spruce (*Picea abies* (L.) H. Karst), silver fir (*Abies alba* Mill.), European beech (*Fagus sylvatica* L.), and sycamore maple (*Acer pseudoplatanus* L.) – and is parameterized for four elevational zones (submontane SM, upper montane UM, high montane HM, and subalpine SA).

The basic entity of the model are cohorts of identical trees that populate cells with known locations whose size allows to accommodate one large individual (40 – 100 m^2^, depending on the elevational zone). The cells are characterized by the state variables species, stem number, diameter at breast height (DBH), height, and height to the crown base. ProForM simulates the demographic processes regeneration, growth, and mortality for each cohort with a temporal resolution of one year, taking competition among neighboring cells into account. It is assumed that the climate is constant and that there are no biotic or abiotic disturbances with impacts beyond single-tree mortality. The model is designed to simulate forest stands of around one to two hectares in size. The model can be initialized either with a fully characterized stand or from bare ground. To assess the protective quality of a simulated stand, its characteristics are compared to so-called “profiles” that describe the minimal and ideal state of a protection forest for a given site type and natural hazard (snow avalanche, landslide/erosion/debris flow, rockfall, and torrents/floods, derived from NaiS). From this comparison, numerical indices in six categories (species mixture, vertical structure, horizontal arrangement, stability of “support trees”, seedlings, and saplings) are calculated that range from −1 (insufficient) across 0 (minimal profile met) to 1 (ideal profile met). Simulated forest dynamics were successfully validated against multiple long-term forest inventory plots in Switzerland (Schmid et al., 2023).

An exhaustive model description is available in Schmid et al. (2023). Here, only the parameters relevant for this study are explained briefly. The two user-defined parameters *site quality* and *regeneration quality*, both dimensionless integer values between 1 (poor) and 5 (good), are used to characterize the aggregated abiotic and biotic growth conditions. *Site quality* influences the growth rates of cohorts larger than 12 cm DBH, whereas *regeneration quality* influences the germination probability, the number of trees at ingrowth (i.e., at a tree height = 130 cm), and the growth of cohorts with DBH < 12 cm. When a new cohort germinates, its species is chosen randomly based on a probability calculated from the stand’s current species composition and user-defined parameters characterizing the species composition of the surrounding forest, as well as the *species pressure*, a measure of the influence of the surrounding stand (e.g., due to the presence of seed trees) vs. the simulated stand itself.

### 2.2. Model development

For this study, several submodels of ProForM were added or developed further, resulting in model version 1.1. Specifically, we (1) added four new management types to the model, (2) improved the algorithm for identifying canopy gaps, (3) adapted the profiles for assessing the protective quality, and (4) slightly relaxed the rules for the assessment process. On a structural level, the process of assessing the protective quality was split into two, with the identification and analysis of gaps now being a separate process. The technical details of the model changes are described in Supplementary material S1. The model code in R (R Core Team, 2021) is freely available on github (Schmid et al., 2024) and includes scripts to run all simulations carried out for this study and to reproduce the graphs of Supplementary materials S3 and S5 as well as Fig. 25. The four modifications are described in detail below.

#### 2.2.1. Management types

To extend the range of possible applications of ProForM, four new management types were implemented in model v1.1: single tree selection (STS), group selection (GRS), slit cuts (SC), and cable yarding (CAB). They share the following parameters that can be specified by the user to control intervention timing and intensity: year of the first intervention, interval between interventions in years, intervention intensity in percent of basal area reduced per intervention, minimum DBH in cm of trees eligible for harvesting, and the minimum basal area shares of each of the four species at the time of intervention below which they will not be eligible for harvesting. When an intervention is scheduled in the model, a list of eligible harvest units (trees, cohorts, or groups of cohorts with spatial arrangements depending on management type and user input) is produced. It is then ordered based on multiple stand characteristics using user-specified weights and a random factor. The weighting factors include species (increasing the selection probability of abundant species to even out species distribution of the remaining stand; not available in CAB), basal area (increasing the selection probability of harvest units with large basal area to spatially concentrate tree removals), crown ratio (increasing the selection probability of harvest units with short crowns to increase the stability of the remaining stand; not available in CAB), abundance of regeneration within the unit (increasing the selection probability of harvest units with advance regeneration to promote existing regeneration; not available in STS), and regeneration in neighboring cells (increasing the selection probability of harvest units with little regeneration in adjacent cells to prevent large continuous gaps from forming; not available in STS). Harvest units are then iteratively selected from the ordered list until the specified management intensity is reached while respecting a user-specified spatial buffer (0-2 cells) between selected units. The process of ordering and iteratively selecting cells to be harvested is largely based on the Uneven-Aged Management Algorithm of the model Samsara2 (Lafond et al., 2014).

The spatial structure of harvest units in each management type can be specified by the user. GRS aggregates groups of one to four adjacent cells (each the size of one large tree) in all possible spatial configurations (Fig. S1.1). In SC, the harvest units consist of slits oriented diagonally to the slope, having a user-defined width and length expressed as the number of cells (Fig. S1.2). In CAB, the user can define up to three columns of cells (directly downslope) in which the cable lines are positioned. Harvest units then consist of slits perpendicular to the cable cells with a user-defined width and length in number of cells (Fig. S1.3). Per intervention, only units belonging to one cable line are eligible for harvesting (rotating regularly between cable lines). Previous to harvesting these units, all trees within the cable slit are removed. In STS, harvest units consist of single trees, and therefore no spatial aggregation is applied.

#### 2.2.2. Gap identification

In the previous version of ProForM, canopy gaps were identified as continuous areas of cells containing no trees with DBH ≥ 12 cm. This caused areas of irregular shapes, e.g., two larger gap areas connected by a string of single cells, to be identified as one single gap (Fig. S1.4b). In practice, such an example would most likely be identified as two separate gaps. The improved gap recognition algorithm in ProForM v1.1 only allows groups of three triangularly arranged cells with no trees with DBH ≥ 12 cm to be part of a gap (Fig. S1.4c).

#### 2.2.3. NaiS profiles

In model version 1.0, there were two sets of NaiS profiles within the elevational zones HM and SA, differentiating between drier and moister site types, with differing threshold (i.e., minimal and ideal) values. As the differences were only marginal, these sets were unified into one per elevational zone, using mean values for the previously different thresholds. Furthermore, the threshold values for the index capturing the complexity of the vertical canopy structure were recalculated. In this index, the share of cohorts in four diameter classes is assessed to ensure a sustainable diameter distribution. The previous threshold formulation used the required canopy cover of saplings and thicket for the lowest diameter class and divided the remaining share equally over the three upper classes. However, preliminary simulations showed that stands in an equilibrium state, i.e., with a sustainable, reverse-J-shaped diameter distribution, had a considerably larger canopy cover in the lowest class than minimally required. Consequently, the shares of the upper three classes were well below the threshold values, and thus an insufficient index value (i.e., below the minimum required) was awarded, even though the diameter distribution was evidently sustainable. Therefore, we derived new threshold values for the upper three classes from stable states of long-term simulations with regular management interventions, differentiating between low and high site qualities. The threshold values for the lowest class remained unchanged, as they were based on theoretical calculations of the minimum necessary amount of regeneration (Brang and Duc, 2002; Frehner et al., 2005; Schmid et al., 2023).

#### 2.2.4. Relaxed assessment rules for protective quality

Preliminary assessments of the protective quality of simulated stands revealed that it was judged overly rigorously, i.e., index values were distinctly lower than experts would expect based on stand characteristics. A detailed analysis of the subindex values in these simulations indicated three main sources of this underestimation, and thus they were slightly relaxed to yield a more realistic assessment result. (1) In many cases, a single gap was found to be present in the stand, but to exceed the threshold value for the maximally allowed gap area in the minimal profile. Given that this gap does not exceed the threshold area by much, most practitioners would not judge this as a violation of the minimal profile. We therefore relaxed the rule for reaching the minimal profile (but not for the ideal profile) in the index for horizontal arrangement, allowing for one gap per stand to exceed the threshold size. (2) With the profile for landslide/erosion/debris flow, the maximally allowed gap size is increased according to NaiS (Frehner et al., 2005) if there is “ensured regeneration” within a gap. In the previous model version, this was assessed based on the cover of regeneration within a gap and its species composition. In many cases, gaps were judged to not contain “ensured regeneration” due to an inadequate species composition. As it is very unlikely, and in practice often unnecessary, to have a balanced species composition on such a small scale of a single gap, this criterion was dropped from the assessment, leaving the “ensured regeneration” to be decided only based on the regeneration cover within the gap. (3) The index for seedlings previously checked the seedling cover within gaps and the number and identity of the species present. It often yielded pessimistic assessments due to the first criterion. In ProForM, germination does not depend on light availability (i.e., competition), i.e., will certainly happen in every empty cohort but is solely driven by model parameters and not the current stand composition. This subindex was therefore dropped from the assessment. However, by assessing the species identity of seedlings, a lack of seedlings is still punished.

### 2.3. Simulation experiments

Short-term (ST) simulations were conducted to study the effects of management on the protective function of typical forest stands over a time scale that is relevant for forest planning. We initialized the model with 36 different stands, each 1 ha in size. These resulted as a combination of three elevational zones (upper montane (UM), high montane (HM), and subalpine (SA)), three stand types (young, well-structured, and mature), two *site qualities* (“good” and “medium” with parameter values of 5 and 3, respectively), and two *regeneration qualities* (“good” and “low”, the latter due to strong browsing pressure, with parameter values of 5/3 and 1, respectively). The detailed methodology for determining the initialization stands is described below (cf. section “Model initialization”). Forest dynamics of each stand were simulated over 90 years, each simulation using one of 96 management regimes. These are a combination of six management types, four return intervals, and four intervention intensities. The management regimes are described in detail below (section “Management regimes”). As a reference, each stand was also simulated without management (NOM). The protective quality throughout the simulation period was then assessed, once for the natural hazard snow avalanches, and once for landslide/erosion/debris flow (detailed profiles can be found in Supplementary material S2). This resulted in a total of 3’492 ST simulations of forest dynamics and 6’984 assessments of the protective quality.

Long-term (LT) simulations were conducted to study the effects of management on the protective function of forest stands in a state of dynamic equilibrium. In general, the simulation experiments followed the same setup as for the ST simulations in terms of stands, management regimes, and assessments of the protective quality. The differences were that (1) only one stand type per elevational zone was used, (2) the simulations were run for 1’000 years, and (3) only the last 100 years of the simulation, i.e., when forest dynamics had reached an equilibrium, were used to assess the protective quality. The reason for not differentiating between stand types was that the initial stand configuration does not have an influence on the equilibrium. In total, 1’164 LT simulations of forest dynamics and 2’328 assessments of the protective quality were carried out. In all simulations, i.e., ST and LT, the following parameters were kept identical: random seed, aspect (235°), slope (36°), and species pressure of the surrounding stand (50%).

### 2.4. Model initialization

Forty-eight fully characterized stands were needed to initialize the model for our simulation experiments. Yet, it was not possible to identify inventories of 48 real stands that would have matched the requirements of our study (i.e., approximate size, species combination, site conditions, and data quality). Therefore, the stands used to initialize the model were created using spin-up simulations. This had the added benefit that there would not be any model artefacts at the beginning of the simulations due to mismatching data combinations, e.g., cohort DBH and H. Three stand types were used to initialize the ST simulations: (1) young, uniform stands (*init-ST-y*) with considerable scope for silvicultural interventions; (2) well-structured stands (*init-ST-s*) that have a sustainable diameter distribution and a good protective quality; and (3) mature stands (*init-ST-m*) with high stem densities and a lack of regeneration, and therefore a need to induce regeneration by management while maintaining stand stability. To initialize the LT simulations (*init-LT*), stands that are at an equilibrium without management were used.

For each combination of elevational zone, site and regeneration quality, two spin-up simulations were run. The first spin-up simulation started from bare ground (BG spin-up) and ran for 1’000 years without management interventions. Stand characteristics at different time steps of these simulations were extracted and used as *init-ST-y*, *init-ST-m*, and *init-LT*. The time steps selected for extracting *init-ST-y* and *init-ST-m* were determined based on expert opinion considering stand characteristics such as diameter distribution, basal area, stem density, and species composition. Preliminary BG spin-up simulations in the elevational zone UM did not yield young stands (*init-ST-y*) with a diverse species composition, as coniferous species clearly outperformed the deciduous ones in the early stages of the simulation that was characterized by barely any competition. Therefore, one intervention was scheduled between the years 60 and 160, depending on site conditions, to promote deciduous species. These spin-up simulations were then run for an additional 1’000 years without further interventions. For *init-LT*, the last time step of the BG spin-up simulations was used. The second spin-up simulation was initialized with *init-LT* and run for 1’000 years with regular management interventions. The management regime consisted of the management type GRS with aggregates of three cells, return periods of 20 to 45 years and intervention intensities of 15 to 30% basal area removal per intervention. These combinations were chosen based on expert opinion, considering elevational zone and site conditions, with the aim of representing a typical regime applied to protection forests in practice. Out of the last 100 simulation years (when equilibrium had been reached), the time step with the best protective quality was extracted and used as *init-ST-s*. To condense the protective quality, which is assessed in Pro-ForM by six indices, into a single numerical score, the mean of all indices per year was calculated for both natural hazards, and added up. The characteristics of all stands used for model initialization are described in detail in Supplementary material S3.

The parameter values for the species composition of the surrounding forest were determined in preliminary simulations. For each combination of elevational zone (except for SA with only spruce), site, and regeneration quality, the parameter value combinations were determined manually. For stands with a good regeneration quality, the goal was to produce stands with a close-to-ideal composition according to the NaiS profiles when a typical management scheme is applied. Therefore, we ran simulations from bare ground over 1’000 years, applying a GRS management type with aggregates of three cells, return periods of 20 to 30 years, and intervention intensities of 20 to 30% basal area removal per intervention, and assessed the species composition over the last 200 years. We then compared the mean species shares of trees above and below the threshold DBH of 12 cm to the NaiS profiles and iteratively adjusted the parameters for species shares of the surrounding forest in order to approximate the ideal profile as good as possible. Stands with low *regeneration quality* were meant to represent conditions with high browsing pressure. Ungulates have preferences for certain tree species, which can lead to their loss (Heuze et al., 2005). In the case of the four species parameterized in ProForM, silver fir and sycamore maple are clearly preferred over spruce and beech (Kupferschmid and Brang, 2010). However, this aspect is not implemented in ProForM yet, as a low regeneration quality decreases regeneration growth and abundance equally for all species. Therefore, the species shares of silver fir and sycamore maple were set to 0% in simulations with the poorest regeneration quality, resulting in pure spruce stands in the elevational zones HM and SA. For the elevational zone UM, the same iterative process of simulations and assessments of species composition as described above was conducted with the aim of reaching a balanced composition between spruce and beech.

### 2.5. Management regimes

A set of 96 management regimes was compiled for this study. We combined six management types, four return intervals (10, 20, 30, and 40 years), and four intervention intensities (10, 20, 30, and 40% basal area reduction per intervention). The management types differ in the degree of spatial aggregation of the removed trees and were, in increasing order of aggregation: single-tree selection (STS), group selection with aggregates of two (GRS1) and four cells (GRS2), cable yarding with an approximate distance between cable lines of 30 m (CAB1) and 50 m (CAB2), and slit cuts (SC) with slits of approximately 1.5 times 0.5 tree heights.

The minimum diameter for trees to be harvested was set to 12 cm DBH, and the species shares below which a species is exempt from the current intervention were set at the minimal threshold values defined in the ideal NaiS profiles. The factors by which potential harvest units are ordered and selected during each intervention were set in the following, decreasing order: basal area, regeneration within the harvest unit, crown ratio, regeneration in neighboring cells (the latter two with equal weights), and species. The first intervention was scheduled for simulation year 1 in all cases. The detailed model settings for each management type are provided in Supplementary material S4.

### 2.6. Statistical analysis

We analyzed the effects of site (elevational zone, stand type, site and regeneration quality) and management regime (management type, return interval, and intervention intensity) on the protective quality using Boosted Regression Trees (BRT) and beta regression. To quantify the aggregated protective effect of each stand, two sets of target variables were derived from the six indices of protective quality, with one target variable each based on the minimal NaiS profile and one on the ideal profile.

In a first step, separate statistical models for each explanatory variable were built for ST and LT simulations and both natural hazards, i.e., snow avalanches (A) and landslides/ersion/debris flow (LED), using all explanatory variables and the complete set of simulation results (“full models”, 16 models per method). In a second step, only those simulations with good regeneration quality were retained (“reduced models”). From the ST simulation outputs, reduced models were built for each combination of natural hazard (A and LED), elevational zone (UM, HM, and SA), and stand type (*init-ST-y, init-ST-s*, and *init-ST-m*), yielding 71 models per method. From the LT simulation outputs, the reduced models were built for each combination of natural hazard and elevational zone, yielding 24 models per method. The analyses were carried out using the R statistics software version 4.0.5 (R Core Team, 2021). The code to reproduce the analysis and all figures is freely available on github (Schmid, 2024).

The first set of target variables for the statistical analysis was the share of indices meeting either the minimal or the ideal profile (*sMP_abs_* and *sIP_abs_*, respectively), i.e., the “absolute” protective effect of a stand. For every time step in the analysis period (time steps 901 to 1’000 for LT and 1 to 90 for ST simulations), the share of the six indices meeting the minimal profile (index value ≥ 0) and the ideal profile (index value = 1), respectively, was calculated and the resulting shares averaged over the analysis period. The second set of target variables was the difference between *sMP_abs_* and *sIP_abs_* of the simulations with management regimes applied and the corresponding NOM simulation (*sMP_diff_* and *sIP_diff_*, respectively), i.e., the impact of management on the protective function. In a preparatory step, missing values in the set of indices had to be removed. Missing values can occur in two indices of protective quality: In the index for seedlings when there are no gap cells present at a certain time step and thus the availability of regeneration cannot be checked; and in the index for the stability of support trees when no conifers are among the support trees, as in the NaiS profiles threshold values are defined for spruce and silver fir only. In these cases, an index value of zero, corresponding to the minimal profile, was enforced.

Boosted Regression Trees (BRT) were used to assess the relative importance of the explanatory variables on the protective effect of a stand. BRT are a machine learning technique that combines regression tree models and boosting algorithms (Elith et al., 2008); they are increasingly used in ecological studies (e.g., Schuler et al., 2019; Seltmann et al., 2021; Szwagrzyk et al., 2021; Thom and Seidl, 2021). A distinct advantage of BRT is that they can handle different types and distributions of target and explanatory variables without prior transformation, as well as diverse and complex types of data relationships, including non-linearities and interactions, without the need of specifying them *a priori*. BRT quantify the relative significance of the explanatory variables based on the frequency of their selection for tree splitting and the model improvement as a result of these splits. To set up the BRT models, we followed Elith et al. (2008) and used the R libraries *gbm* (Greenwell et al., 2022) and *dismo* (Hijmans et al., 2022). We used a fixed random seed, a tree complexity of 5 for the full and 3 for the reduced models, and a bag fraction of 0.75. The learning rate was chosen individually for each model in the range of 0.001 and 0.1, so as to have at least 1’000 trees.

We applied beta regression models (Cribari-Neto and Zeileis, 2010) to further identify the effects of the individual explanatory variables. Beta regression can be used to model linear effects of explanatory variables on target variables whose values lie within the standard unit interval (0, 1). As *sMP_abs_* and *sIP_abs_* can take values of 0 and 1, we used the transformation (*y* · (*n* − 1) + 0.5)/*n*, with *y* being the target variable and *n* the sample size (Smithson and Verkuilen, 2006). *sMP_diff_* and *sIP_diff_* take values between −1 and 1, and were therefore transformed by applying (*y* + 1)/2 (Cribari-Neto and Zeileis, 2010). The full models were built according to Eq. (1), the reduced models according to Eq. (2), and included an interaction term between the return period (*RP*) and the intervention intensity (*Int*) as well as the squares of both these predictors. A logit-link function was applied.

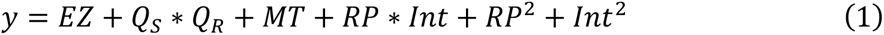

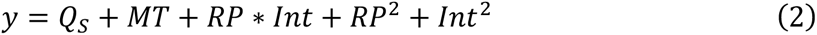

with *y* being the target variable, *EZ* the elevational zone, *Q_S_* the site quality, *Q_R_* the regeneration quality, and *MT* the management type. The models were calculated with the R library *betareg* (Cribari-Neto and Zeileis, 2010). We then calculated and visualized the marginal effects of the predictor variables using the R library *ggeffects* (Lüdecke, 2018).

## 3. Results

First, we will present the results of the full BRT and beta regression models, describing the importance and effect of the variables explaining the protective quality of forest stands across all *elevational zones*, *site* and *regeneration qualities*, and management regimes. The second part of this section contains the results of the reduced models where only simulations with good *regeneration quality* were considered, separately for each elevational zone and initialization stand type, followed by a summary of the management recommendations. Finally, we present the species-specific growth behavior of ProForM that helps to explain certain model outputs.

### 3.1. Importance and effects of site and management in full models

According to the BRT, by far the most important variables for explaining the absolute protective quality of stands in short-term simulations (ST) were *elevational zone* and *regeneration quality*, with mean variable importance values across natural hazards and the type of NaiS-profiles of 60 and 33%, respectively (Fig. 1a). The most important variables for explaining the impact of management on the protective quality were *elevational zone* (mean importance of 29% across natural hazards and NaiS profiles), *regeneration quality* and *initialization stand type* (both 22% importance, Fig. 1b). The importance of the management-related variables *type, interval*, and *intensity* as well as *site quality* was low for both types of response variables, with maximum mean importance values of 2% for the absolute protective quality and 9% for the impact of management.

**Fig. 1.**
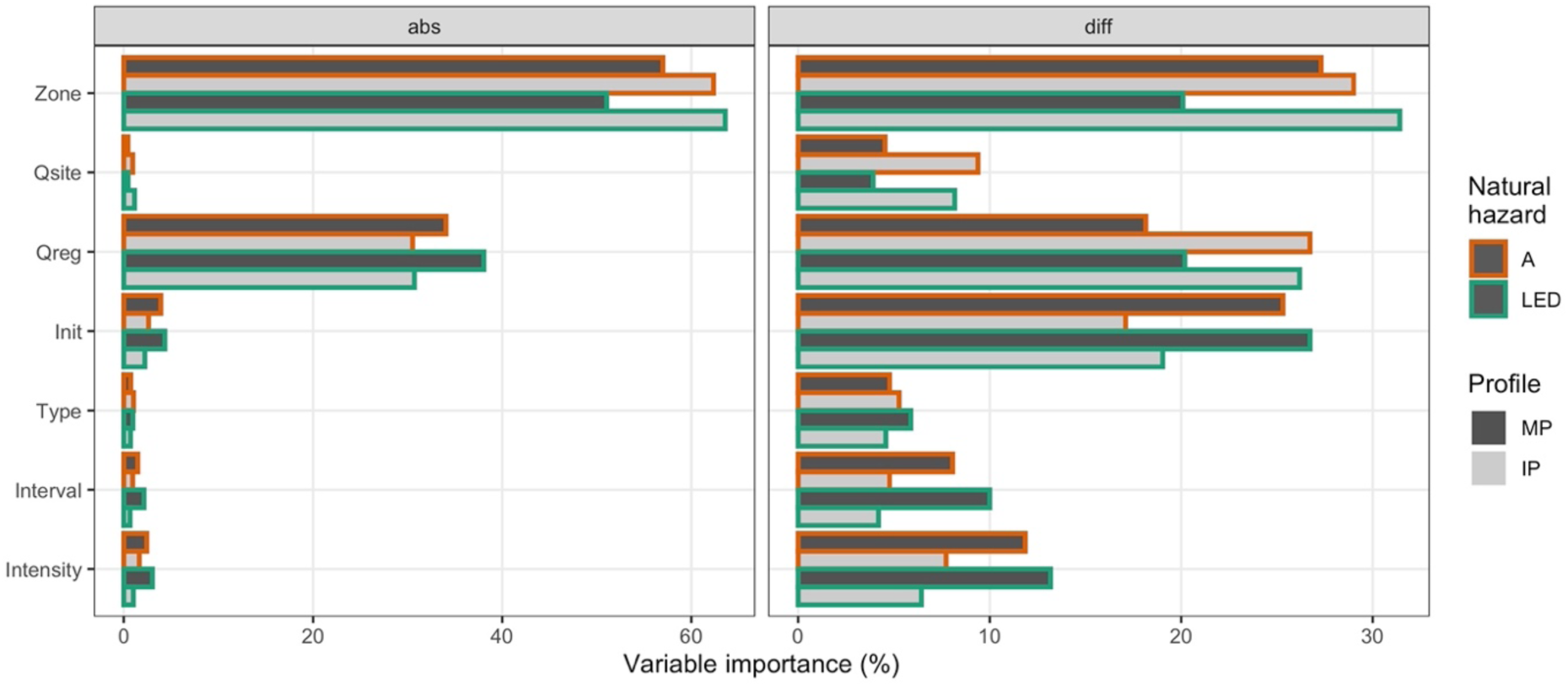
Variable importance in in BRT of short-term simulations explaining the absolute protective quality (“abs”, left panel) and the effect of management on the protective quality (difference in protective quality compared to NOM; “diff”, right panel) using all explanatory variables (full models). Bar colors differentiate minimal profile (MP) and ideal profile (IP), outline colors the natural hazards snow avalanches (A) and landslide/erosion/debris flow (LED). Explanatory variables are ordered by mean variable importance. Abbreviations of explanatory variables: Zone = elevational zone; Qreg = regeneration quality; Init = initialization stand type; Qsite = site quality; Type = management type.

Similarly, in long-term simulations (LT) *elevational zone* and *regeneration quality* were the most important variables in the BRT models explaining both the absolute protective quality as well as the impact of management (Fig. 2). The mean variable importance across natural hazards and NaiS profiles of *elevational zone* was 57% for the absolute protective quality and 33% for the impact of management. The importance of *regeneration quality* was 30% and 42%, respectively. Again, the management-related variables as well as *site quality* were of minor importance. The cross-validation metrics of all BRT models using the full data set (“full models”) are given in Supplementary material S7, Table S7.1.

**Fig. 2.**
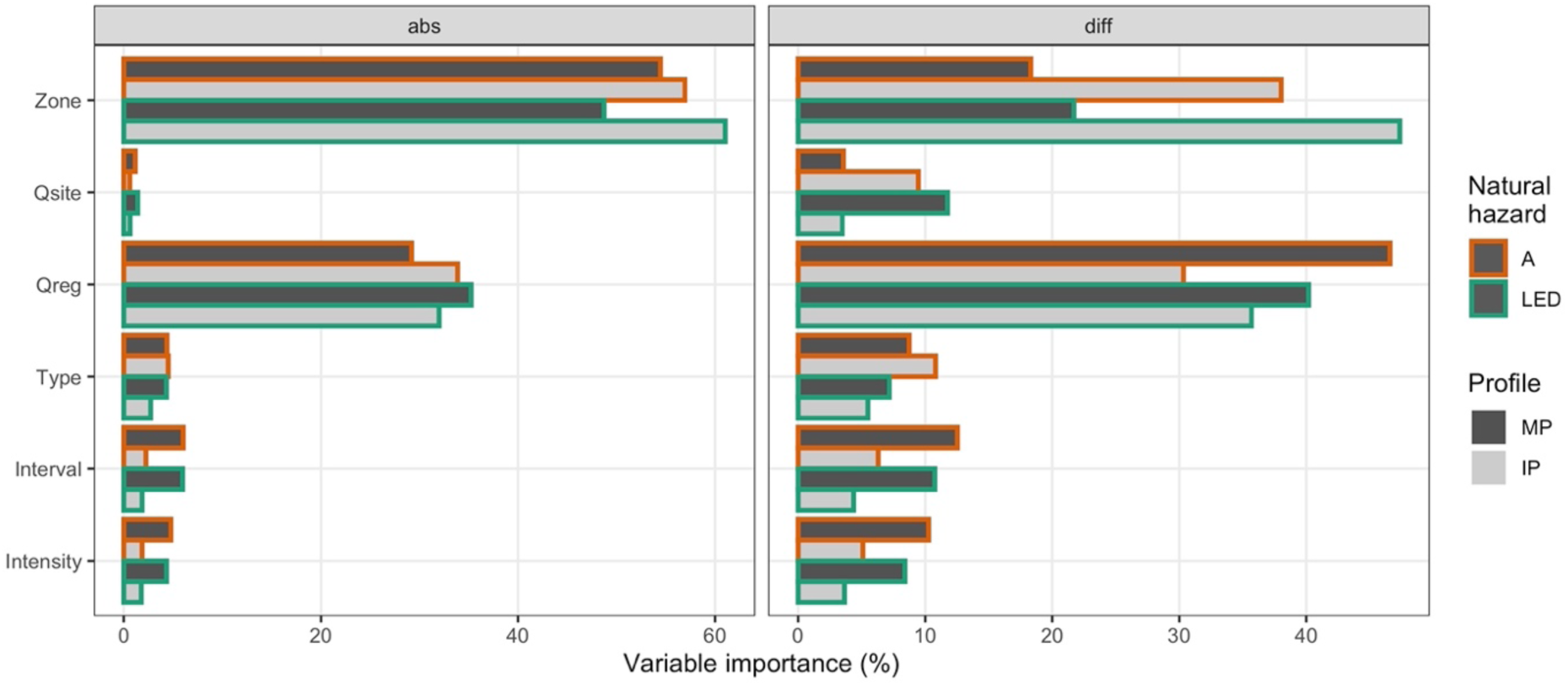
Variable importance in BRT on long-term simulation results explaining the absolute protective quality (“abs”, left panel) and the effect of management on the protective quality (difference in protective quality compared to NOM; “diff”, right panel) using all explanatory variables (full models). Bar colors differentiate minimal profile (MP) and ideal profile (IP), outline colors the natural hazards snow avalanches (A) and landslide/erosion/debris flow (LED). Explanatory variables are ordered by mean variable importance. Abbreviations of explanatory variables: Zone = elevational zone; Qreg = regeneration quality; Qsite = site quality; Type = management type.

The most important variables influencing the protective quality of a stand according to the BRT models can thus not be altered by forest management on a stand level, as they are either given by the stand’s location (*elevational zone*), the regional ungulate densities (*regeneration quality*), or the management history (*initialization stand type*).

Marginal effects of the beta regression models of the full dataset (model statistics in Table S7.2) revealed a clear negative effect of low *regeneration quality* (Fig. 3, S7.1, and S7.3). Furthermore, analyses of the assessment results of pairs of simulations that differed only in *regeneration quality* showed that the main reason was the lower number of tree species present, which resulted in lower index values for “species mixture”, “seedlings”, and “saplings and thicket”, as well as a lower canopy cover and therefore lower index values for “horizontal arrangement”.

**Fig. 3.**
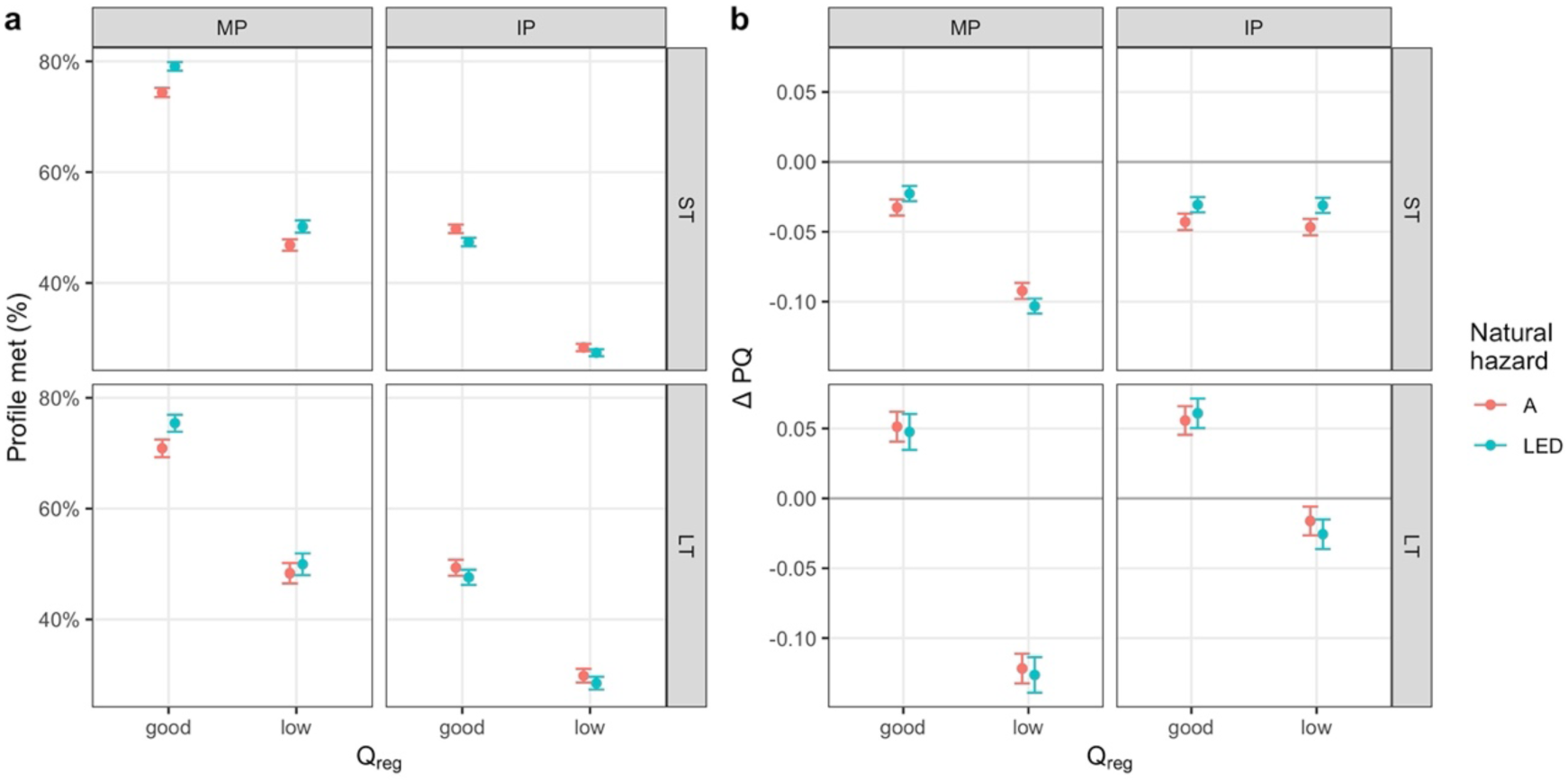
Marginal effects (mean and 95% confidence interval) of **regeneration quality** in beta regression models explaining the absolute protective quality (a) and effect of management on the protective quality (difference in protective quality compared to NOM, ΔPQ; b) using all explanatory variables (full models). Panel columns separate the NaiS-profile (MP = minimal profile; IP = ideal profile), panel rows the simulation type (ST = short-term; LT = long-term simulations), and color the natural hazards snow avalanches (A) and landslide/erosion/debris flow (LED).

The mean marginal effects of management (*type*, *intensity*, and *interval*; Figs. S7.2 and S7.4) on the protective quality were almost exclusively negative. A further differentiation of these marginal effects by fixing *regeneration quality* at either of its two possible values (“normal” or “low”) revealed that management interventions – regardless of *type*, *interval*, or *intensity* – had a negative effect on the protective quality when regeneration was strongly hindered in both ST and LT simulations and for both the minimal (Fig. 4) and the ideal profile (Fig. 5). The only exception were two slightly positive mean marginal effects on the impact of management in LT simulations when the ideal profile was considered (Fig. 5). Yet, management effects in stands with a normal *regeneration quality* were always positive in LT simulations and considerably less negative in ST simulations. The main reasons were – again inferred from comparisons of individual simulation results – that while species composition (with effects on the indices “species mixture”, “seedlings”, and “saplings and thicket”) could not be improved through management, the already often critical values of canopy cover were decreased and gap sizes increased by the interventions, leading to an overall negative effect of management.

**Fig. 4.**
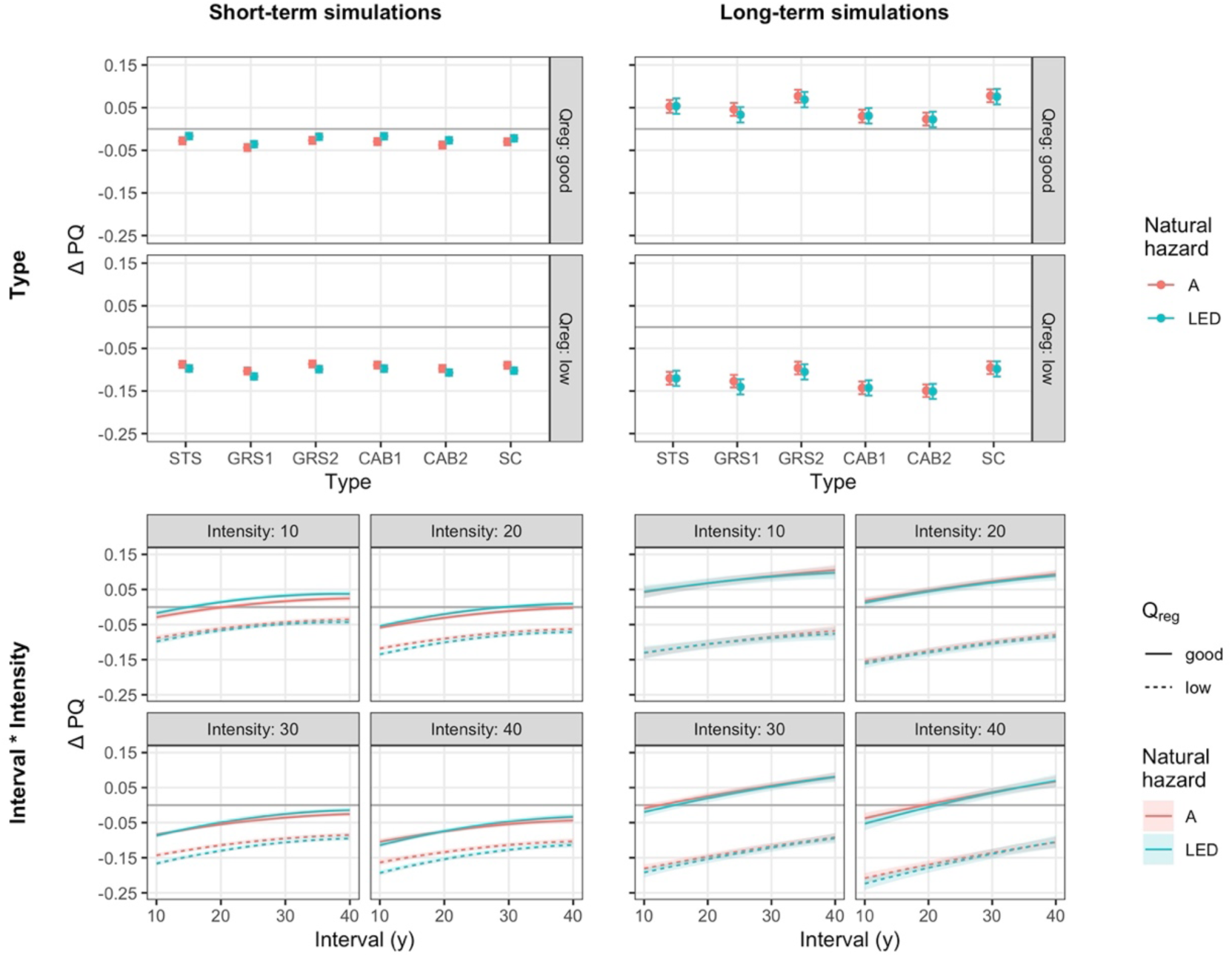
Marginal effects (mean and 95% confidence interval) of **management type** (upper row) and the interaction of **interval and intensity** (lower row) in beta regression models explaining the effect of management on the protective quality (difference in protective quality compared to NOM, ΔPQ) assessed by the **minimal profile** using all explanatory variables (full models). The natural hazards snow avalanches (A) and landslide/erosion/debris flow (LED) are differentiated by color, the two levels of regeneration quality (Q_reg_) by panels in the upper row and by line type in the lower row.

**Fig. 5.**
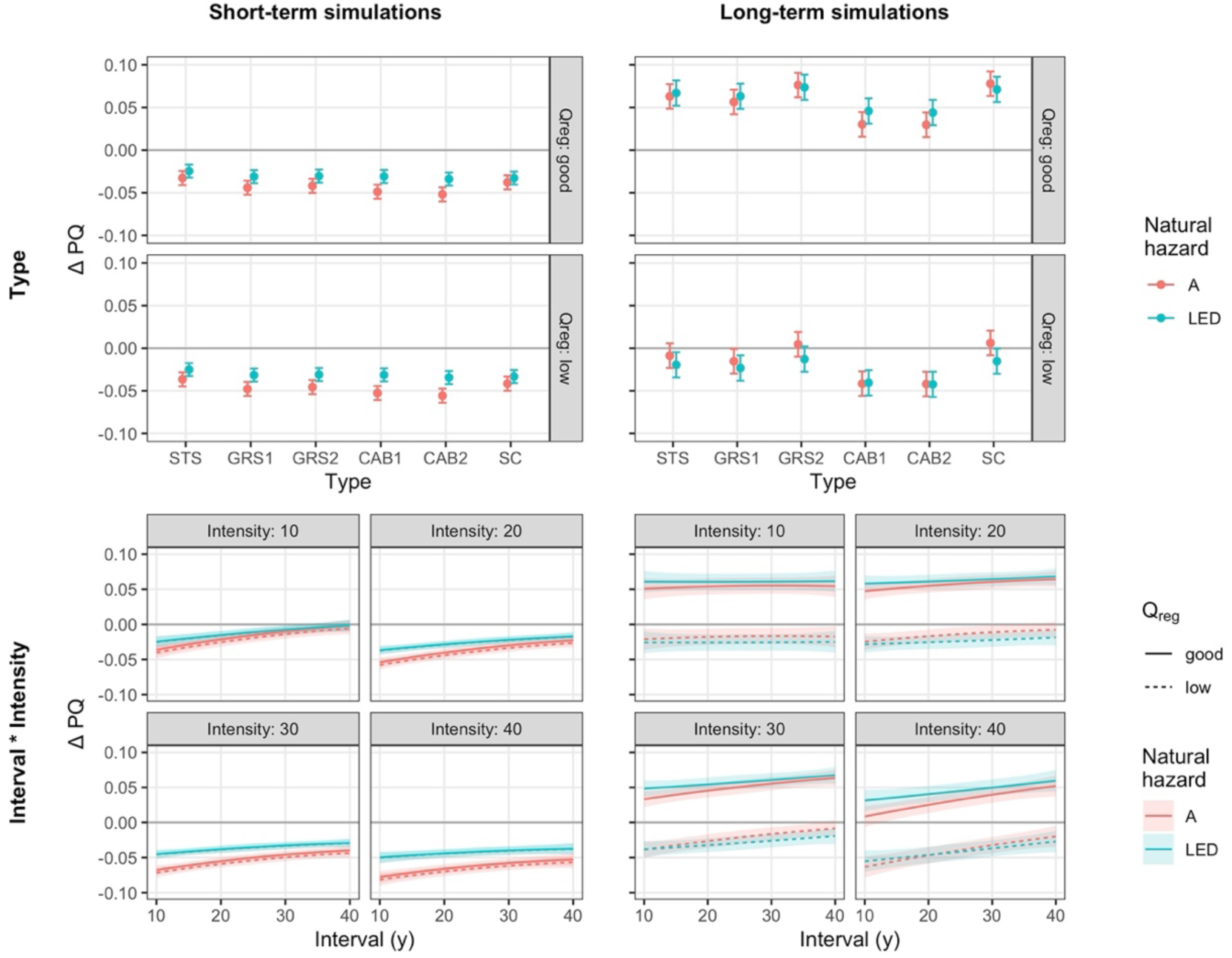
Marginal effects (mean and 95% confidence interval) of **management type** (upper row) and the interaction of **interval and intensity** (lower row) in beta regression models explaining the effect of management on the protective quality (difference in protective quality compared to NOM, ΔPQ) assessed by the **ideal profile** using all explanatory variables (full models). The natural hazards snow avalanches (A) and landslide/erosion/debris flow (LED) are differentiated by color, the two levels of regeneration quality (Q_reg_) by panels in the upper row and by line type in the lower row.

### 3.2. Effect of management on protective quality in reduced models

The reduced models are based only on simulations with a good *regeneration quality,* i.e., a low browsing pressure. To be able to better disentangle the role of management for the protective quality, we built separate models for each *elevational zone* and *initialization stand type*, thus removing two important explanatory variables from the models that cannot be influenced by forest managers. The cross-validation metrics of these BRT models are provided in Table S7.3, the statistics of the beta regression models in Table S7.4.

As our main goal was to identify suitable management regimes for enhancing or at least maintaining the protective quality of stands, below we focus on the analysis of the impact of management on the protective quality, i.e., the difference to the corresponding simulations without management (NOM). The change in protective quality (ΔPQ) depicted by *sMP_diff_* and *sIP_diff_* quantifies how much smaller or larger the share of yearly index values is that fulfill the respective NaiS-profile, compared to the NOM simulation. If an index, e.g., “species mixture”, is constantly below the minimal profile in NOM but at the minimal profile in the simulation with management, this causes a *sMP_diff_* of +0.17 (i.e., 1/6), as there are six indices in total. The basis for interpreting these relative values are the protective qualities achieved in NOM simulations, which are presented at the beginning of each section. Further insights into the underlying reasons for the effect of certain management regimes on the protective quality were drawn from individual model outputs along with their protective qualities (cf. Supplementary material S5) and aggregated visualizations of the protective qualities per model (cf. Supplementary material S6).

#### 3.2.1. Upper montane zone

In the upper montane zone, NOM led to a relatively poor protective quality, with *sMP_abs_* of roughly 60% and *sIP_abs_* below 40% across all simulation types and initialization stand types, NaiS-profiles, *site qualities*, and natural hazards (Fig. 6). Across all simulations, the “species mixture” rarely fulfilled the minimal profile, with shares of *Abies alba* being above and those of *Fagus sylvatica* below their respective threshold values according to the NaiS profiles. This translated into the index for “seedlings” often not reaching the minimal profile either, as *Abies alba* seedlings dominated regeneration in the few openings. The indices for “vertical structure” and “horizontal arrangement” were at or close to the ideal profile. Apart from the young stands in ST simulations, the “stability of support trees” (measured by their average crown length) did not usually fulfill the requirements for the minimal profile, as these unmanaged stands were relatively dense.

**Fig. 6.**
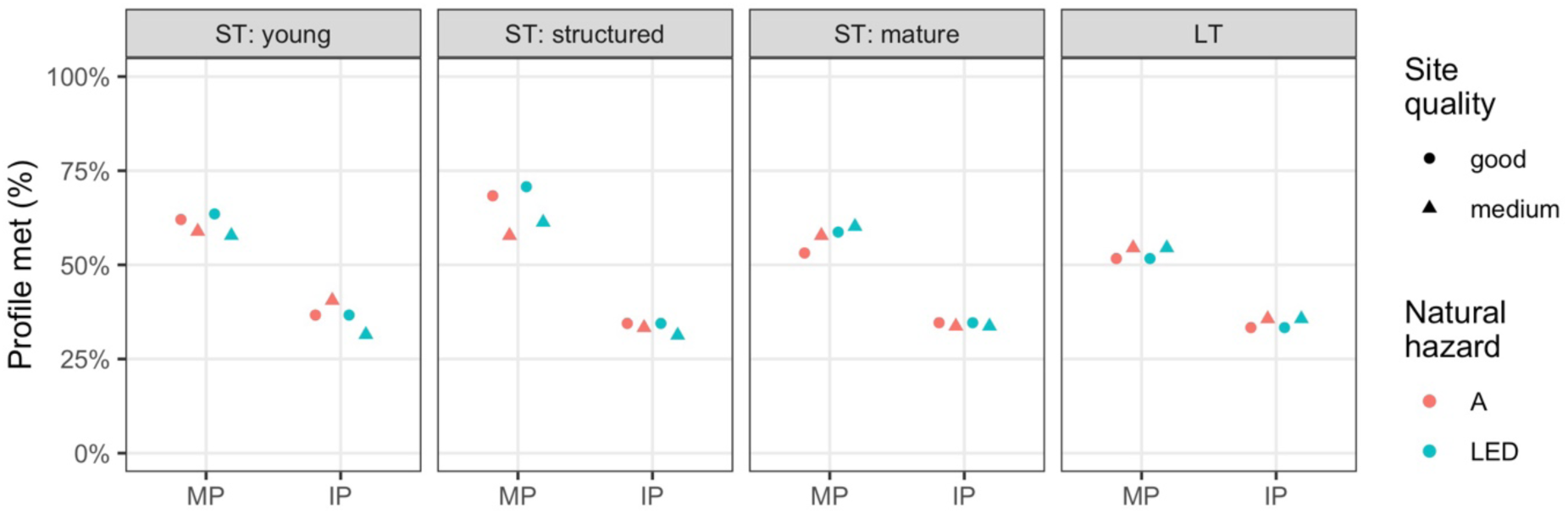
Protective quality of simulations without management (NOM) of elevational zone UM by initialization stand type (ST = short-term; LT = long-term; panels), and NaiS-profile (MP = minimal profile, sMP_abs_; IP = ideal profile, sIP_abs_). Symbols differentiate the site quality, colors the natural hazards snow avalanches (A) and landslide/erosion/debris flow (LED).

According to the BRT, the three management-related variables *management type, interval,* and *intensity* had a similar mean importance for explaining the impact of management on the protective quality across all UM models, while *site quality* was of lower importance (Fig. S7.7). The beta regression models revealed that the effect of forest management on the protective quality strongly depended on the stand type used for the initialization and the simulation length (Figs. 7-10). While the degree of fulfillment of the minimal profile was increased by management in some stand types, the already low fractions of the ideal profile being fulfilled in NOM simulations were further reduced by almost all management regimes, for the following reasons. (1) The species-related indices (“species mixture” and the two regeneration-related indices) could not – if at all – be improved to reach the ideal profile and did therefore not impact *sMP_diff_.* (2) The indices for “vertical structure” and “horizontal arrangement” were already at or close to the ideal profile in NOM simulations and could therefore hardly be improved by management, but only maintained or reduced (e.g., by opening gaps that exceeded the area thresholds). (3) The “stability of support trees” was improved by management, yet at the cost of reducing other indices (i.e., “vertical structure” and “horizontal arrangement”) to a higher degree, and thus leading to an overall deterioration of *sIP_abs_*. However, as even the best-performing management regimes were unable to reach 100% fulfillment of the minimal profile, the effect of management with regard to the ideal profile was of minor importance in UM. The UM beta regression models had a mean pseudo-R^2^ of 0.47 (mean of 0.53 for the minimal-profile-models and 0.42 for the ideal-profile-models, Table S7.4).

**Fig. 7.**
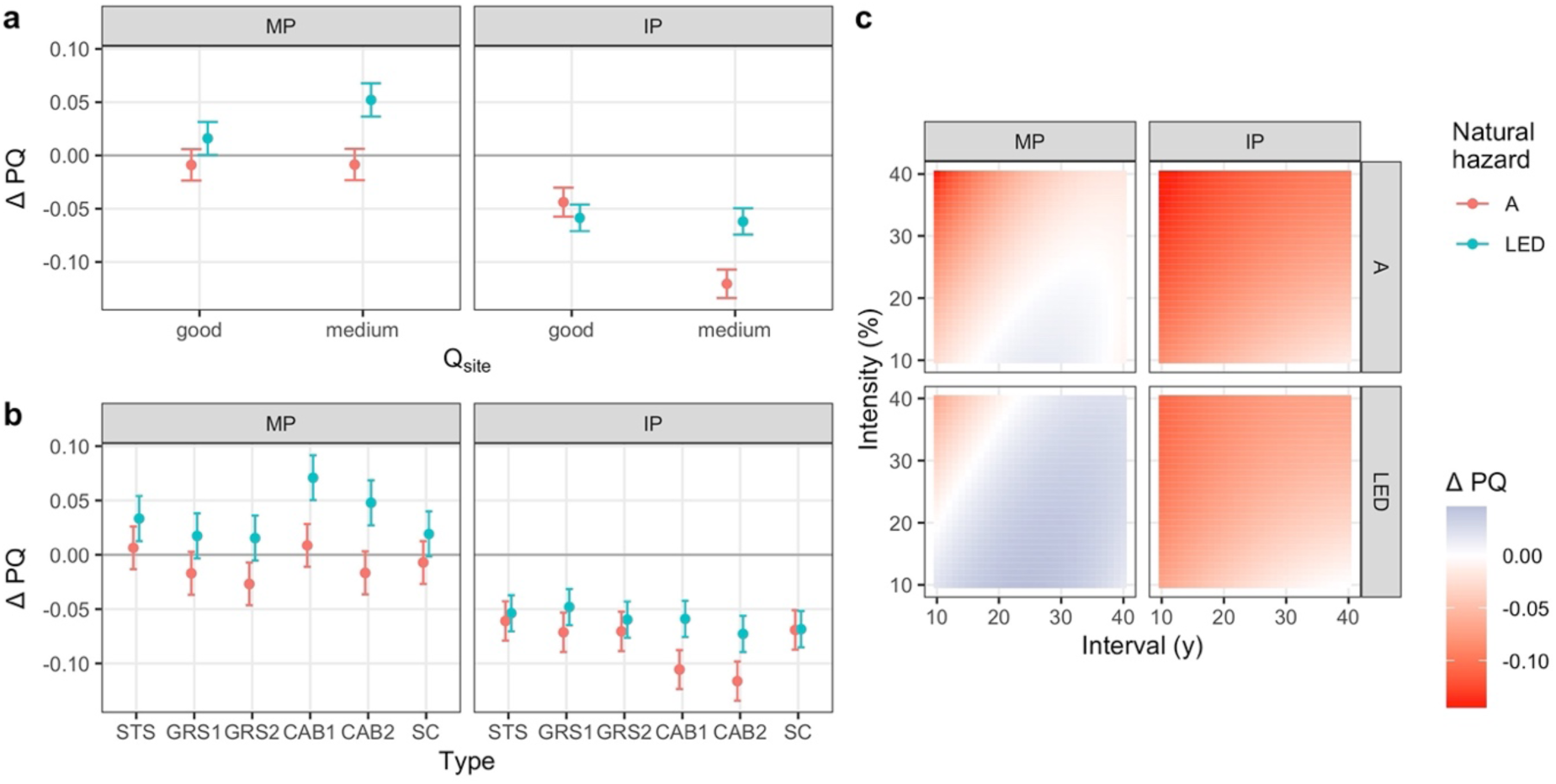
Marginal effects in reduced beta regression models explaining the effect of management on the protective quality (difference in protective quality compared to NOM, ΔPQ) in elevational zone **UM**, **short-term** simulations initialized with a **young stand**. a: mean effects and 95% confidence interval of site quality; b: mean effects and 95% confidence interval of management type; c: mean effects of interval and intensity. Separated by NaiS-profiles (MP = minimal profile; IP = ideal profile) and natural hazard (A = snow avalanches; LED = landslide/erosion/debris flow).

**Fig. 8.**
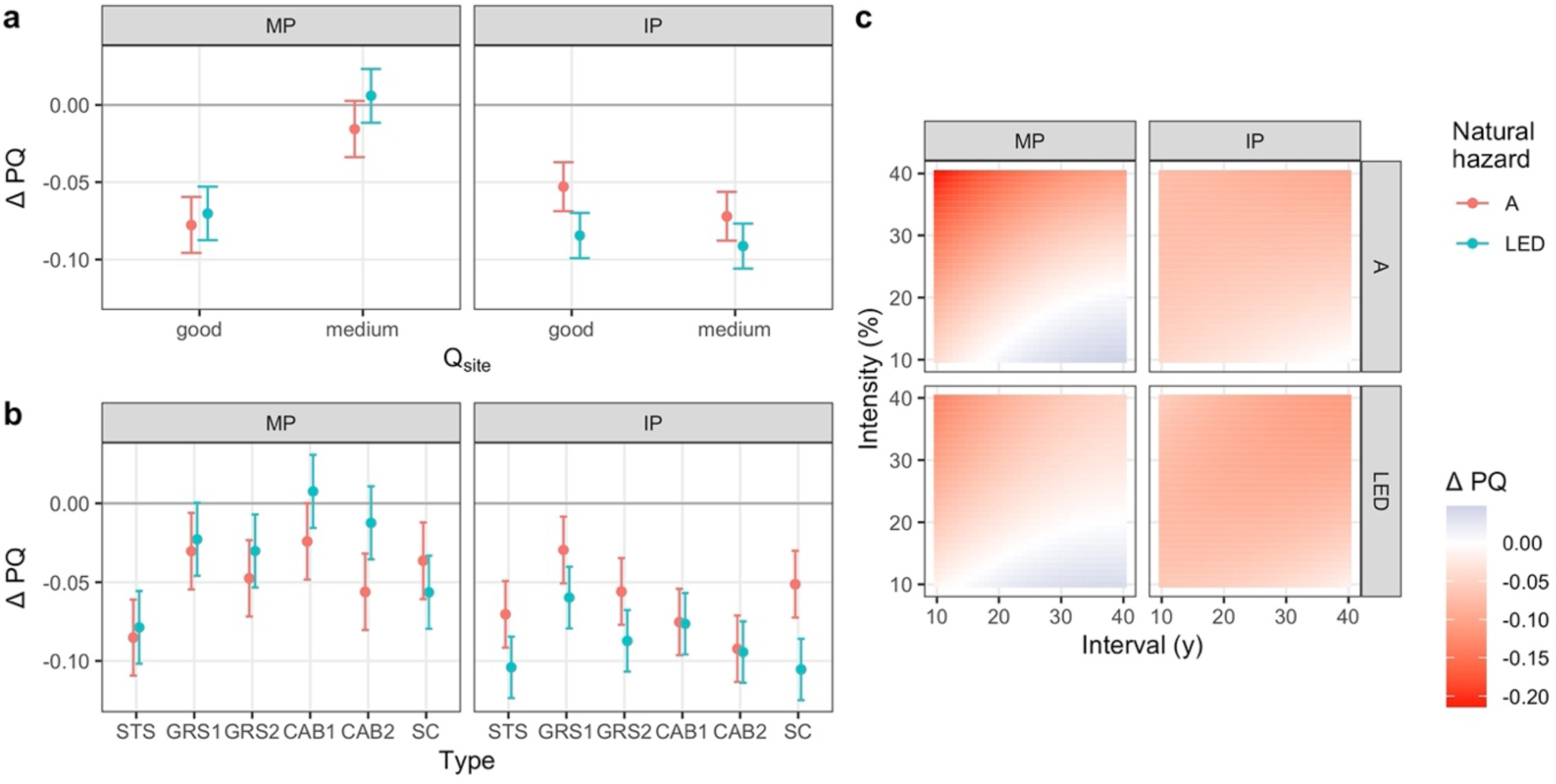
Marginal effects in reduced beta regression models explaining the effect of management on the protective quality (difference in protective quality compared to NOM, ΔPQ) in elevational zone **UM**, **short-term** simulations initialized with a **well-structured stand**. a: mean effects and 95% confidence interval of site quality; b: mean effects and 95% confidence interval of management type; c: mean effects of interval and intensity. Separated by NaiS-profiles (MP = minimal profile; IP = ideal profile) and natural hazard (A = snow avalanches; LED = landslide/erosion/debris flow).

**Fig. 9.**
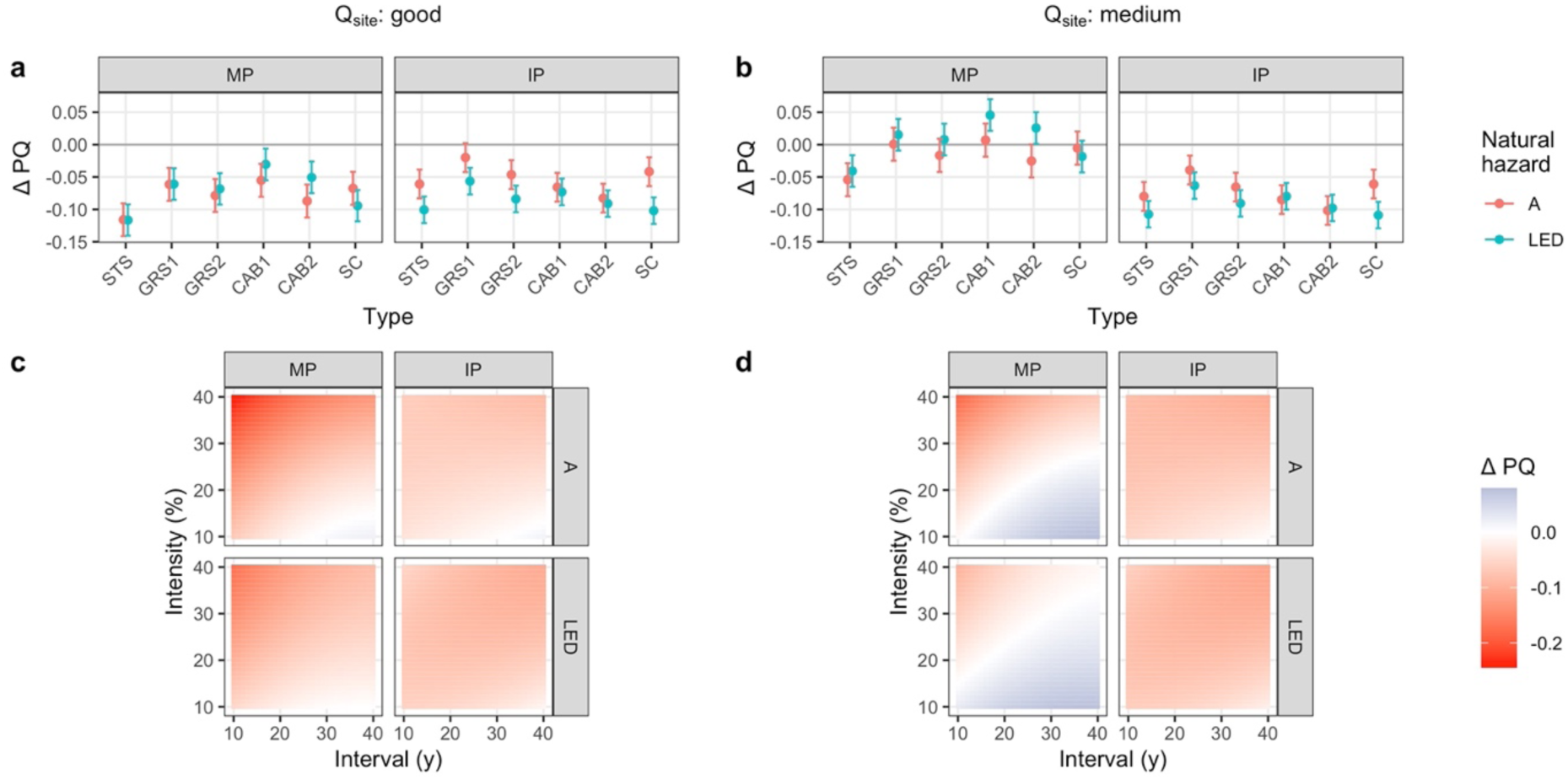
Marginal effects in reduced beta regression models explaining the effect of management on the protective quality (difference in protective quality compared to NOM, ΔPQ) in elevational zone **UM**, **short-term** simulations initialized with a **well-structured stand** with **site quality held fixed** at “good” (left column) and “medium” (right column). a & b: mean effects and 95% confidence interval of management type; c & d: mean effects of interval and intensity. Separated by NaiS-profiles (MP = minimal profile; IP = ideal profile) and natural hazard (A = snow avalanches; LED = landslide/erosion/debris flow).

**Fig. 10.**
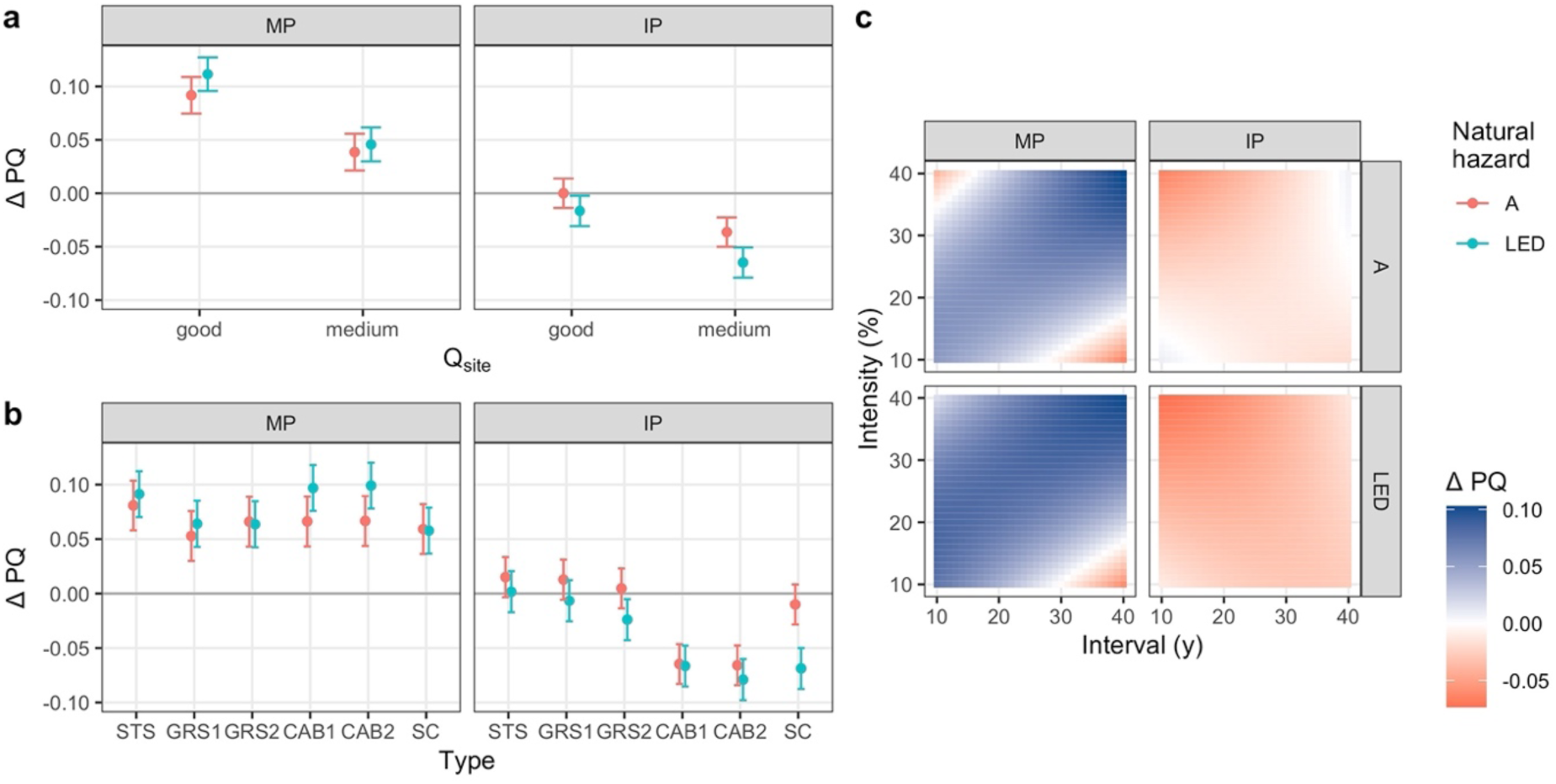
Marginal effects in reduced beta regression models explaining the effect of management on the protective quality (difference in protective quality compared to NOM, ΔPQ) in elevational zone **UM**, **short-term** simulations initialized with a **mature stand**. a: mean effects and 95% confidence interval of site quality; b: mean effects and 95% confidence interval of management type; c: mean effects of interval and intensity. Separated by NaiS-profiles (MP = minimal profile; IP = ideal profile) and natural hazard (A = snow avalanches; LED = landslide/erosion/debris flow).

Protective quality was improved more (or reduced less) in stands protecting against landslides/erosion/debris flow (LED) than against avalanches (A), especially regarding the minimal profile. The main reason for this was the considerably lower threshold values for allowable gap sizes in the profile for avalanche protection forests. However, as snow avalanches are of low importance at these elevations (800-1400 m a.s.l), we will focus on LED below.

##### 3.2.1.1. Young stands

Unmanaged young stands in UM almost never fulfilled the minimal profiles for the species-related indices. “Species mixture” was constantly below the minimal profile due to an overabundance of *Abies alba*, and the index for “seedlings” – if there were any gaps – most of the time as well. “Saplings and thicket” fulfilled the minimal profile for a few decades on medium *site quality*, but this index was rated below the minimal profile most of the time due to a lack of *Abies alba* in the regeneration. The paradox situation of having too much *Abies alba* in the mature stand while having too little in the regeneration arose from the removal of conifers during the spin-up simulations aimed at creating an initialization stand not completely dominated by coniferous species. The indices for “vertical structure” and “horizontal arrangement” improved throughout the simulations from below the minimal towards the ideal profile, while “stability of support trees” was mostly ideal over time.

On average, all management types had a positive effect on the fulfillment of the minimal profile in simulations of young stands protecting against LED, but a negative effect on the fulfillment of the ideal profile (Fig. 7). The fulfillment of the minimal profile was increased primarily through an improved species mixture among the regeneration, leading to close-to-ideal ratings of the index for “saplings and thicket”, while the “species mixture” of adult trees remained dominated by *Abies alba* beyond the threshold values of the minimal profile. The relatively large gaps formed through CAB allowed for a more balanced species composition among seedlings, thereby increasing the corresponding index value above the minimal profile more often than in the case of smaller openings created by single-tree (STS) or group selection (GRS). Most combinations of *interval* and *intensity* had a mean positive effect of similar magnitude, only frequent (10-20 years interval) and strong interventions (30-40% basal area removal) led to a decrease in the protective quality. In the latter cases, management reduced the basal area and canopy cover so strongly that the “vertical structure” and “horizontal arrangement” rarely met the requirements of the minimal profile. The negative effect of management on fulfilling the ideal profile resulted mostly from a slightly lower “vertical structure” and “horizontal arrangement”, while the other indices did not reach the ideal profile. Only interventions at low intensity and long intervals led to almost no decrease in *sIP_abs_*.

Based on these results, the most promising management regime for young stands would be to intervene only two or three times over the course of the simulated 90 years, using a low intensity and concentrating the logging on a few gaps. This would increase the fulfillment of the minimal profile through a more balanced species composition in the regeneration while not compromising the naturally favorable development of the horizontal and vertical stand structure.

##### 3.2.1.2. Well-structured stands

Well-structured stands that were left unmanaged had a relatively good “vertical structure”, i.e., diameter distribution (lying between the minimal and the ideal profile) and an often ideal “horizontal arrangement” due to the absence of large gaps. The “stability of support trees” varied over time and often did not fulfil the minimal profile, while the species composition of seedlings in the few gaps was usually not diverse enough to fulfill the minimal profile. The indices for “species mixture” and “saplings and thicket” depended strongly on the *site quality* of the simulated stands. On good *site quality*, the “species mixture” was close to ideal but there was not enough *Abies alba* among the regeneration to fulfill the minimal profile. On medium *site quality*, the opposite was found: “species mixture” was below the minimal profile due to too little *Fagus sylvatica*, but the index for “saplings and thicket” was close to ideal. The differences in species composition arose mainly from the different initialization stands (cf. Figs. S3.5 and S3.7). Overall, stands on good *site quality* had a higher protective effect without management than those on medium *site quality*.

The differences in the stand attributes between *site qualities* led to different effects of management on the protective quality, with improvements in the fulfillment of the minimal profile only possible in stands with medium *site quality* (Fig. 8a).

On good *site quality* (Fig. 9a/c), all management regimes led to either no change or a mean decrease in the protective quality. The main reasons were that (1) interventions skewed the diameter distribution too much towards smaller tree sizes, (2) gaps exceeding the threshold size were created, and (3) canopy cover dropped below the threshold of the ideal profile with stronger interventions. On medium *site quality*, however, the degree of fulfillment of the minimal profile was increased by all management types except for STS and SC (slit cuts), with CAB1 performing best (Fig. 9b), and interventions with low *intensity* being generally more favorable (Fig. 9d). Compared to stands with good *site quality*, the “stability of support trees” and the species distribution among seedlings was improved more. Still, these interventions generally lowered the degree to which the ideal profile was fulfilled. While the “stability of support trees” and the species mixture among seedlings was improved so as to fulfill the minimal profile, the indices for “vertical structure” and “horizontal arrangement” were reduced from the ideal to the minimal profile.

Even though the possibilities of improving the fulfillment of the minimal profile were considerably larger in stands with medium *site quality*, the recommended management regimes for well-structured stands would be similar for both *site qualities*: Either not intervening at all or carrying out interventions with a low overall *intensity* at long *intervals*, with the removals concentrated on few gaps. This would lead to no or only a small decrease in the protective quality on good sites while improving the protective quality on medium sites, as potentially positive effects on species composition of regeneration and on stand stability are balanced with negative effects on horizontal stand structure by avoiding large canopy openings.

##### 3.2.1.3. Mature stands

Mature stands that were left unmanaged for 90 years had an ideal “vertical structure” and “horizontal arrangement”, i.e., a well-balanced diameter distribution and few to no gaps. The index for “saplings and thicket” was rated slightly below the ideal profile due to the species composition, while the share of regeneration was well above the ideal threshold throughout the entire simulation. The “species mixture” of adult trees did not fulfill the minimal profile due to too little *Fagus sylvatica* on good *site quality* and an overabundance of *Abies alba* on medium sites, and species diversity within the few gaps mostly did not fulfill the minimal profile, regardless of *site quality*. The largest trees had relatively short crowns, and therefore the “stability of support trees” was rated below the minimal profile.

Forest management increased the degree of fulfillment of the minimal profile across *site types* and irrespective of the management type, with the exception of the extreme cases of very frequent strong or infrequent fine interventions (Fig. 10). The positive effect of management was more pronounced in stands with a good *site quality*. There, the degree of fulfillment of the ideal profile could also be increased somewhat by dispersed, small openings (management types STS and GRS; Fig. 11a). On medium *site quality*, the fulfillment of the ideal profile decreased with virtually all management regimes (Fig. 11b). The main difference between *site qualities* was that gaps opened through interventions were closed faster due to higher regeneration growth rates on good sites. Overall, management interventions generally improved the crown length of support trees and often the number of species among seedlings. However, the “species mixture” of adult trees was only improved sufficiently to reach the minimal profile when intense interventions were made.

**Fig. 11.**
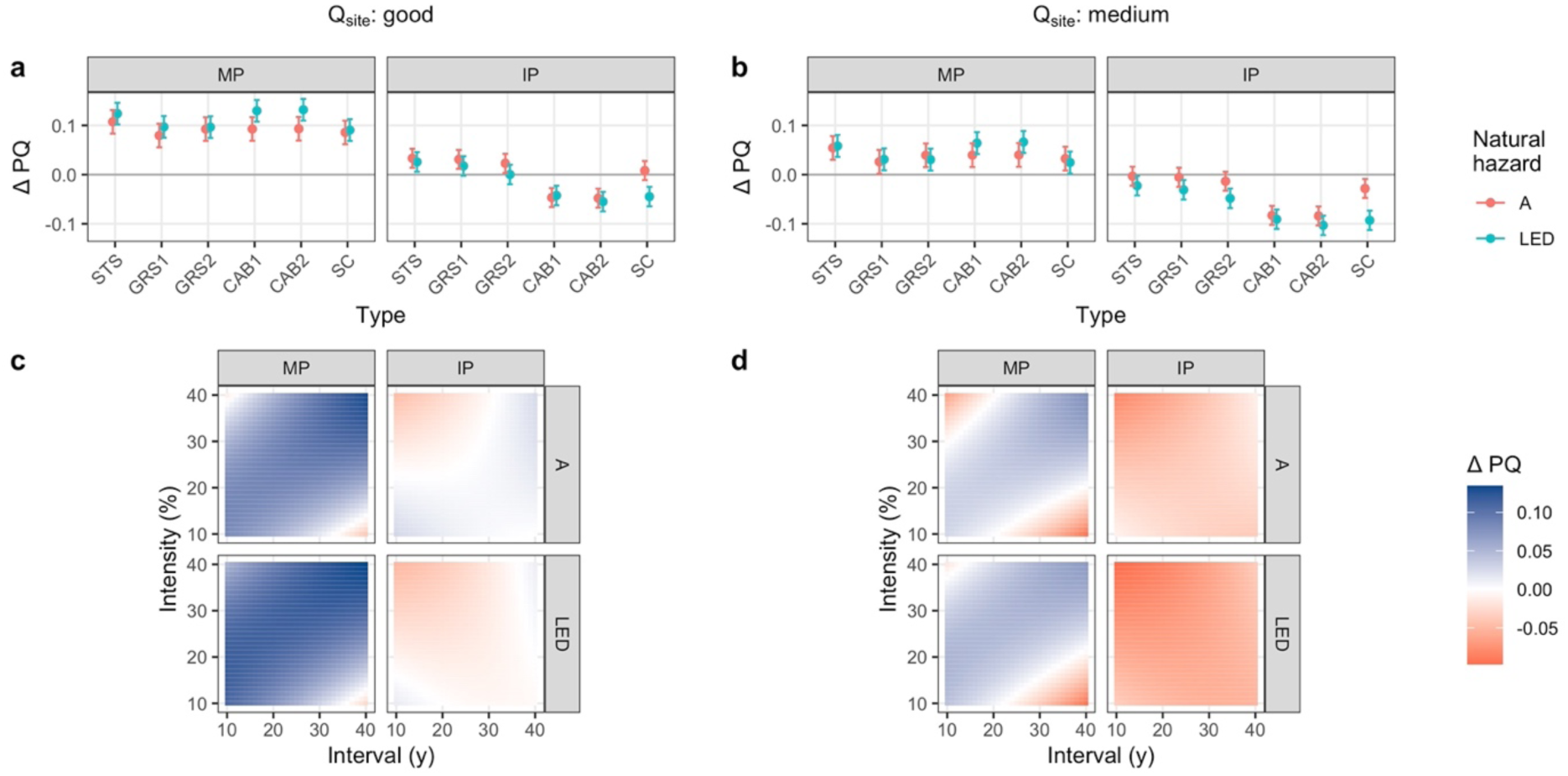
Marginal effects in reduced beta regression models explaining the effect of management on the protective quality (difference in protective quality compared to NOM, ΔPQ) in elevational zone **UM**, **short-term** simulations initialized with a **mature stand** with **site quality held fixed** at “good” (left column) and “medium” (right column). a & b: mean effects and 95% confidence interval of management type; c & d: mean effects of interval and intensity. Separated by NaiS-profiles (MP = minimal profile; IP = ideal profile) and natural hazard (A = snow avalanches; LED = landslide/erosion/debris flow).

The management regime to be recommended for mature stands based on these results is mainly determined by avoiding negative effects on the fulfillment of the ideal profile, i.e., not reducing the quality of the diameter distribution and not creating gaps that exceed the threshold size for too long. As such gaps close more quickly on good sites, the available management options are broader than on medium sites. However, the fulfillment of the ideal profile is reduced least on both *site qualities* with low-intensity interventions and when creating only small openings (STS or GRS). Such management regimes can also increase the fulfillment of the ideal profile, mainly through increased stability.

##### 3.2.1.4. Long-term simulations

In LT simulations in the elevational zone UM, forest management improved the fulfillment of the minimal profile in a stronger fashion than in the other simulation types (Fig. 12). All *management types* and most combinations of *intensity* and *interval* led to an improvement. The main deficits of NOM simulations were the “species mixture” (almost complete dominance by *Abies alba*), the short crowns of support trees and the few species present among seedlings. A more balanced species composition of adult trees could be reached with either frequent and/or strong interventions, while the species composition among seedlings improved under most management regimes. The “stability of support trees” was greatly improved by management, regardless of the regime applied. However, most simulated management regimes led to a decrease in the fulfillment of the ideal profile. Negative effects – even if only temporary – on the “vertical structure” and “horizontal arrangement” that were ideal over time in NOM could not be compensated by other indices such as “species mixture” or the “stability of support trees” reaching the ideal profile at times. The least negative effects on the fulfillment of the ideal profile were achieved by management regimes creating relatively small openings (STS or GRS) with low overall intensities.

**Fig. 12.**
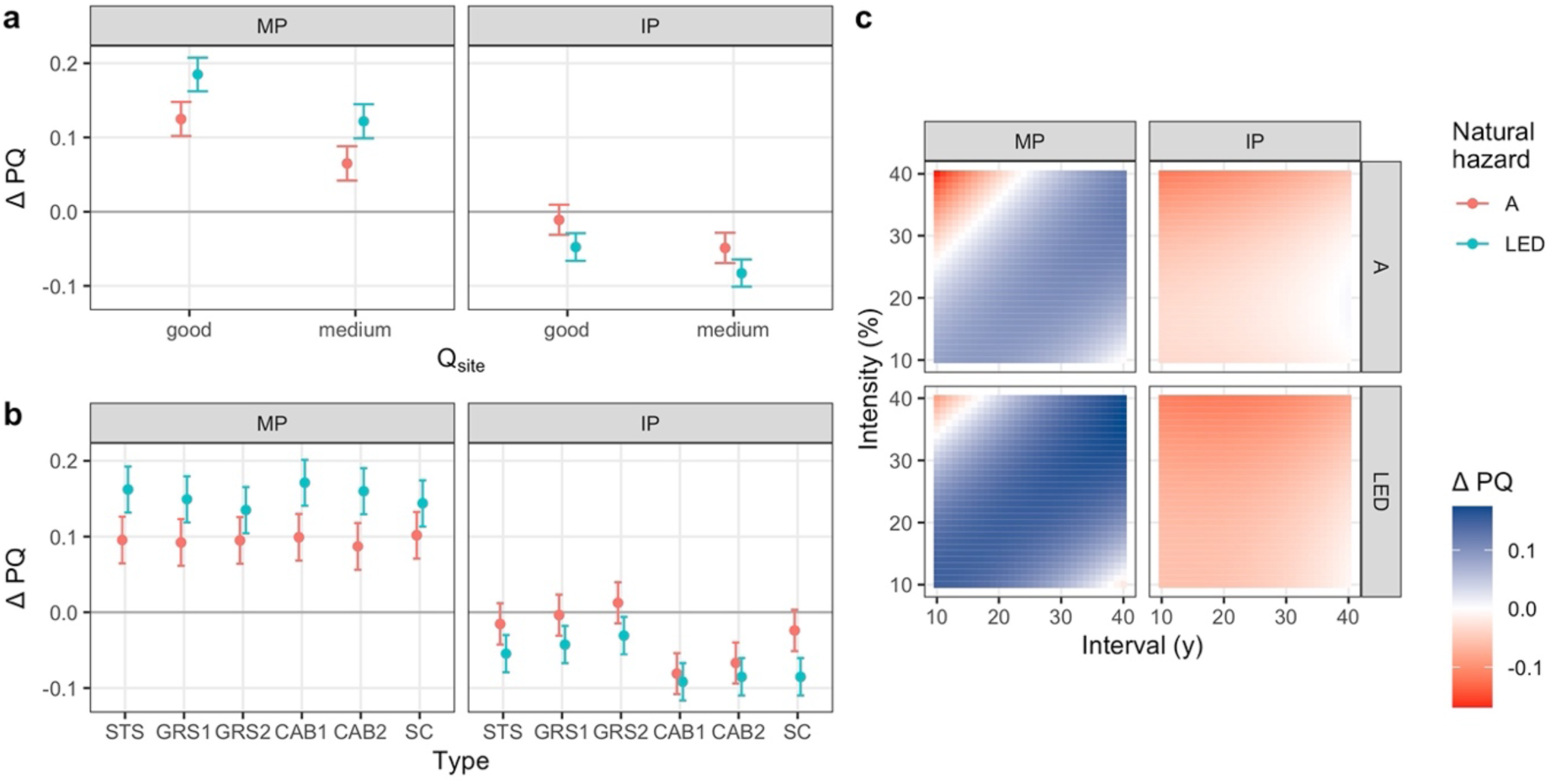
Marginal effects in reduced beta regression models explaining the effect of management on the protective quality (difference in protective quality compared to NOM, ΔPQ) in elevational zone **UM**, **long-term** simulations. a: mean effects and 95% confidence interval of site quality; b: mean effects and 95% confidence interval of management type; c: mean effects of interval and intensity. Separated by NaiS-profiles (MP = minimal profile; IP = ideal profile) and natural hazard (A = snow avalanches; LED = landslide/erosion/debris flow).

Therefore, the recommended management regime in the long-term consists of small openings at a low overall intensity, as this hampers the fulfillment of the ideal profile least while greatly improving the fulfillment of the minimal profile, mainly due to a more balanced species composition and longer crowns of the support trees.

#### 3.2.2. High montane zone

In the high montane zone, the protective quality of all NOM simulations ranged between ca. 70-100% *sMP_abs_* (higher in ST than in LT simulations) and 40-80% *sIP_abs_* (^F^ig. 13). In ST simulations, the minimal profiles were nearly completely fulfilled across time, with the only exception being the “stability of support trees”, i.e., mean crown length. LT simulations achieved a lower protective quality due to the “species mixture” being dominated by *Abies alba* and thus never fulfilling the minimal profile for this index.

**Fig. 13.**
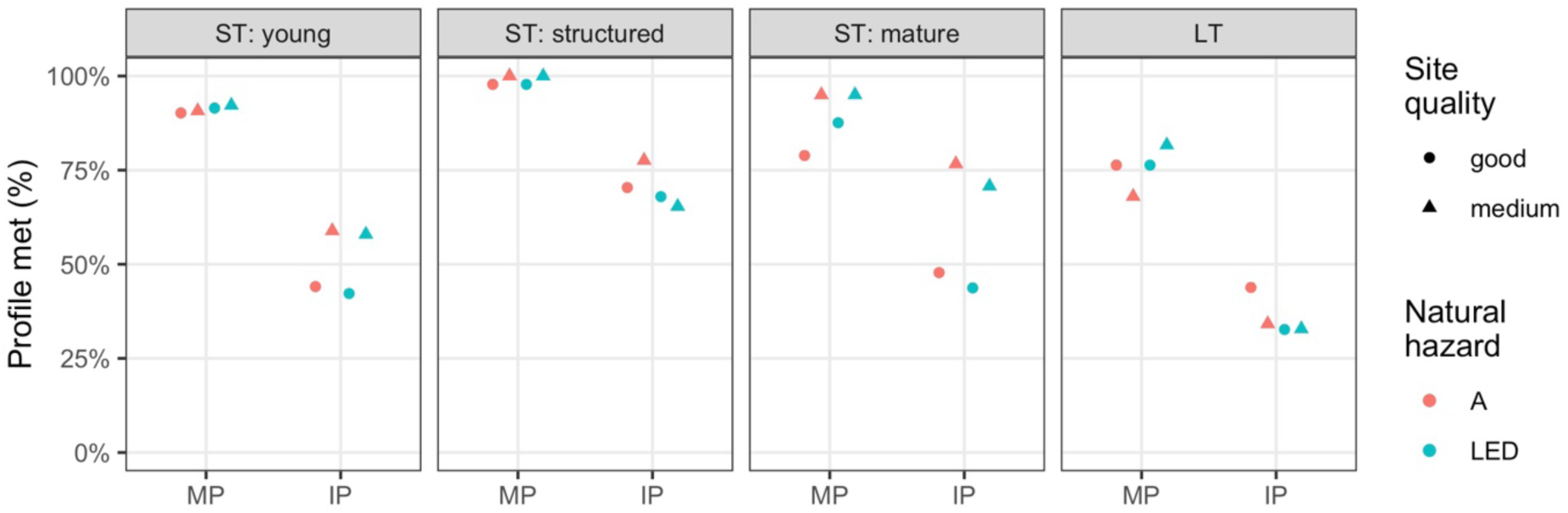
Protective quality of simulations without management (NOM) of elevational zone HM by initialization stand type (ST = short-term; LT = long-term; panels), and NaiS-profile (MP = minimal profile, sMP_abs_; IP = ideal profile, sIP_abs_). Symbols differentiate the site quality, colors the natural hazards snow avalanches (A) and landslide/erosion/debris flow (LED).

The mean variable importance across all HM models according to BRT was relatively similar for all four explanatory variables, ranging between 22 and 28%, but it differed between the simulation types and NaiS profiles (Fig. S7.7). The strongest deviations were found for *site quality*, which was of almost no importance in young stands when the minimal profile was taken as reference, but it was by far the most important variable determining *sIP_diff_* in mature stands.

The effects of forest management on the protective quality according to the beta regression models showed a clear dependence on the initialization stand type and the simulation length (Figs. 14-18). The prevalent natural hazard hardly affected the results except for ST simulations initialized with well-structured stands, where the reduction of protective quality was stronger in stands protecting against avalanches than those protecting against landslides/erosion/debris flow. The HM beta regression models had a mean pseudo-R^2^ of 0.67 (mean of 0.79 for the minimal profile models and 0.55 for the ideal profile models, Table S7.4).

**Fig. 14.**
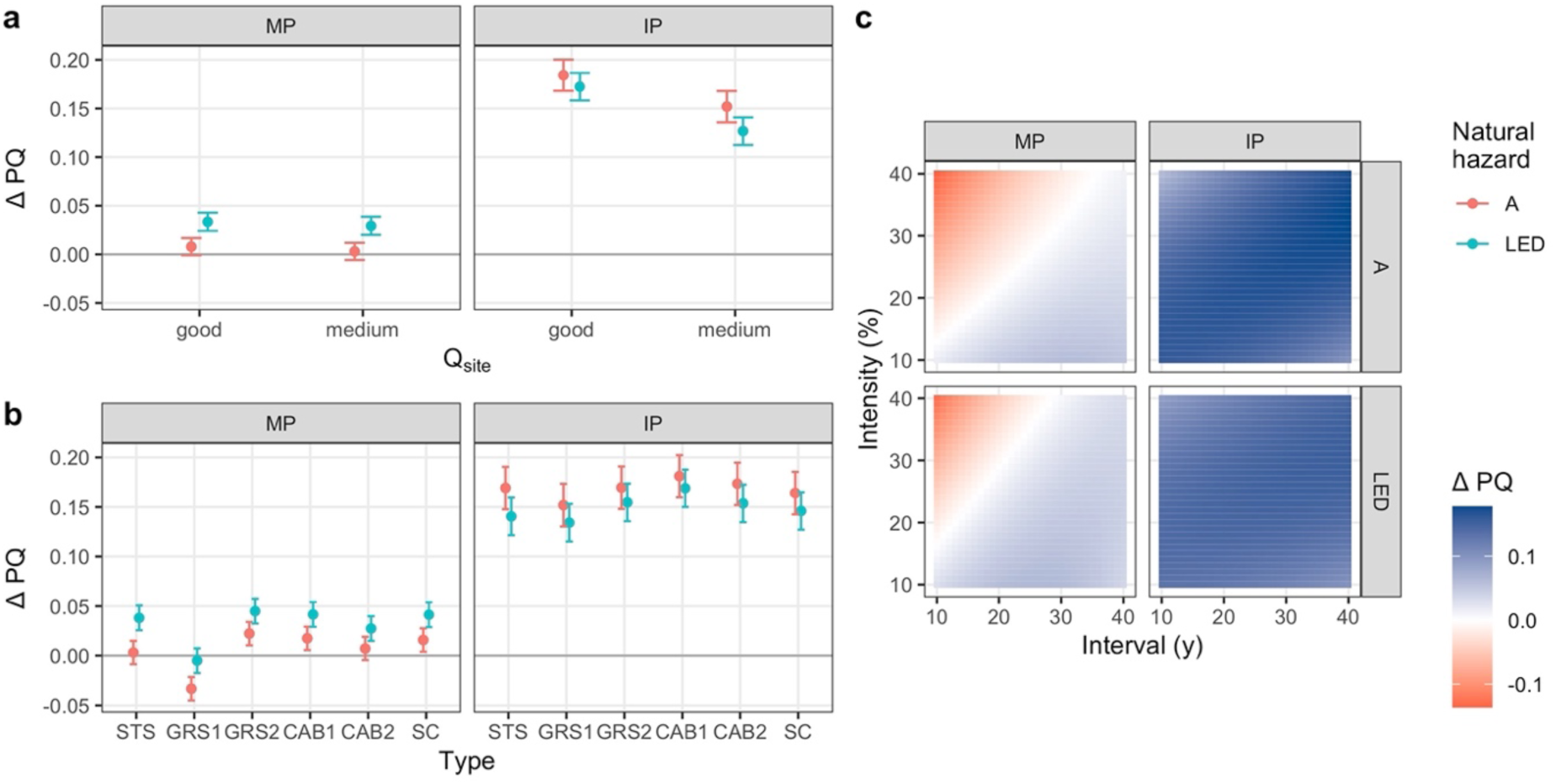
Marginal effects in reduced beta regression models explaining the effect of management on the protective quality (difference in protective quality compared to NOM, ΔPQ) in elevational zone **HM**, **short-term** simulations initialized with a **young stand**. a: mean effects and 95% confidence interval of site quality; b: mean effects and 95% confidence interval of management type; c: mean effects of interval and intensity. Separated by NaiS-profiles (MP = minimal profile; IP = ideal profile) and natural hazard (A = snow avalanches; LED = landslide/erosion/debris flow).

**Fig. 15.**
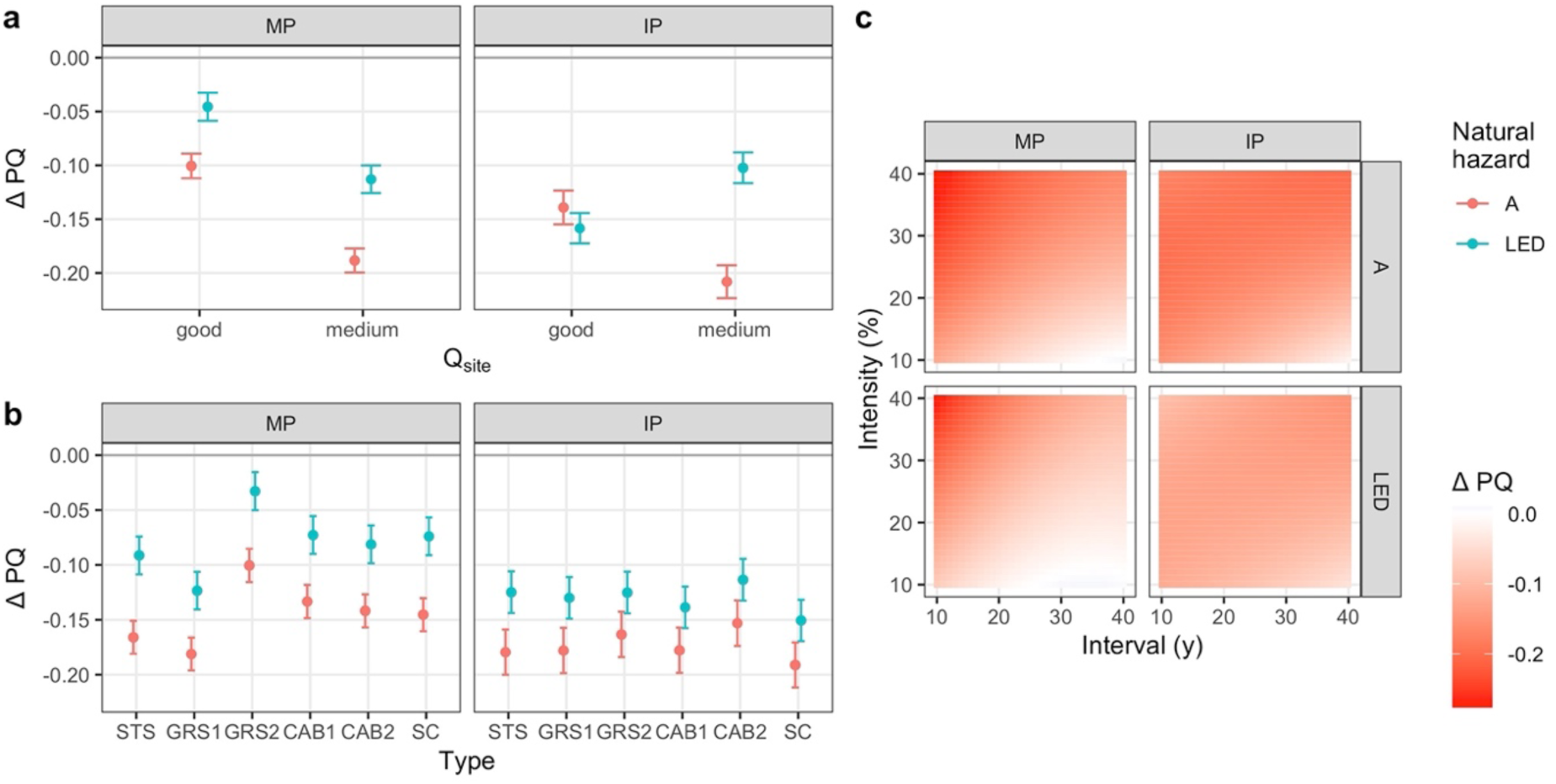
Marginal effects in reduced beta regression models explaining the effect of management on the protective quality (difference in protective quality compared to NOM, ΔPQ) in elevational zone **HM**, **short-term** simulations initialized with a **well-structured stand**. a: mean effects and 95% confidence interval of site quality; b: mean effects and 95% confidence interval of management type; c: mean effects of interval and intensity. Separated by NaiS-profiles (MP = minimal profile; IP = ideal profile) and natural hazard (A = snow avalanches; LED = landslide/erosion/debris flow).

**Fig. 16.**
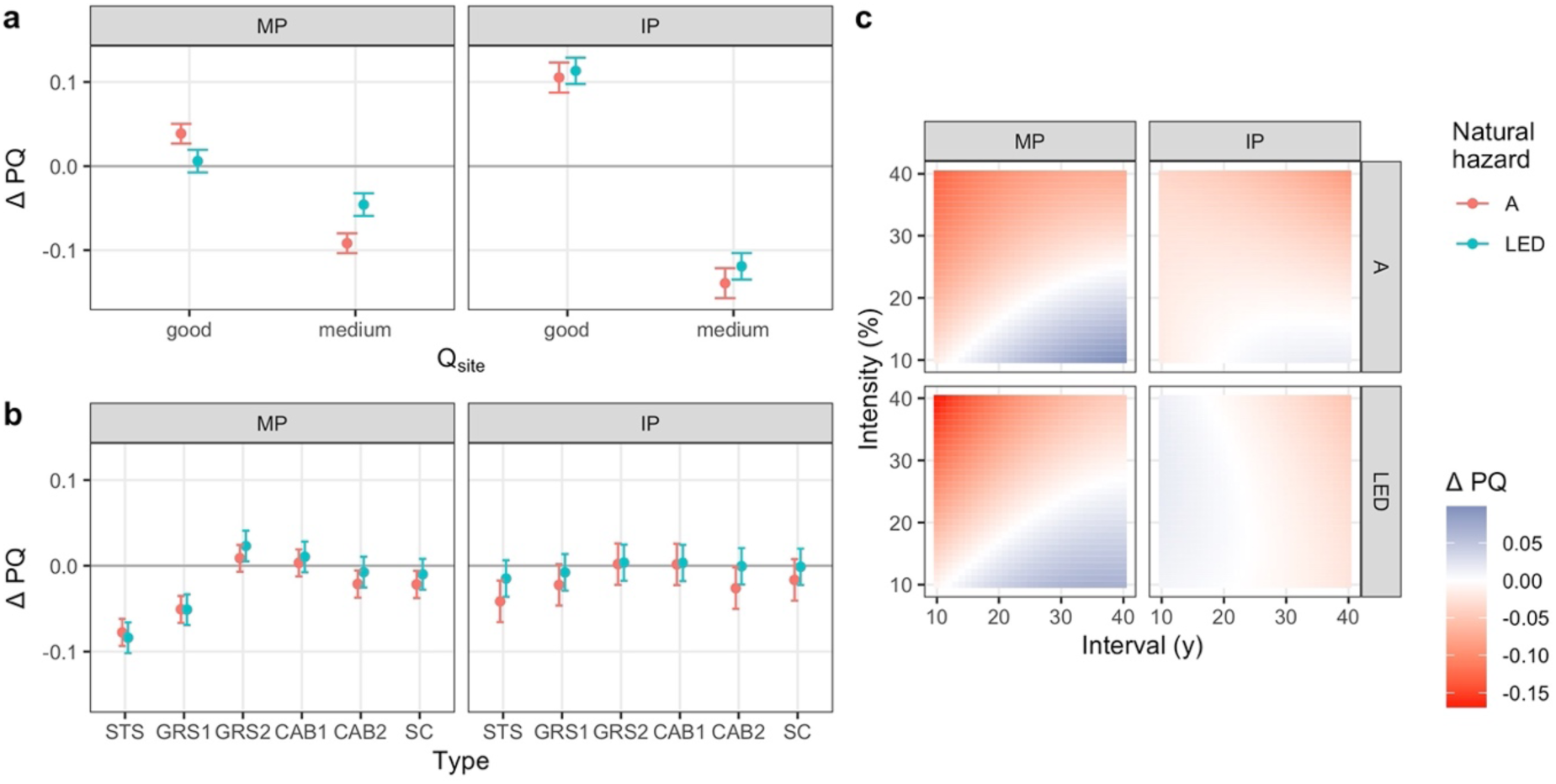
Marginal effects in reduced beta regression models explaining the effect of management on the protective quality (difference in protective quality compared to NOM, ΔPQ) in elevational zone **HM**, **short-term** simulations initialized with a **mature stand**. a: mean effects and 95% confidence interval of site quality; b: mean effects and 95% confidence interval of management type; c: mean effects of interval and intensity. Separated by NaiS-profiles (MP = minimal profile; IP = ideal profile) and natural hazard (A = snow avalanches; LED = landslide/erosion/debris flow).

**Fig. 17.**
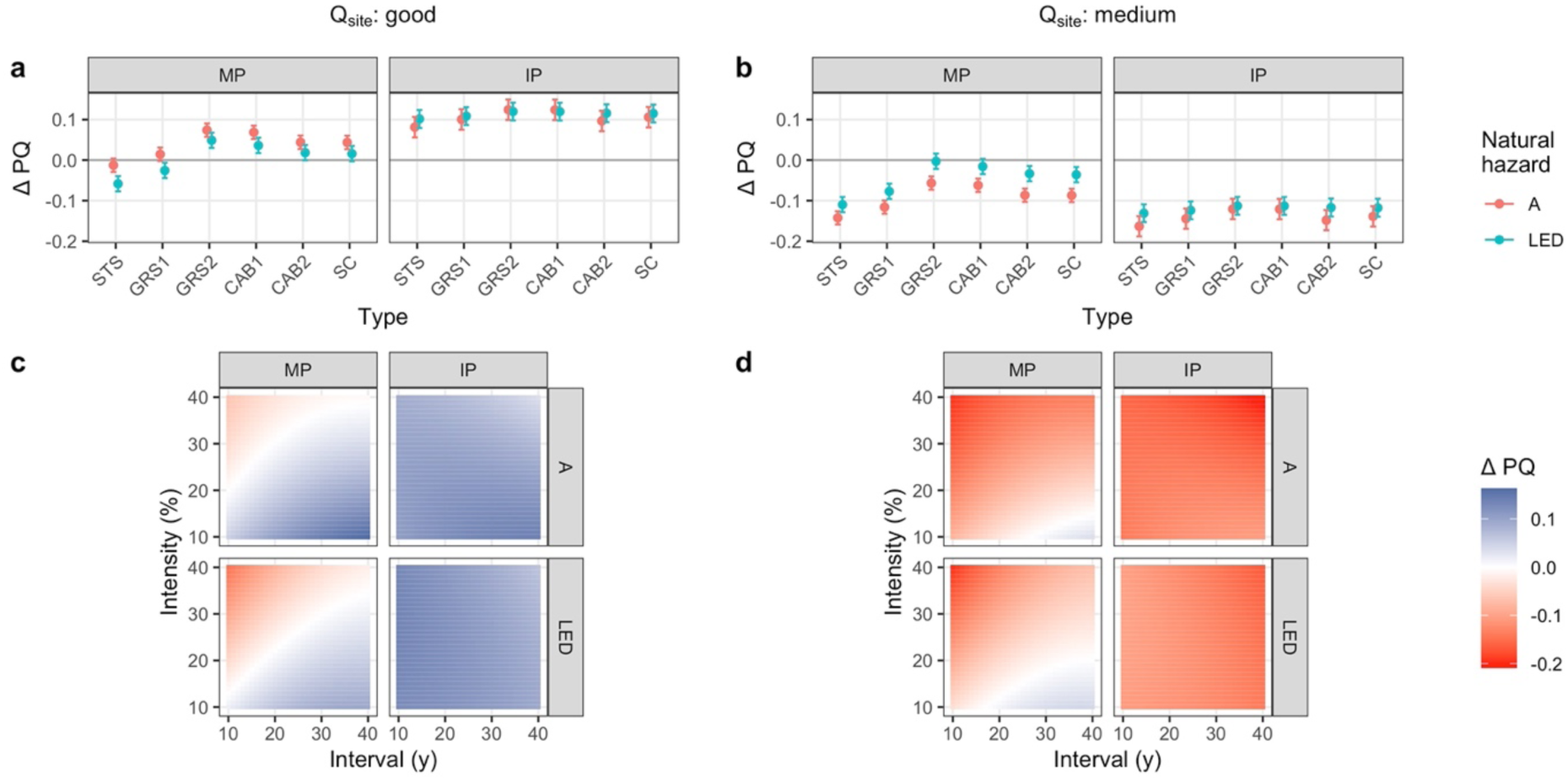
Marginal effects in reduced beta regression models explaining the effect of management on the protective quality (difference in protective quality compared to NOM, ΔPQ) in elevational zone **HM**, **short-term** simulations initialized with a **mature stand** with **site quality held fixed** at “good” (left column) and “medium” (right column). a & b: mean effects and 95% confidence interval of management type; c & d: mean effects of interval and intensity. Separated by NaiS-profiles (MP = minimal profile; IP = ideal profile) and natural hazard (A = snow avalanches; LED = landslide/erosion/debris flow).

**Fig. 18.**
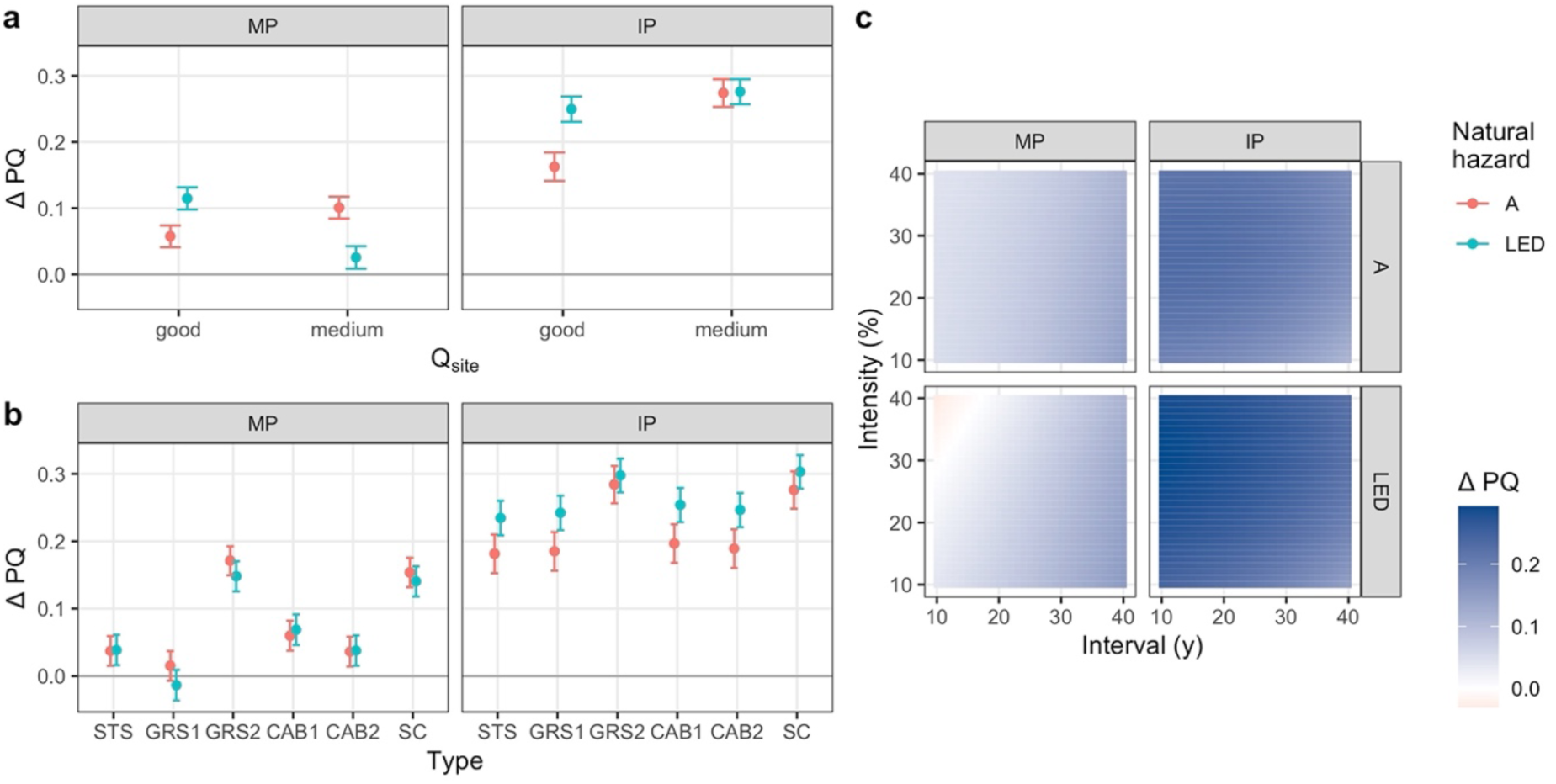
Marginal effects in reduced beta regression models explaining the effect of management on the protective quality (difference in protective quality compared to NOM, ΔPQ) in elevational zone **HM**, **long-term** simulations. a: mean effects and 95% confidence interval of site quality; b: mean effects and 95% confidence interval of management type; c: mean effects of interval and intensity. Separated by NaiS-profiles (MP = minimal profile; IP = ideal profile) and natural hazard (A = snow avalanches; LED = landslide/erosion/debris flow).

##### 3.2.2.1. Young stands

In ST simulations initialized with a young stand, forest management had a positive mean effect on the protective quality across almost all *management types* and combinations of *interval* and *intensity* (except for interventions with high frequency and intensity) (Fig. 14), even though *sMP_abs_* in NOM simulations was already around 90% (^F^ig. 13). Compared to these simulations, improvements towards fully reaching the minimal profile were theoretically possible in the “horizontal arrangement” during the first ca. 20 years of the simulation, and in the “stability of support trees” and for “saplings and thicket” towards the end of the simulation. Individual simulation results showed that the improvements were realized in the latter two indices while the remaining indices almost never dropped below the minimal profile.

The management type GRS1 performed worst, with slightly negative mean effects in both natural hazards. Improvements in the protective quality were still realized with GRS1 by less frequent and less intense interventions, but the negative effects on the protective quality were stronger than with other *management types* due to a slightly lower canopy cover (Figs. S6.9 and S6.10). The fulfillment of the ideal profile was improved with all simulated management regimes. Across almost all management regimes, even those that led to a decrease in *sMP_abs_*, the “species mixture” plus at least one further index were improved and fulfilled the ideal profile more often than under NOM. Overall, these results suggest that almost any management regime except for those with strong interventions at short intervals lead to an improvement of the protective quality.

##### 3.2.2.2. Well-structured stands

In NOM simulations initialized with a well-structured stand, the minimal profile was virtually always met and the ideal profile fulfilled to a relatively high degree (Fig. 13). Systematic deficits in fulfilling the entire ideal profile existed in the species composition of “saplings and thicket” (slightly more *Picea abies* than *Abies alba*) and in the “stability of support trees” in stands with good *site quality*. A positive effect of management on the protective quality measured by the minimal profile was not possible. All but very fine interventions at long intervals lowered certain indices at least temporarily and thus reduced the protective quality (Fig. 15c). The negative effect of management in avalanche protection forests was stronger (Fig. 15a/b) due to the stricter gap size thresholds compared to LED. The fulfillment of the ideal profile was also lowered by management, as opening gaps decreased the index value for “horizontal arrangement” and the slightly shifted diameter distribution was classified as less than ideal. The species distribution of “saplings and thicket” could only be shifted towards more *Abies alba* and therefore improved by relatively strong and frequent interventions which, however, had multiple negative effects on other indices. Therefore, not intervening at all during at least 90 years would be the best management option in well-structured stands.

##### 3.2.2.3. Mature stands

In ST simulations initialized with mature stands, *site quality* played a large role. This was evident from its high importance in BRT (Fig. S7.7), the different signs of the mean effects in beta regression models (Fig. 16) and the substantial difference in protective quality in NOM simulations (Fig. 13). The role of *site quality* was more pronounced when taking the ideal NaiS profile as a reference. The main difference in the protective quality of NOM simulations between the *site qualities* were in the “vertical structure” and “horizontal arrangement”. While the “vertical structure” mostly fulfilled the ideal profile on medium *site quality*, it only achieved the minimal profile on good *site quality* since the diameter distribution was more strongly dominated by the largest and the smallest trees with intermediate diameter classes being underrepresented. “Horizontal arrangement” remained almost ideal over time on medium *site quality* but dropped below the minimal profile within 30-40 years on good *site quality* due to both a decrease in canopy cover and an increase in gaps exceeding the threshold size. The different development of the unmanaged stands on either *site quality* led to very different effects of management on the protective quality (Fig. 17).

In stands with good *site quality*, all *management types* except for those creating the smallest openings (STS and GRS1) and all “reasonable” combinations of *interval* and *intensity* (i.e., *intensity* in % not being much higher than *interval* in years) improved the protective quality in terms of both the minimal and the ideal profile being fulfilled (Fig. 17a/c). Mainly “species mixture”, “horizontal arrangement”, and “stability of support trees” were improved by management. The best results were achieved with intermediate-sized openings (GRS2 and CAB1) and long return periods (≥ 30 years). On sites with medium *site quality*, the unmanaged stands provided a high degree of protection (Fig. 13), which was lowered by virtually all simulated management regimes (Fig. 17b/d). Even though the fulfillment of the minimal profile was improved to up to 100% with a low percentage of basal area removal (due to better “stability of the support trees”), it came at the cost of reducing the share of indices fulfilling the ideal profile (mainly the “vertical structure” and “horizontal arrangement”). Therefore, the best strategy would be not to intervene at all during at least 90 years.

##### 3.2.2.4. Long-term simulations

LT simulations led to the lowest protective quality of unmanaged stands within the high montane zone (Fig. 13). The major deficits in terms of the minimal profile were the “species mixture” (complete dominance by *Abies alba*), the “stability of support trees” on good *site quality* and the slightly too low canopy cover to fulfill the minimal threshold in avalanche protection forests on medium *site quality*. The stands were two-layered with mainly very small and very large trees, while the intermediate size classes were underrepresented. Forest management had a strong positive effect on the protective quality (Fig. 18). With all management regimes, a more balanced “species mixture” that at least fulfilled the minimal profile could be produced. Medium-sized openings as realized with *management types* GRS2 and SC led to the best overall protective quality, while small (STS and GRS1) and large openings (CAB2) produced deficits in the “vertical structure” and “horizontal arrangement”. The recommended long-term management regime would thus consist of medium-sized openings, while the options regarding intensity and interval are very large.

#### 3.2.3. Subalpine zone

In the NOM simulations of the subalpine zone, the minimal profile was met by ca. 65-85% of the indices over time (*sMP_abs_*), the ideal profile by 50-70% (*sIP_abs_*) (^F^ig. 19). In all simulations except for the ST simulations initialized with a young stand, the index for “horizontal arrangement” was mostly below the minimal profile. The main reason was the low canopy cover (ca. 40%) plus at times up to three gaps per hectare exceeding the threshold size values. Simultaneously, support trees had crown lengths of only 25 to 60%, thus not meeting the minimal profile threshold of 70% either. In LT simulations, the largest (and thus among the oldest) trees even had below-average crown lengths. As *Picea abies* is the only species included in the SA simulations, the indices of “species mixture” and “seedlings” by definition fulfilled the ideal profile fully over time. The index for “saplings and thicket” also fulfilled the ideal profile over time in all simulations except for those initialized with a young stand where the area covered by regeneration dropped below the threshold values towards the end of the simulations. A successful management regime thus had to improve or maintain “vertical structure” and “horizontal arrangement” as well as stand stability.

**Fig. 19.**
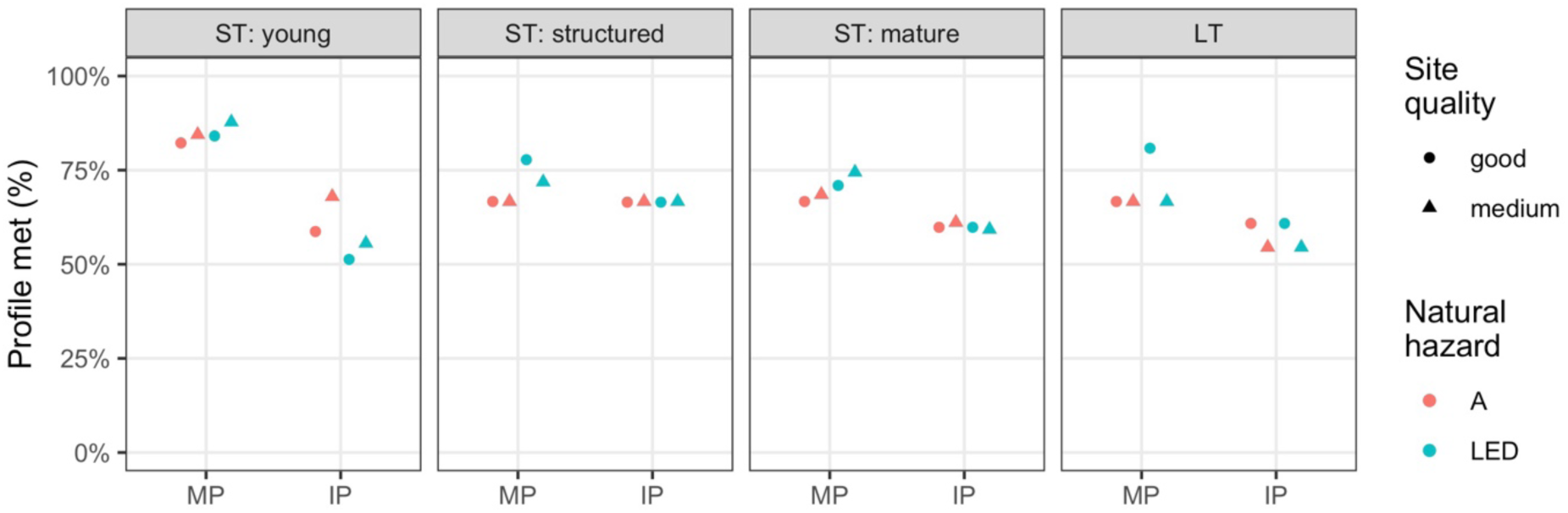
Protective quality of simulations without management (NOM) of elevational zone SA by initialization stand type (ST = short-term; LT = long-term; panels), and NaiS-profile (MP = minimal profile, sMP_abs_; IP = ideal profile, sIP_abs_). Symbols differentiate the site quality, colors the natural hazards snow avalanches (A) and landslide/erosion/debris flow (LED).

In the BRT models of the subalpine zone, the management-related variables had similar mean importance, while *site quality* had a lower average importance of ca. 20% (Fig. S7.7). However, in one of the LT models (natural hazard LED & response variable *sIP_diff_*), *site quality* was disproportionately important. The effect of management according to the beta regression models depended strongly on the initialization stand type and the natural hazard (Figs. 20-23). The minimal NaiS profiles differ for the two natural hazards in terms of allowed gap size (750 m^2^ for A, 600 m^2^ for LED in the case of insufficient regeneration) and the required degree of canopy cover (50% for A, 40% for LED). The SA beta regression models had a mean pseudo-R^2^ of 0.64 (mean of 0.53 for the minimal-profile-models and 0.75 for the ideal-profile models, Table S7.4)

**Fig. 20.**
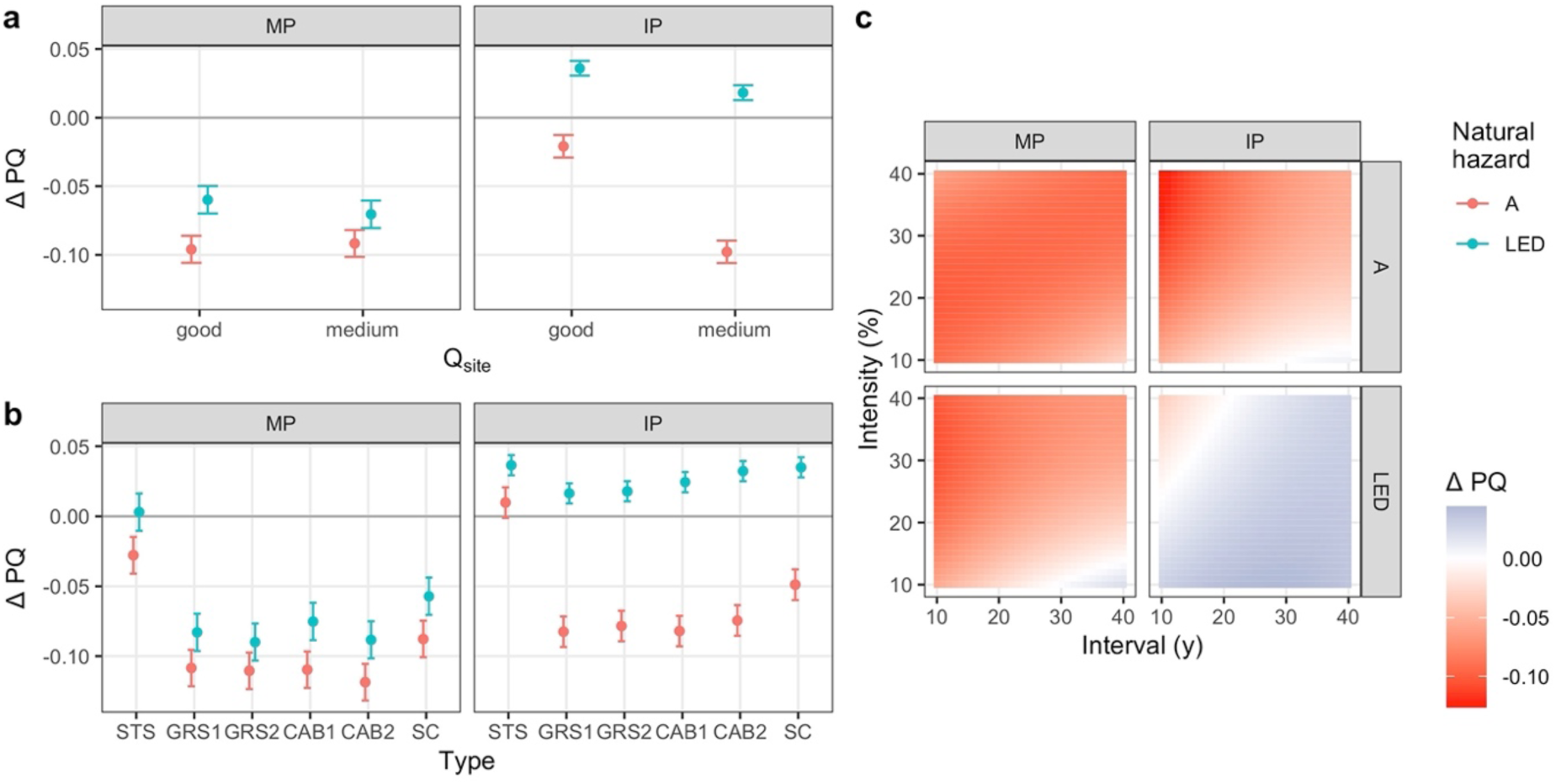
Marginal effects in reduced beta regression models explaining the effect of management on the protective quality (difference in protective quality compared to NOM, ΔPQ) in elevational zone **SA**, **short-term** simulations initialized with a **young stand**. a: mean effects and 95% confidence interval of site quality; b: mean effects and 95% confidence interval of management type; c: mean effects of interval and intensity. Separated by NaiS-profiles (MP = minimal profile; IP = ideal profile) and natural hazard (A = snow avalanches; LED = landslide/erosion/debris flow).

**Fig. 21.**
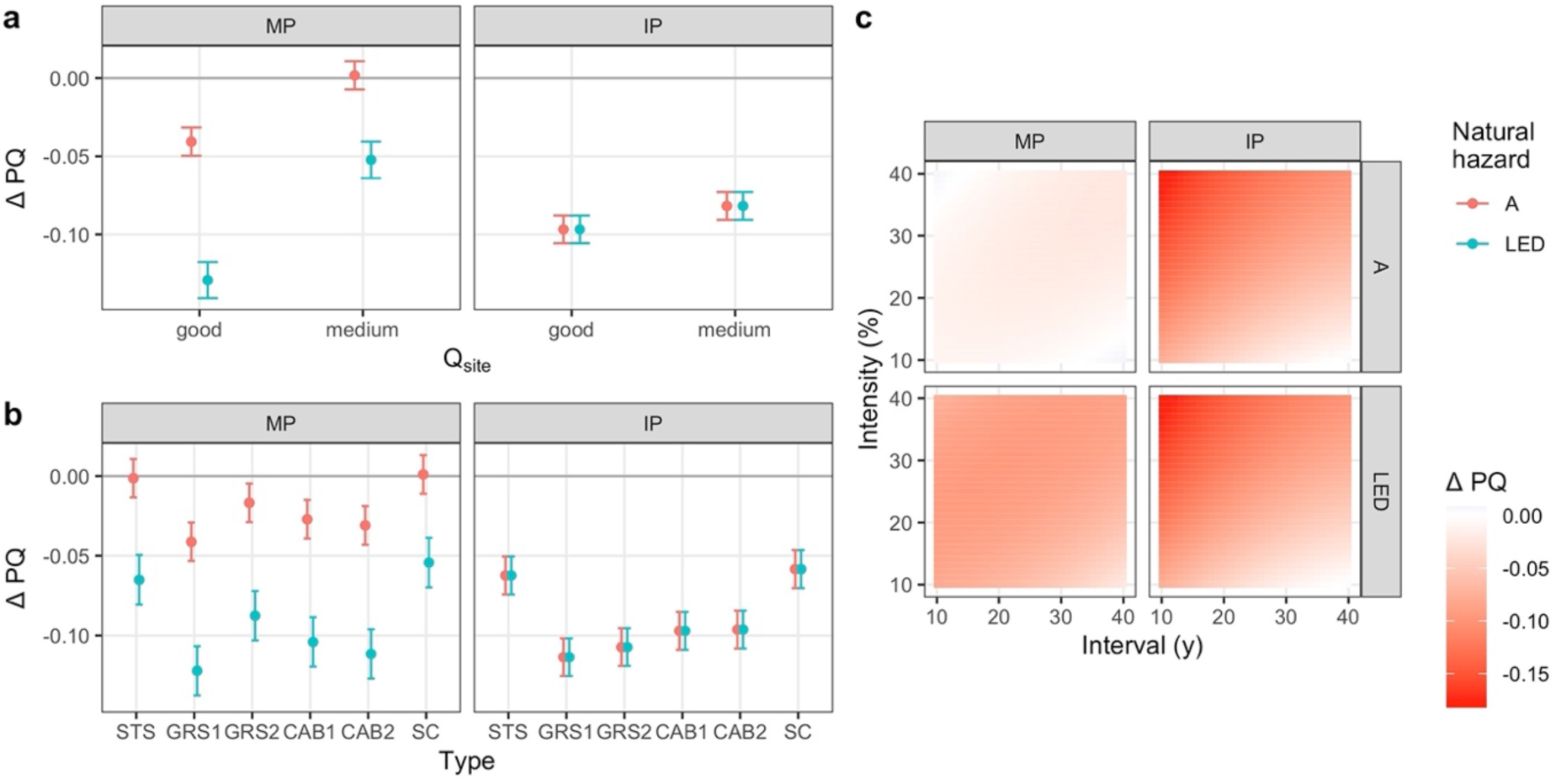
Marginal effects in reduced beta regression models explaining the effect of management on the protective quality (difference in protective quality compared to NOM, ΔPQ) in elevational zone **SA**, **short-term** simulations initialized with a **well-structured stand**. a: mean effects and 95% confidence interval of site quality; b: mean effects and 95% confidence interval of management type; c: mean effects of interval and intensity. Separated by NaiS-profiles (MP = minimal profile; IP = ideal profile) and natural hazard (A = snow avalanches; LED = landslide/erosion/debris flow).

**Fig. 22.**
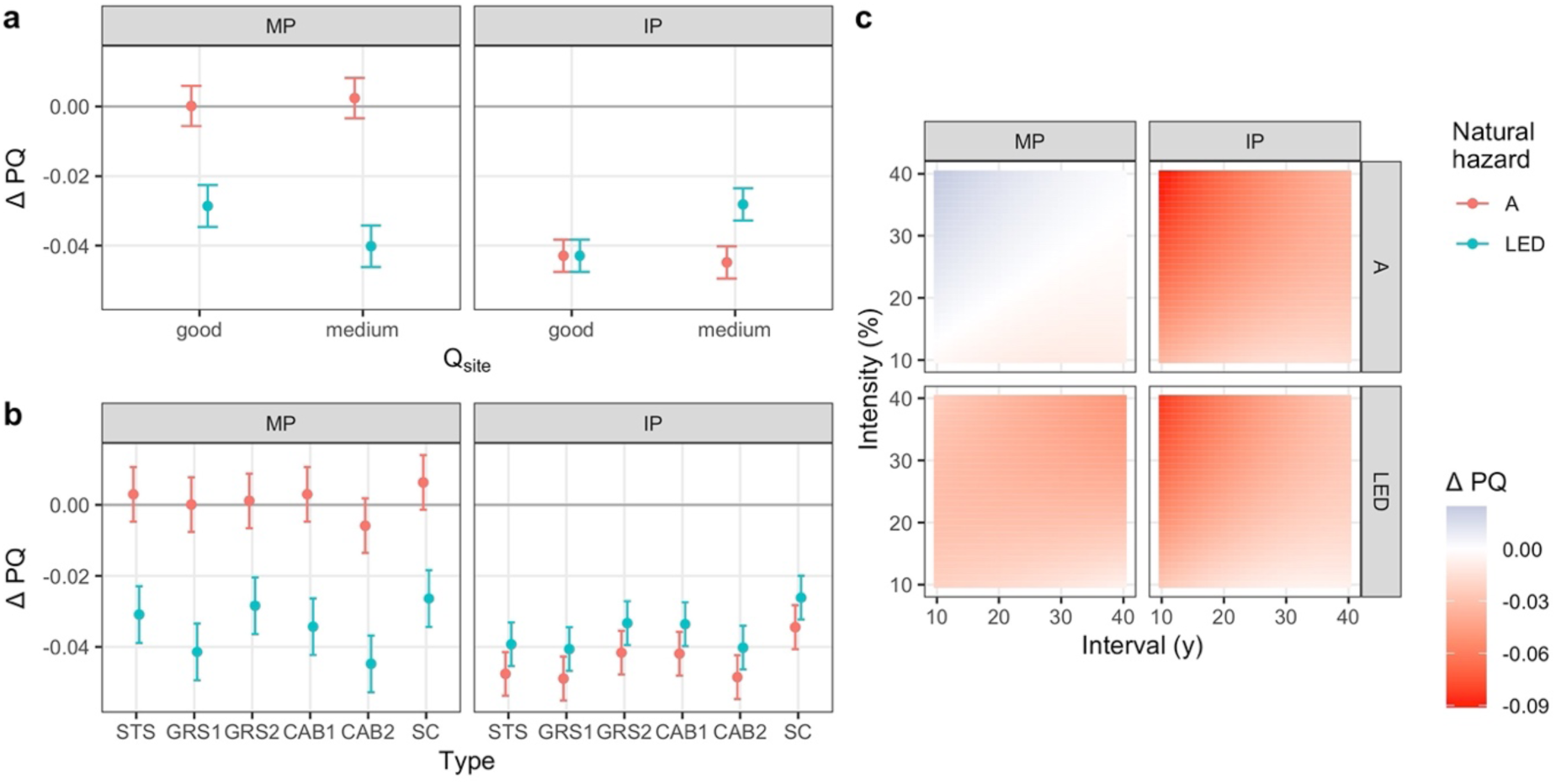
Marginal effects in reduced beta regression models explaining the effect of management on the protective quality (difference in protective quality compared to NOM, ΔPQ) in elevational zone **SA**, **short-term** simulations initialized with a **mature stand**. a: mean effects and 95% confidence interval of site quality; b: mean effects and 95% confidence interval of management type; c: mean effects of interval and intensity. Separated by NaiS-profiles (MP = minimal profile; IP = ideal profile) and natural hazard (A = snow avalanches; LED = landslide/erosion/debris flow).

**Fig. 23.**
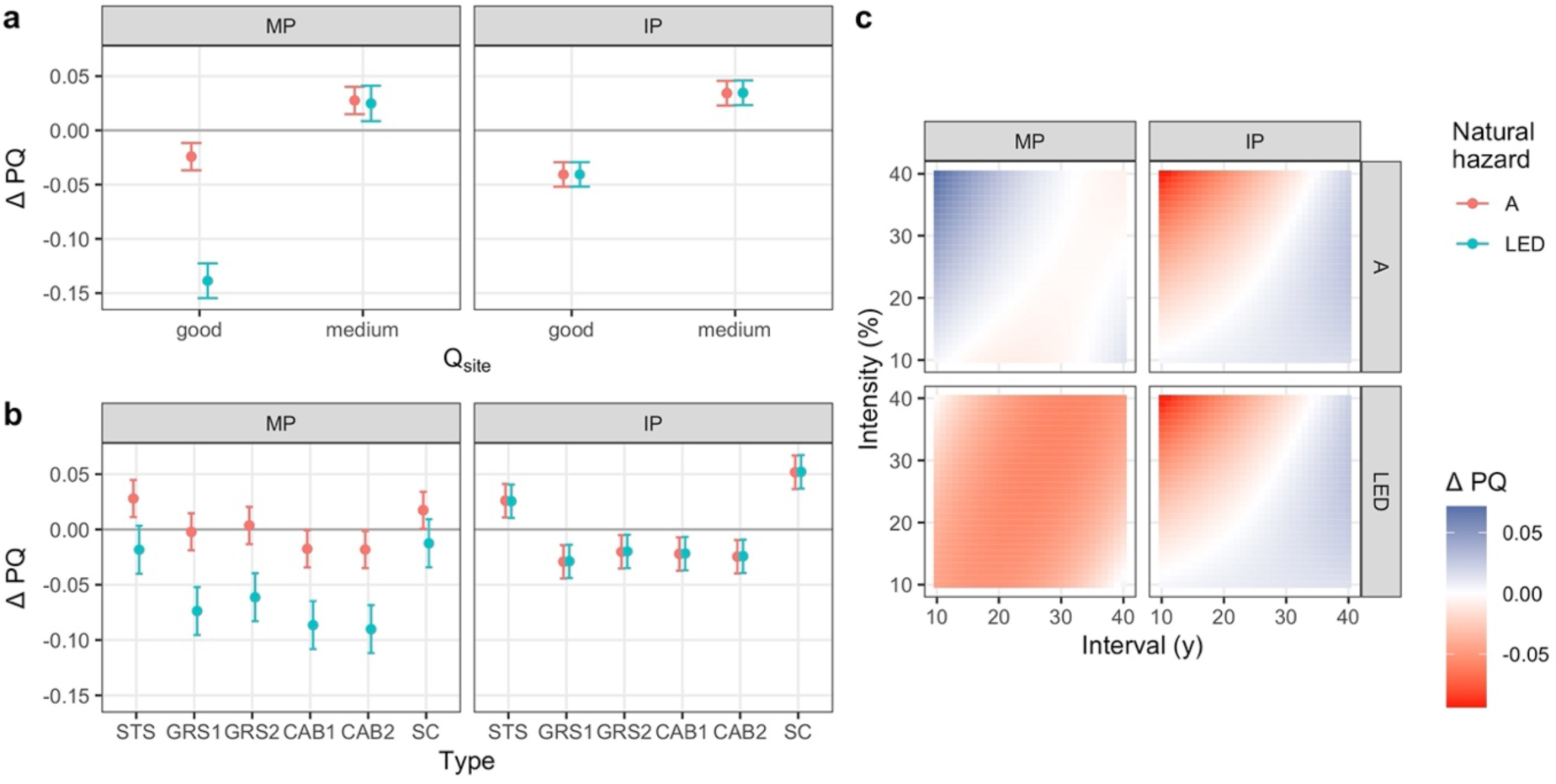
Marginal effects in reduced beta regression models explaining the effect of management on the protective quality (difference in protective quality compared to NOM, ΔPQ) in elevational zone **SA**, **long-term** simulations. a: mean effects and 95% confidence interval of site quality; b: mean effects and 95% confidence interval of management type; c: mean effects of interval and intensity. Separated by NaiS-profiles (MP = minimal profile; IP = ideal profile) and natural hazard (A = snow avalanches; LED = landslide/erosion/debris flow).

##### 3.2.3.1. Young stands

In SA avalanche protection forests initialized with young stands, protective quality was lowered by practically all simulated management regimes (Fig. 20). In LED protection forests, however, the fulfillment of the minimal profile was increased by infrequent, light interventions, and that of the ideal profile by most regimes. Interventions generally led to an improvement of “vertical structure” and the cover of “saplings and thicket”. However, even the light and infrequent interventions worsened “horizontal arrangement” by the formation of gaps and the lowering of canopy cover. Even though this index was reduced under both natural hazards, the overall effect on the ideal profile differed as the “horizontal arrangement” of unmanaged avalanche protection forests was deemed ideal, while that of unmanaged LED protection forests was deemed worse due to slightly differing threshold values. In all cases, the *management type* STS outperformed the other types. By removing only single trees and thus avoiding the formation of gaps, “horizontal arrangement” was barely worsened. Even without management, there was just enough regeneration towards the end of the simulation to fulfill the minimal profile. Therefore, these spatially fine interventions that hardly produce additional regeneration had no adverse effect on the protective quality. Based on these results and over the course of 90 years, STS with low intensity and long intervals is the recommended management regime for LED protection forests, while no interventions would be best in A protection forests.

##### 3.2.3.2. Well-structured stands

Forest management generally reduced the protective quality of well-structured stands compared to unmanaged ones (Fig. 21). Besides “horizontal arrangement” and “stability of support trees”, all indices in NOM simulations were rated as ideal. However, both canopy cover and the mean crown ratio of the support trees were only around 40%. Management interventions inevitably lowered canopy cover even further. The mean crown ratio of support trees was improved only by management regimes that led to a complete removal of the old stand within a few decades, which evidently led to a decrease in the overall protective quality. The minimal threshold for canopy cover in LED protection forests is 40%, which was partially fulfilled without management, while the threshold of 50% for avalanche protection forests was not. Due to these different threshold values, the further reduction through interventions led to a decrease in *sMP_abs_* in LED protection forests. Some management regimes, especially *management types* STS and SC in avalanche protection forests with medium *site quality*, led to a positive *sMP_diff._* However, this “positive” effect was produced by very strong and frequent interventions, resulting in stands with very low basal area and canopy cover (Fig. S6.19). These results suggest that no interventions are necessary in well-structured SA stands.

##### 3.2.3.3. Mature stands

When mature SA stands were left unmanaged, the initially extremely dense stands (basal areas of 60 to 80 m^2^/ha) experienced a strong decrease in basal area, stem number and canopy cover, the latter down to around 30%. Large gaps appeared, leading to a high cover of regeneration, while the remaining stand was dominated by few large trees. “Vertical structure” was considered ideal for the first half of the simulations and minimal for the remainder, while the crown ratios were below the threshold of the minimal profile for the entire duration of the simulation. LED protection forests scored slightly higher regarding *sMP_abs_* than avalanche protection forests (Fig. 19) because “horizontal arrangement” still fulfilled the minimal profile in the first decades of the simulation due to different threshold values for canopy cover. Strong and frequent interventions led to an increase in the mean crown ratio of the support trees, at the cost of worsening “vertical structure” and “horizontal arrangement” (i.e., strong reduction of basal area and large parts of the stand converted to regeneration). In the statistical models, this translated into a positive overall effect on *sMP_diff_* in avalanche protection forests only (Fig. 22), as “horizontal arrangement” was already below the minimal profile in NOM simulations. The slight negative effect in both the minimal and ideal profiles of more fine and less frequent interventions resulted from the faster deterioration of “vertical structure” and “horizontal arrangement” compared to NOM, while the “stability of support trees” could not be increased substantially. Overall, the “naturally” decreasing protective quality of unmanaged stands could hardly be increased, irrespective of *site quality* and *management type*, leaving NOM as the recommended management option.

##### 3.2.3.4. Long-term simulations

In LT simulations of SA stands, the impact of management on the protective quality strongly depended on stand *site quality*. This was evident from its high variable importance in BRT for *sMP_diff_* and the natural hazard LED (Fig. S7.7), and the different signs of the effect for the two levels in the beta regression models (Fig. 23). LED protection forests had a higher degree of fulfillment of the minimal profile in stands with good *site quality*, with the difference lying in the index for “horizontal arrangement”. Canopy cover was higher on good sites and just above the threshold of 40% to fulfill the minimal profile of LED, but not sufficient for the 50% required in avalanche protection forests. This difference led to a generally negative effect of management on good *site quality* but a positive one on medium *site quality* (Fig. 24).

**Fig. 24.**
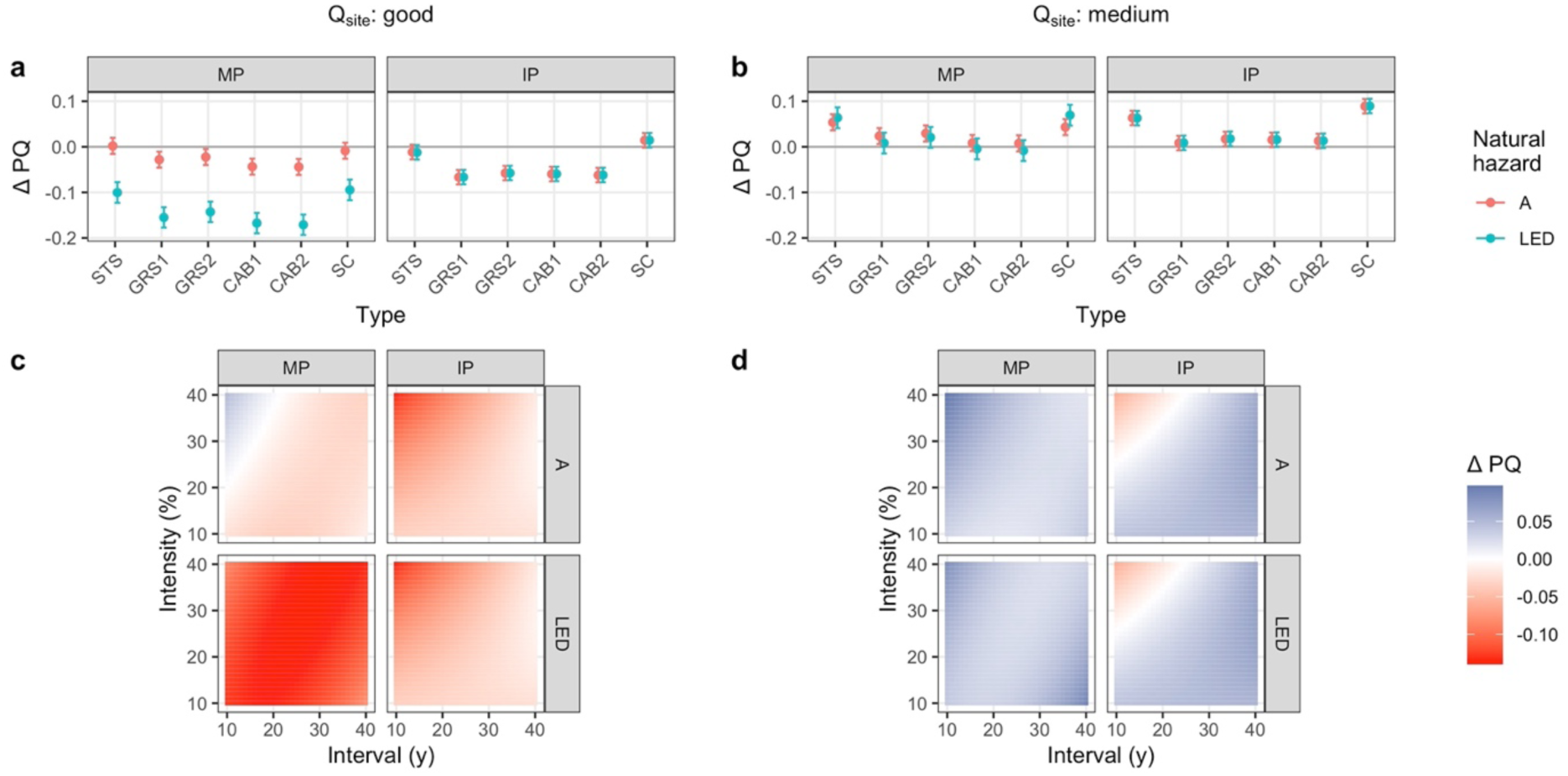
Marginal effects in reduced beta regression models explaining the effect of management on the protective quality (difference in protective quality compared to NOM, ΔPQ) in elevational zone **SA**, **long-term** simulations with **site quality held fixed** at “good” (left column) and “medium” (right column). a & b: mean effects and 95% confidence interval of management type; c & d: mean effects of interval and intensity. Separated by NaiS-profiles (MP = minimal profile; IP = ideal profile) and natural hazard (A = snow avalanches; LED = landslide/erosion/debris flow).

In stands with good *site quality*, a positive impact of management was only possible in avalanche protection forests with strong and frequent interventions (Fig. 24c). In these simulations, the index for “vertical structure” did not drop below the minimal profile, the one for “stability of support trees” was increased, and “horizontal arrangement” remained below the minimal profile. Even though this was rated as an overall increase in protective quality, the resulting stands with very low basal area and canopy cover would not be considered “good” protection forests in practice. In LED protection forests, *sMP_abs_* was generally lowered by management, mainly due to the inevitable reduction in canopy cover and thus the index for “horizontal arrangement” being rated below the minimal profile. The fulfillment of the ideal profile was lowered with both natural hazards due to at least slight decreases in the rating of “vertical structure”. Only with the *management type* SC and light interventions at long intervals, the fulfillment of the ideal profile could be increased (Fig. 24a, S6.23, and S6.24), as this led to an ideal “vertical structure” over time. These management regimes also led to little or no decrease in the fulfillment of the minimal profile and would therefore be the recommended regime, based on these simulation results.

In stands with a medium *site quality*, most management regimes led to an increase in the protective quality for both natural hazards and both the minimal and ideal profile (Fig. 24b/d). The degree of fulfillment of the minimal profile was increased most by frequent and rather strong interventions (with best outcome using *management type* STS) mainly by increasing the mean crown lengths of support trees but leading to stands with very low basal area and thus having the same problem as described above. However, management of type SC (except if frequent and strong interventions were used) had a positive effect on the fulfillment of the ideal profile, and a neutral or slightly positive effect on the fulfillment of the minimal profile (Fig. 24b/d, S6.23, and S6.24). With long return periods, “vertical structure” remained ideal throughout the simulation, while “horizontal arrangement” could also be improved somewhat. In the long term, the best-performing management regime would be SC at long intervals, almost regardless of the intervention intensity.

### 3.3. Management recommendations

The management recommendations based on the analysis of the beta regression models and individual simulation results are summarized in Table 1. For more detailed explanations, see the respective sections and the Discussion.

**Table 1.**
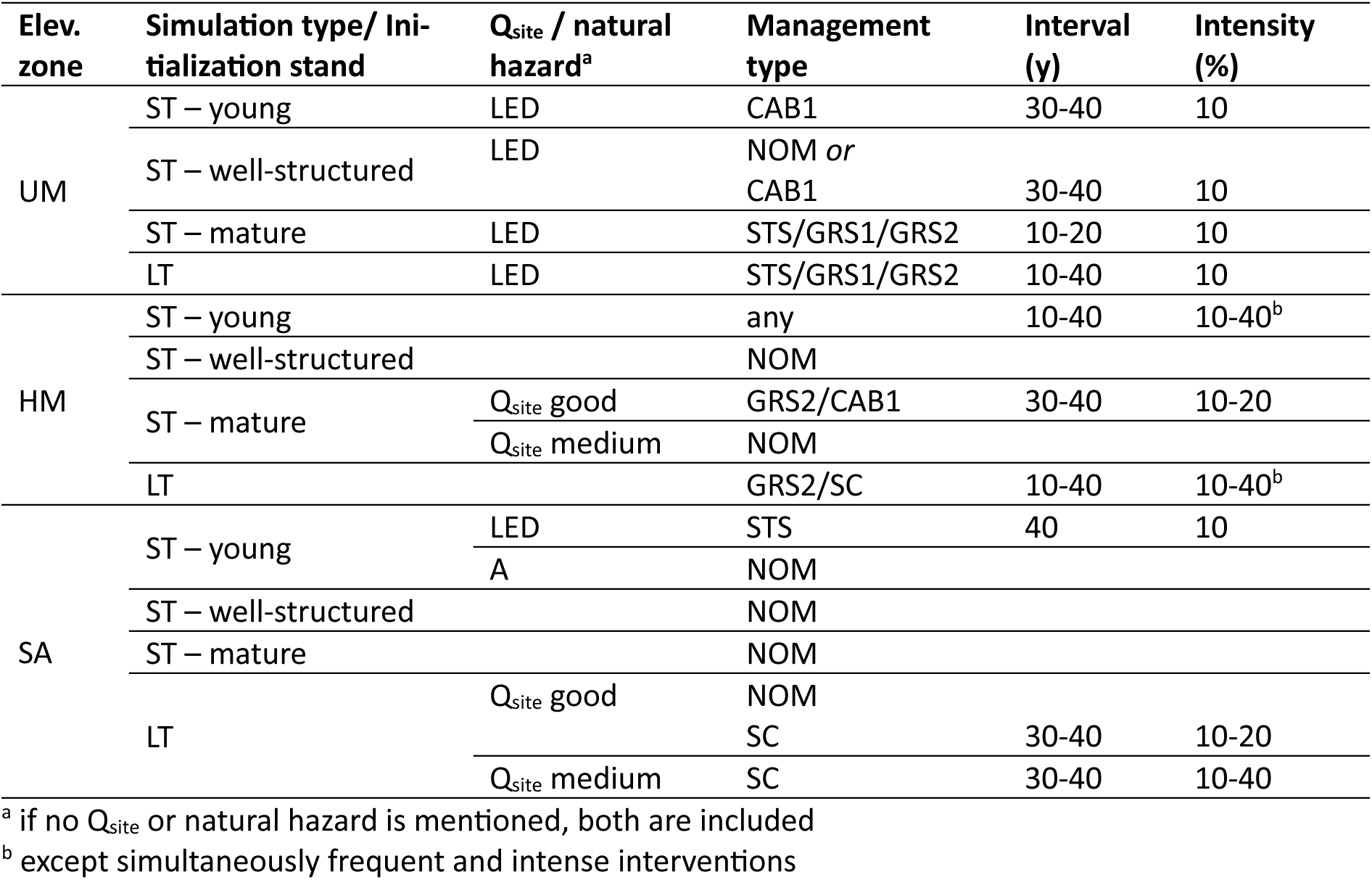
Summarized management recommendations.

### 3.4. Species performance in ProForM

In order to understand why *Abies alba* dominated the species composition in elevational zones UM and HM, we analyzed the maximum growth rates in ProForM and the realized maximum cohort ages in the LT-NOM simulations. In the elevational zone UM, the coniferous species clearly outperformed the deciduous species both in the maximum height growth rates of regeneration (DBH < 12 cm, Fig. 25a) and the maximum diameter growth rates of adult trees (DBH ≥ 12 cm, Fig. 25b). Furthermore, the 95%-quantiles of maximum realized cohort age were also larger for the coniferous than the deciduous species, with the maximum age of *Fagus sylvatica* being by far lowest among the four species (Fig. 25c). While *Abies alba* had slightly lower maximum height growth rates than *Picea abies* in the stage of regeneration, its maximum diameter growth rate as an adult tree as well as the maximum realized cohort age were higher. In the elevational zone HM, *Picea abies* had higher maximum height growth rates up to a DBH of 12 cm, but *Abies alba* had higher maximum diameter growth rates as adult trees and a larger maximum cohort age.

**Fig. 25.**
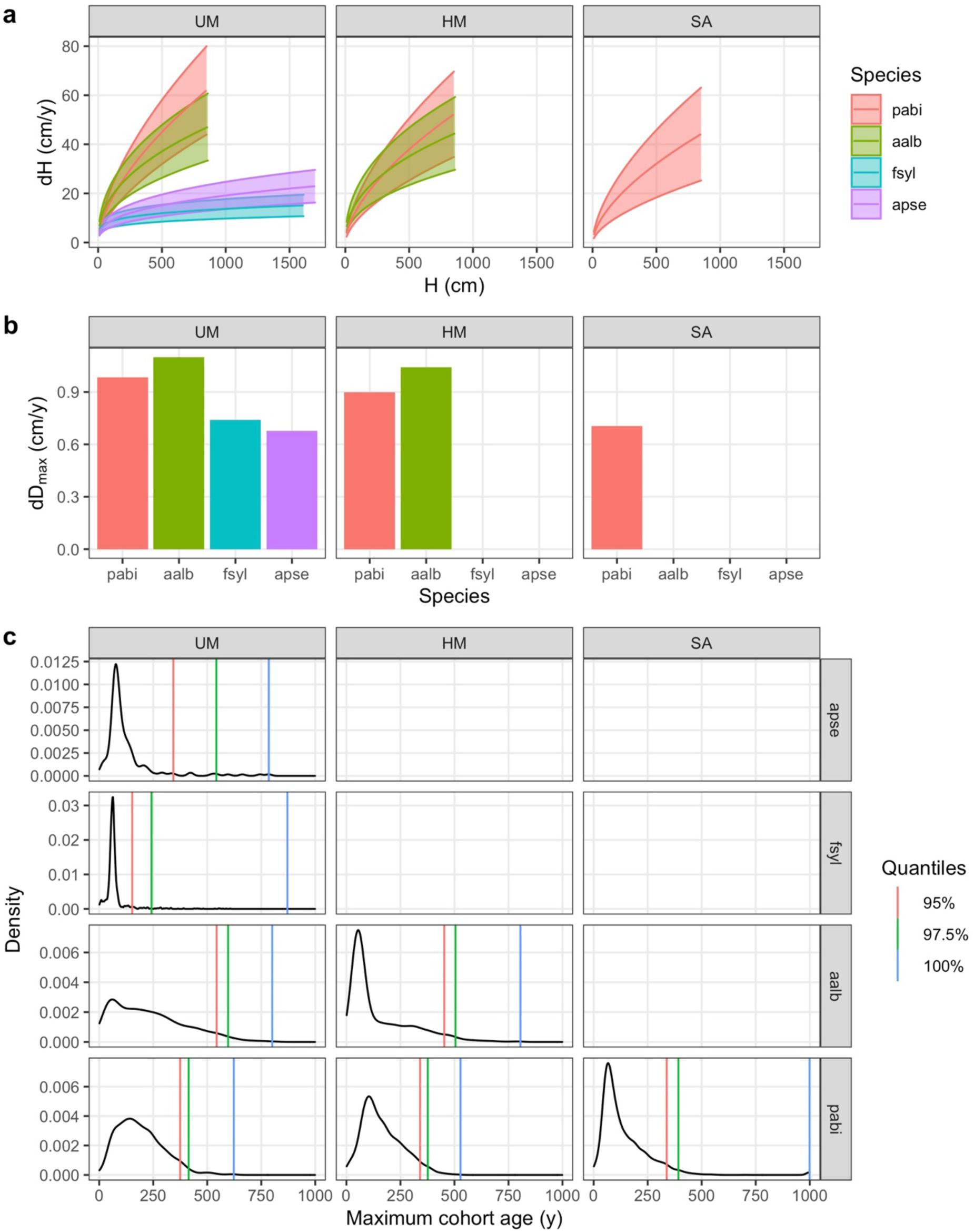
Species performance in ProForM. (a) Maximum annual height growth (dH) as a function of tree height (H) of regeneration (DBH < 12 cm) per species and elevational zone. The upper ribbon limits correspond to growth rates at good, the middle line to medium, and the lower ribbon limit to low regeneration quality. The maximum height (H) depicted per species corresponds to the height at which a DBH of 12 cm is reached. (b) Maximum annual diameter growth rates (dD_max_) of adult trees (DBH ≥ 12 cm) per species and elevational zone. (c) Density distributions of maximum cohort ages in long-term-simulations without management (NOM) per species and elevational zone, and the respective quantiles.

## 4. Discussion

In this study, we analyzed the effect of management and regeneration failure on the level of provision of the ecosystem service “protection against gravitational natural hazards” of mountain forest stands using a dynamic simulation model. This was done across large environmental gradients of climate (elevational zone), growth conditions, and browsing pressure, and using different initial stand conditions. We furthermore applied a series of management regimes differing in return interval, intervention intensity, and the spatial aggregation of removed trees. We derived short- and long-term management recommendations under the assumptions of (1) a constant climate and (2) the absence of large disturbances. Many simulation studies have investigated similar aspects regarding the protective effect of mountain forests (Moos et al. (2023) provide a good overview), but to our knowledge an extensive comparison of management strategies for protection forests at the stand scale has not been done to date, and this is the gap we are filling with the present analyses.

The following discussion is structured into four parts. First, we discuss the effects of high browsing pressure on the protective quality and its consequences for management options. Second, the management recommendations for stands with unhindered regeneration are discussed in general and for each of the three elevational zones. Third, the consequences of the most important methodological choices are examined and, finally, we discuss the reasons and possible solutions for deficiencies in ProForM that were revealed by this study.

### 4.1. Consequences of high browsing pressure on the protective quality

The joint analysis of the simulation results with “good” and “low” regeneration quality, i.e., low and very high levels of ungulate browsing, clearly revealed that a severely hindered regeneration process (germination and growth) leads to a much lower protective quality compared to situations with low browsing pressure, both in the short and long term. The effects of regeneration failure were evident in most stand characteristics, especially the species composition of both young and adult trees due to the selectivity of browsing regarding tree species (Didion et al., 2009; Motta, 1998; Ramirez et al., 2018; Schulze et al., 2014). Further effects were unsustainable diameter distributions and gaps – formed by natural or anthropogenic processes – that remained open for a long time. These findings are in line with previous studies, all having found negative effects of browsing on the protective quality (Didion et al., 2009; Wehrli et al., 2007b; Woltjer et al., 2008) or stand resilience (Thrippleton et al., 2020).

Unfortunately, unsustainably high levels of browsing damage on a local to regional level and the absence of viable regeneration of *Abies alba* are a reality in many regions across the Alps (Brang, 2017; Kupferschmid et al., 2015; Senn and Suter, 2003). Our results underline that the effect of severe browsing is stronger than the one of forest management, as previously reported for rockfall protection forests (Woltjer et al., 2008). Furthermore, we found that there was almost no possibility of substantially counteracting these negative effects through thinning or logging. In reality, under such conditions forest managers have to resort to protecting regeneration through fences or chemically, and/or planting artificial regeneration – management options that were not included in our study. Thus, our results support the view that high browsing levels can hinder regeneration to the point that the quality of protection forests is insufficient, independent of the sylvicultural treatment applied.

In addition, several tree species that are expected to gain in importance in mountain forests due to climate change, e.g., *Acer* spp. or *Quercus* spp., are heavily browsed (Kupferschmid et al., 2015), which poses a serious problem for adapting mountain forests to a future climate at reasonable cost (Brang, 2017). For maintaining functioning protection forests, it is therefore of utmost importance to manage ungulate densities on a regional scale to a level that allows for all main tree species to naturally regenerate in sufficient densities.

### 4.2. Management recommendations for protection forests

#### 4.2.1. General considerations

As management options under heavy browsing turned out to be severely limited in our simulations, we only discuss the low-browsing results here. They clearly show that successful protection forest management is highly site-specific. In the full models, elevational zone and the initialization stand type were among the most important explanatory variables. The effects of management type, intervention intensity, and return interval differed strongly among the site-specific reduced models. However, the changes in protective quality induced by management were relatively small, with most mean effect sizes in the statistical models per elevational zone and initialization stand type being smaller than ± 0.2 index values, corresponding to about one index newly fulfilling the minimal profile throughout the entire simulation period, or not fulfilling it any more. In most cases, the net change in protective quality resulted from simultaneous improvements (or deteriorations) of the indices of protective quality. Hence, suitable management regimes in protection forests always have to be tailored to the current stand composition and the site-specific growth conditions, and there is no silver bullet for a suitable management approach (Moos et al., 2023). This is in line with the approach of NaiS (Frehner et al., 2005), where (a) target states are indicated, rather than prescribing management regimes, thus requiring local foresters to adjust the management interventions to the current stand characteristics, and (b) target states are differentiated by groups of site types, i.e., site conditions.

Regardless of the natural hazard considered, effective protection forests should feature a high canopy cover and contain only small gaps. Inevitably, many management interventions reduce the protective quality at least temporarily (Brang et al., 2006). Yet, in our study the recommended management regime sometimes differed based on whether the stand was protecting against avalanches or landslides/erosion/debris flow. The assessment criteria for these two natural hazards differ in the allowed gap size and the required canopy cover, albeit to a relatively small extent. Maroschek et al. (2015) reported similar findings in their simulation study, where the same sylvicultural treatment led to a better protective effect against avalanches than against landslides. The sensitivity of the protective quality to small variations in threshold values highlights that the target profiles can be quite restrictive for forest management in some cases.

The NaiS profiles (Frehner et al., 2005) are meant to describe sustainable (or even self-sustaining) stand compositions and structures that are close to a presumed natural state. Hence, stands fulfilling at least the minimal profile in all aspects, i.e., indices, should need little to no management interventions to remain in that state, especially under the assumption of a constant climate and the absence of major disturbances (Brang, 2001; Dorren et al., 2004). Suitable management regimes should, however, be able to improve the protective quality of stands that currently do not fulfil these prerequisites. Consequently, we had expected that not intervening would be a suitable management regime for well-structured stands in short-term simulations as well as in long-term simulations. This was true for the well-structured stands over 90 years across all elevational zones. However, for the long-term simulations, no management was among the recommended regimes only in the subalpine zone. The main reason for the clear improvement of the protective quality through suitable management regimes in the upper and high montane zones was the dominance of *Abies alba* in the absence of management, which will be discussed in detail further below.

As also expected, our results showed that the protective quality of young and mature stands could be mostly improved with management across elevational zones. In contrast, many studies have identified no management as the best strategy for protection forests across a large geographical range and varied stand conditions (Irauschek et al., 2017a, 2017b; Maroschek et al., 2015; Mina et al., 2017; Pardos et al., 2017; Scheidl et al., 2020). There however, the protective effect was measured without considering aspects of stand resistance and resilience and disturbances were not included in the simulations. We believe that incorporating characteristics such as regeneration, species composition, and diameter distribution in assessing a protection forests quality in simulation studies is crucial to derive realistic management recommendations for forestry practice. This is especially true when using simulation models that operate on a stand scale, in which disturbances are usually not explicitly depicted.

#### 4.2.2. Upper montane zone

In the upper montane zone, our results suggest that low-intensity interventions are best suited to increase or maintain the protective quality across all stand types. While long intervals of 30 to 40 years were ideal for young and well-structured stands (or no management at all in the latter case), shorter intervals are appropriate in mature stands. In the equilibrium of long-term simulations, the choice of interval does not seem to be decisive. Tree removals should be concentrated in larger gaps in young and well-structured stands, but in smaller gaps in mature stands and at equilibrium; this is in line with Rammer et al. (2015), who found a positive impact of management compared to NOM, especially when opening slit-shaped gaps.

Species composition is crucial in the simulation results for this zone. Species composition not matching the targets in most simulations is an important reason for the low protective quality achieved in UM compared to the other elevational zones. It is also the main reason for the surprisingly large positive effect of management in the long-term simulations. Over the course of roughly a century, the species composition is determined to a large extent by the initialization state, as replacing the entire stand within a short time is no viable option in protection forests due to the large gaps that would be created. However, in young stands we would have expected it to be possible to obtain a larger shift in species composition while maintaining a suitable stand structure. This could have been improved by giving more weight to a balanced species composition in the management settings as it was the least important weighing factor in the settings used across all management regimes. The effects of management in well-structured stands differed strongly depending on *site quality*, which can be attributed mainly to different species shares in the respective initialization stands. The preference for large gaps in young and well-structured stands results from a better species distribution among seedlings in larger gaps, which was not necessary to improve in mature stands as they already had a balanced species composition. The importance of species composition (mainly regarding stand resistance and resilience) in protective forests has been postulated earlier (Brang, 2001) and is corroborated by our findings.

However, our results revealed three methodological issues that are discussed in detail further below: (1) The dominance of *Abies alba* is surprisingly strong, both in very young stands and in the long term; (2) the species composition of the initialization stands should have been more similar between stands of the same type but different *site quality;* and (3) choosing different weighting factors to select trees to be removed by management based on the initial stand may have led to better protective qualities. Despite this, we deem the management recommendations to be realistic, as they lead to increased stand stability while maintaining a relatively closed canopy and improving or maintaining an already sustainable diameter distribution. With “better” species composition in all initialization stands, the management type would probably have been less important. However, many protection forests do not currently fulfill the minimal profile in that regard either, and hence our choice of initialization stands was not unrealistic.

#### 4.2.3. High montane zone

In the high montane zone, our results suggest that no management interventions are necessary over 90 years to maintain the protective quality in well-structured stands, which includes the mature stand with medium *site quality*, as there are no signs of natural decay throughout this period. Starting from a high protective quality and a sustainable stand structure with uninhibited regeneration, this corresponds to our expectations. Previous simulation of regions with comparable climatic characteristics (but including not only well-structured stands) have also reported NOM to achieve the highest protective effect against natural hazards under historic climate (Irauschek et al., 2017b, 2017a; Maroschek et al., 2015). However, and as discussed above, their results were probably influenced by the fact that no measures of resistance or resilience were incorporated in the assessment of the protective quality.

In young HM stands, practically any management regime led to an improvement of stand structure and stability. It is important to seize this opportunity, as in spruce-dominated forests stand stability can be improved substantially only by management in the young phase (Bebi et al., 2013; Stritih et al., 2021). The irrelevance of the choice of management type can be explained by there being little need to foster additional regeneration throughout most of the simulation time as the regeneration cover did not fall below the minimal thresholds in NaiS. However, creating groups (patches) of new regeneration should be started already at a young stand age in order to reach an uneven-aged stand structure in the long term (Bebi et al., 2013). Therefore, creating medium-sized openings (preferably slit-shaped) would probably be the best choice (Glanzmann et al., 2019; Lafond et al., 2014; Streit et al., 2009).

In naturally decaying stands, i.e., the mature stands with good *site quality*, low-intensity interventions at long intervals with removals concentrated on medium-sized gaps provide the best balance between gap size, diameter distribution and stand stability. Under similar conditions, Courbaud et al. (2001) found that group selection led to a more balanced stand structure than single-tree selection. High stand stability is quite important in these stands (but hard to improve by management), as they are prone to large biotic and abiotic disturbances (Ott et al., 1997). Yet, disturbances are not considered in ProForM, i.e., the stands “survive” for a long time even without management. In practice, however, this risk has to be taken into account and interventions should therefore be either more intense than suggested by our model in order to facilitate a faster transition to a younger stand, or not carried out at all in order not to further destabilize the system (Bebi et al., 2013; Stritih et al., 2021). Not intervening in decaying stands can have the additional benefit of an increasing number of lying logs that help to stabilize the snowpack and thereby lower the risk of avalanche release while simultaneously improving the growing conditions for regeneration in their vicinity. In contrast, weakened or recently fallen *Picea abies* trees are preferred host trees for bark beetles which increases the risk of further outbreaks in the remaining stand (Stadelmann et al., 2013).

One reason for the strong positive effect of management in the long-term simulations is, as already seen in UM, species-related. In the absence of management, *Abies alba* completely dominates the stand (cf. section “species imbalance” below), while management over a long time can substantially change the species shares. However, the recommended management regime with medium-sized gaps not only improves the species composition but also stand stability while maintaining a balanced diameter distribution; this is thus realistic.

#### 4.2.4. Subalpine zone

In the simulations of subalpine stands, canopy cover was unexpectedly low, to the point that even without management, the threshold of 50% to fulfill the requirements in avalanche protection forests was sometimes not reached. This also led to negative effects of management in LED protection forests, as the respective threshold of 40% canopy cover was usually fulfilled without management, but reduced below that level with interventions. This was most evident in well-structured stands. Another issue arising from the low canopy cover is that penalties for exceedingly large gap sizes were ineffective, as canopy cover was too low and thus the index for “horizontal arrangement”, incorporating both gap size and canopy cover, was already below the minimal profile. Another common feature of these simulations were unexpectedly short crowns and thus low stand stability, even in stands with many naturally occurring gaps (cf. section “low crown rates” below). Because the height of the crown base cannot be lowered in *Picea abies*, it is difficult to increase the mean crown length of the support trees except through management regimes with high intervention intensities and short return intervals. This led to the situation that these regimes were regarded as beneficial by the statistical models in mature stands and over the long term, even though the resulting stand structure with a very low basal area would in reality not provide a good protection against any natural hazard. These are clearly anomalous behaviors of ProForM.

Despite these shortcomings, the management recommendation of not intervening in well-structured stands and in the long-term on good *site quality* is realistic and corresponds to our expectations under the assumption that regeneration is not hindered. In young stands, the recommendation of not intervening or practicing single-tree selection at long intervals and low intensity are more questionable. While it might be the case that even after 90 years there is still sufficient regeneration to fulfill the minimal profile, there is a risk of producing a uniform stand in the longer term, and a management regime that improves vertical and horizontal stand structure would most likely be more beneficial. Bebi et al. (2013) stressed that interventions to enhance stand stability should take place in an early developmental stage when crowns still are long, as the risk of destabilizing the stand by an intervention increases with stand age. Guidelines for forest practitioners on the treatment of young mountain forest stands also suggest forming spatial clusters through one rather strong intervention (so called “Rottenpflege”, Glanzmann et al., 2019) where entire tree clusters are removed to induce new regeneration. In this case, a simulated management regime that either would not have had its first intervention directly at the beginning of the simulation or had not followed a regular schedule could have performed better. However, such regimes were not included in our simulations. Irregular management regimes might have also produced positive results in mature stands, where none of the simulated regimes led to an improvement compared to the naturally decreasing protective function without management. This illustrates the practical difficulties of managing mature and instable stands, as it is extremely difficult to increase their stability through interventions that bear the risk of further destabilizing the stand (Bachofen and Zingg, 2005; Bebi et al., 2013).

### 4.3. Methodological considerations

#### 4.3.1. Initialization stands

For this study, we needed a total of 48 initialization stands, most of which were supposed to represent “typical” situations of current mountain protection forests in Switzerland. A key challenge was that a “typical” stand is not clearly defined, and thus a selection based on expert judgement was necessary. Another challenge was that for different stand types – young, well-structured, and mature – initialization stands for two site and regeneration qualities each were required. We used “spin-up” simulations with the respective stand parameters and manually selected time steps for the initialization. The advantage of this approach was that any number of initialization stands could be created and that there was no risk of model artifacts (i.e., drift) at the start of the simulations due to, e.g., mismatching h/d ratios of trees or sudden mortality events due to locally exceedingly high tree densities. However, subjectively choosing stands that are similar across site and regeneration qualities in terms of their diameter distribution, abundance or lack of regeneration, and species composition was quite difficult, and identifying stands that are identical in at least some of these criteria was impossible. The simulation results showed a considerable dependency of the role of management on the initialization state. Due to the necessity of using non-identical initialization stands, a detected effect of, e.g., site quality, may actually have been due to a slightly different species composition of the initialization stands; this methodological problem was inherent to our approach.

There are a number of alternative methods that could have been used to generate initialization stands. One option would be to run a single spin-up simulation per stand type with a fixed set of parameters. This would ensure identical initial conditions for all simulations per site type. However, in reality stand characteristics such as basal area, stem numbers, or the share of regeneration differ in stands of high or medium site quality or with good or low regeneration quality; this would most likely have led to drift in the simulation due to these non-matching properties. Another option would be to run separate spin-up simulations but adjust and align key characteristics such as the species composition manually. However, many differences would still persist. A third option would be to use a stand generator, e.g., the structure generator STRUGEN (Pretzsch, 1997). Following the premise in the development of ProForM of using empirical data as far as possible, approaches based on inventory data such as the Swiss NFI (Mey et al., 2023, 2021) could be a viable method as well. However, further model development would be necessary to translate the output of a stand generator into the spatially explicit format required by ProForM. Finally, inventory data, e.g., from the Experimental Forest Management (EFM) network of WSL (Forrester et al., 2019) could be used, which however was not feasible for this study due to the large number of initialization stands required. All of these alternative methods for deriving initialization stands contain the risk of producing model artifacts. Yet, simulations with ProForM based on initialization data from EFM plots (Schmid et al., 2023) have not produced such effects and thus might be worth considering in future model applications.

#### 4.3.2. Management regimes

In order to conduct a systematic comparison of the impact of the management variables *type*, *interval* and *intensity* across different stand types using statistical models, we created a factorial set of management regimes that contained all possible combinations of the variable levels. To ensure comparability, all other model parameters relating to management were identical across stand types and elevational zones, i.e., time of the first intervention, minimum DBH of trees eligible for harvest as well as spatial buffer settings and weights of different stand characteristics to select the trees for harvesting that are specific to each management type. Hence, these management regimes are rather generic and not tailored to the individual stand and elevational zone. While this was the only way to compare the influence of the selected management variables across many stand types, two shortcomings became evident in the simulation results that should be addressed in future model applications. First, although interventions scheduled at regular intervals may be a viable option in well-structured stands and stands that are in a (pseudo-)equilibrium, young and mature stands often benefit from irregular interventions and/or interventions of varying intensities. Second, the weighting of stand characteristics that we used for selecting the trees to be removed reflected the management goals in well-structured stands and in the long-term: the primary focus was on removing large trees, the secondary focus on fostering existing regeneration patches, the tertiary focus was on not connecting existing gaps and removing trees with low crown lengths, and finally fostering an overall balanced species composition. However, in practice, weights may be chosen differently depending on specific stand attributes. For example, changing the species mixture in young stands would likely be the top priority, at least for the first two or three interventions. However, developing such nuanced management regimes for each initialization stand was beyond the scope of this study, as the possible variable combinations for a systematic comparison would have been too large to handle, and we prioritized maintaining comparability across site types.

#### 4.3.3. Quantifying stand protective quality

Determining a protection forest’s quality within the NaiS framework (Frehner et al., 2005) consists of assessing the current and estimating the future state of a stand in seven categories, six of which were implemented as indices in ProForM. For each index (and in some cases further subindices), two threshold levels are defined: the minimal and the ideal profile (MP and IP, respectively). The differentiation on how strongly a certain characteristic diverges from one of the threshold states is left to the practitioner’s judgement. This framework was challenging to translate into a numerical scale for a single index, and even more so at the aggregated level. Our approach was to use two aggregated numerical indices, one for the MP and one for the IP, which integrate the indices along time. While this lowered the level of complexity of the analysis considerably, challenges remained. First, using two response variables doubled the number of statistical models to be built and analyzed. Second, for arriving at a conclusion regarding an overall beneficial management regime, a prioritization has to be carried out by expert judgement in cases where a certain management regime has a positive impact on one but a negative impact on the other target variable. Third, in such an aggregation, stand characteristics that deviate strongly from the MP and that would be prioritized over other stand characteristics in any practical application, such as an extremely low basal area, had the same negative weight in our scheme as if they did not fulfill the MP by a slight margin.

In preliminary analyses, we tested an alternative response variable. Instead of calculating the share of individual indices meeting either profile over time, we calculated the share of years in which either profile was fulfilled by every index. However, in simulations of the elevational zones UM and SA, even the MP was often not fully fulfilled. Hence, this variable could not be used to produce meaningful statistical analyses of the underlying factors. To weigh indices with a very low index value properly, metrics based on distances to the profile threshold could be applied. However, the lower end of the scale would have to be defined clearly, which it is not the case in the NaiS framework. A further aggregation into a single “super-index” describing the protective quality in an integral way would not be a solution, but would pose additional difficulties. One example would be how to avoid one index at the ideal state compensating for the lack of reaching the minimal profile of another index. A stand that fulfills the minimal profile across all indices should be valued more highly than one that fulfills the ideal profile in half but fails to fulfill the minimal profile in the other half of the indices.

In previous studies that used a valuation of the protective function based (at least to some extent) on NaiS, two main approaches were taken. The first approach is using categorical quality descriptions for either one or multiple stand characteristics but, in the latter case, without aggregation (e.g., Cordonnier et al., 2008; Irauschek et al., 2017b; Wehrli et al., 2007b). The second approach is to create numerical indices integrating multiple stand characteristics such as stem densities in different diameter classes, average DBH, canopy cover, leaf area index, or the share of coniferous species (Briner et al., 2013; Elkin et al., 2013; Lafond et al., 2017; Mina et al., 2017; Seidl et al., 2019). Most of these indices focused on stand qualities required for the current protective effect against a certain natural hazard, but none of them included aspects of resistance or resilience. These challenges reinforce the approach taken in NaiS to refrain from a single numerical index for describing a protection forest’s quality, as the human brain is more capable of prioritizing and evaluating trade-offs between individual stand characteristics to arrive at an overall impression, compared to a computer following a strict set of quantitative rules.

With the deficiencies remaining in our target variables, a strictly model-driven optimization of management regimes was not possible. Instead, we derived management recommendations based on the simultaneous interpretation of the statistical model’s outcomes and selected underlying simulation results. With a single and more reliable target variable, management regimes could be iteratively optimized, or the best management regime could be identified automatically from a larger number of predefined simulations, as done for other ecosystem services (e.g., Altamirano-Fernández et al., 2023; Buongiorno et al., 2012; Díaz-Yáñez et al., 2021; Foppert and Maker, 2024). Such techniques would also allow for a larger set of varying management parameters to be investigated.

### 4.4. Needs for further research

ProForM was calibrated and validated with long time series of repeated inventories (Schmid et al., 2023). Simulations of stands in the three elevational zones UM, HM, and SA suggested a high accuracy for rendering key stand attributes such as basal area, stem numbers, diameter distributions, and species composition. In the submontane elevational zone (SM), the development of the diameter distribution and the species composition could not be tracked accurately enough, which, together with difficulties in the spin-up simulations, was the main reason for not including this elevational zone in the current study. In the updated model version 1.1 (Schmid et al., 2024), the algorithms for forest dynamics were not modified.

Here, we used ProForM for two purposes that it was not specifically designed nor validated for: (1) simulations from bare ground in spin-up simulations, and (2) long-term simulations without management, reaching an equilibrium state of the forest. While it is not necessarily a goal to accurately portray the process of reforestation or the potential natural vegetation (PNV) with ProForM, the deficiencies revealed above have implications also for short-term simulations of managed stands and should be addressed in the future, as discussed below.

#### 4.4.1. Species imbalance

In long-term simulations without management in the elevational zones UM and HM, *Abies alba* came to dominance in terms of basal area. This results from the combination of two factors: *Abies alba* has the highest maximum diameter growth rate and, more importantly, the lowest mortality rate (Fig. 25). With germination probabilities in the model depending on the current species shares in the stand, this leads to a positive feedback and thus over time to almost monospecific stands. Truly validating PNV simulations is not possible, as no empirical data are available (Bugmann, 1996). However, likely models of the climax vegetation as proposed by Ellenberg (1986) or in NaiS (Frehner et al., 2005; Frey et al., 2021) suggest that pure *Abies alba* stands are unrealistic in both UM and HM. To calibrate and validate ProForM to more accurately portray species composition in the long term, model comparisons with, e.g., ForClim (Huber et al., 2020) could be used. Maximum diameter growth rates and empirical mortality functions (Hülsmann et al., 2018) would have to be calibrated simultaneously (c.f. Cailleret et al., 2020).

The height growth functions of regeneration that were derived from empirical data exhibited considerable and, unfortunately, unrealistic differences between coniferous and deciduous species (Fig. 25a). In ProForM, the coniferous species clearly outperform the broadleaved ones. Combined with the higher slenderness ratio of the broadleaved species, reaching a DBH of 12 cm only at much larger tree heights than the conifers and therefore being treated as “regeneration” for a longer time in ProForM, this leads to a strong imbalance of competition, strongly favoring conifers in the early simulation when initializing from bare ground. This effect was clearly visible in preliminary spin-up simulations in the (later discarded) elevational zone SM as well as for UM, where management interventions removing most conifers had to be included to obtain young initialization stands that had a substantial share of broad-leaved species. The regression models used to simulate regeneration height growth of *Fagus sylvatica* and *Acer pseudoplatanus* in ProForM exhibited higher errors and lower R^2^ and were based on fewer observations than those for *Picea abies* and *Abies alba* (Schmid et al., 2023). Prior to further model applications, these growth functions must be re-evaluated and improved.

#### 4.4.2. Low canopy cover

Long-term unmanaged simulations in HM and SA led to very low canopy cover in equilibrium states with values of ca. 45 to 55% in HM (medium and good site quality, respectively) and 38 to 42% in SA. Canopy cover is calculated in two steps in ProForM. First, the share of cells containing trees of at least 12 cm DBH is calculated. Then, these values are reduced for elevational zones HM and SA to 90% and 75%, respectively (no reduction in UM), based on the assumption that one large tree occupying one cell does not cover the entire cell area with its canopy. This reduction might be too strong, as cells that are populated by multiple smaller individuals may well be filled completely by tree crowns. However, even a smaller reduction would yield canopy covers of less than 60% in HM and 50% in SA, which we still consider too low.

A further contributing factor to the low canopy cover could be an unrealistic distribution of basal area across the diameter classes. The total basal area in equilibrium of 60 to 75 m^2^/ha in HM (medium and good site quality, respectively) and 40 to 55 m^2^/ha in SA seems to be realistic when compared to data from managed stands of HM and SA (Bachofen and Zingg, 2005; Forrester et al., 2021). However, in ProForM there are multiple large trees at the maximum DBH that live for several hundred years (cf. Fig. 25c, especially in SA). These trees strongly reduce growth in the adjacent cells through the basal-area-based competition indices, which tend to lead to gaps remaining open for a long time. The fact that some areas of several 100 m^2^ in unmanaged long-term simulations remain “gaps” throughout the entire assessment period of 100 years supports this hypothesis. The fact that many individuals survive for a long time at the maximum diameter, at which the mortality probability is at its maximum due to no further diameter growth, again points at an underestimation of the mortality rate, especially for *Abies alba* that dominates LT simulations in HM (cf. section “species imbalance”). However, due to their relatively complex structure compared to other model functions in ProFroM, adjusting or calibrating the empirical mortality functions (Hülsmann et al., 2018) would be difficult using the procedure employed in the calibration of ProForM v1.0 (Schmid et al., 2023). Combined with the problem that there is little to no measured data available on tree mortality that have not been used in developing these functions, an automated calibration does not seem feasible either. A possible solution might be to modify the mortality probability with a newly introduced parameter.

Lastly, canopy cover was not among the emerging patterns validated in Schmid et al. (Schmid et al., 2023), as there were no data for the stands used. Estimates of canopy cover could be inferred from historical aerial photographs for at least some of the inventories and used for the calibration and validation of the suggested new parameter. It is questionable, however, whether these time series would yield conclusive results, as all these stands were managed and canopy cover thus mostly determined by management and not by natural mortality. Calibration would thus have to rely either on PNV descriptions or outputs of other models (cf. section “species imbalance”).

#### 4.4.3. Low crown ratios

In unmanaged long-term simulations of the subalpine zone, the crown ratios of the 100 dominant trees per ha were surprisingly low (20-40%), even though canopy cover was low, as discussed above. On long-term monitoring plots of managed subalpine stands, crown ratios of 42 to 77% were reported (Bachofen and Zingg, 2005), clearly exceeding our results. This may result from locally high competition index values due to several trees at a very high or even the maximum DBH being present. One would expect these large trees to have large crowns, as they probably dominated locally. However, these trees have crown ratios even below the stand average. This suggests that the purely theory-based algorithm for calculating the change in height to crown base might have calibration deficits in SA, as only one stand was used for calibration and one for validation, both being plantations (Schmid et al., 2023) and thus not containing data on large and old trees. A recalibration of this algorithm with additional inventory data would certainly help to resolve this issue.

#### 4.4.4. Assessment of protective function: the gap challenge

One of the main challenges in the assessment of the protective function in ProForM is the identification of gaps. In model version 1.0 (Schmid et al., 2023), gaps were defined as continuous areas of cells with trees of DBH ≤ 12 cm. In version 1.1, an additional rule was introduced, i.e., that only groups of three adjacent cells fulfilling this criterion could form part of a gap, leading to a more realistic gap detection. Further improvements to the gap detection algorithm may be possible, e.g., by implementing a moving window approach searching for gaps of different shapes and sizes, as done in PICUS (Irauschek and Lexer, 2018; Maroschek et al., 2015), thus eliminating the identification of oddly shaped areas as gaps.

A further change from ProForM v1.0 to v1.1 was the relaxation of the threshold for fulfilling the minimal profile, allowing for one gap to exceed the size limits stipulated by NaiS (Frehner et al., 2005). Unfortunately, this pragmatic solution for a previously systematic underestimation of the protective quality led to unrealistic results in some SA simulations with very low canopy cover. There, most “gap cells” ended up being connected and were thus categorized as one extremely large gap. However, due to the relaxed assessment rule, these stands still fulfilled the minimal profile in terms of gap length and width. Such situations could be avoided by tightening the new rule to, e.g., allowing one gap exceeding the minimal profile, but only by 50%; or by abolishing this relaxation altogether.

## 5. Conclusions

Our study has shown the pivotal importance of regeneration for the short- and long-term protective quality of mountain forests. In the extreme case in which the already slow regeneration growth in mountain forests is strongly reduced by browsing and in which some of the key tree species cannot regenerate successfully, the protective quality is seriously deteriorated and there is no possibility to improve it through forest management apart from highly expensive measures such as planting and subsequently protecting the regeneration: this is a “dead end” for sylviculture. This underlines the need for keeping ungulate populations to a level where tree regeneration of all species required for warranting the protection function is not significantly hindered.

For stands with a better regeneration growth, we derived management recommendations to improve or sustainably maintain the protective quality. We found general patterns between forest types, i.e., elevational zones. In the mixed forests of the upper montane zone, interventions with a low intensity are generally preferred. In the spruce-fir-forests of the high montane zone, a wide range of intervals and intensities can be applied as long as trees are removed in groups. In subalpine spruce-forests, either interventions at long intervals are suitable, or no interventions are necessary at all.

Even more than the forest types, the initial stand conditions are decisive for the management recommendations. In the absence of large disturbances, well-structured stands with a good protective quality do not need to be managed actively across all three elevational zones. However, many current mountain forest stands originate from spontaneous re-forestation or homogenous planting activities in the late 19^th^ and early 20^th^ centuries and are thus not well-structured. Our results therefore do not imply that management of current mountain protection forests in the European Alps would not be necessary altogether. Our simulations showed that the protective quality of young as well as mature stands can be enhanced considerably by management in most cases. While the specific recommendations differ by stand type and elevational zone, generally, return intervals of 30 to 40 years and removals of only 10 to 20% of the stand basal area are deemed sufficient to improve or maintain the protective quality.

Further research needs with regards to ProForM were identified, with the most pressing being the species-specific improvement of the growth and mortality rates which currently lead to an unrealistic dominance of conifers, especially *Abies alba*. We suggest to scrutinize the regeneration growth submodel and to conduct an additional model “validation” with long-term simulations, using either descriptions of potential natural vegetation or outputs from other simulation models as benchmarks. We assume that improving the mortality submodel will also improve the current underestimation of canopy cover in the high montane and subalpine zone. Finally, the submodel for calculating crown length should be recalibrated for the subalpine zone with additional inventory data.

## CRediT authorship contribution statement

**Ueli Schmid**: Conceptualization, Methodology, Software, Formal analysis, Investigation, Writing - Original Draft, Visualization. **Monika Frehner**: Conceptualization, Methodology, Writing - Review & Editing, Supervision, Funding acquisition. **Harald Bugmann**: Conceptualization, Methodology, Resources, Writing - Review & Editing, Supervision, Funding acquisition.

## Acknowledgements

We thank Christoph Bigler for his support with the statistical analysis. The Swiss Federal Office for the Environment FOEN generously supplied funding for this study under contract number 00.0186.PZ/Q041–2010.

## List of abbreviations

A: natural hazard *snow avalanches*
BRT: Boosted Regression Trees
CAB: management type *cable yarding*
DBH: diameter at breast height
ES: ecosystem service
GRS: management type *group selection*
HM: elevational zone *high montane*
IP: *ideal profile*, thresholds for protective quality assessment
LED: natural hazard landslides, erosion, debris flow
LT: simulation type *long term*
NaiS: framework for assessing the protective quality in Swiss forestry practice
NOM: management regime *no management*
MP: *minimal profile*, thresholds for protective quality assessment
SA: elevational zone *subalpine*
SC: management type *slit cuts*
SM: elevational zone *submontane*
sIP_abs_: target variable *share of indices meeting the ideal profile* (protective quality)
sIP_diff_: target variable *difference between* sIP_abs_ of simulation with management and the corresponding NOM simulation (impact of management on protective quality)
sMP_abs_: target variable share of indices meeting the minimal profile (protective quality)
sMP_diff_: target variable difference between sMP_abs_ of simulation with management and the corresponding NOM simulation (impact of management on protective quality)
ST: simulation type *short term*
STS: management type *single tree selection*
UM: elevational zone *upper montane*
ΔPQ: *difference in protective quality*, either *sMP_diff_* or *sIP_diff_* (impact of management on protective quality)

