## Supplementary material S1 for "Management recommendations for Alpine protection forests: the importance of regeneration quality and initial stand composition"

### Supplementary material S1: Description of the ProForM model v1.1

**Note:** The model description for ProForM v1.0 can be found in the Supplementary material S1 of Schmid et al. (2023). It follows the ODD (Overview, Design concepts, Details) protocol for describing individual- and agent-based models (Grimm et al., 2006), as updated by Grimm et al. (2020). The present document uses the same structure, but contains text only in those sections that experienced changes and additions between ProForM v1.0 and v1.1. If not stated otherwise, the sections of this document are either additions or replace the ones in the previous model documentation.

The computer code of the model is available on github (Schmid et al., 2024).

### 1. Purpose and patterns

*No changes*

### 2. Entities, state variables, and scales

*No changes*

### 3. Process overview and scheduling

The first process of **forest dynamics and management** remains unchanged between v1.0 and v1.1, except for the addition of new management algorithms to determine the removal of trees in management intervention $dN_{mgm}$ (step 5 in the schedule; cf. management sub-models).

The **assessment of the protective quality of the simulated stand** remains essentially the same in v1.1 as in v1.0 only that the process is divided now into two parts – preparation (i.e., gap recognition) and assessment (index calculation) – that can be run separately. In the up-dated schedule below, steps 3 and 4 of the assessment, i.e., the actual comparison of stand properties to the requirements of the profile, are repeated in a loop for each time step.

*Part 1 – preparation*

1. Load simulation outputs
2. Calculate hexagon metrics height and size (as in submodel init Simulation grid)
3. Restrict simulation outputs to state variables 1 (spring)
4. Create subset of simulation results with largest cohort (largest *D*) per cell
5. Identify gaps and calculate area, length, and width (submodel nais Gaps)
6. Aggregate and save outputs

*Part 2 – assessment*

1. Load preparation outputs
2. Load site- and natural-hazard-specific profiles (submodel nais Profiles)
3. Calculate stand characteristics and indices
    a. Mixture (submodel nais Mixture)
    b. Vertical structure (submodel nais Vertical structure)
    c. Horizontal arrangement (submodel nais Horizontal arrangement)
    d. Support trees (submodel nais Support trees)
    e. Seedlings (submodel nais Seedlings)
    f. Saplings & thicket (submodel nais Saplings and thicket)
4. Aggregate and save outputs

### 4. Design concepts

*No changes*

#### 5. Initialization

*No changes*

#### 6. Input data

The only input data that externally drive model dynamics throughout a simulation are the management instructions.

For the management algorithms **RDC tree** (submodel mgm RDC tree) and **RDC cohort** (submodel mgm RDC cohort), the instructions consist of a table defining the time steps in which interventions take place ($step_{mgm}$), the minimal diameter (*Dmin*) for each intervention and species ($Spc_{mgm}$), and the share of basal area to be removed per species ($BAshare_i$) from each diameter class (*Dclass*).

For the management algorithms **STS** (single tree selection, submodel mgm STS), **GRS** (group selection, submodel mgm GRS), **SC** (slit cuts, submodel mgm SC), and **CAB** (cable yarding, submodel mgm CAB), the instructions are similar. They consist of a table defining the management settings in four categories: (1) timing and intensity of interventions, (2) minimum species shares to be maintained, (3) spatial layout of harvest units, and (4) weighting factors for tree selection. If not stated otherwise below, the settings apply to all four management algorithms. For the detailed meaning and impact of the settings, refer to the respective management submodel.

1. Category "timing and intensity of interventions"
    a. time step of the first intervention ($interv_{first}$)
    b. interval between interventions in years ($interv_{interval}$)
    c. share of total basal area reduction per intervention ($interv_{BAred}$)
    d. minimal DBH in cm of trees eligible for harvest($interv_{Dmin}$)
2. Category "minimum species shares to be maintained"
    a. Share of stand basal area per species at the time of management, below which they are not eligible for harvest ($BAmin_i$)
3. Category "spatial layout of harvest units"
    a. Number of cells to be aggregated to groups ($mgmunit_{agg}$, only in GRS)
    b. Slit length in cells ($mgmunit_{SL}$, only in SC and CAB)
    c. Slit width in cells ($mgmunit_{SW}$, only in SC and CAB)
    d. Up to three grid columns containing cable lines ($mgmunit_{CL1-3}$, only in CAB)
    e. Initial buffer width in number of cells to be left unmanaged around interventions ($mgmunit_{buff}$)

4. Category "weighting factors for tree selection"
    a. Weight given to tree species (*mgmw*$_{spc}$, not in CAB)
    b. Weight given to basal area (*mgmw*$_{BA}$)
    c. Weight given to crown ratio (*mgmw*$_{CR}$, not in CAB)
    d. Weight given to regeneration within harvest unit (*mgmw*$_{RegUnit}$, not in STS)
    e. Weight given to regeneration in cells neighboring harvest unit (*mgmw*$_{RegNeighb}$, not in STS)

### 7. Submodels

#### Data

*No changes*

#### Submodel group "initialization"

##### init Calculate parameters

This submodel remains largely identical between v1.0 and v1.1. The only changes concern the determination and addition of management parameters from the management instruction to the final list of parameters due to the newly added management algorithms. The list below replaces the one bullet point in the original model documentation concerned with management parameters.

- Determination and addition of management parameters to the final list of parameters, depending on the selected management type (*mgm*$_{type}$)
    - If *mgm*$_{type}$ is either RDC tree or RDC cohort:
        - Time steps in which a management intervention take place (*step*$_{mgm}$), specified in the management instructions.
    - If *mgm*$_{type}$ is either STS, GRS, SC, or CAB:
        - Determination of time steps in which management interventions take place (*step*$_{mgm}$), based on *interv*$_{first}$ and *interv*$_{interval}$ (specified in the management instructions) and the total simulation time (specified in the simulation settings).
        - Addition of remaining management instructions to the final list of parameters.

##### init Simulation grid

*No changes*

##### init Neighbors

*No changes*

##### init Data structure

*No changes*

#### Submodel group "auxiliary variables"

##### aux Stage

*No changes*

##### aux Basal area

*No changes*

##### aux Volume

*No changes*

##### aux Crown ratio

*No changes*

##### aux CI height

*No changes*

##### aux CI diameter

*No changes*

##### aux CI mortality

*No changes*

##### aux CI crown length

*No changes*

##### aux Species selection

*No changes*

#### Submodel group "forest dynamics"

##### dyn Height growth regeneration

*No changes*

##### dyn Height adult trees

*No changes*

##### dyn Height growth all

*No changes*

##### dyn Diameter regeneration

*No changes*

##### dyn Diameter growth adult trees

*No changes*

##### dyn Diameter growth all

*No changes*

##### dyn Ingrowth

*No changes*

##### dyn Mortality

*No changes*

##### dyn Crown base change

*No changes*

##### flag Germination

*No changes*

##### flag Cohort death

*No changes*

##### flag Advance regeneration

*No changes*

#### Submodel group “update state variables”

##### update Species

*No changes*

##### update Stem number

*No changes*

##### update Diameter

*No changes*

##### update Height

*No changes*

##### update HCB

*No changes*

#### Submodel group “management”

##### mgm RDC tree

*No changes*

##### mgm RDC cohort

*No changes*

#### Common features of management submodels STS, GRS, SC, and CAB

The management submodels STS, GRS, SC, and CAB, all new in v1.1, share two basic mechanisms, which are implemented slightly differently in each submodel: the ranking of management units and a spatial buffer between selected management units.

##### Ranking of management units

All four submodels work with a list of management units (trees, cells, or groups of cells), which is based on the model's list of cohorts with their current state and auxiliary variables. In a first step, all cohorts that are not eligible for harvest are removed from the list. This can be a) due to their tree species, if the species' current share of basal area is at or below the minimum species share $BAmin_i$ (defined in the management settings), or b) due to the tree's DBH, if it is at or below the minimum diameter $interv_{Dmin}$ (defined in the management settings).

In a second step, depending on the management algorithm, the list of cohorts is organized in management units. For each unit, a selection probability $P_{sel}$ is calculated, and the list of management units is ranked in descending order of $P_{sel}$. The selection probability is calculated according to Eq. S1.1 and is based on the Uneven-Aged Management Algorithm developed for the model Samsara2 (Lafond et al., 2014).

$$P_{sel} = \prod_{y} {P_y}^{mgmw_y} \cdot rn \qquad (S1.1)$$

with $P_y$ being the selection weight probability of criterion $y$, $mgmw_y$ its associated weight defined in the management settings, and $rn$ a pseudo random number of uniform distribution between 0 and 1 assigned to each management unit. Depending on the selected management type, different criteria are evaluated. The value of the weights can be chosen freely by the user, but the following values are recommended: 0, if the criterion should be irrelevant; 1, to use it as an orientation (equal weight as random number); 10, to place a focus on it; and 100, to prioritize this criterion.

The selection weight probabilities $P_y$ are calculated for each management unit. The species proportion selection weight probability $P_{spc}$ is calculated as the current basal area share of species $i$ over the maximum value of the basal area shares of all four species (Eq. S1.2). If management units contain multiple species, $P_{spc,i}$ of the most abundant species in terms of basal area within the unit is assigned to the entire unit. Together with $mgmw_{spc}$, the partial selection probability based on the unit's species is calculated with a large selection probability assigned to units of abundant species, and a large weight leading to a more balanced species composition after management.

$$P_{spc,i} = \frac{BAcurr_i}{\max(BAcurr)} \qquad (S1.2)$$

The basal area selection weight probability $P_{BA}$ is calculated as the management unit's basal area over the maximum unit basal area (Eq. S1.3). Together with $mgmw_{BA}$, the partial selection probability based on the unit's basal area is calculated with a large selection probability assigned to units with large basal area, and a large weight leading to more aggregated and less spread-out tree removals.

$$P_{BA,i} = \frac{BA_{unit,i}}{\max(BA_{unit})} \qquad (S1.3)$$

The crown ratio selection weight probability $P_{CR}$ depends on the management unit's crown ratios, or rather their stem ratios (i.e., 1-CR). It is calculated as the mean stem ratio (average weighted by basal area) over the maximum stem ratio among the units (Eq. S1.4). Together with $mgmw_{CR}$, the partial selection probability based on the unit's crown ratio is calculated with a large selection probability assigned to units with low crown ratios, and a large weight leading to a remaining stand with a higher average crown ratio after management.

$$P_{CR} = \frac{\left(1 - CR_{unit,i}\right)}{\max(1 - CR_{unit})} \qquad (S1.4)$$

The selection weight probability for regeneration within the unit $P_{RegUnit}$ depends on the abundance of regeneration within the management unit. It is calculated as the unit's share of cohorts containing regeneration (stages 1-3) over the maximum value of this criterion among units (Eq. S1.5). Together with $mgmw_{RegUnit}$, the partial selection probability based on the unit's regeneration share is calculated with a large selection probability assigned to units with high shares of regeneration, and a large weight leading to more removal of units containing advance regeneration and thus fostering it.

$$P_{RegUnit} = \frac{s_{regu,i}}{\max\left(s_{regu}\right)} \qquad (S1.5a)$$

$$s_{regu} = \frac{ncoh_{reg}}{ncoh_{tot}} \qquad (S1.5b)$$

with $s_{regu}$ being the share of cohorts within a management unit containing regeneration, $ncoh_{reg}$ the number of cohorts with regeneration, and $ncou_{tot}$ the total number of cohorts per unit.

The selection weight probability for regeneration in cells neighboring the harvest unit $P_{Reg\text{-}Neighb}$ depends on the abundance of regeneration in the unit's immediate neighborhood, i.e., its adjacent cells. It is calculated as the neighbor's share of cells stocked solely with regeneration (only cohorts in stages 1-3) over the maximum value of this criterion among units (Eq. S1.6). Together with $mgmw_{RegNeighb}$, the partial selection probability based on the neighbors's regeneration share is calculated with a large selection probability assigned to units with small shares of regeneration among its neighbors, and a large weight leading to more removal of units surrounded mostly by adult trees and thus less enlargements of existing gaps.

$$P_{RegUnit} = \frac{(1 - s_{regn,i})}{\max(1 - s_{regn})} \quad (S1.6a)$$

$$s_{regn} = \frac{ncells_{reg}}{ncells_{tot}} \quad (S1.6b)$$

with $s_{regn}$ being the share of cells among the management unit's immediate neighbors containing only regeneration, $ncells_{reg}$ the number of cells containing only regeneration among those neighbors, and $ncells_{tot}$ the total number of neighboring cells.

**Spatial buffer**

In the management settings, the user can define a buffer value $mgmunit_{buff}$ with possible values of 0, 1, or 2. The buffer represents the width (in number of cells) of the ring around management units already selected for harvest that is initially exempt from harvest. If the target basal area cannot be reached this way, the actual buffer value is reduced by one and the unit selection procedure is repeated. Large buffer values are meant to avoid the spatial clumping of harvested units and therefore the formation of overly large gaps. In the CAB algorithm, the buffer value is reduced in steps of 0.5, see submodel mgm CAB for details.

mgm STS

This submodel (new to v1.1) is called if a management intervention is scheduled for the current time step and the management type "STS" (single tree selection) is specified in the simulation settings. It determines the number of trees per cohort to be removed through management ($dN_{mgm}$). It uses the management instructions passed to the model as input data (see section Input data) and a partially randomized process to determine how many, if any, trees per cohort should be removed.

The submodel takes the current list of cohort state information (cf. submodel init Data structure) and the management instructions as inputs. The output is the list of cohort state information with updated values for the number of trees to be harvested ($dN_{mgm}$). To rank the list of units for harvest, the species, basal area, and crown ratio are taken into account. As only single trees are harvested, the criteria for regeneration within and around the harvested individual cannot be applied.

**STS algorithm**

1. List of all cohorts with their current state and auxiliary variables is retrieved
2. Current species shares and derived values are calculated.
    a. Calculation of the basal area share of all species of the current state 2 (*$BAcurr_i$*).
    b. Comparison to the minimum species shares (*$BAmin_i$*) and decision, whether species is eligible for harvest (eligible if current basal area share > *$BAmin_i$*).
    c. Calculation of the species proportion selection weight probability* ($P_{spc,i}$).
3. Cohorts exempt from harvest are removed from list of cohorts to get list of potential harvest cohorts.
    a. Removal of all cohorts of tree species not eligible for harvest based on the minimum basal area criterion.
    b. Removal of all cohorts that do not fulfill the minimum diameter criterion (*$interv_{Dmin}$*).
    c. Stop algorithm, if no cohorts are eligible for harvest and return $dN_{mgm}$ = 0 for all cohorts.
4. Selection probabilities are calculated and selection list is ranked.
    a. Calculation of selection weight probabilities ($P_{BA}$ *and* $P_{CR}$) per tree*.
    b. Turn list of potential harvest cohorts into list of potential harvest trees
    c. Calculate uniformly distributed random number *rn* between 0 and 1 for each tree on the list.
    d. Calculate selection probability $P_{sel}$ for each tree*.
    e. Order list by descending selection probability $P_{sel}$ and add rank corresponding to it.
5. Harvest operation is prepared.
    a. Addition of column “harvested” to each tree on the selection list, initially set to FALSE.
    b. Calculation of target basal area $BA_{target}$ as the product of current total stand basal area (including trees not eligible for harvest) and the desired share of basal area reduction $interv_{BAred}$.
    c. Initialize the buffer state ($S_{buff}$) to the desired initial value $mgmunit_{buff}$.
    d. Initialize empty vector for collecting cells on which a tree has been selected for harvest.
    e. Initialize empty vector for collecting cells within buffer ring 1 (immediate neighbors of cells on which a tree has been selected for harvest).
    f. Initialize vector for collecting cumulated basal area of trees selected for harvest with initial value 0 and Boolean ($TBA_{reached}$) indicating if $BA_{target}$ has been reached as FALSE.

6. Harvest operation is carried out in a while-loop with the condition of $S_{buff} \geq 0$.
    a. Select trees to be harvested in for-loop through the ranked selection list .
        i. If tree has already been selected for harvest, jump to next list entry.
        ii. If tree lies within buffer zone of an already harvested tree, jump to next list entry.
            1. With $S_{buff}$ = 2, the buffer zone consists of all cells with trees already selected for harvest plus those in buffer ring 1 around them.
            2. With $S_{buff}$ = 1, the buffer zone consists of all cells with trees already selected for harvest.
            3. With $S_{buff}$ = 0, there is no buffer zone and multiple trees on one cell can be harvested.
        iii. Mark current tree to be selected for harvest.
        iv. Add current cell to vector of cells with trees having been selected for harvest.
        v. Add current cell's immediate neighbors to vector of cells within buffer ring 1.
        vi. Add updated cumulated basal area of trees selected for harvest to collection vector.
        vii. If current cumulated basal area of trees selected for harvest exceeds $BA_{target}$, break for-loop.
    b. If current cumulated basal area of trees selected for harvest exceeds $BA_{target}$, break while-loop.
    c. Reduce $S_{buff}$ by 1.
7. Decision whether last selected tree for harvest should be retained in harvest list.
    a. Calculate absolute difference between last two entries of vector with cumulated harvested basal area and $BA_{target}$.
    b. If difference is smaller for second to last entry, remove last tree to be selected from harvest.

*For the calculation of the weights, cf. section Common features of management submodels STS, GRS, SC, and CAB.

The information on the number of trees harvested per cohort is then assembled and linked to the current list of cohorts, updating the number of trees to be harvested ($dN_{mgm}$).

##### mgm GRS

This submodel (new to v1.1) is called if a management intervention is scheduled for the current time step and the management type "GRS" (group selection) is specified in the simulation settings. It determines the number of trees per cohort to be removed through management ($dN_{mgm}$). It uses the management instructions passed to the model as input data (see section Input data) and a partially randomized process to determine how many, if any, trees per cohort should be removed.

The submodel takes the current list of cohort state information (cf. submodel init Data structure), the spatial cell information (cf. submodel init Simulation grid and submodel init Neighbors) and the management instructions as inputs. The output is the list of cohort state information with updated values for the number of trees to be harvested ($dN_{mgm}$). It consists of two algorithms: the grouping and the GRS algorithm. The grouping algorithm identifies all possible management units, i.e., the groups, and is only run once, at the time step of the first management intervention. The GRS algorithm defines which units are harvested and is called at each intervention time step. To rank the list of groups for harvest, all criteria described in the ranking process are used.

##### Grouping algorithm

The grouping algorithm returns a list of all theoretically possible management units, i.e., groups (including overlapping groups). The list contains information on the cells and cohorts included in the group and their respective neighboring cells in ring 1 (immediately surrounding the group cells) and ring 2 (immediately surrounding cells of ring 1). Group size ($mgmunit_{agg}$) can be defined as 1, 2, 3, or 4 cells in the management settings with cell areas equaling approximately the canopy area of one large tree (39 – 100 m$^2$, depending on the selected *stratum*). Groups of size one correspond to the individual cells in the simulation grid. Groups of two and of four cells can have three different shapes, groups of three cells can have two different shapes (Fig. S1.1). The number of potential management units is thus a multiple of the number of cells in the stand with the number of possible shapes.

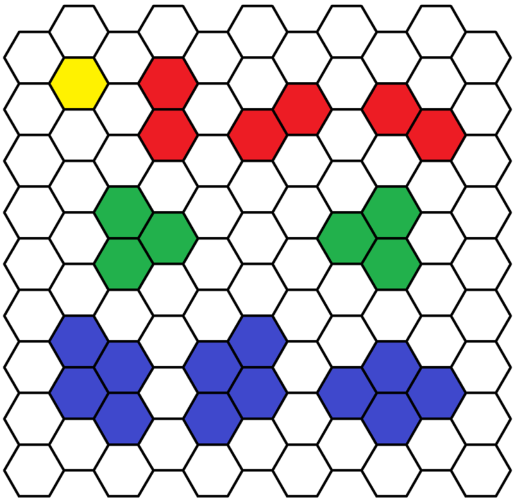

***Fig. S1.1*** *All spatial arrangements of groups returned by the grouping algorithm of the submodel mgm GRS depending on the selected group size (identified by color).*

**GRS algorithm**

1. List of all cohorts with their current state and auxiliary variables is retrieved
2. Current species shares and derived values are calculated for each cohort.
    a. Calculation of the basal area share of all species of the current state 2 ($BAcurr_i$).
    b. Comparison to the minimum species shares ($BAmin_i$) and decision, whether species is eligible for harvest (eligible if current basal area share > $BAmin_i$).
    c. Calculation of the species proportion selection weight probability* ($P_{spc,i}$).
3. Cohorts exempt from harvest are first identified and then removed from list of co-horts to get list of potential harvest cohorts.
    a. All cohorts of tree species that do not fulfill the minimum basal area criterion.
    b. All cohorts that do not fulfill the minimum diameter criterion ($interv_{Dmin}$).
    c. Stop algorithm, if number of cohorts not eligible for harvest equals the total number of cohorts in the stand and return $dN_{mgm}$ = 0 for all cohorts.
    d. Remove previously identified cohorts from list of management units.
4. Characterization of management units
    a. Retrieve state and auxiliary variables of cohorts, organized by management unit.
    b. Identify dominant tree species per unit as the one with the largest share of basal area
    c. Calculate unit's total basal area
    d. Calculate unit's mean crown ratio as a mean across cohorts, weighted by ba-sal area.
    e. Calculate share of regeneration (cohorts in stages 1-3) within each unit..
    f. Calculate share of regeneration (cells with all cohorts in stages 1-3) among neighbors of ring 1 of each unit.
5. Selection probabilities are calculated and selection list is ranked.
    a. Calculation selection weight probabilities ($P_{BA}$, $P_{CR}$, $P_{RegUnit}$, *and* $P_{RegNeighb}$) and random number *rn* per unit*.
    b. Calculate selection probability $P_{sel}$ for each unit*.
    c. Order list by descending selection probability $P_{sel}$ and add rank corresponding to it.

6. Harvest operation is prepared.
    a. Addition of empty column "harvested" to each unit on the selection list.
    b. Calculation of target basal area $BA_{target}$ as the product of current total stand basal area (including trees not eligible for harvest) and the desired share of basal area reduction $interv_{BAred}$.
    c. Initialize the buffer state ($S_{buff}$) to the desired initial value $mgmunit_{buff}$.
    d. Initialize empty vectors for collecting groups and cells that have been selected for harvest.
    e. Initialize empty vectors for collecting cells within buffer rings 1 and 2 of groups that have been selected for harvest.
    f. Initialize vector for collecting cumulated basal area of groups selected for harvest with initial value 0 and Boolean ($TBA_{reached}$) indicating if $BA_{target}$ has been reached as FALSE.
7. Harvest operation is carried out in a while-loop with the condition of $S_{buff} \geq 0$.
    a. Select groups to be harvested in for-loop through the ranked selection list.
        i. If any cell within the current group has already been selected for harvest, jump to next list entry.
        ii. If any cells within the current group lie within buffer zone of an already harvested group, jump to next list entry. $S_{buff}$ indicates the number of rings of neighbors that are being checked.
        iii. Mark current group to be selected for harvest.
        iv. Add current group, its cells, and its neighbors in rings 1 and 2 to the corresponding collection vectors.
        v. Add updated cumulated basal area of groups selected for harvest to collection vector.
        vi. If current cumulated basal area of groups selected for harvest exceeds $BA_{target}$, break for-loop.
    b. If current cumulated basal area of groups selected for harvest exceeds $BA_{target}$, break while-loop.
    c. Reduce $S_{buff}$ by 1.
8. Decision whether last selected group for harvest should be retained in harvest list.
    a. Calculate absolute difference between last two entries of vector with cumulated harvested basal area and $BA_{target}$.
    b. If difference is smaller for second to last entry, remove last group to be selected from harvest.

*For the calculation of the weights, cf. section Common features of management submodels STS, GRS, SC, and CAB.

The information on the number of trees harvested per cohort is then assembled and linked to the current list of cohorts, updating the number of trees to be harvested ($dN_{mgm}$).

#### mgm SC

This submodel (new to v1.1) is called if a management intervention is scheduled for the current time step and the management type "SC" (slit cuts) is specified in the simulation settings. It determines the number of trees per cohort to be removed through management ($dN_{mgm}$). It uses the management instructions passed to the model as input data (see section Input data) and a partially randomized process to determine how many, if any, trees per cohort should be removed.

This submodel is largely identical to GRS. The only difference lies in the **grouping algorithm** which builds slit-shaped groups oriented diagonally to the slope. The slit dimensions are user-defined in the management settings and characterized by their width (downslope; $mgmunit_{SW}$) and length (diagonally upwards; $mgmunit_{SL}$) in number of cells (Fig. S1.2). For the rest of the algorithm, refer to the management submodel GRS.

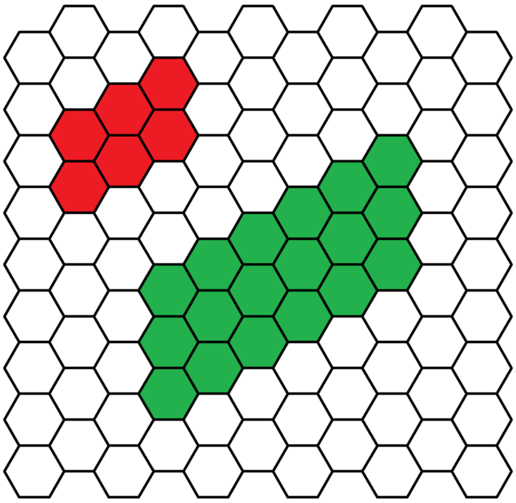

***Fig. S1.2*** *Two examples of slit-shaped groups returned by the grouping algorithm of the submodel mgm SC. The red group has a width of 2 and a length of 3 cells, the green one a width of 3 and a length of 6 cells.*

#### mgm CAB

This submodel (new to v1.1) is called if a management intervention is scheduled for the current time step and the management type "CAB" (cable yarding) is specified in the simulation settings. It determines the number of trees per cohort to be removed through management ($dN_{mgm}$). It uses the management instructions passed to the model as input data (see section Input data) and a partially randomized process to determine how many, if any, trees per cohort should be removed.

The submodel takes the current list of cohort state information (cf. submodel init Data structure), the spatial cell information (cf. submodel init Simulation grid and submodel init Neighbors) and the management instructions as inputs. The output is the list of cohort state information with updated values for the number of trees to be harvested ($dN_{mgm}$). It consists of three algorithms: (1) identification of cable cells (cable algorithm), (2) identification of all possible management units, i.e., slits along the cable lines (grouping algorithm), and (3) the CAB algorithm, defining which units are harvested. The first two algorithms are only run once, at the time step of the first management intervention. The CAB algorithm is called at each intervention time step. To rank the list of units for harvest, the basal area, the regeneration within each unit as well as the regeneration in cells surrounding the unit are taken into account. As the management units can be relatively large, the criteria for (dominant) species and mean crown ratio make little sense. For the same reason, the slits are not necessarily harvested entirely, but iteratively from the cable outwards. Thus, If the target basal area is reached and harvesting stops, the last slit might only be halfway harvested.

Up to three yarding cables can be defined but only one cable is "active" per intervention and all harvesting occurs in horizontal slits along this cable. The cells through which the active cable runs are completely harvested and, in each intervention, another cable is activated, following a regular pattern.

**Cable algorithm**

The cable algorithm identifies the cells through which each of the yarding cables run. The number of cables (one to three) and their position within the simulation grid (as the grid columns) are defined in the management settings ($mgmunit_{CL1\text{-}3}$).

**Grouping algorithm**

The grouping algorithm returns a list of all theoretically possible management units, i.e., slits along cable lines (including overlapping slits). The list contains information on the cells and cohorts included in the slit and their respective neighboring cells in ring 1 (immediately surrounding the group cells) and ring 2 (immediately surrounding cells of ring 1). Each slit is "attached" to a cable line and is positioned either to its left or right. The total number of theoretically possible units is thus the product of the number of rows in the simulation grid, the number of cables defined in the management settings, and the two sides to each cable. Each slit and the cells making it up are successively identified in three nested loops over (1) the yarding cables, (2) the sides to the cable, and (3) the cells along the current cable. Within these nested loops, the horizontal slits are "built" first along the slit length (number of cells outwards from the cable line; $mgmunit_{SL}$) and then along the slit width (number of cells downslope within each slit; $mgmunit_{SW}$) defined in the management settings (Fig. S1.3). Finally, the cells within each group are ordered by distance to the cable.

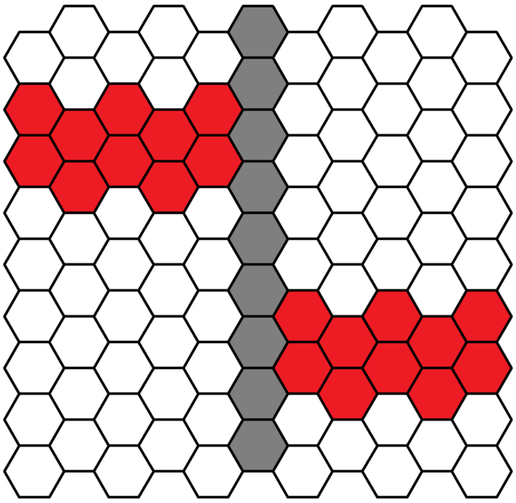

***Fig. S1.3*** *Two of the possible slits (red) along a single cable line (gray). The cable line is positioned in the 6th column of the simulation grid and the slits have a length of five and a width of two cells.*

#### CAB algorithm

1. List of all cohorts with their current state and auxiliary variables is retrieved
2. Current species shares and derived values are calculated for each cohort.
   a. Calculation of the basal area share of all species of the current state 2 ($BAcurr_i$).
   b. Comparison to the minimum species shares ($BAmin_i$) and decision, whether species is eligible for harvest (eligible if current basal area share > $BAmin_i$).
3. Cohorts exempt from harvest are first identified and then removed from list of cohorts to get list of potential harvest cohorts.
   a. All cohorts of tree species that do not fulfill the minimum basal area criterion.
   b. All cohorts that do not fulfill the minimum diameter criterion ($interv_{Dmin}$).
   c. Reduce list to groups belonging to the active cable.
   d. Stop algorithm, if no cohort is eligible for harvest and return $dN_{mgm}$ = 0 for all cohorts.
   e. Remove previously identified cohorts from list of management units and remove units without cohorts eligible for harvest.
4. Characterization of management units
   a. Calculate unit's total basal area.
   b. Calculate share of regeneration (cohorts in stages 1-3) within each unit.
   c. Calculate share of regeneration (cells with all cohorts in stages 1-3) among neighbors of ring 1 of each unit.

5. Selection probabilities are calculated and selection list is ranked.
    a. Calculation selection weight probabilities ($P_{BA}$, $P_{RegUnit}$, *and* $P_{RegNeighb}$) and random number *rn* per unit*.
    b. Calculate selection probability $P_{sel}$ for each unit*.
    c. Order list by descending selection probability $P_{sel}$ and add rank corresponding to it.
6. Harvest operation is prepared and cells along cables are harvested.
    a. Calculation of target basal area $BA_{target}$ as the product of current total stand basal area (including trees not eligible for harvest) and the desired share of basal area reduction $interv_{BAred}$.
    b. Harvest all cohorts with H ≥ 130 cm among cable cells.
    c. Initialize vector for collecting cumulated basal area of groups selected for harvest with initial value corresponding to the basal area harvested along the cable.
    d. Addition of empty column "harvested" to each unit on the selection list.
    e. Initialize the buffer state ($S_{buff}$) to the desired initial value $mgmunit_{buff}$.
    f. Initialize empty vector for collecting cells that have been selected for harvest.
    g. Initialize empty vectors for collecting cells within buffer rings 1 and 2 of groups that have been selected for harvest.
    h. Initialize Boolean ($TBA_{reached}$) indicating if $BA_{target}$ has been reached as FALSE.
    i. Calculate basal area per cell for the entire simulation grid including only cohorts eligible for harvest.

7. Harvest operation is carried out in a while-loop with the condition of $S_{buff} \geq 0$.
   a. Select slits to be harvested in for-loop through the ranked selection list.
      i. If any cell within the current slit has already been selected for harvest, jump to next list entry.
      ii. If too many cells** within the current slit lie within buffer zone of already harvested slits, jump to next list entry.
      iii. Mark current slit to be selected for harvest.
      iv. Add neighbors in rings 1 and 2 of current slit to the corresponding collection vectors.
      v. Iteratively mark cells of slit to be harvested in loop over ordered cell list per slit (from cable outwards)
         1. Retrieve cell basal area (excluding cohorts exempt from harvest)
         2. Calculate absolute values of current and potential difference (including the focus cell) of harvested basal area to target basal area.
         3. If potential difference is smaller than current difference, add focus cell and updated cumulative harvested basal area to corresponding collection vectors.
         4. Else, $BA_{target}$ is exceeded. Break loop over cells per slit
      vi. If current cumulated basal area of groups selected for harvest exceeds $BA_{target}$, break for-loop.
   b. If current cumulated basal area of groups selected for harvest exceeds $BA_{target}$, break while-loop.
   c. Reduce $S_{buff}$ by 0.5.

*For the calculation of the weights, cf. section Common features of management submodels STS, GRS, SC, and CAB.

**In contrast to the other management algorithms using a buffer, the actual buffer value is reduced in steps of 0.5 instead of 1 if the target basal area cannot be reached with the initial buffer value applied. If the actual buffer value is an integer, all slits are excluded from potential harvest that lie within the buffer of the specified with, potentially excluding slits on the same elevation but on the opposite side of the cable. If the actual buffer value is 1.5 or 0.5, slits are only excluded if the number of cells within the buffer exceeds the slit length, i.e., slits at the same altitude but on different sides of the cable are still eligible for harvest.

The information on the number of trees harvested per cohort is then assembled and linked to the current list of cohorts, updating the number of trees to be harvested ($dN_{mgm}$).

#### Submodel group “protective quality”

##### nais Gaps

This submodel first identifies and then characterizes gaps (i.e., cells without cohorts with $D \geq$ 12 cm) in the simulation output of a particular time step. There are two changes from v1.0 to v1.1: The first change concerns the identification of continuous gaps; the description below replaces the corresponding paragraph in the original model documentation. The second change is that all gap characteristics are calculated so that the protective function can later be assessed with any of the natural hazard profiles.

For cells to belong to a gap, two prerequisites have to be fulfilled: a) the larger cohort per cell needs to have $D < 12$ cm, making it a potential gap cell, and b) the cell must be part of a triangularly aligned group of three potential gap cells, making it a gap cell. First, all possible groups of three cells are identified, analogous to the grouping algorithm in submodel mgm GRS. Then, those groups consisting of only potential gap cells (and their respective cells) are identified. Finally, continuous gaps are identified and numbered, following the same algorithm as in v1.0. In a loop over all cells, gap IDs are assigned in a hierarchical way to continuous gaps. If the target cell is a gap cell and if no neighboring cells in the first ring (cf. submodel init Neighbors) have been assigned a gap ID (yet), a new gap ID is assigned to that cell. If only one of the neighbors has a gap ID, the same number is assigned to the target cell. If multiple neighbors have gap IDs already assigned, the smallest of them is assigned to the target cell as well as all other gap cells in the same gap. As this procedure potentially leads to non-continuous gap IDs, continuous IDs are re-assigned at the end. As the neighbors of each cell have been identified across simulation grid edges (toroidal space), gaps can also spread across these edges.

The new prerequisite of potential gap cells having to be part of a triangular group of gap cells was introduced in v1.1 to increase realism in gap recognition. For example, the algorithm now recognizes two larger gap areas that are connected only by a “string” the width of one cell as two separate gaps, and long narrow canopy openings are not classified as gaps any more (Fig. S1.4).

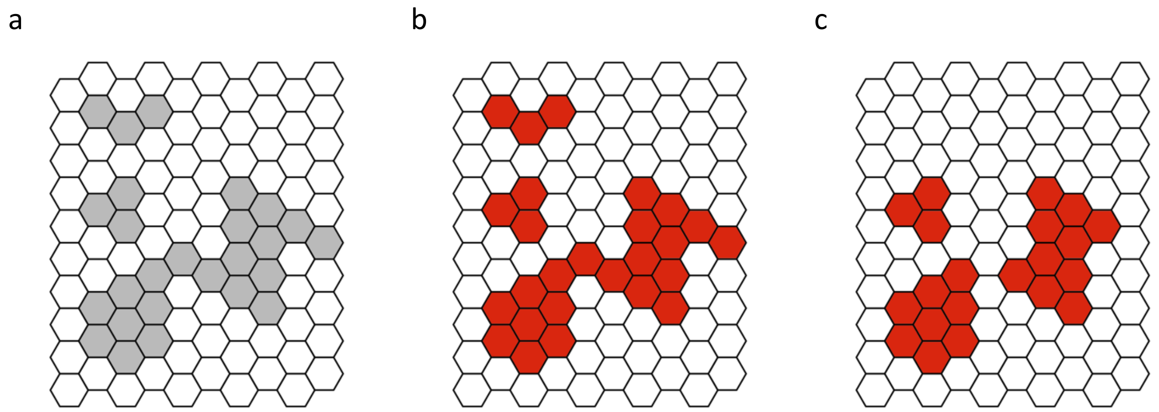

***Fig. S1.4*** *Illustration of the change in the gap recognition algorithm from v1.0 to v1.1. a) Potential gap cells with the largest cohort having D < 12 cm. b) Result of the gap recognition algorithm in v1.0, where all potential gap cells become part of gaps. c) Result of the gap recognition algorithm in v1.1, where the three potential gap cells on the top left are not recognized as a gap and where the potential gap cells in the lower portion of the grid are identified as two separate gaps.*

##### nais Profiles

This submodel generates the minimal and ideal profile to assess the protective function of a simulated stand from different profile elements that depend on the selected site type group, i.e., the simulation's *stratum*, the natural hazard, and, if applicable, the slope and rockfall scenario. In v1.1, the two original profiles for *strata* HM and SA (HM1/HM2 and SA1/SA2, respectively) were merged into one per *stratum*. The new threshold values are only reported in the respective submodel sections if they previously differed between the two groups per *stratum*. Furthermore, the simulation's *site quality* is used to determine the threshold values for the index for vertical structure (cf. submodel nais Vertical structure).

##### Common features of index-submodels

*No changes*

#### nais Mixture

Apart from the merging of the two HM and SA profiles and thus new threshold values (Table S1.1), the submodel remains identical to v1.0.

***Table S1.1*** *Minimal and ideal profiles for the index mix.*

| Stratum - Natural hazard | Species | Minimal profile | Ideal profile |
|---|---|---|---|
| SM | fsyl | 30 – 100% | 50 – 90% |
| | apse | 1 – 70% | 10 – 50% |
| | conifers | 0 – 30% | 0% |
| UM | fsyl | 30 – 80% | 40 – 60% |
| | apse | 1 – 60% | 10 – 30% |
| | aalb | 10 – 60% | 30 – 50% |
| | pabi | 0 – 30% | 0 – 20% |
| UM-LED | fsyl | 30 – 80% | 40 – 60% |
| | apse | 1 – 60% | 10 – 30% |
| | aalb | 20 – 60% | 30 – 50% |
| | pabi | 0 – 30% | 0 – 20% |
| UM-A | fsyl | 30 – 80% | 40 – 60% |
| | apse | 1 – 60% | 10 – 30% |
| | aalb | 10 – 60% | 30 – 50% |
| | pabi | 0 – 30% | 0 – 20% |
| | conifers | 30 – 70% | 30 – 70% |
| HM | aalb | 30 – 90% | 50 – 70% |
| | pabi | 10 – 70% | 30 – 50% |
| SA | pabi | 100% | 100% |

#### nais Vertical structure

In v1.1, the threshold values for calculating the subindex for diameter classes (*vs-DC*) have been recalculated as described below. The calculation of the remaining subindices (*vs-RFSN* and *vs-RFBA*) and of the aggregated index (*vs*) remains identical.

The prerequisites for fulfilling the first **subindex *vs-DC*** are defined in NaiS as there being "enough trees with development potential" in 2-3 diameter classes per hectare. The diameter classes are defined as follows: D1 (< 12 cm DBH), D2 (12 – 30 cm), D3 (30 – 50 cm), and D4 (≥ 50 cm). It is, however, not defined how many trees are enough and what exactly differentiates trees with and without development potential. In v1.0 as in v1.1, the threshold values for class D1 ($RV_{D1}$) equal those for the canopy cover of saplings and thicket ($Tst_{CC}$ as used for the subindex *sapthi-CC*, cf. submodel nais Saplings and thicket) in the minimal or ideal profile, respectively. In v1.0, the threshold values for classes D2 – D4 ($RV_{D2}$, $RV_{D3}$, $RV_{D4}$) were then calculated by equally dividing the remaining share of canopy cover and by further reducing them by a stratum-specific reduction factor ($RF_D$) for the minimal profile. However, simulation analyses showed that stands in an equilibrium state had a considerably larger canopy cover in the lowest class than minimally required and the shares of the upper three classes were well below the threshold values. Consequently, an insufficient index value was awarded even though the diameter distribution was evidently sustainable. Therefore, we derived new threshold values for the upper three classes from stable states of long-term simulations with regular management interventions.

The simulations used to determine the diameter class thresholds were initialized with an equilibrium state of a long-term simulation without management run for 1’000 years. For each of the four strata, combinations of *site* and *regeneration quality* of 3/3 and 5/5 were simulated. Equal species shares were enforced in the regeneration. We applied different management regimes, all of type GRS with aggregates of three cells, with a factorial combination of return intervals (*t*) of 10, 20, 30, and 40 years and of intervention intensities (*i*) of 10, 20, 30, and 40% basal area reduction. The combinations *t*10/*i*30, *t*10/*i*40, and *t*20/*i*40 were deemed unrealistic and thus omitted in the subsequent analysis of the resulting diameter class distribution. The minimum *D* for harvesting was set to 20 cm and the minimum species shares to the lower mixture thresholds for the minimal profile (cf. submodel nais Mixture) and maximum weight was awarded to basal area.

For each of these simulations, the mean cohort share per diameter class in the last 200 simulation years was calculated. For each stratum and for good and for medium sites, respectively, the median of these cohort shares across the respective simulations was then calculated and used as threshold values for D2-4, differentiating between sites with *site quality* ≤ 3 and >3 (Table S1.2). No reduction factor is applied any more. To fulfill the minimal or ideal profile, the reference values in at least 2 or 3 classes have to be reached, respectively.

***Table S1.2**: Minimal and ideal profiles for the subindex vs-DC.*

| Stratum | Site quality | Diameter class | Minimal profile | Ideal profile |
|---|---|---|---|---|
| SM | ≤ 3 | D1 | 5% | 13% |
| | | D2 | 22% | 22% |
| | | D3 | 16% | 16% |
| | | D4 | 15% | 15% |
| | >3 | D1 | 5% | 13% |
| | | D2 | 38% | 38% |
| | | D3 | 18% | 18% |
| | | D4 | 7% | 7% |
| UM | ≤ 3 | D1 | 7% | 12% |
| | | D2 | 20% | 20% |
| | | D3 | 14% | 14% |
| | | D4 | 13% | 13% |
| | >3 | D1 | 7% | 12% |
| | | D2 | 27% | 27% |
| | | D3 | 15% | 15% |
| | | D4 | 15% | 15% |
| HM | ≤ 3 | D1 | 7% | 10% |
| | | D2 | 17% | 17% |
| | | D3 | 16% | 16% |
| | | D4 | 24% | 24% |
| | >3 | D1 | 7% | 10% |
| | | D2 | 18% | 18% |
| | | D3 | 17% | 17% |
| | | D4 | 23% | 23% |
| SA | ≤ 3 | D1 | 12% | 16% |
| | | D2 | 17% | 17% |
| | | D3 | 12% | 12% |
| | | D4 | 13% | 13% |
| | >3 | D1 | 12% | 16% |
| | | D2 | 20% | 20% |
| | | D3 | 14% | 14% |
| | | D4 | 14% | 14% |

##### nais Horizontal arrangement

In v1.1, the rules for the minimal profile in the index for horizontal arrangement, *ha*, were slightly relaxed for the natural hazards snow avalanches and landslides/erosion/debris flow (subindices *ha-g-A* and *ha-ga-LED*) as described below. The remaining subindices and the overall structure of the submodel did not change.

In the relaxed rules for the **subindex *ha-g-A***, a subindex value of 0 (corresponding to the minimal profile) is awarded even if one gap does not fulfill the minimal profile. The rest of the algorithm remains identical, notably also the prerequisites for a subindex value of 1 to be assigned. The rule for assigning a subindex value of 0 was relaxed because in practice, a single gap (slightly) exceeding the threshold value would in many cases be deemed acceptable.

The rules for the **subindex *ha-ga-LED*** were relaxed in two ways. First, as in *ha-g-A* and for the same reason, if one gap exceeds the area threshold for the minimal profile, a subindex value of 0 is still assigned. Second, the rule for determining whether a gap contains “ensured regeneration” and hence the area threshold values are doubled, was relaxed by dropping the prerequisites for species composition within the gap and only focusing on the canopy cover of regeneration in the same way as in v1.0. The reason for dropping the species composition within a gap as a condition for determining whether its regeneration is “ensured” was that not every gap needs to feature a balanced species composition and that the overall species composition of the regeneration is already checked in submodel nais Saplings and thicket.

##### nais Support trees

*No changes*

##### nais Seedlings

In v1.1, the rules for assessing the index for seedlings, *seedl*, were relaxed by removing the subindex checking the seedling cover in gaps, *seedl-GC*. The reasoning behind this change was that germination in ProForM does not depend on light availability, and the appearance of seedlings in an empty cohort thus solely depends on model parameters but not on current stand characteristics. However, a complete lack of seedlings still leads to a low overall index value via the subindex *seedl-sp*.

##### nais Saplings and thicket

Apart from the merging of the two HM and SA profiles and thus new threshold values for the regeneration canopy cover in subindex *sapthi-CC* (Table S1.3) and the species composition in subindex *sapthi-sp* (cf. submodel nais Mixture, Table S1.1), this submodel remains identical to v1.0.

***Table S1.3*** *Minimal and ideal profiles for the subindex sapthi-CC*

| Stratum | Stand characteristic | Minimal profile | Ideal profile |
|---|---|---|---|
| SM | Regeneration cover | 5% | 13% |
| UM | Regeneration cover | 7% | 12% |
| HM | Regeneration cover | 7% | 10% |
| SA | Regeneration cover | 12% | 16% |

#### Model calibration

*No changes*
