## Supplementary material S2 for "Management recommendations for Alpine protection forests: the importance of regeneration quality and initial stand composition"

### Supplementary material S2: NaiS-profiles

#### 1. Upper montane elevational zone (UM)

##### 1.1. Snow avalanches (A)

***Table S2.1*** *NaiS-profile implemented in ProForM v1.1 for the upper montane elevational zone, natural hazard snow avalanches.*

| Index | Minimal profile | Ideal profile |
|---|---|---|
| Mixture | Basal area shares of trees ≥ 12 cm DBH:<br>*Fagus sylvatica*: 30-80%<br>*Acer pseudoplatanus*: 1-60%<br>*Abies alba*: 10-60%<br>*Picea abies*: 0-30%<br>*Coniferous species:* 30-70% | Basal area shares of trees ≥ 12 cm DBH:<br>*Fagus sylvatica*: 40-60%<br>*Acer pseudoplatanus*: 10-30%<br>*Abies alba*: 30-50%<br>*Picea abies*: 0-20%<br>*Coniferous species:* 30-70% |
| Vertical structure | Share of cohorts per diameter class,<br>2 or more of the following classes fulfilled:<br>*Site quality:* *≤ 3* *> 3*<br>DBH < 12 cm: ≥ 7% ≥ 7%<br>DBH 12-30 cm: ≥ 20% ≥ 27%<br>DBH 30-50 cm: ≥ 14% ≥ 15%<br>DBH ≥ 50 cm: ≥ 13% ≥ 15% | Share of cohorts per diameter class,<br>3 or more of the following classes fulfilled:<br>*Site quality:* *≤ 3* *> 3*<br>DBH < 12 cm: ≥ 12% ≥ 12%<br>DBH 12-30 cm: ≥ 20% ≥ 27%<br>DBH 30-50 cm: ≥ 14% ≥ 15%<br>DBH ≥ 50 cm: ≥ 13% ≥ 15% |
| Horizontal arrangement | Gap length[a]: ≤ 50 m (at 36° slope)<br>Gap width[a]: ≤ 5 m, if gap length is exceeded<br>Canopy cover: > 50% | Gap length: ≤ 40 m (at 36° slope)<br>Gap width: ≤ 5 m if gap length is exceeded<br>Canopy cover: > 50% |
| Support trees<br>(Thickest trees in 4 diameter classes, total of 100 $ha^{-1}$) | Mean crown length:<br>*Abies alba*: ≥ 67%<br>*Picea abies*: ≥ 50% | Mean crown length:<br>*Abies alba* & *Picea abies*: ≥ 67% |
| Seedlings<br>(10-40 cm height) | ≥ 2 of the following species present as seedlings on cells without any trees < 40 cm height:<br>*Fagus sylvatica*, *Acer pseudoplatanus*, *Abies alba* | All of the following species present as seedlings on cells without any trees < 40 cm height:<br>*Fagus sylvatica*, *Acer pseudoplatanus*, *Abies alba* |
| Saplings and thicket<br>(40 cm height to 12 cm DBH) | Share of cells with saplings: ≥ 7%<br>Species shares (via cohorts) within saplings analogous to mixture (minimal profile) | Share of cells with saplings: ≥ 12%<br>Species shares (via cohorts) within saplings analogous to mixture (ideal profile) |

[a] one gap exceeding these limits allowed to still fulfill the minimal profile

#### 1.2. Landslide/erosion/debris flow (LED)

***Table S2.2*** *NaiS-profile implemented in ProForM v1.1 for the upper montane elevational zone, natural hazard landslide/erosion/debris flow.*

| **Index** | **Minimal profile** | **Ideal profile** |
|---|---|---|
| Mixture | Basal area shares of trees ≥ 12 cm DBH:<br>*Fagus sylvatica*: 30-80%<br>*Acer pseudoplatanus*: 1-60%<br>*Abies alba*: 20-60%<br>*Picea abies*: 0-30% | Basal area shares of trees ≥ 12 cm DBH:<br>*Fagus sylvatica*: 40-60%<br>*Acer pseudoplatanus*: 10-30%<br>*Abies alba*: 30-50%<br>*Picea abies*: 0-20% |
| Vertical structure | Share of cohorts per diameter class,<br>2 or more of the following classes fulfilled:<br>*Site quality:* *≤ 3* *> 3*<br>DBH < 12 cm: ≥ 7% ≥ 7%<br>DBH 12-30 cm: ≥ 20% ≥ 27%<br>DBH 30-50 cm: ≥ 14% ≥ 15%<br>DBH ≥ 50 cm: ≥ 13% ≥ 15% | Share of cohorts per diameter class,<br>3 or more of the following classes fulfilled:<br>*Site quality:* *≤ 3* *> 3*<br>DBH < 12 cm: ≥ 12% ≥ 12%<br>DBH 12-30 cm: ≥ 20% ≥ 27%<br>DBH 30-50 cm: ≥ 14% ≥ 15%<br>DBH ≥ 50 cm: ≥ 13% ≥ 15% |
| Horizontal arrangement | Gap area[a]: ≤ 600 $m^2$<br>≤ 1’200 $m^2$, with ensured regeneration[b]<br>Canopy cover: > 40% | Gap area: ≤ 400 $m^2$<br>≤ 800 $m^2$, with ensured regeneration[c]<br>Canopy cover: > 60% |
| Support trees<br>(Thickest trees in 4 diameter classes, total of 100 $ha^{-1}$) | Mean crown length:<br>*Abies alba*: ≥ 67%<br>*Picea abies*: ≥ 50% | Mean crown length:<br>*Abies alba* & *Picea abies*: ≥ 67% |
| Seedlings<br>(10-40 cm height) | ≥ 2 of the following species present as seedlings on cells without any trees < 40 cm height:<br>*Fagus sylvatica*, *Acer pseudoplatanus*, *Abies alba* | All of the following species present as seedlings on cells without any trees < 40 cm height:<br>*Fagus sylvatica*, *Acer pseudoplatanus*, *Abies alba* |
| Saplings and thicket<br>(40 cm height to 12 cm DBH) | Share of cells with saplings: ≥ 7%<br>Species shares (via cohorts) within saplings analogous to mixture (minimal profile) | Share of cells with saplings: ≥ 12%<br>Species shares (via cohorts) within saplings analogous to mixture (ideal profile) |

[a] one gap exceeding these limits allowed to still fulfill the minimal profile
[b] cover of saplings and thicket within gap ≥ 12% (corresponds to the ideal threshold of saplings and thicket)
[c] cover of saplings and thicket within gap ≥ 18% (corresponds to 1.5 times the ideal threshold of saplings and thicket)

### 2. High montane elevational zone (HM)

#### 2.1. Snow avalanches (A)

***Table S2.3*** *NaiS-profile implemented in ProForM v1.1 for the high montane elevational zone, natural hazard snow avalanches.*

| Index | Minimal profile | Ideal profile |
|---|---|---|
| Mixture | Basal area shares of trees ≥ 12 cm DBH:<br>*Abies alba*: 30-90%<br>*Picea abies*: 10-70% | Basal area shares of trees ≥ 12 cm DBH:<br>*Abies alba*: 50-70%<br>*Picea abies*: 30-50% |
| Vertical structure | Share of cohorts per diameter class,<br>2 or more of the following classes fulfilled:<br>*Site quality:* *≤ 3* *> 3*<br>DBH < 12 cm: ≥ 7% ≥ 7%<br>DBH 12-30 cm: ≥ 17% ≥ 18%<br>DBH 30-50 cm: ≥ 16% ≥ 17%<br>DBH ≥ 50 cm: ≥ 24% ≥ 23% | Share of cohorts per diameter class,<br>3 or more of the following classes fulfilled:<br>*Site quality:* *≤ 3* *> 3*<br>DBH < 12 cm: ≥ 10% ≥ 10%<br>DBH 12-30 cm: ≥ 17% ≥ 18%<br>DBH 30-50 cm: ≥ 16% ≥ 17%<br>DBH ≥ 50 cm: ≥ 24% ≥ 23% |
| Horizontal arrangement | Gap length[a]: ≤ 50 m (at 36° slope)<br>Gap width[a]: ≤ 15 m, if gap length is exceeded<br>Canopy cover: > 50% | Gap length: ≤ 40 m (at 36° slope)<br>Gap width: ≤ 15 m, if gap length is exceeded<br>Canopy cover: > 50% |
| Support trees<br>(Thickest trees in 4 diameter classes, total of 100 $ha^{-1}$) | Mean crown length:<br>*Abies alba* & *Picea abies*: ≥ 50% | Mean crown length:<br>*Abies alba* & *Picea abies*: ≥ 67% |
| Seedlings<br>(10-40 cm height) | ≥ 1 of the following species present as seedlings on cells without any trees < 40 cm height:<br>*Abies alba, Picea abies* | All of the following species present as seedlings on cells without any trees < 40 cm height:<br>*Abies alba, Picea abies* |
| Saplings and thicket<br>(40 cm height to 12 cm DBH) | Share of cells with saplings: ≥ 7%<br>Species shares (via cohorts) within saplings analogous to mixture (minimal profile) | Share of cells with saplings: ≥ 10%<br>Species shares (via cohorts) within saplings analogous to mixture (ideal profile) |

[a] one gap exceeding these limits allowed to still fulfill the minimal profile

#### 2.2. Landslide/erosion/debris flow (LED)

***Table S2.4*** *NaiS-profile implemented in ProForM v1.1 for the high montane elevational zone, natural hazard landslide/erosion/debris flow.*

| Index | Minimal profile | Ideal profile |
|---|---|---|
| Mixture | Basal area shares of trees ≥ 12 cm DBH:<br>*Abies alba*: 30-90%<br>*Picea abies*: 10-70% | Basal area shares of trees ≥ 12 cm DBH:<br>*Abies alba*: 50-70%<br>*Picea abies*: 30-50% |
| Vertical structure | Share of cohorts per diameter class,<br>2 or more of the following classes fulfilled:<br>*Site quality: ≤ 3 > 3*<br>DBH < 12 cm: ≥ 7% ≥ 7%<br>DBH 12-30 cm: ≥ 17% ≥ 18%<br>DBH 30-50 cm: ≥ 16% ≥ 17%<br>DBH ≥ 50 cm: ≥ 24% ≥ 23% | Share of cohorts per diameter class,<br>3 or more of the following classes fulfilled:<br>*Site quality: ≤ 3 > 3*<br>DBH < 12 cm: ≥ 10% ≥ 10%<br>DBH 12-30 cm: ≥ 17% ≥ 18%<br>DBH 30-50 cm: ≥ 16% ≥ 17%<br>DBH ≥ 50 cm: ≥ 24% ≥ 23% |
| Horizontal arrangement | Gap area[a]: ≤ 600 $m^2$<br>≤ 1'200 $m^2$, with ensured regeneration[b]<br>Canopy cover: > 40% | Gap area: ≤ 400 $m^2$<br>≤ 800 $m^2$, with ensured regeneration[c]<br>Canopy cover: > 60% |
| Support trees<br>(Thickest trees in 4 diameter classes, total of 100 $ha^{-1}$) | Mean crown length:<br>*Abies alba* & *Picea abies*: ≥ 50% | Mean crown length:<br>*Abies alba* & *Picea abies*: ≥ 67% |
| Seedlings<br>(10-40 cm height) | ≥ 1 of the following species present as seedlings on cells without any trees < 40 cm height:<br>*Abies alba, Picea abies* | All of the following species present as seedlings on cells without any trees < 40 cm height:<br>*Abies alba, Picea abies* |
| Saplings and thicket<br>(40 cm height to 12 cm DBH) | Share of cells with saplings: ≥ 7%<br>Species shares (via cohorts) within saplings analogous to mixture (minimal profile) | Share of cells with saplings: ≥ 10%<br>Species shares (via cohorts) within saplings analogous to mixture (ideal profile) |

[a] one gap exceeding these limits allowed to still fulfill the minimal profile
[b] cover of saplings and thicket within gap ≥ 10% (corresponds to the ideal threshold of saplings and thicket)
[c] cover of saplings and thicket within gap ≥ 15% (corresponds to 1.5 times the ideal threshold of saplings and thicket)

### 3. Subalpine elevational zone (SA)

#### 3.1. Snow avalanches (A)

***Table S2.5*** *NaiS-profile implemented in ProForM v1.1 for the subalpine elevational zone, natural hazard snow avalanches.*

| **Index** | **Minimal profile** | **Ideal profile** |
|---|---|---|
| Mixture | Basal area shares of trees ≥ 12 cm DBH:<br>*Picea abies*: 100% | Basal area shares of trees ≥ 12 cm DBH:<br>*Picea abies*: 100% |
| Vertical structure | Share of cohorts per diameter class,<br>2 or more of the following classes fulfilled:<br>*Site quality:* ≤ 3 / > 3<br>DBH < 12 cm: ≥ 12% / ≥ 12%<br>DBH 12-30 cm: ≥ 17% / ≥ 20%<br>DBH 30-50 cm: ≥ 12% / ≥ 14%<br>DBH ≥ 50 cm: ≥ 13% / ≥ 14% | Share of cohorts per diameter class,<br>3 or more of the following classes fulfilled:<br>*Site quality:* ≤ 3 / > 3<br>DBH < 12 cm: ≥ 16% / ≥ 16%<br>DBH 12-30 cm: ≥ 17% / ≥ 20%<br>DBH 30-50 cm: ≥ 12% / ≥ 14%<br>DBH ≥ 50 cm: ≥ 13% / ≥ 14% |
| Horizontal arrangement | Gap length[a]: ≤ 50 m (at 36° slope)<br>Gap width[a]: ≤ 15 m, if gap length is exceeded<br>Canopy cover: > 50% | Gap length: ≤ 40 m (at 36° slope)<br>Gap width: ≤ 15 m, if gap length is exceeded<br>Canopy cover: > 50% |
| Support trees<br>(Thickest trees in 4 diameter classes, total of 100 $ha^{-1}$) | Mean crown length:<br>*Picea abies*: ≥ 67% | Mean crown length:<br>*Picea abies*: 100% |
| Seedlings<br>(10-40 cm height) | *Picea abies* present as seedlings on cells without any trees < 40 cm height | *Picea abies* present as seedlings on cells without any trees < 40 cm height |
| Saplings and thicket<br>(40 cm height to 12 cm DBH) | Share of cells with saplings: ≥ 12%<br>Species shares (via cohorts) within saplings analogous to mixture (minimal profile) | Share of cells with saplings: ≥ 16%<br>Species shares (via cohorts) within saplings analogous to mixture (ideal profile) |

[a] one gap exceeding these limits allowed to still fulfill the minimal profile

#### 3.2. Landslide/erosion/debris flow (LED)

***Table S2.6*** *NaiS-profile implemented in ProForM v1.1 for the subalpine elevational zone, natural hazard landslide/erosion/debris flow.*

| **Index** | **Minimal profile** | **Ideal profile** |
|---|---|---|
| Mixture | Basal area shares of trees ≥ 12 cm DBH:<br>*Picea abies*: 100% | Basal area shares of trees ≥ 12 cm DBH:<br>*Picea abies*: 100% |
| Vertical structure | Share of cohorts per diameter class,<br>2 or more of the following classes fulfilled:<br>*Site quality:* ≤ 3 / > 3<br>DBH < 12 cm: ≥ 12% / ≥ 12%<br>DBH 12-30 cm: ≥ 17% / ≥ 20%<br>DBH 30-50 cm: ≥ 12% / ≥ 14%<br>DBH ≥ 50 cm: ≥ 13% / ≥ 14% | Share of cohorts per diameter class,<br>3 or more of the following classes fulfilled:<br>*Site quality:* ≤ 3 / > 3<br>DBH < 12 cm: ≥ 16% / ≥ 16%<br>DBH 12-30 cm: ≥ 17% / ≥ 20%<br>DBH 30-50 cm: ≥ 12% / ≥ 14%<br>DBH ≥ 50 cm: ≥ 13% / ≥ 14% |
| Horizontal arrangement | Gap area[a,b]: ≤ 600 $m^2$<br>≤ 1'200 $m^2$, with ensured regeneration[c]<br>Canopy cover: > 40% | Gap area[b]: ≤ 400 $m^2$<br>≤ 800 $m^2$, with ensured regeneration[d]<br>Canopy cover: > 60% |
| Support trees<br>(Thickest trees in 4 diameter classes, total of 100 $ha^{-1}$) | Mean crown length:<br>*Picea abies*: ≥ 67% | Mean crown length:<br>*Picea abies*: 100% |
| Seedlings<br>(10-40 cm height) | *Picea abies* present as seedlings on cells without any trees < 40 cm height | *Picea abies* present as seedlings on cells without any trees < 40 cm height |
| Saplings and thicket<br>(40 cm height to 12 cm DBH) | Share of cells with saplings: ≥ 12%<br>Species shares (via cohorts) within saplings analogous to mixture (minimal profile) | Share of cells with saplings: ≥ 16%<br>Species shares (via cohorts) within saplings analogous to mixture (ideal profile) |

[a] one gap exceeding these limits allowed to still fulfill the minimal profile
[b] larger areas acceptable if gap is slit-shaped and no wider than 20m
[c] cover of saplings and thicket within gap ≥ 16% (corresponds to the ideal threshold of saplings and thicket)
[d] cover of saplings and thicket within gap ≥ 24% (corresponds to 1.5 times the ideal threshold of saplings and thicket)
