## Supplementary material S3 for "Management recommendations for Alpine protection forests: the importance of regeneration quality and initial stand composition"

#### Supplementary material S3: Initialization stands

In the following graphs, the characteristics of the initialization stands are depicted.

The upper left panel contains the identification of the initialization stand as well as key characteristics of the stand:

- Elev. zone = Elevational zone (*UM*, *HM*, or *SA*)
- Stand type = initialization stand type (*init-ST-y*, *init-ST-s*, *init-ST-m*, or *init-LT*)
- $Q_{site}$ = site quality (*good* or *medium*)
- $Q_{reg}$ = regeneration quality (*good* or *hindered*)
- $BA_{adu}$ = Basal area of adult trees (DBH ≥ 12 cm)
- $SN_{reg}$ = Stem number of regeneration (DBH < 12 cm)
- $SN_{adu}$ = Stem number of adult trees (DBH ≥ 12 cm)
- CC = Canopy cover, i.e., share of cells with trees with DBH ≥ 12 cm (without reduction factor used in assessment of protective quality)

The two remaining panels in the upper row show the distribution of basal area and stem numbers among species for regeneration and adult trees, respectively. The lower panel shows the diameter distribution of trees with DBH ≥ 4 cm per species.

The graphs are ordered by elevational zone, initialization stand type, site and regeneration quality.

### 1. Upper montane zone

#### 1.1. Young stand (init-ST-y)

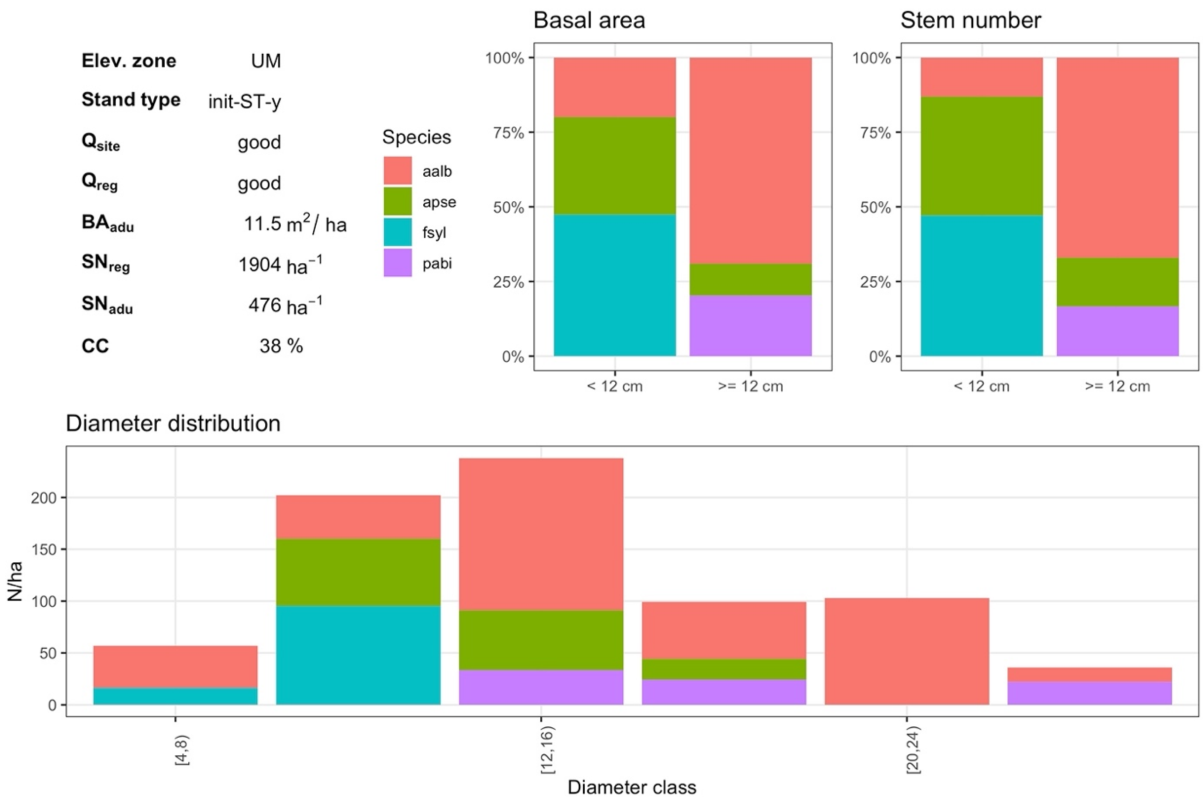


***Fig. S3.1*** *UM init-ST-y with good site quality (5) and good regeneration quality (5)*

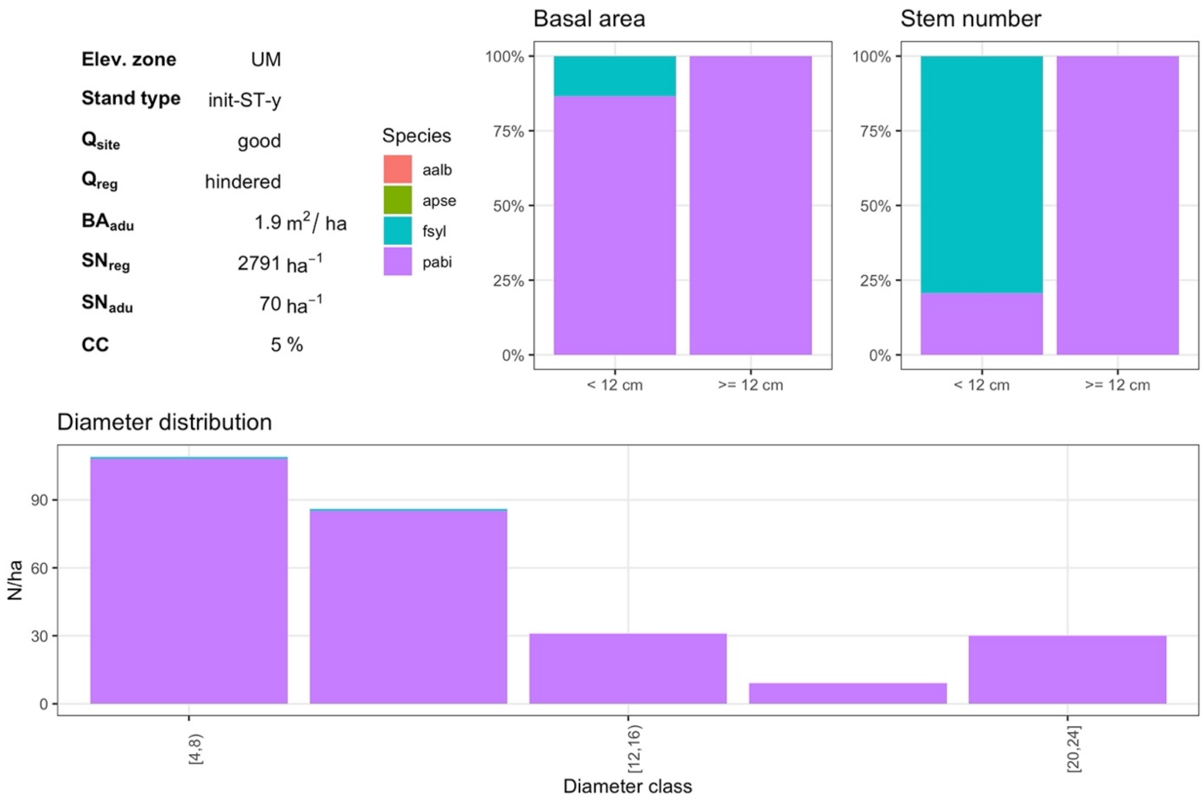


***Fig. S3.2*** *UM init-ST-y with good site quality (5) and hindered regeneration quality (1)*

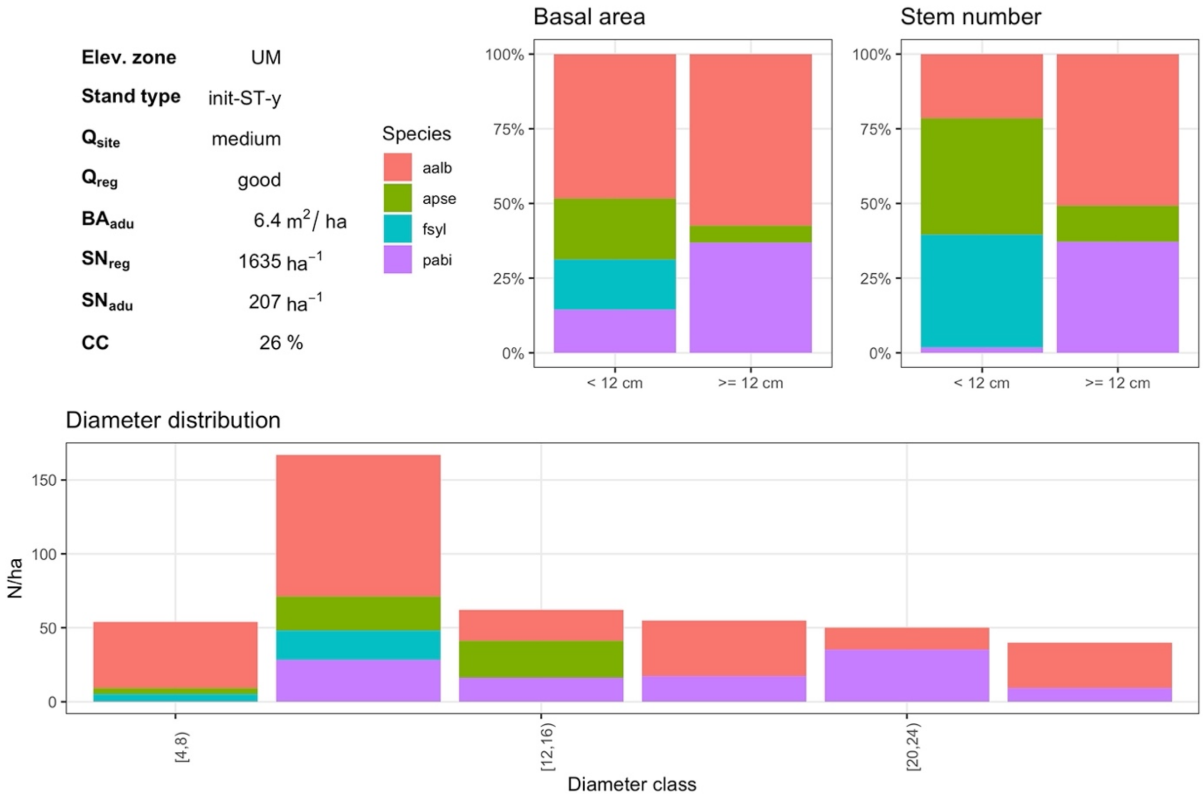


***Fig. S3.3*** *UM init-ST-y with medium site quality (3) and good regeneration quality (3)*

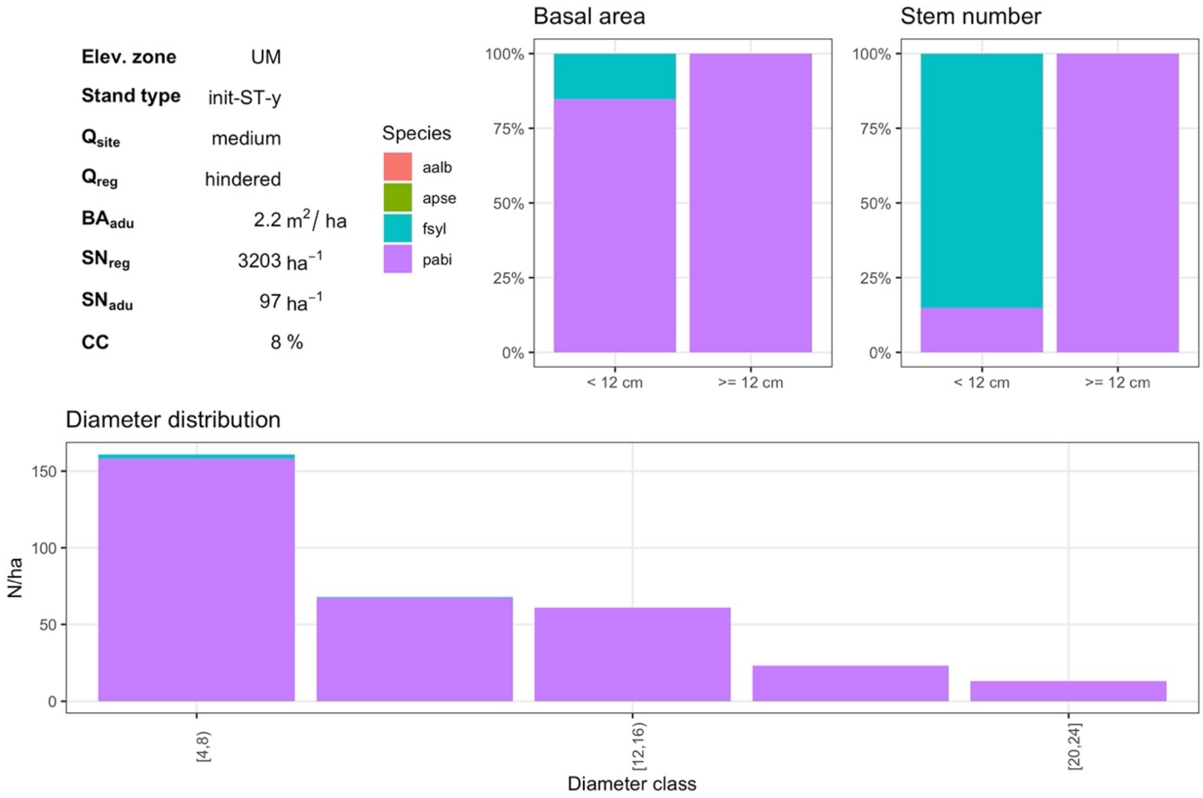


***Fig. S3.4*** *UM init-ST-y with medium site quality (3) and hindered regeneration quality (1)*

#### 1.2. Well-structured stand (init-ST-s)

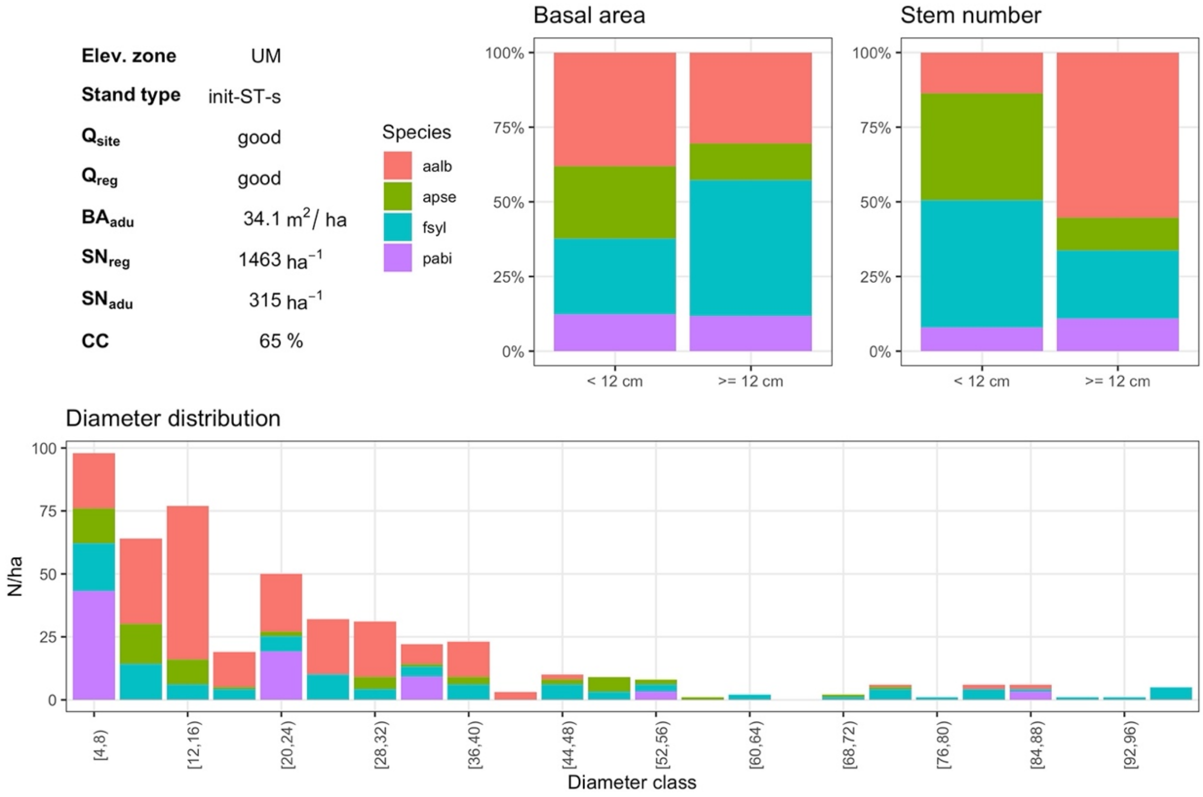


***Fig. S3.5*** *UM init-ST-s with good site quality (5) and good regeneration quality (5)*

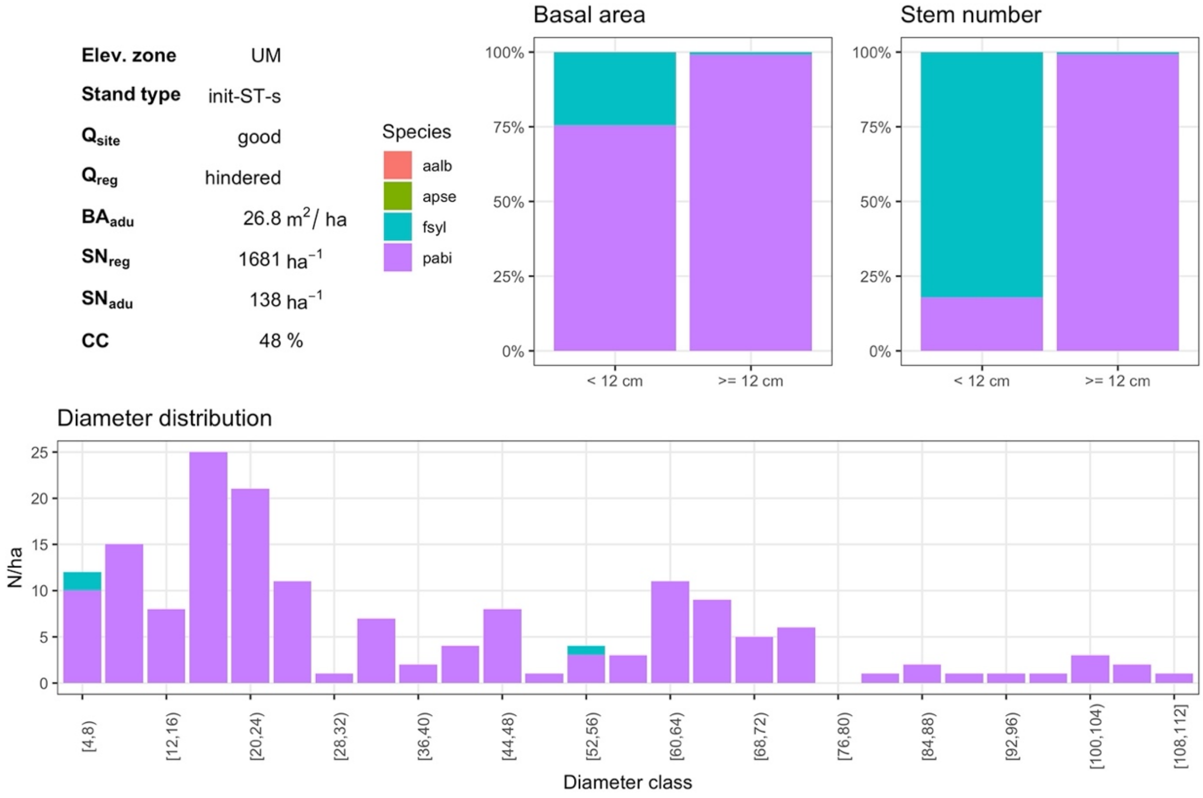


***Fig. S3.6*** *UM init-ST-s with good site quality (5) and hindered regeneration quality (1)*

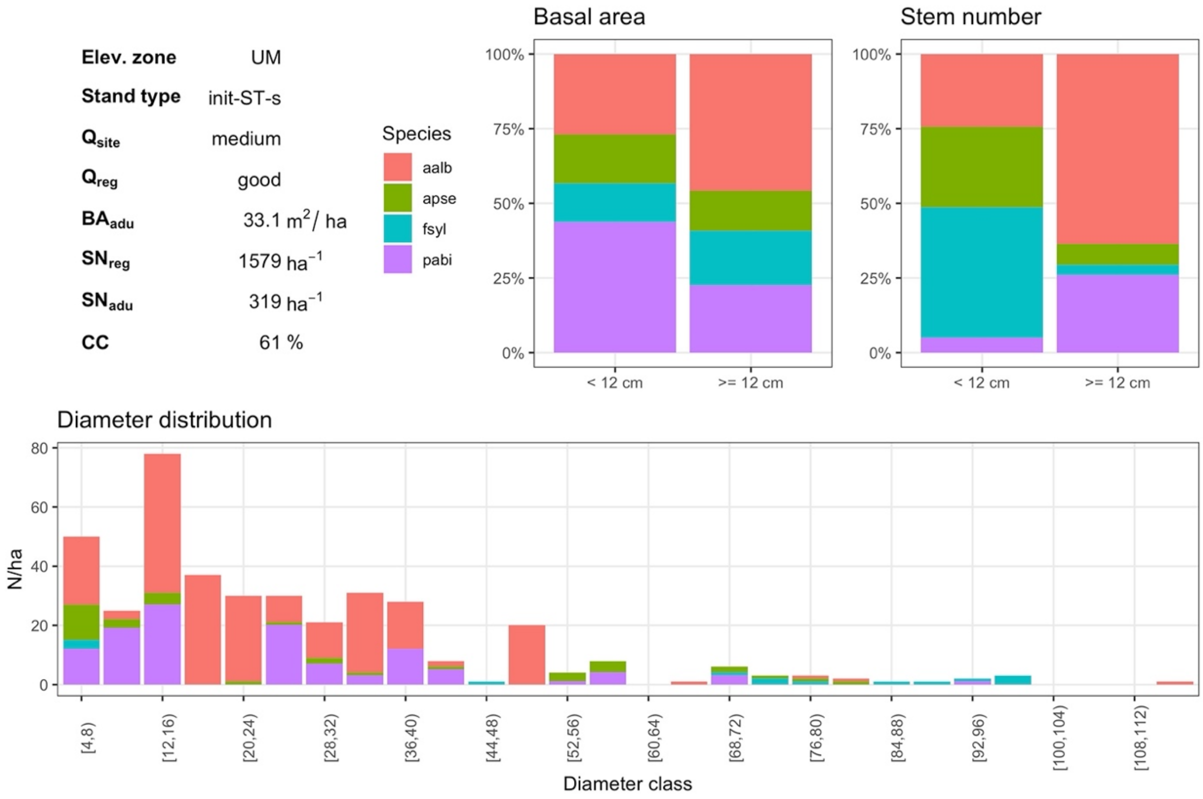


***Fig. S3.7*** *UM init-ST-s with medium site quality (3) and good regeneration quality (3)*

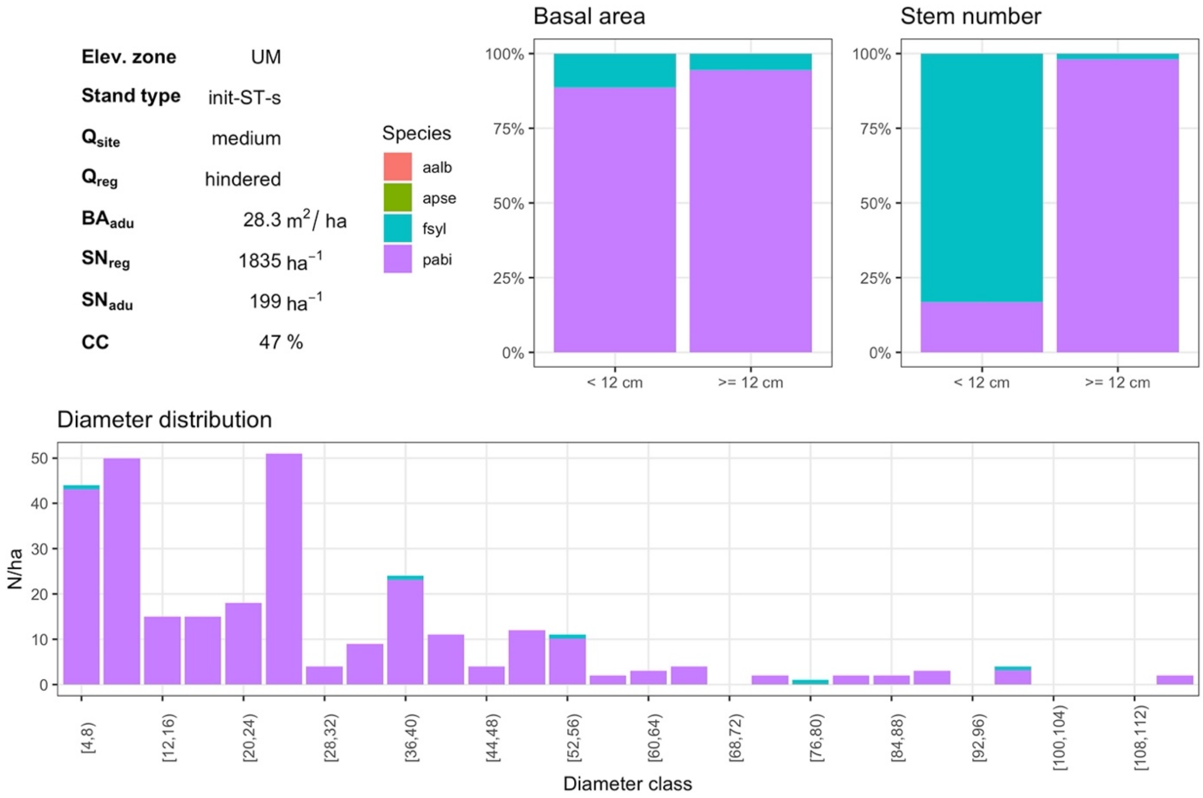


***Fig. S3.8*** *UM init-ST-s with medium site quality (3) and hindered regeneration quality (1)*

#### 1.3. Mature stand (init-ST-m)

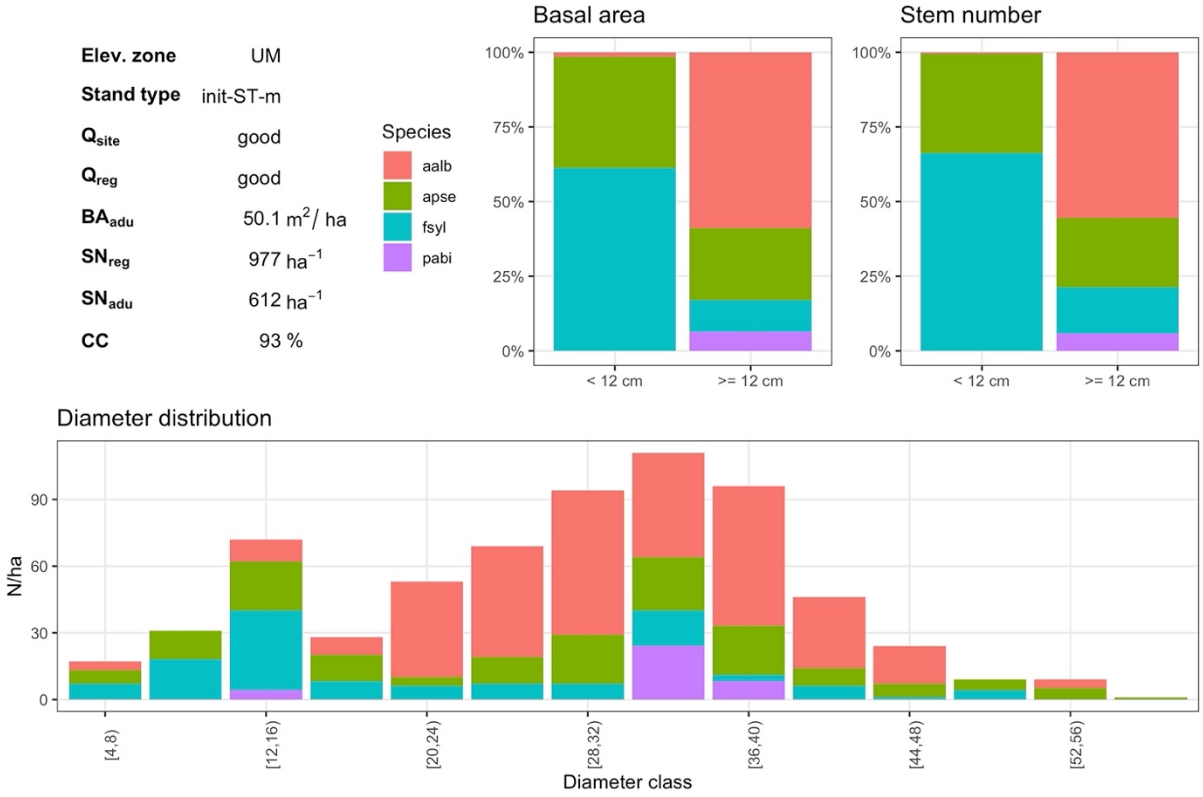


***Fig. S3.9*** *UM init-ST-m with good site quality (5) and good regeneration quality (5)*

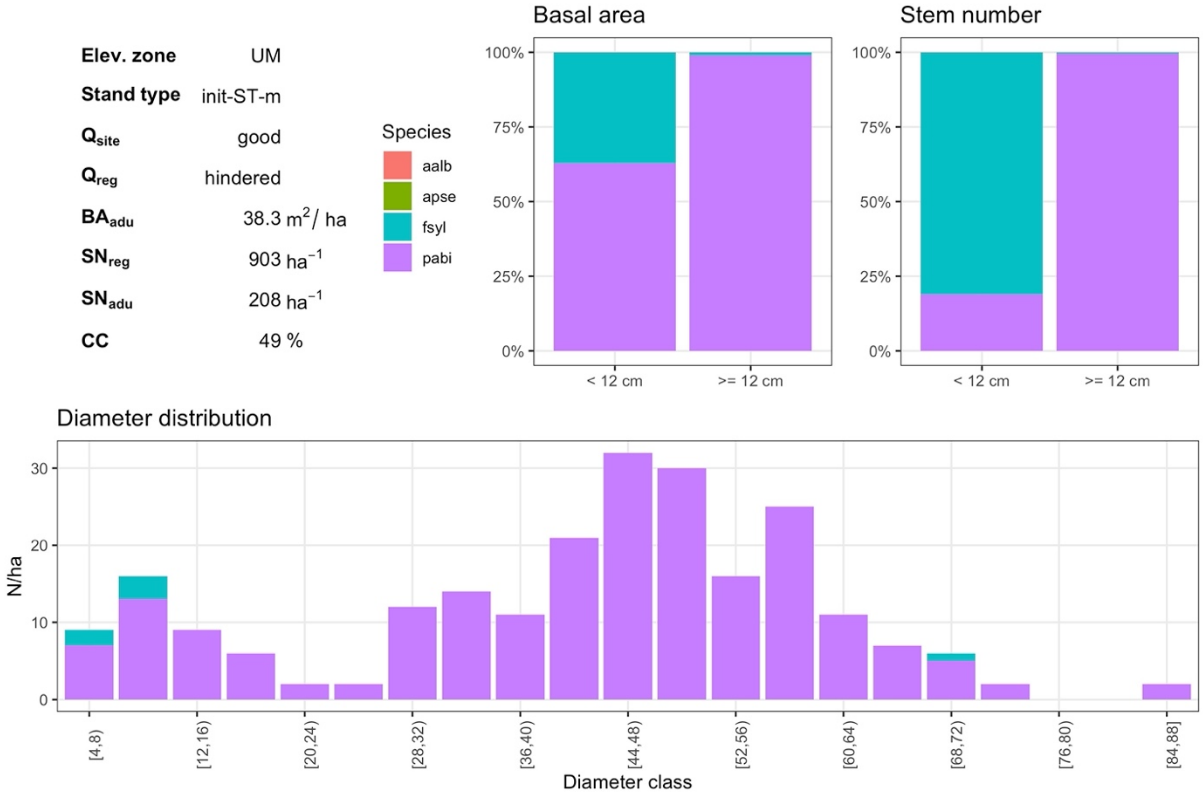


***Fig. S3.10*** *UM init-ST-m with good site quality (5) and hindered regeneration quality (1)*

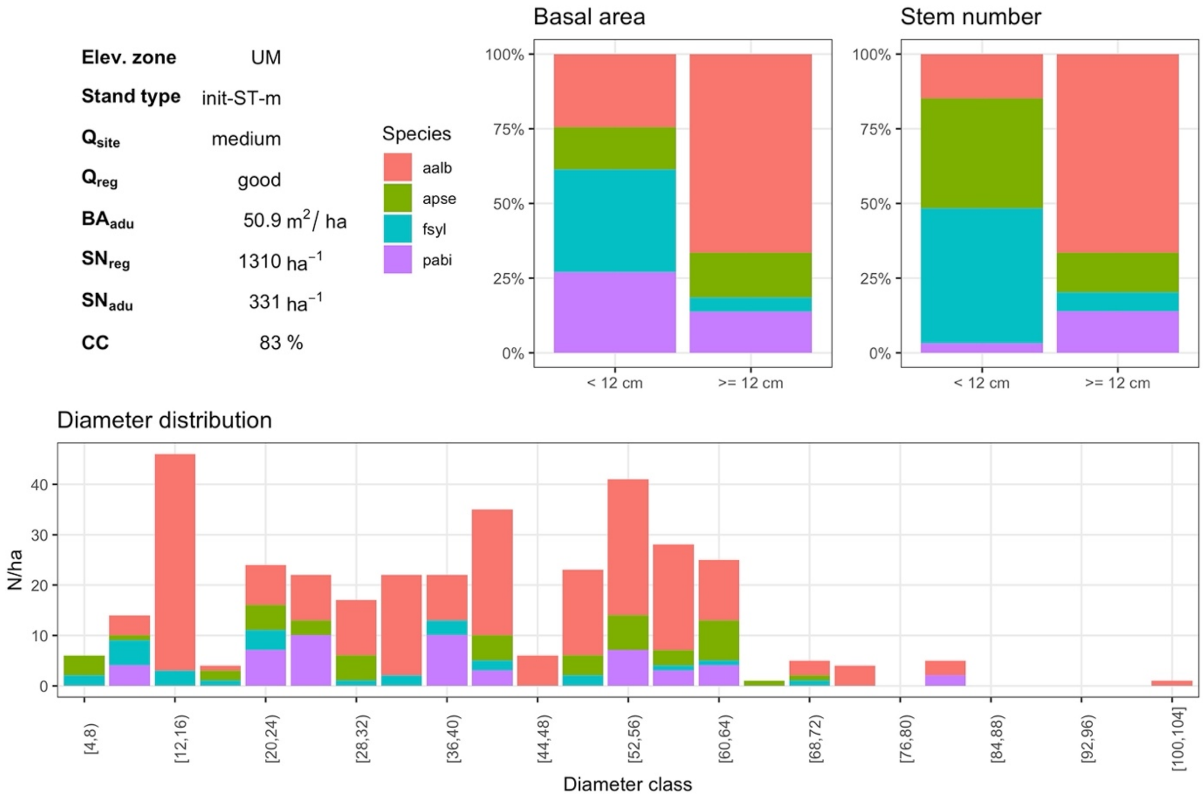


***Fig. S3.11*** *UM init-ST-m with medium site quality (3) and good regeneration quality (3)*

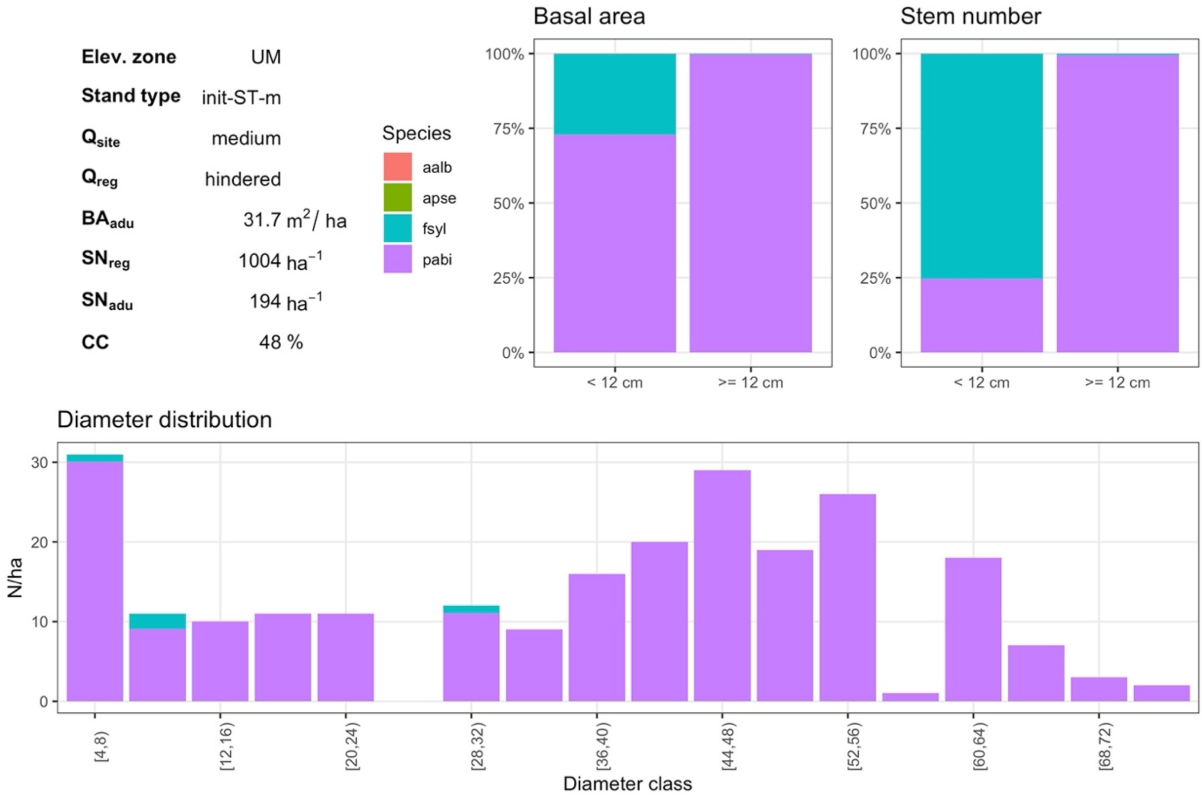


***Fig. S3.12*** *UM init-ST-m with medium site quality (3) and hindered regeneration quality (1)*

#### 1.4. Long-term simulation (init-LT)

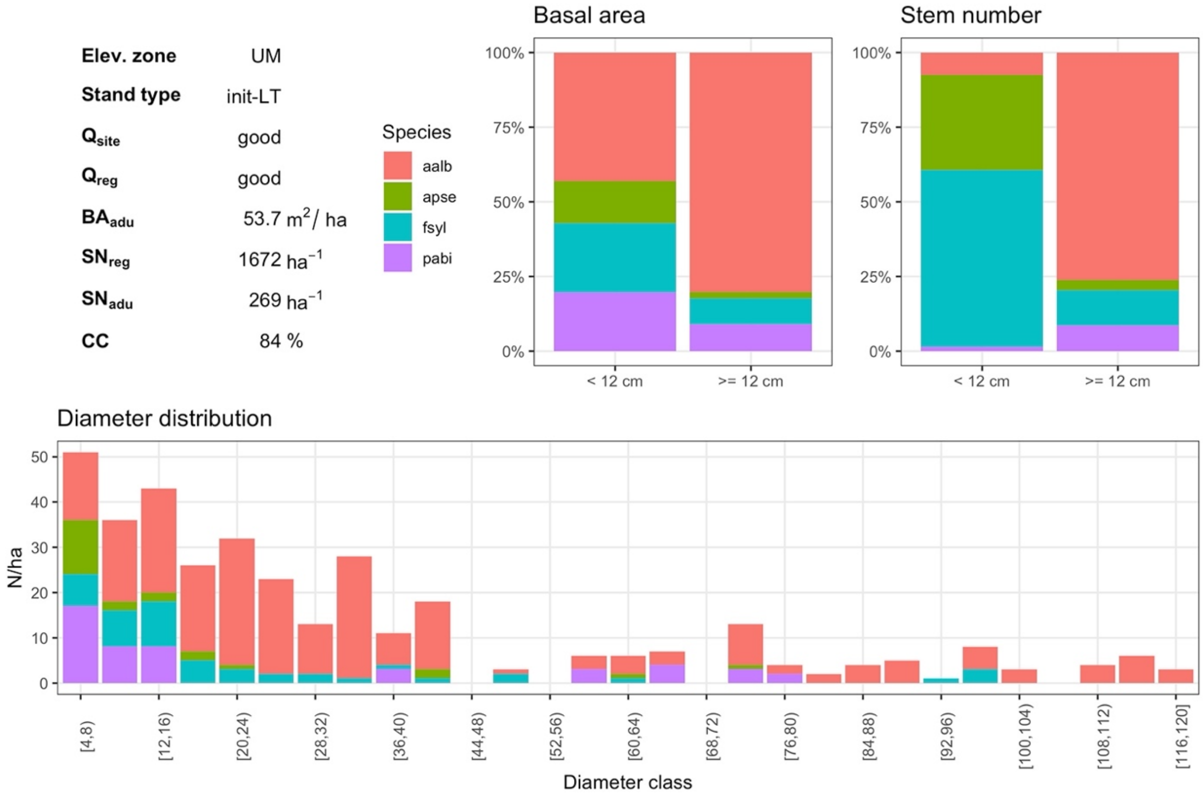


***Fig. S3.13*** *UM init-LT with good site quality (5) and good regeneration quality (5)*

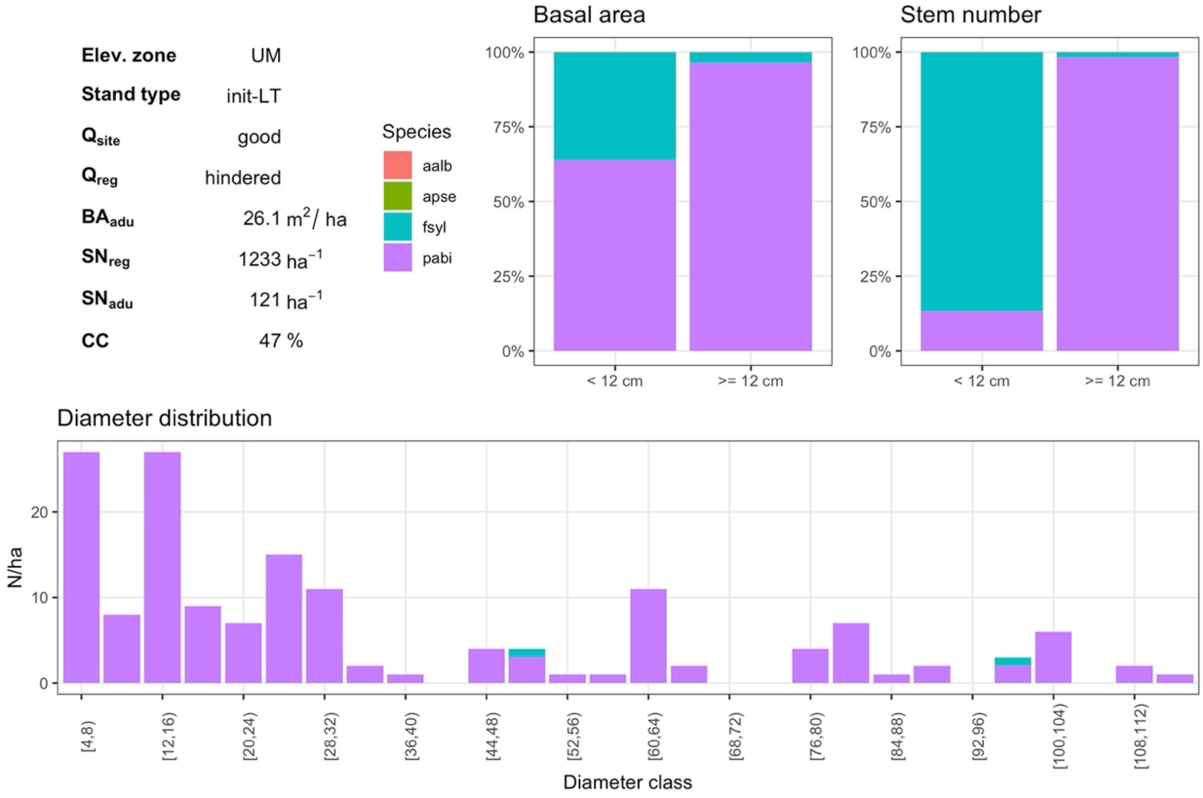


***Fig. S3.14*** *UM init-LT with good site quality (5) and hindered regeneration quality (1)*

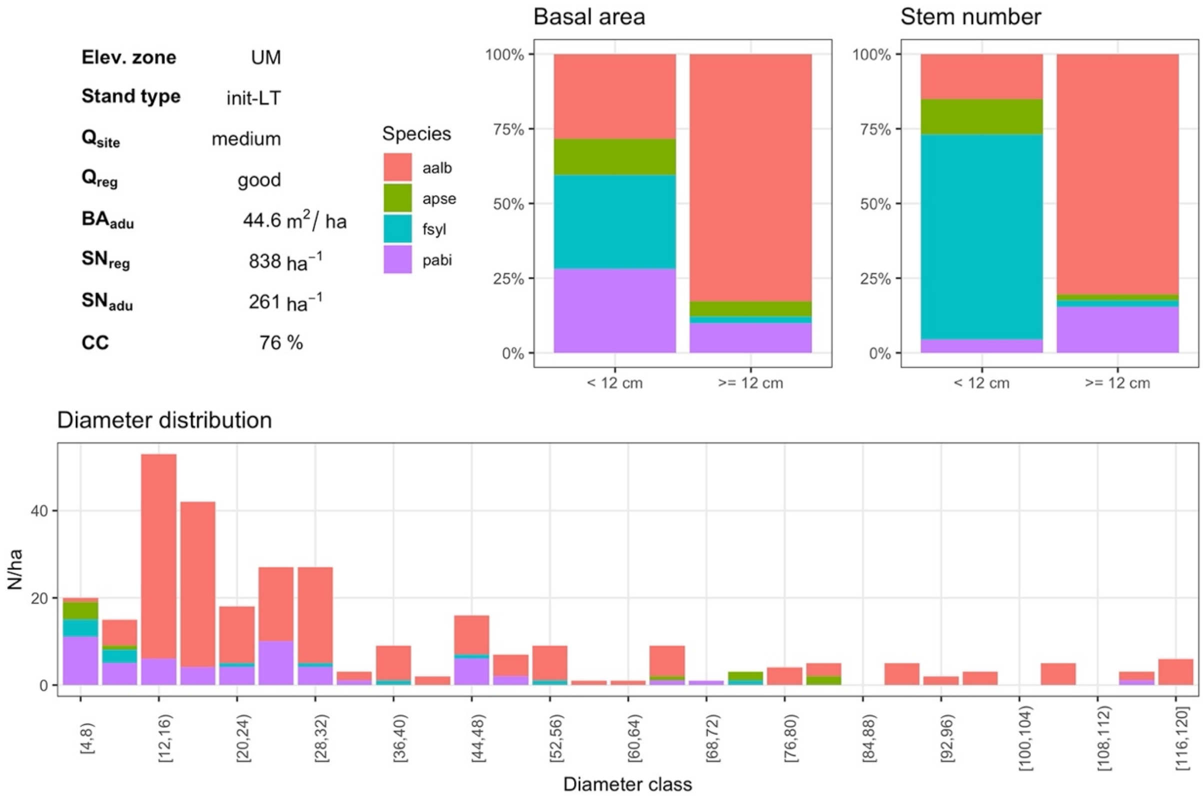


***Fig. S3.15*** *UM init-LT with medium site quality (3) and good regeneration quality (3)*

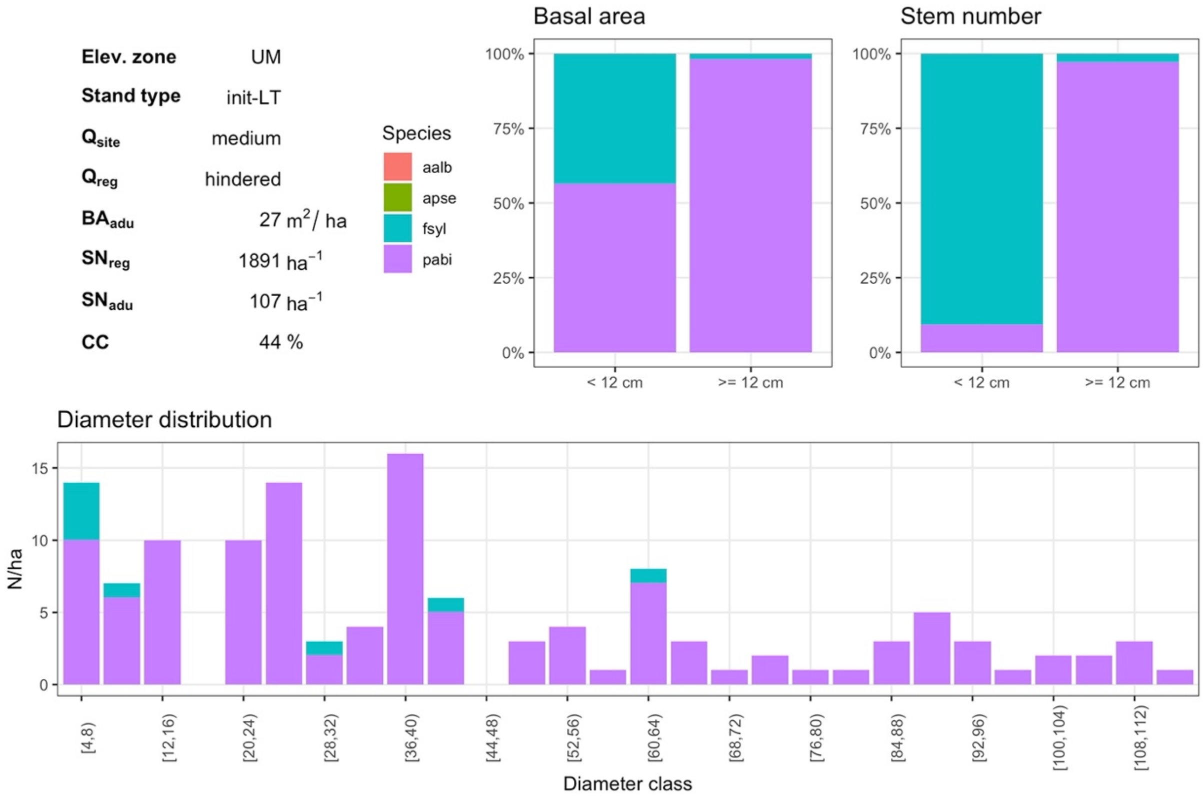


***Fig. S3.16*** *UM init-LT with medium site quality (3) and hindered regeneration quality (1)*

### 2. High montane zone

#### 2.1. Young stand (init-ST-y)

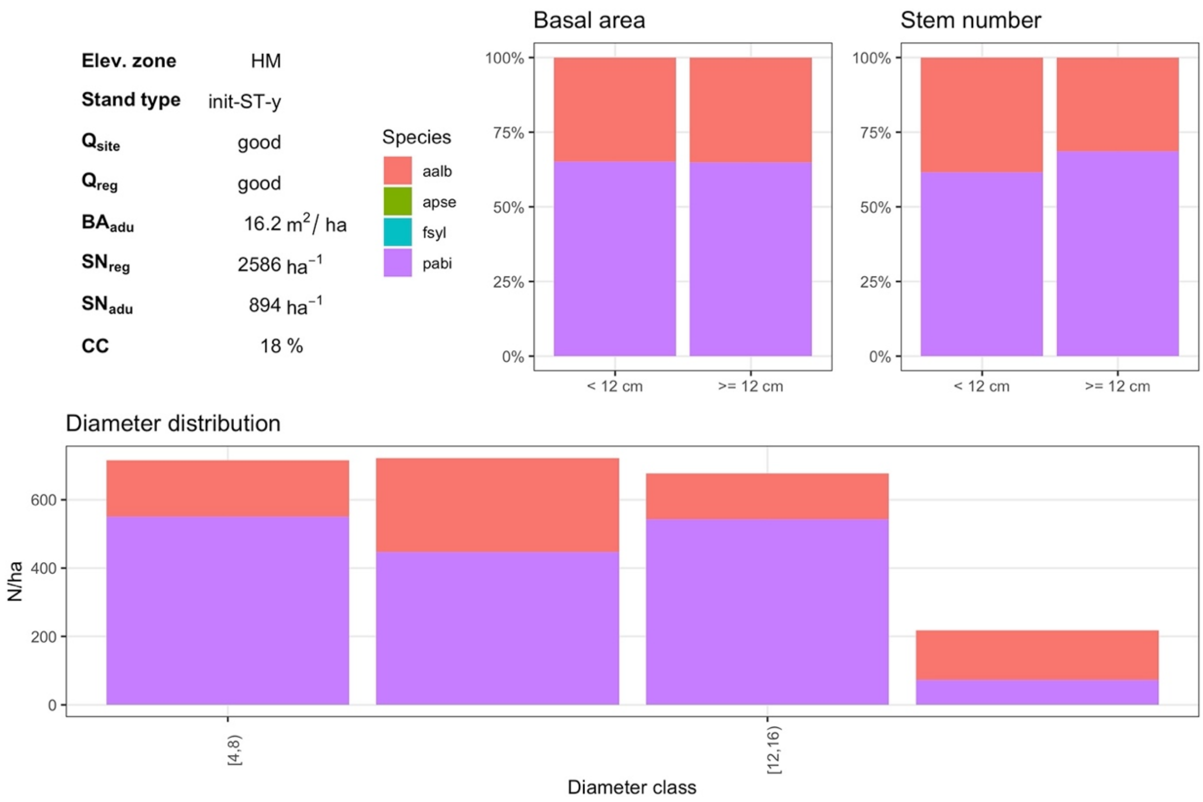


***Fig. S3.17*** *HM init-ST-y with good site quality (5) and good regeneration quality (5)*

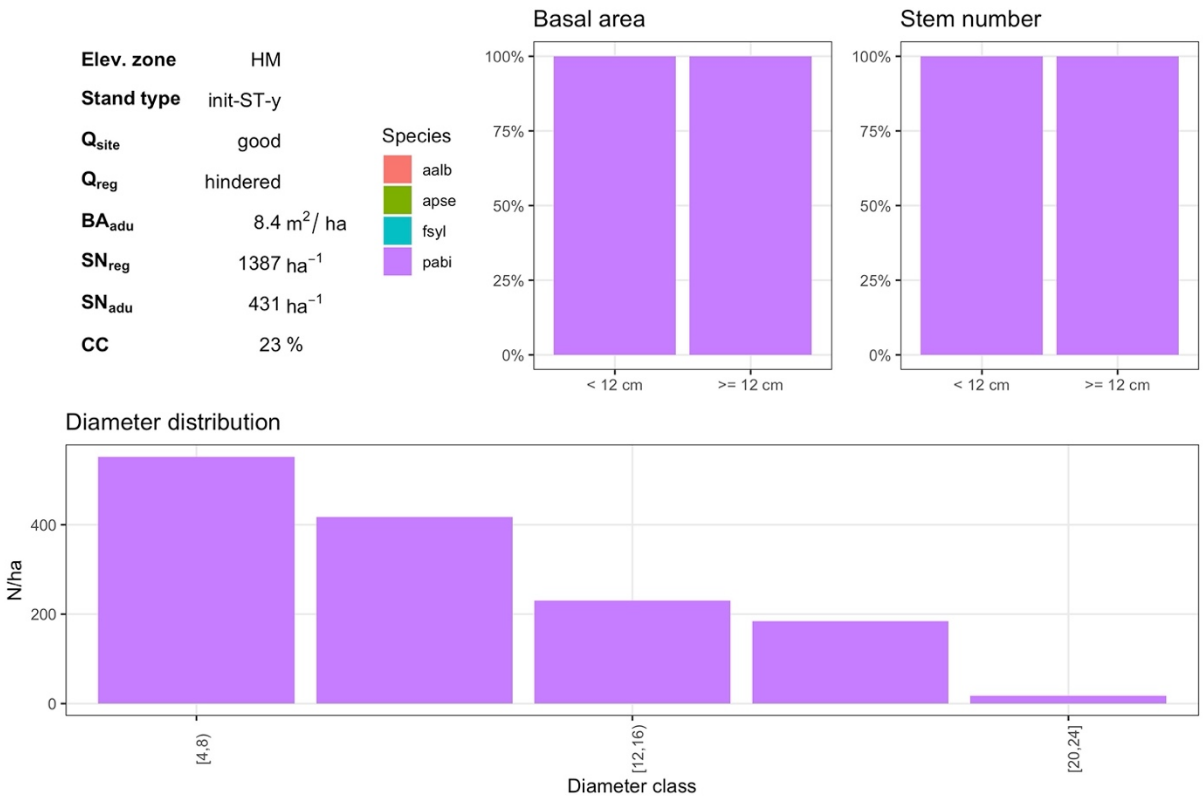


***Fig. S3.18*** *HM init-ST-y with good site quality (5) and hindered regeneration quality (1)*

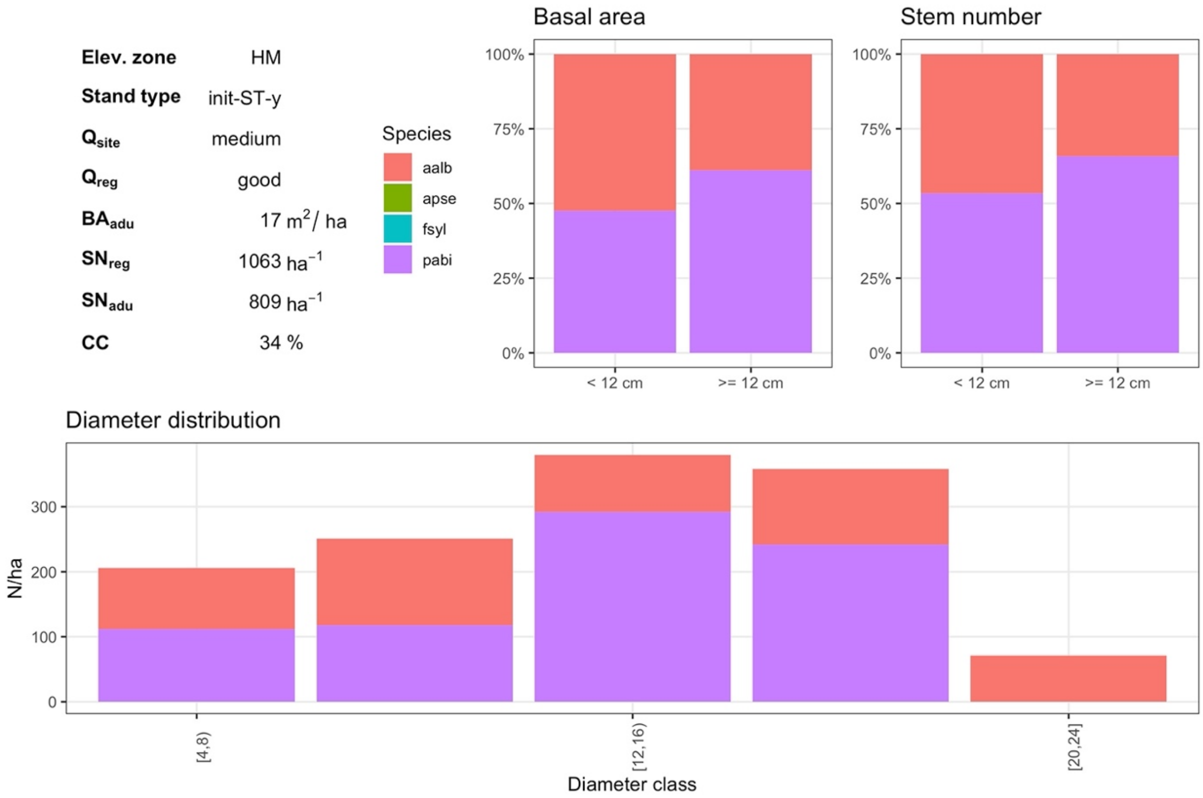


***Fig. S3.19*** *HM init-ST-y with medium site quality (3) and good regeneration quality (3)*

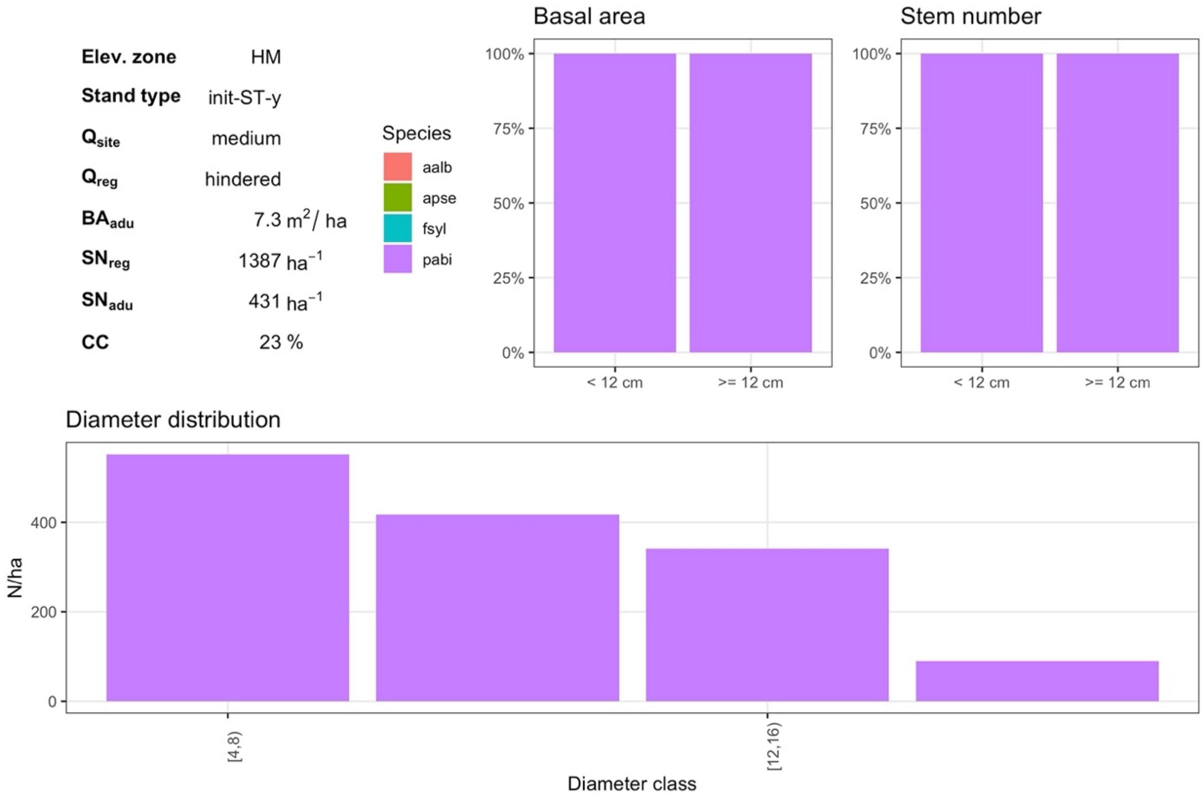


***Fig. S3.20*** *HM init-ST-y with medium site quality (3) and hindered regeneration quality (1)*

#### 2.2. Well-structured stand (init-ST-s)

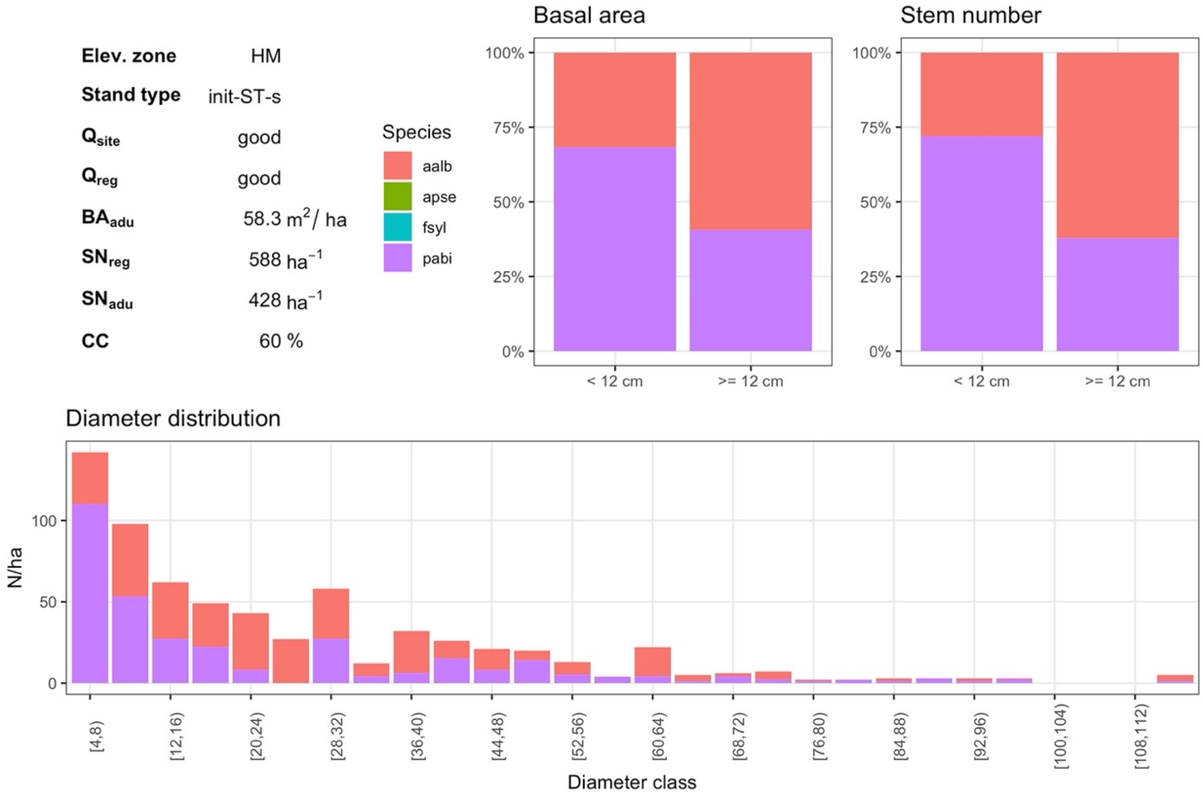


***Fig. S3.21*** *HM init-ST-s with good site quality (5) and good regeneration quality (5)*

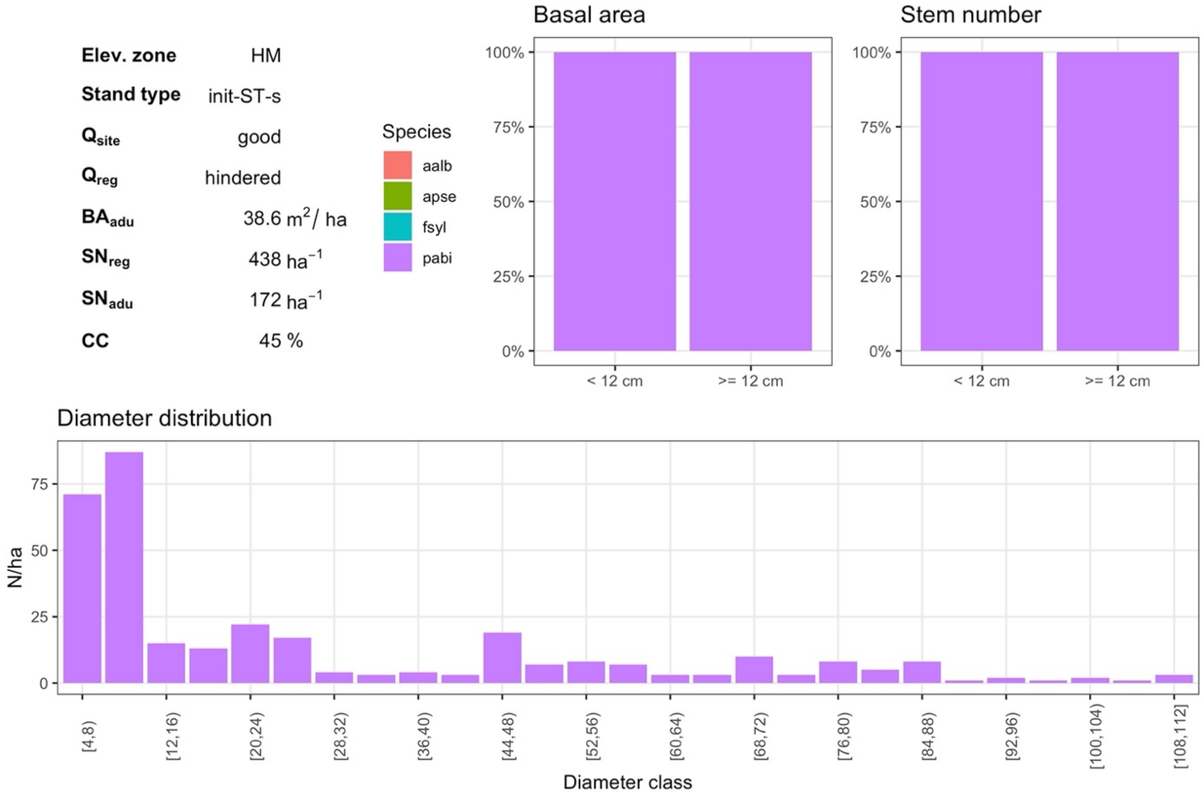


***Fig. S3.22*** *HM init-ST-s with good site quality (5) and hindered regeneration quality (1)*

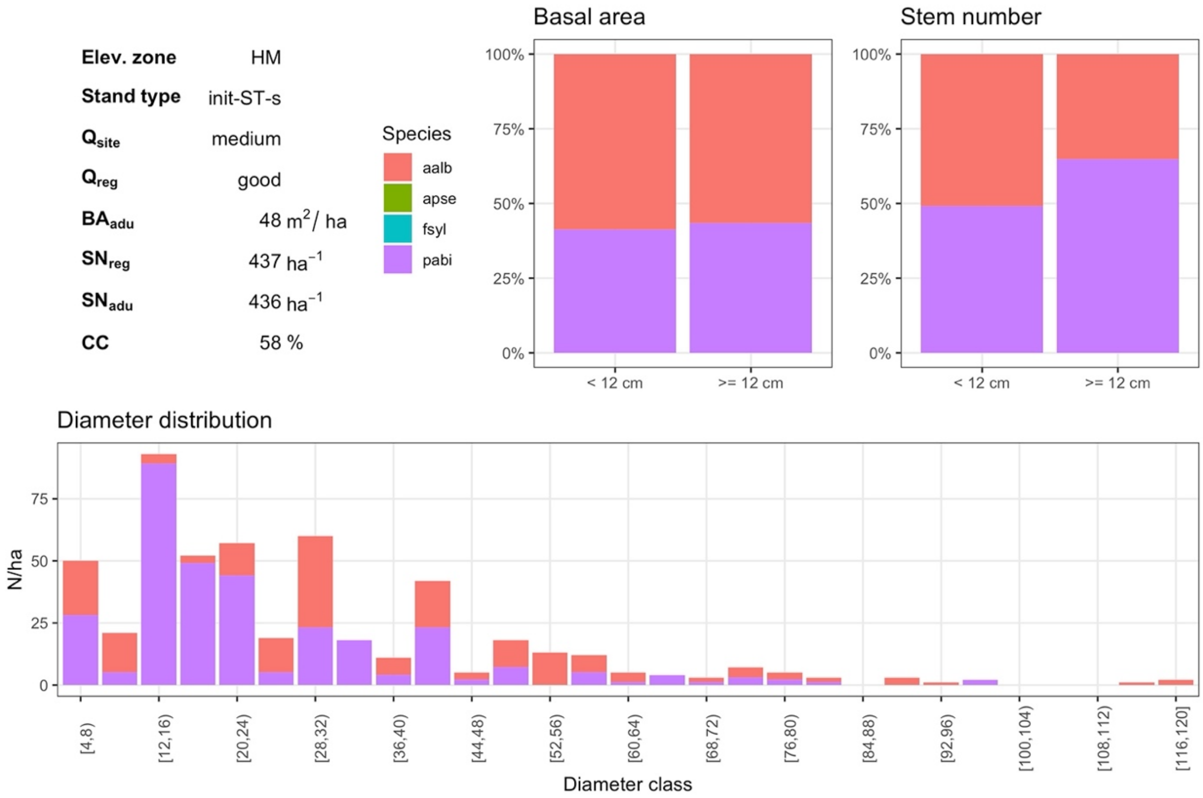


***Fig. S3.23*** *HM init-ST-s with medium site quality (3) and good regeneration quality (3)*

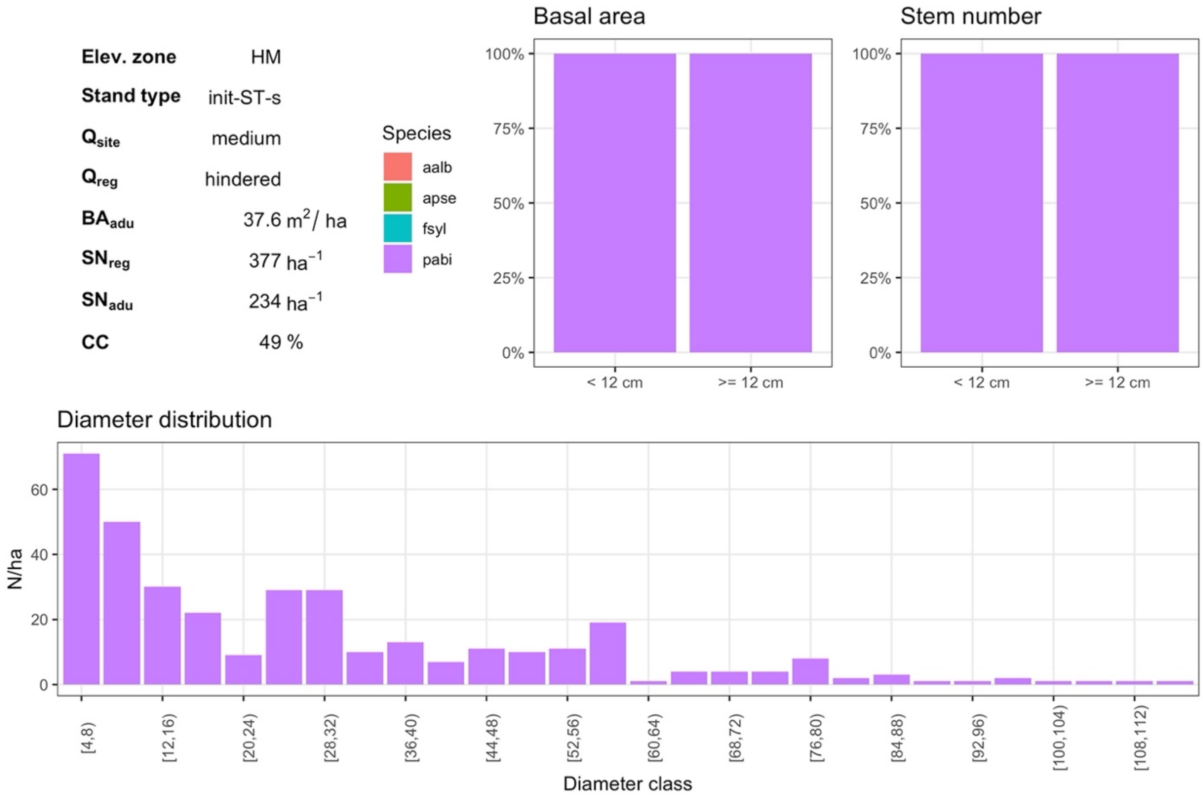


***Fig. S3.24*** *HM init-ST-s with medium site quality (3) and hindered regeneration quality (1)*

#### 2.3. Mature stand (init-ST-m)

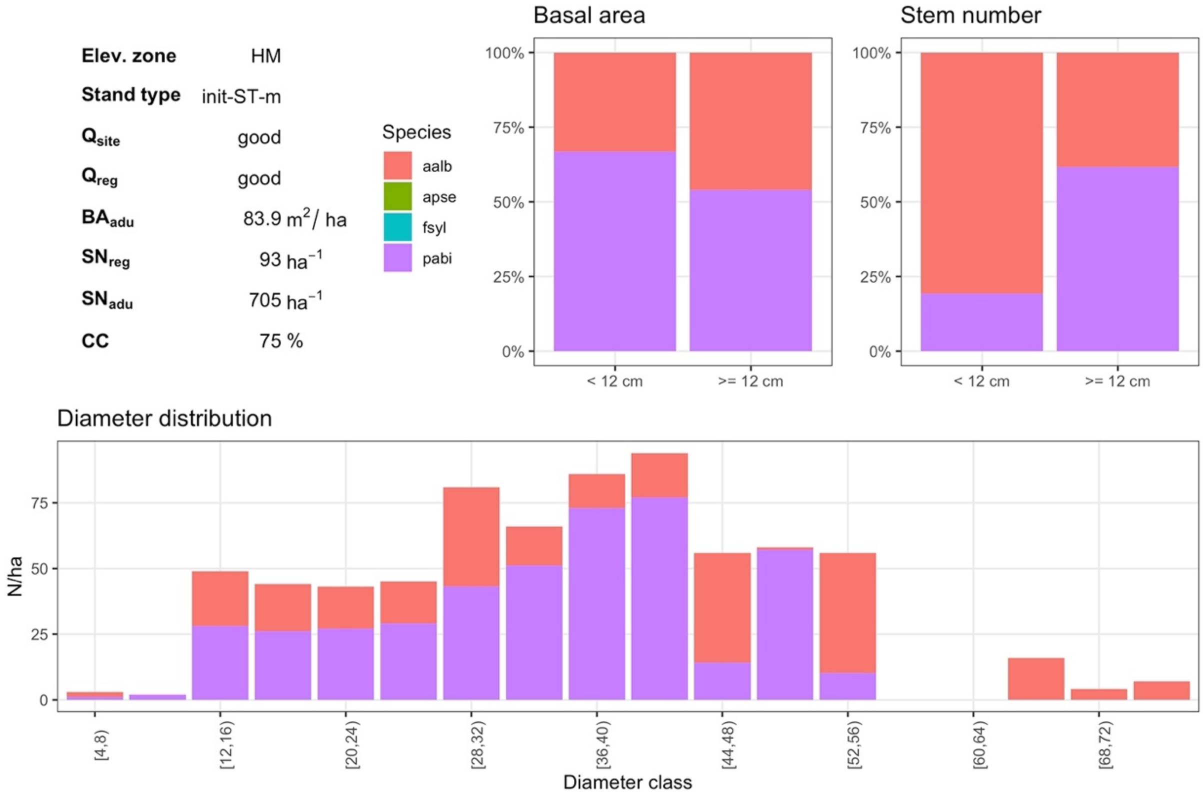


***Fig. S3.25*** *HM init-ST-m with good site quality (5) and good regeneration quality (5)*

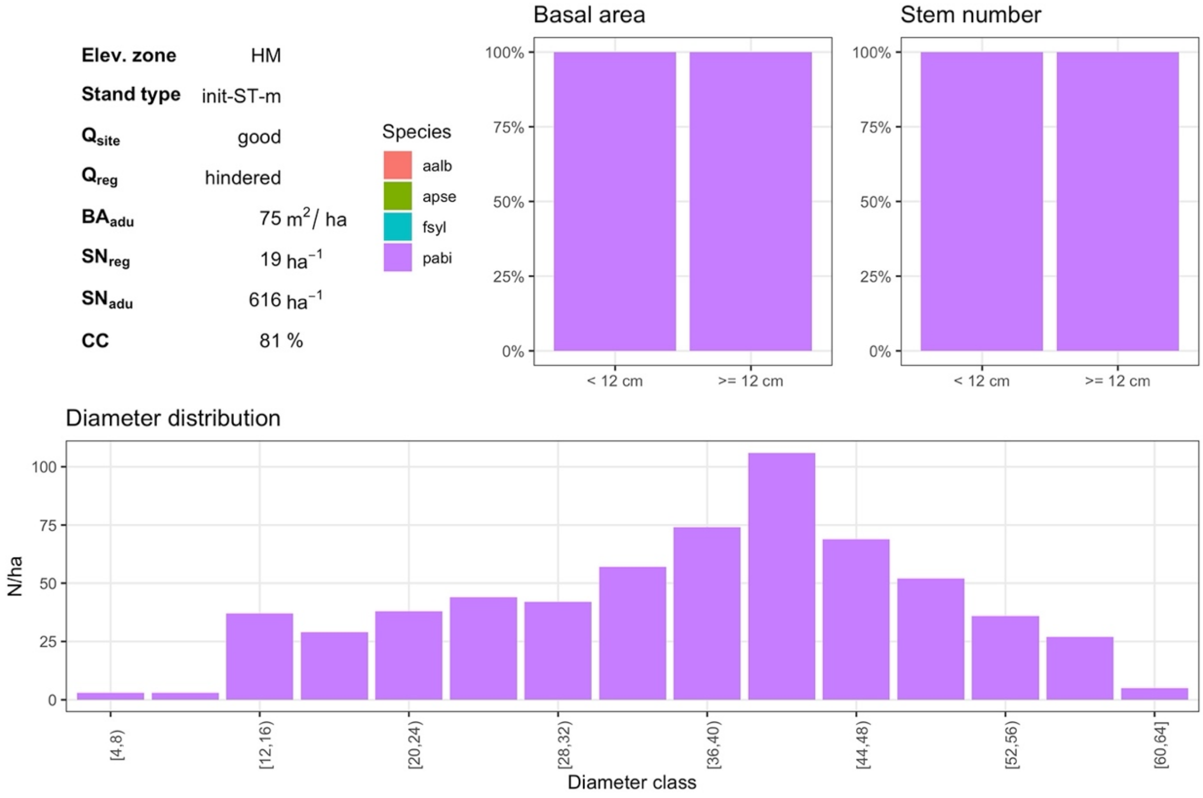


***Fig. S3.26*** *HM init-ST-m with good site quality (5) and hindered regeneration quality (1)*

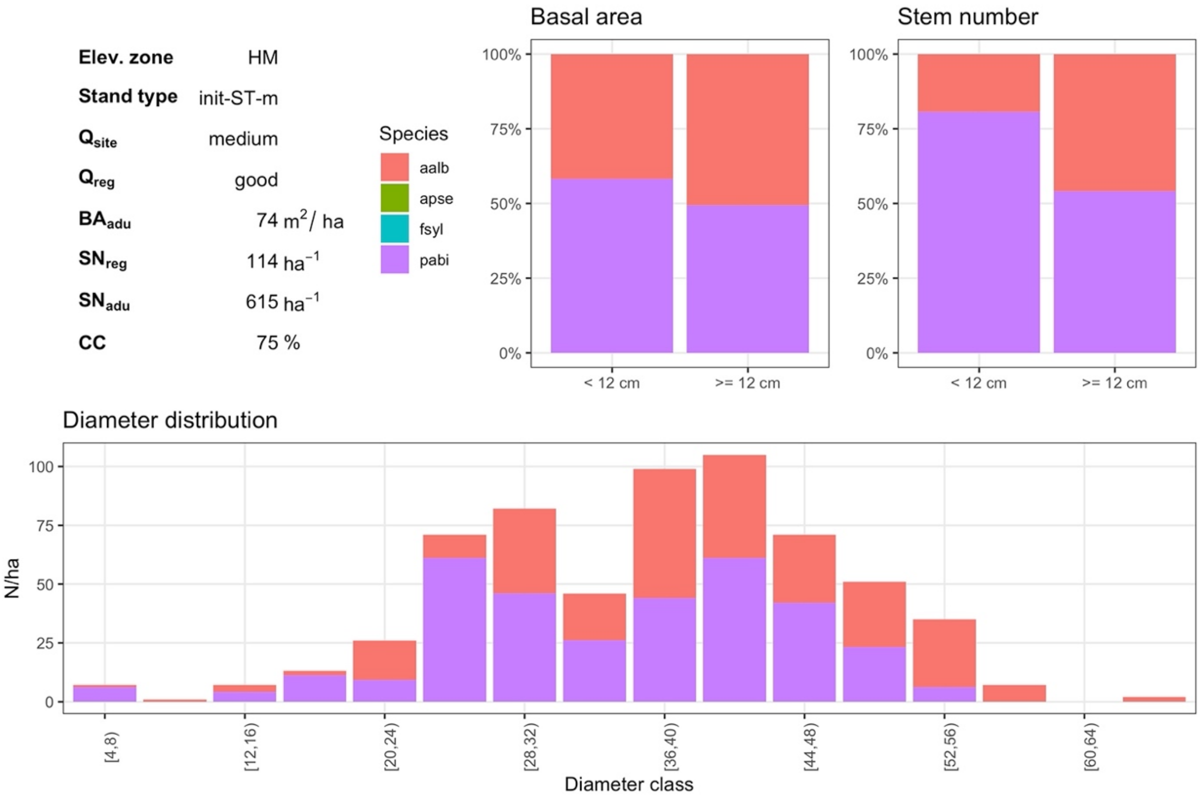


***Fig. S3.27*** *HM init-ST-m with medium site quality (3) and good regeneration quality (3)*

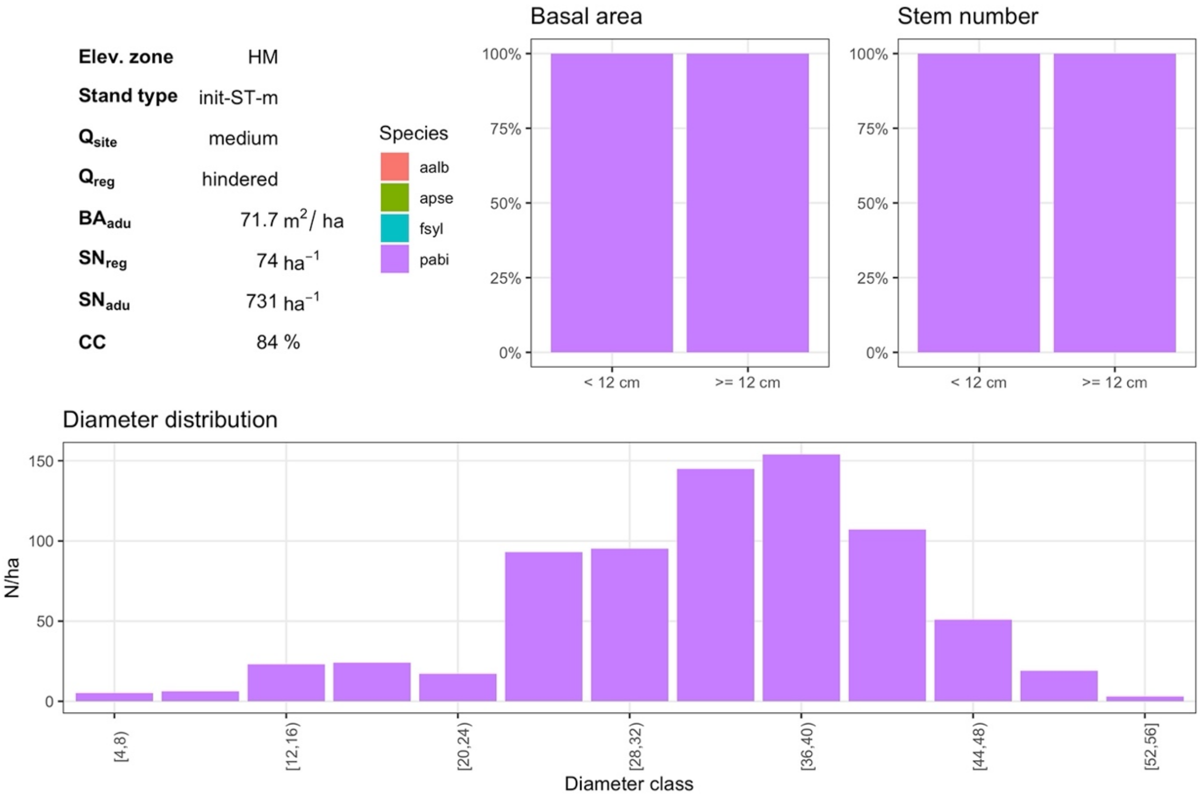


***Fig. S3.28*** *HM init-ST-m with medium site quality (3) and hindered regeneration quality (1)*

#### 2.4. Long-term simulation (init-LT)

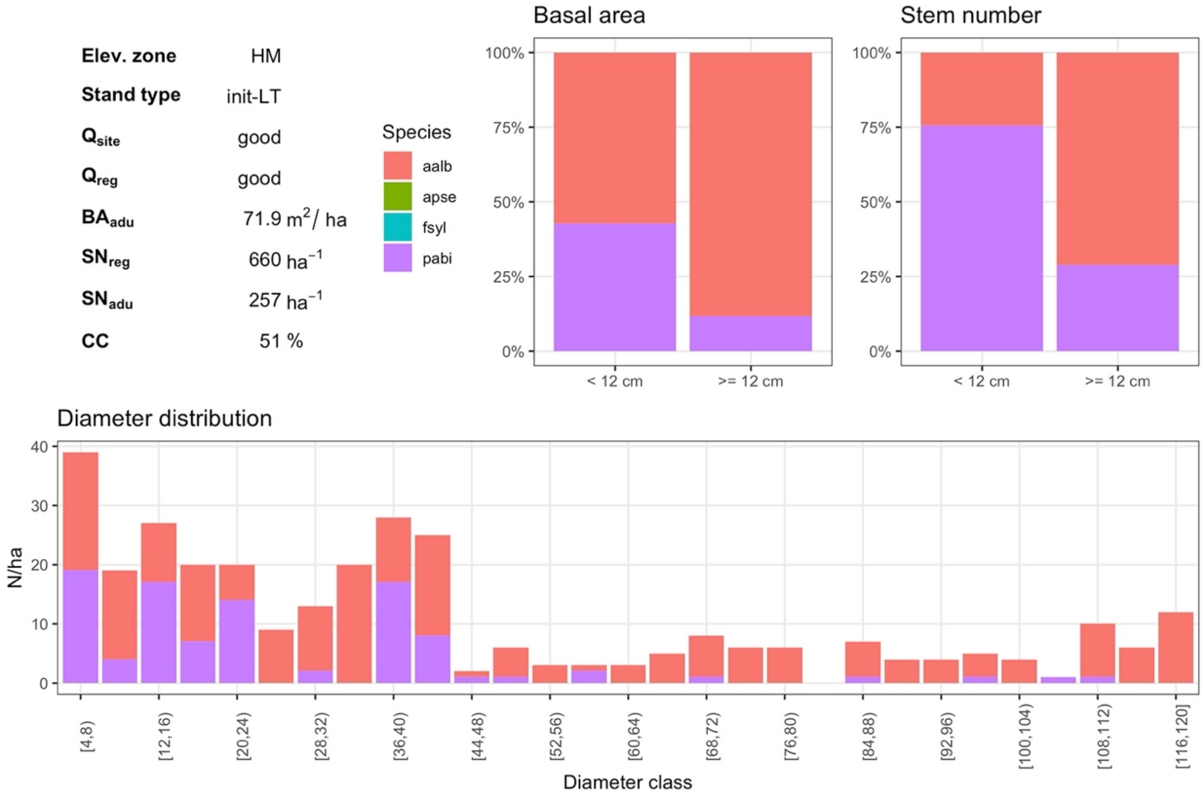


***Fig. S3.29*** *HM init-LT with good site quality (5) and good regeneration quality (5)*

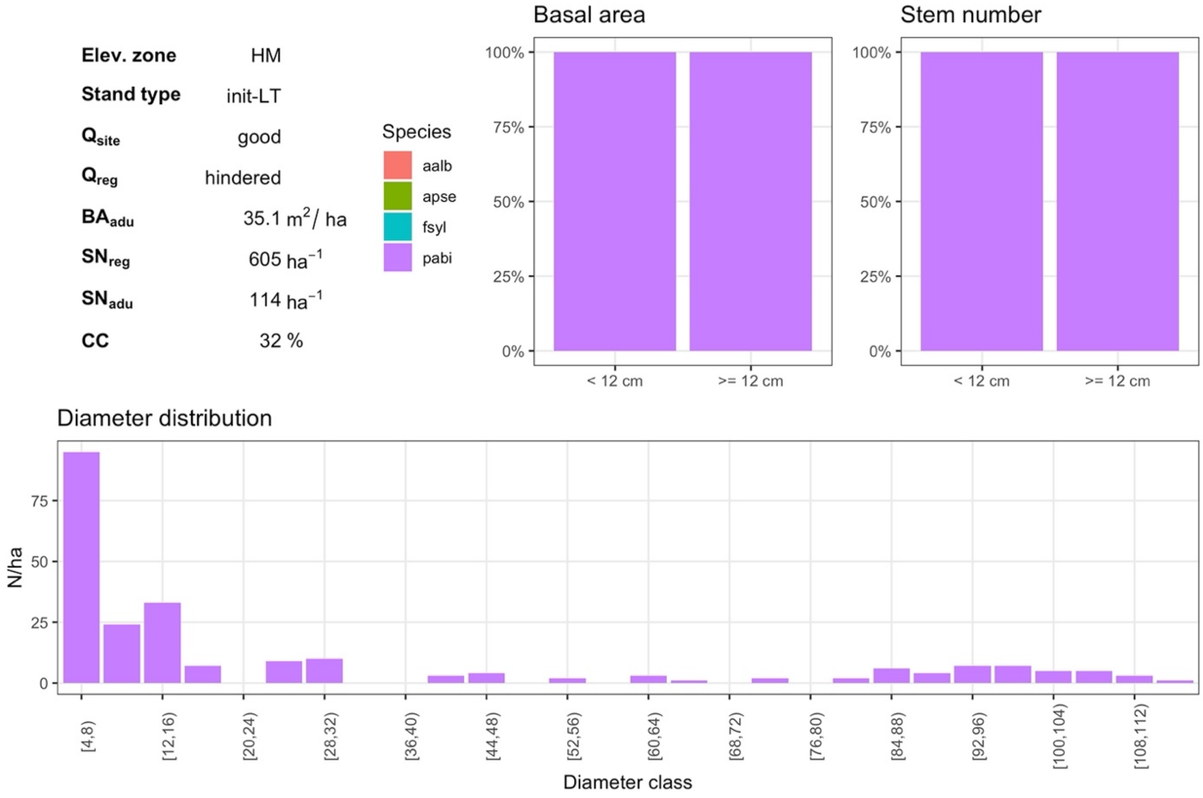


***Fig. S3.30*** *HM init-LT with good site quality (5) and hindered regeneration quality (1)*

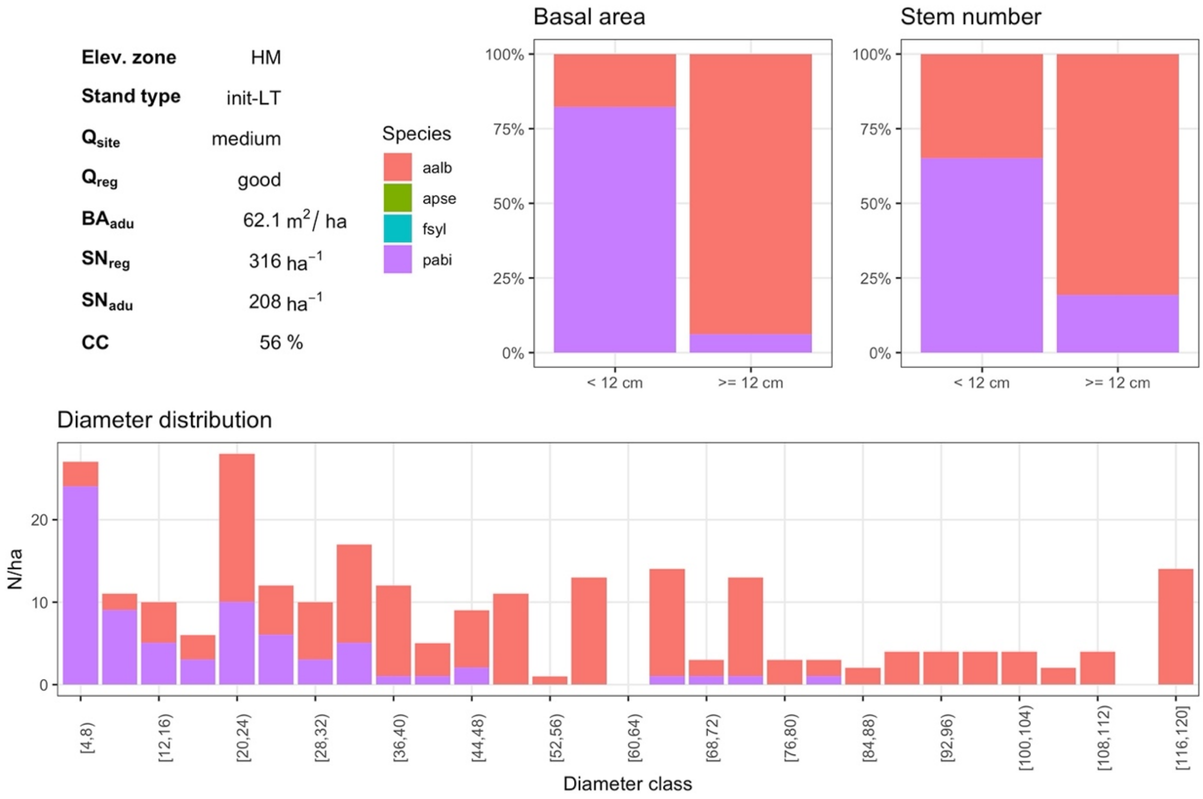


***Fig. S3.31*** *HM init-LT with medium site quality (3) and good regeneration quality (3)*

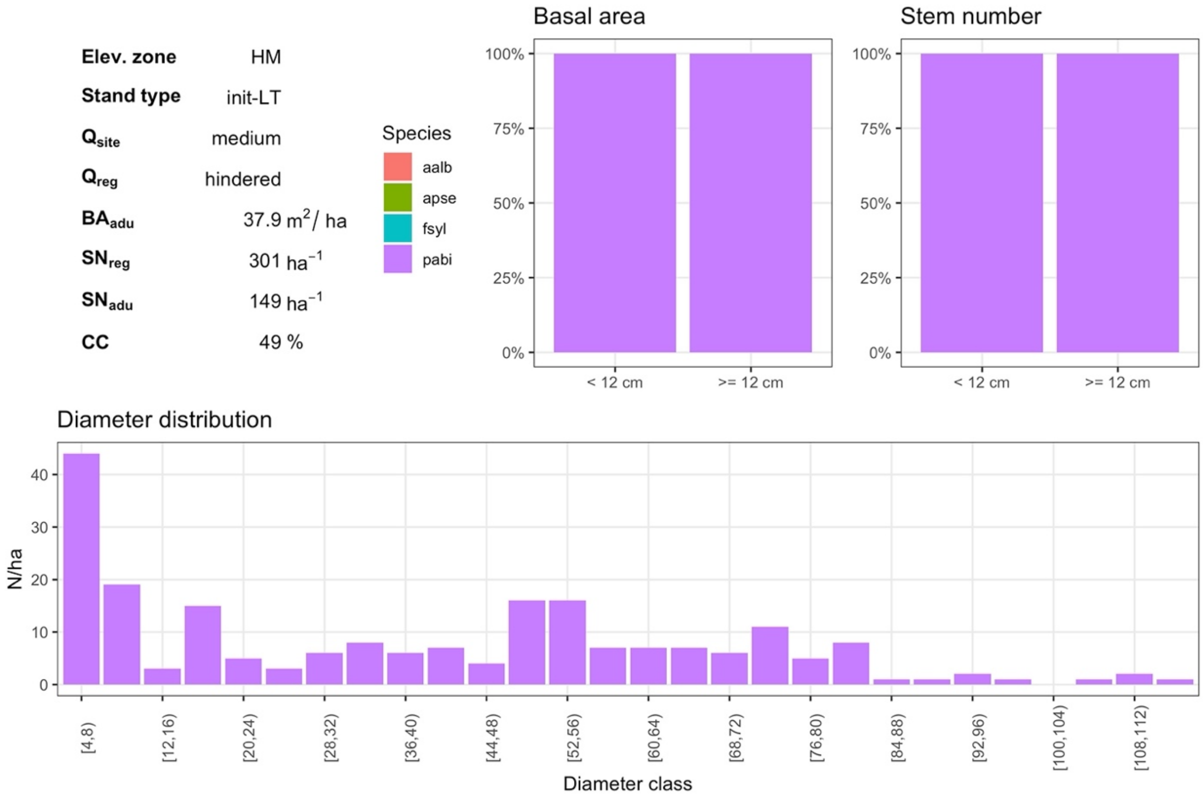


***Fig. S3.32*** *HM init-LT with medium site quality (3) and hindered regeneration quality (1)*

### 3. Subalpine zone

#### 3.1. Young stand (init-ST-y)

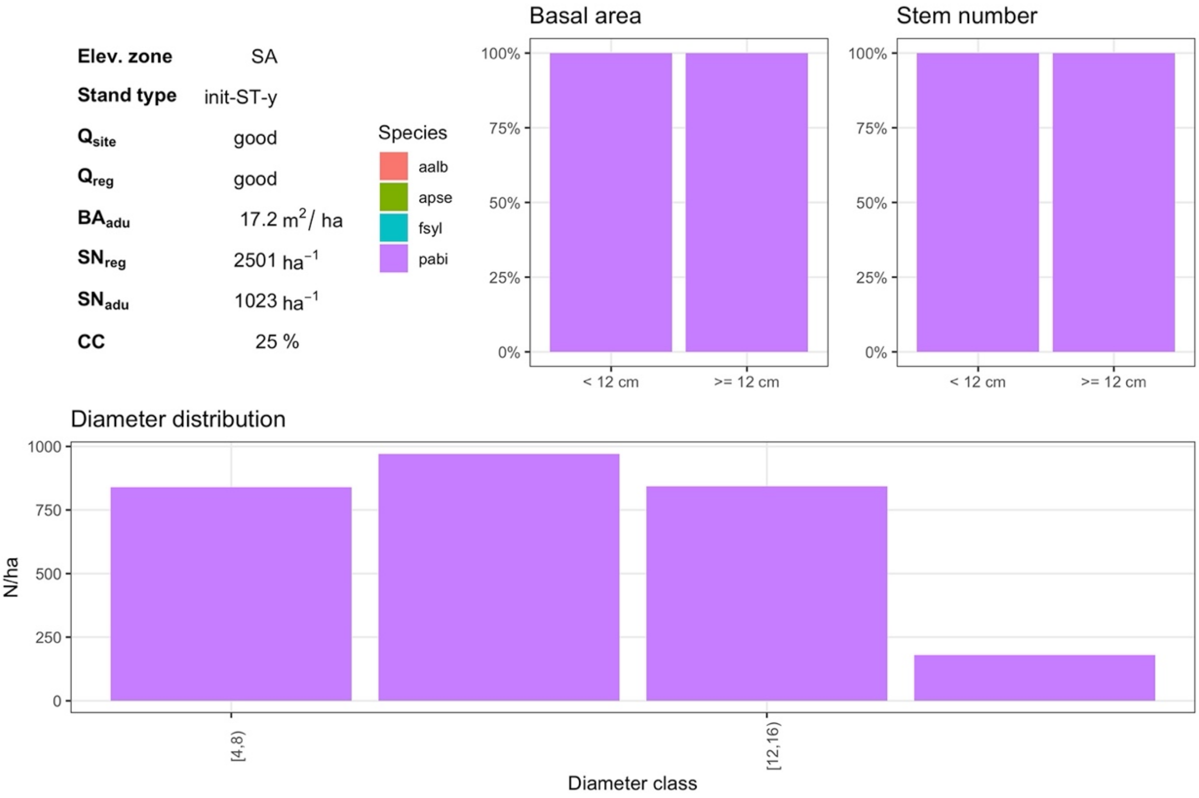


***Fig. S3.33*** *SA init-ST-y with good site quality (5) and good regeneration quality (5)*

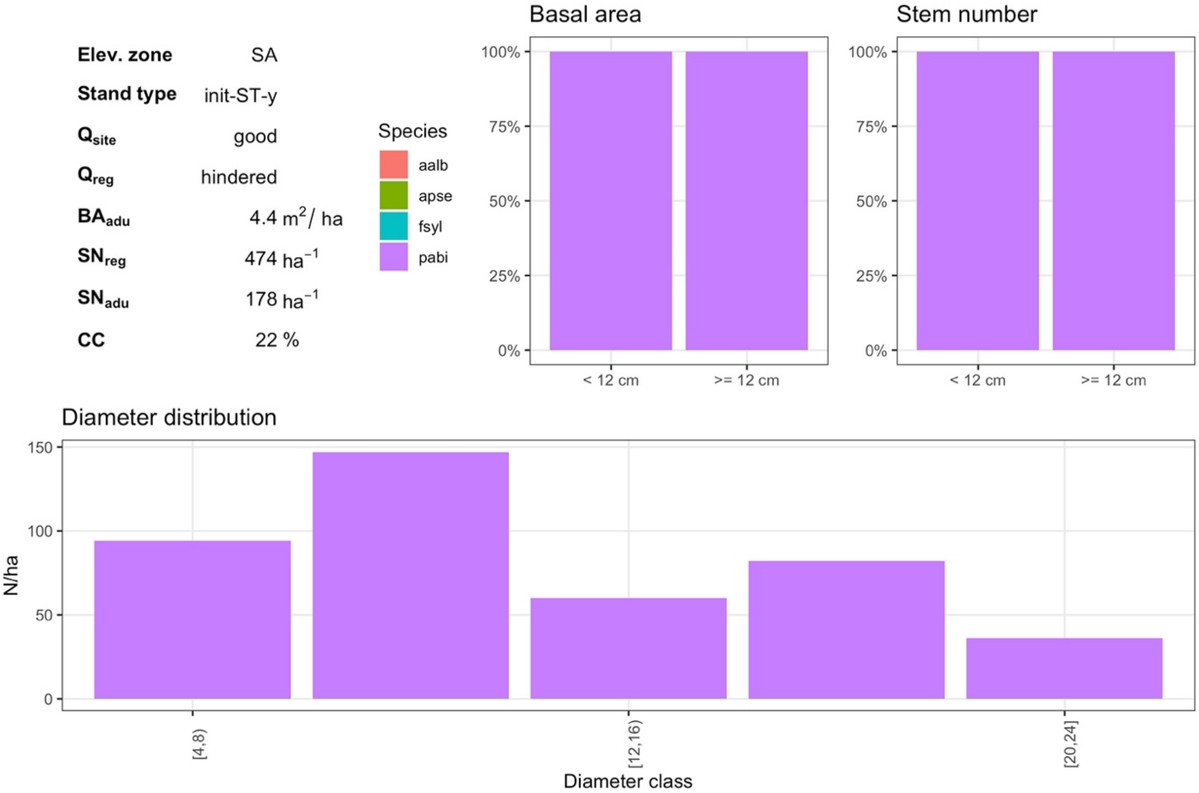


***Fig. S3.34*** *SA init-ST-y with good site quality (5) and hindered regeneration quality (1)*

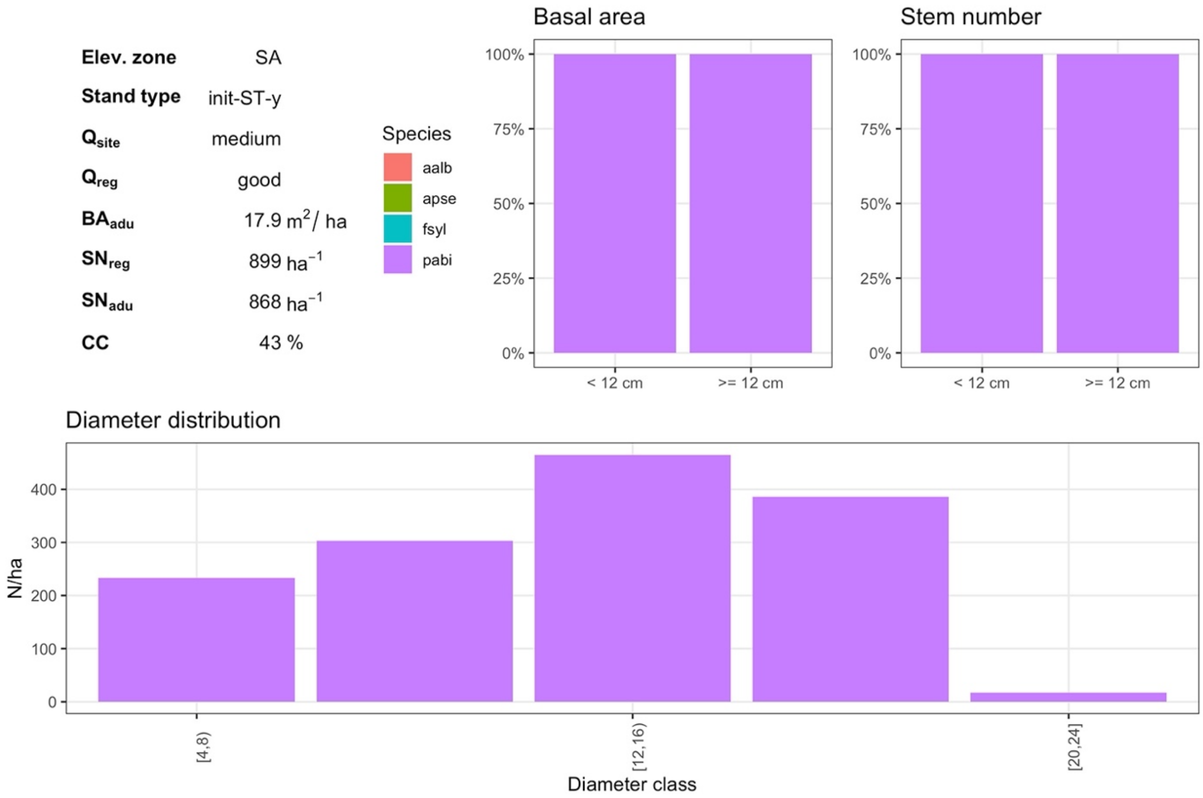


***Fig. S3.35*** *SA init-ST-y with medium site quality (3) and good regeneration quality (3)*

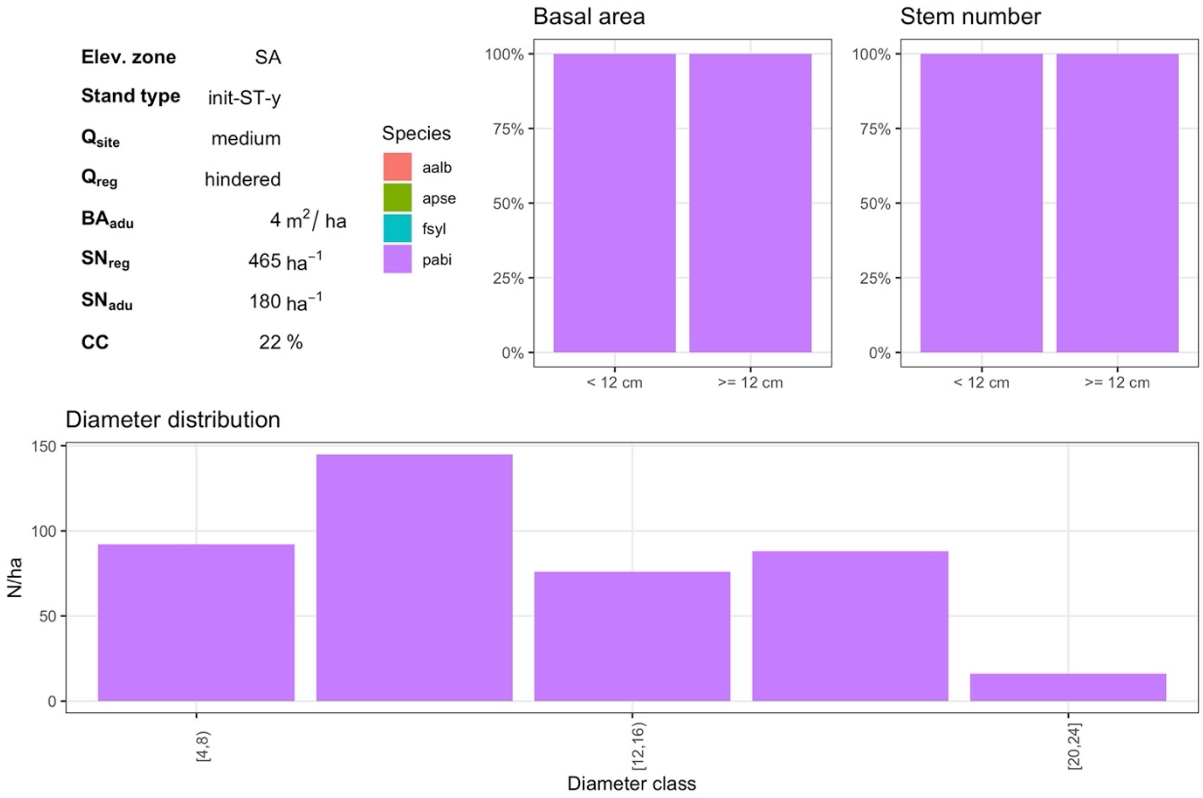


***Fig. S3.36*** *SA init-ST-y with medium site quality (3) and hindered regeneration quality (1)*

#### 3.2. Well-structured stand (init-ST-s)

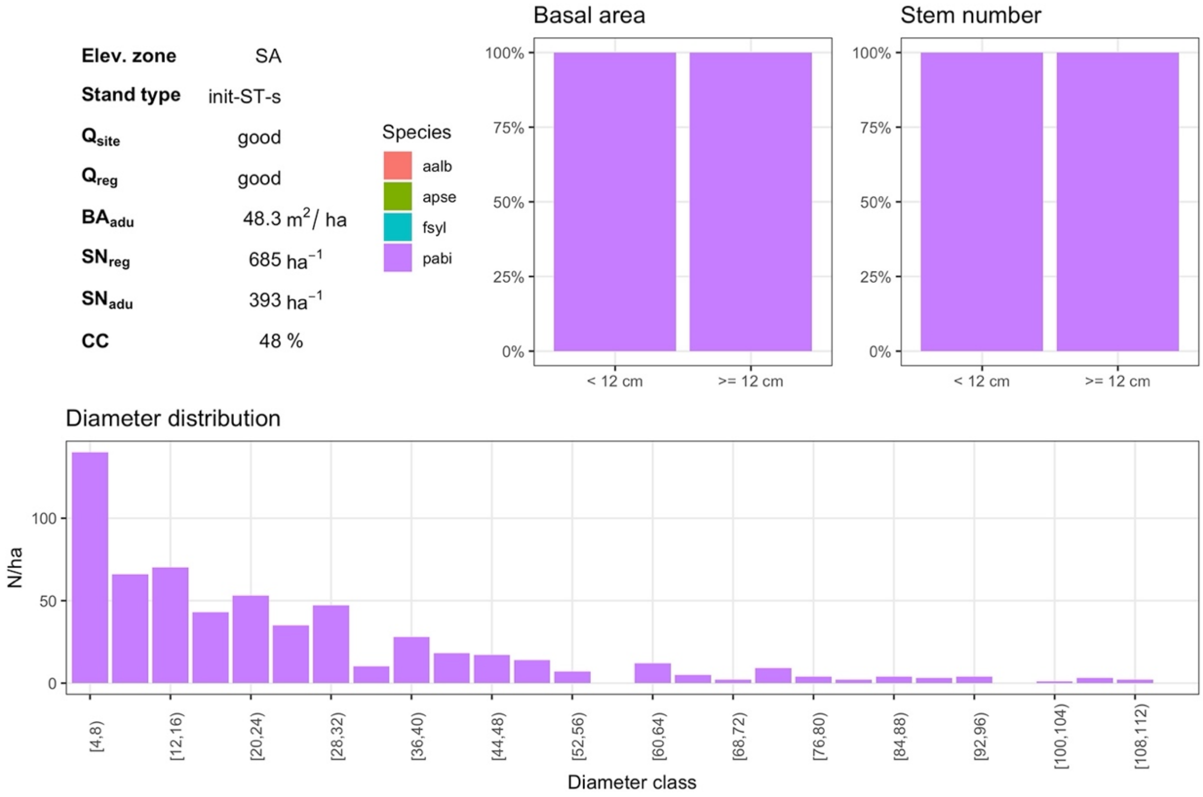


***Fig. S3.37*** *SA init-ST-s with good site quality (5) and good regeneration quality (5)*

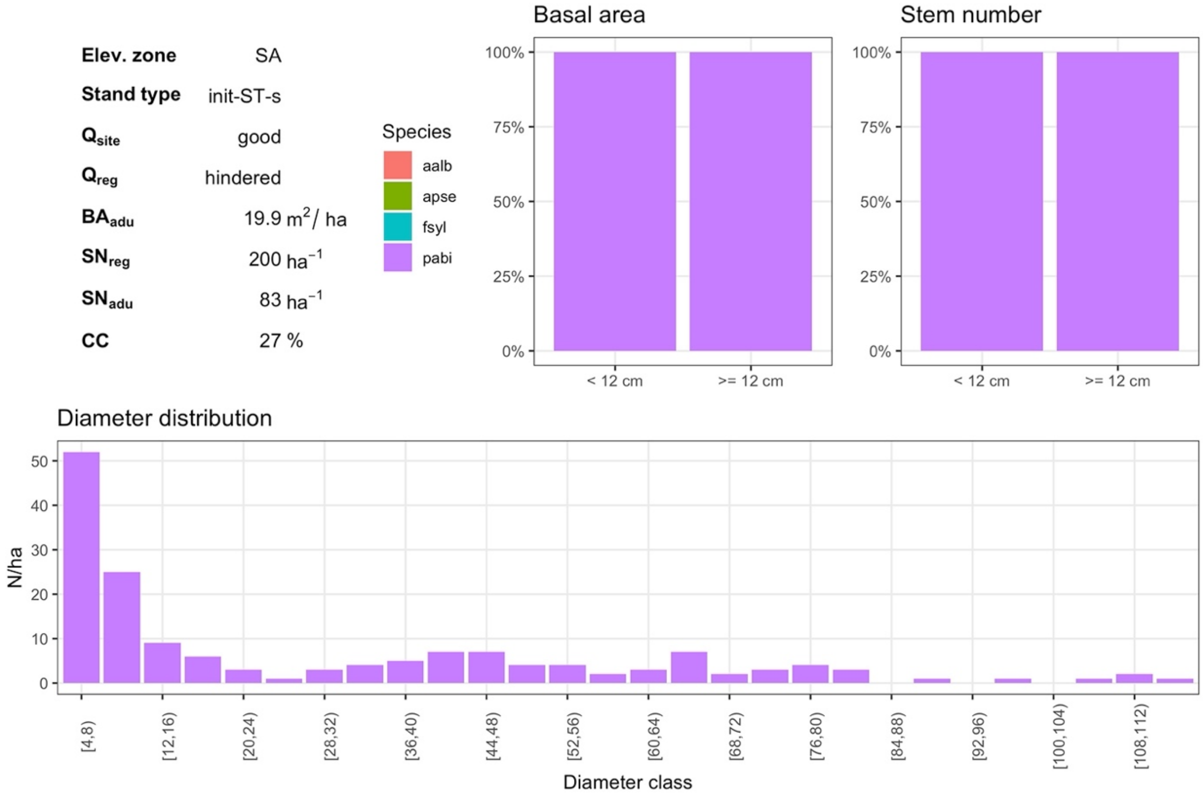


***Fig. S3.38*** *SA init-ST-s with good site quality (5) and hindered regeneration quality (1)*

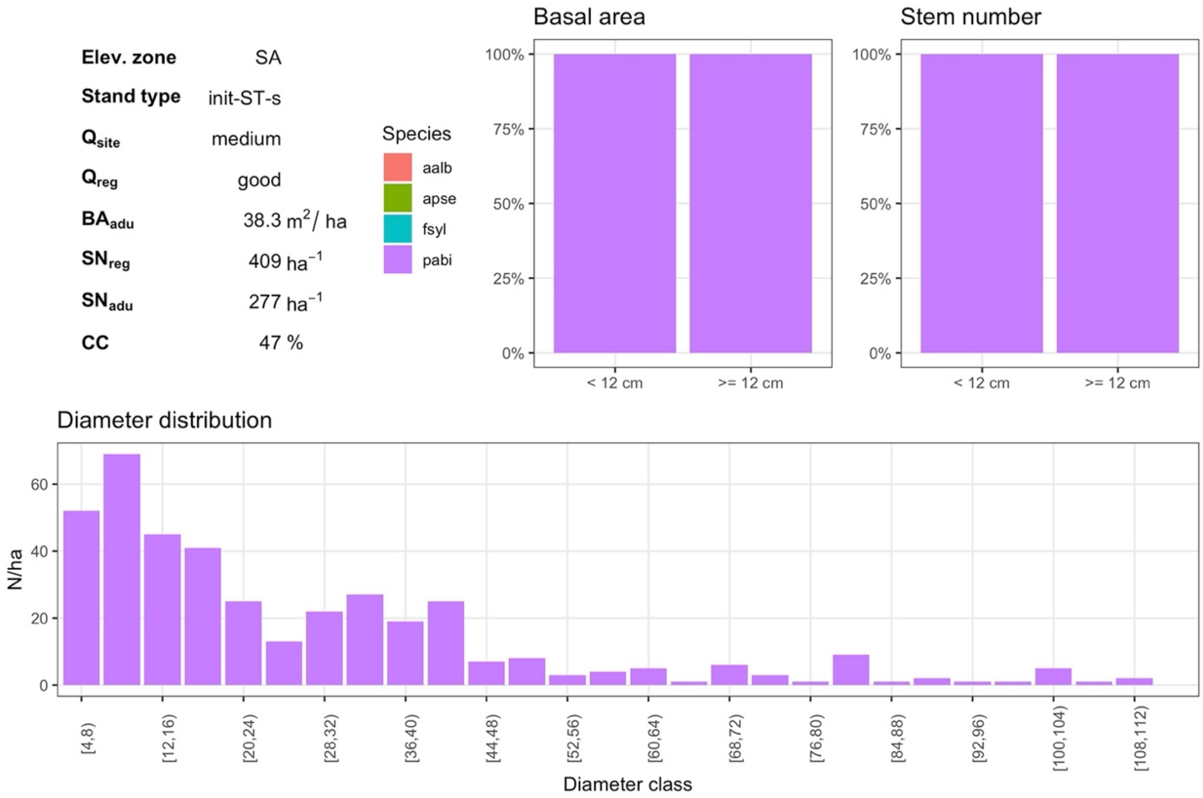


***Fig. S3.39*** *SA init-ST-s with medium site quality (3) and good regeneration quality (3)*

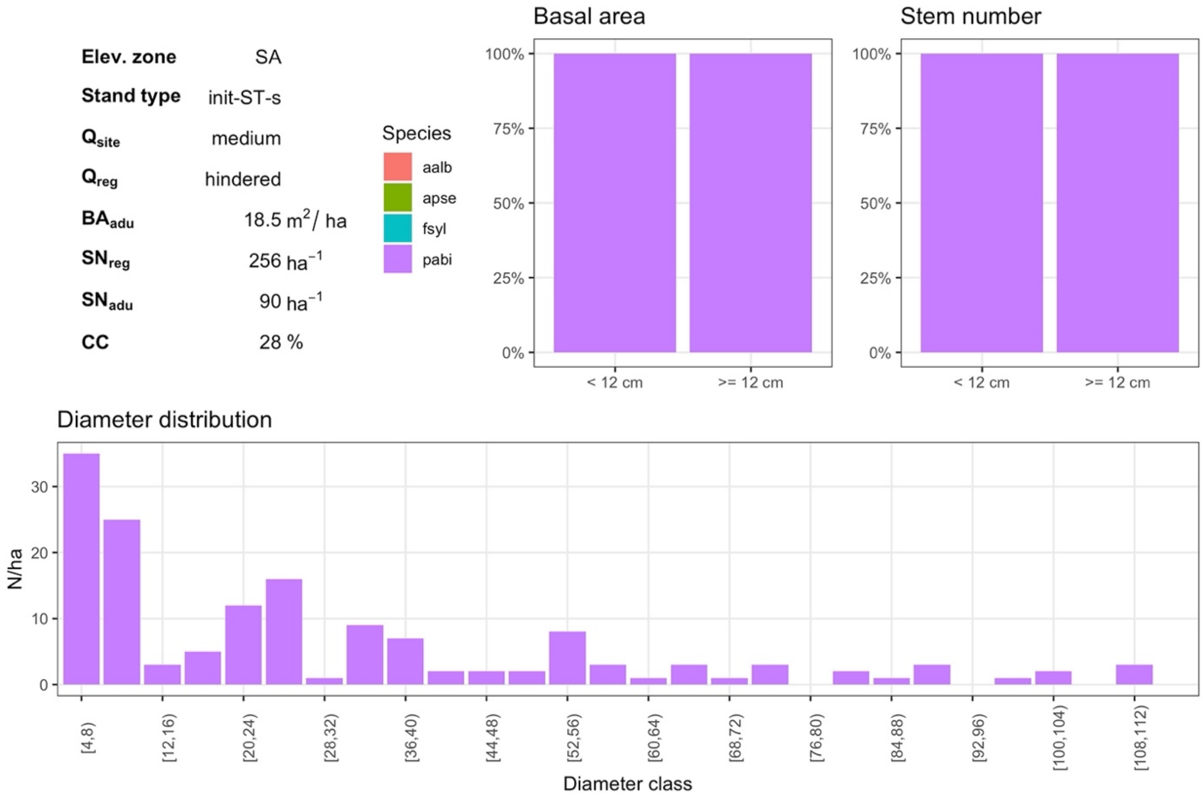


***Fig. S3.40*** *SA init-ST-s with medium site quality (3) and hindered regeneration quality (1)*

#### 3.3. Mature stand (init-ST-m)

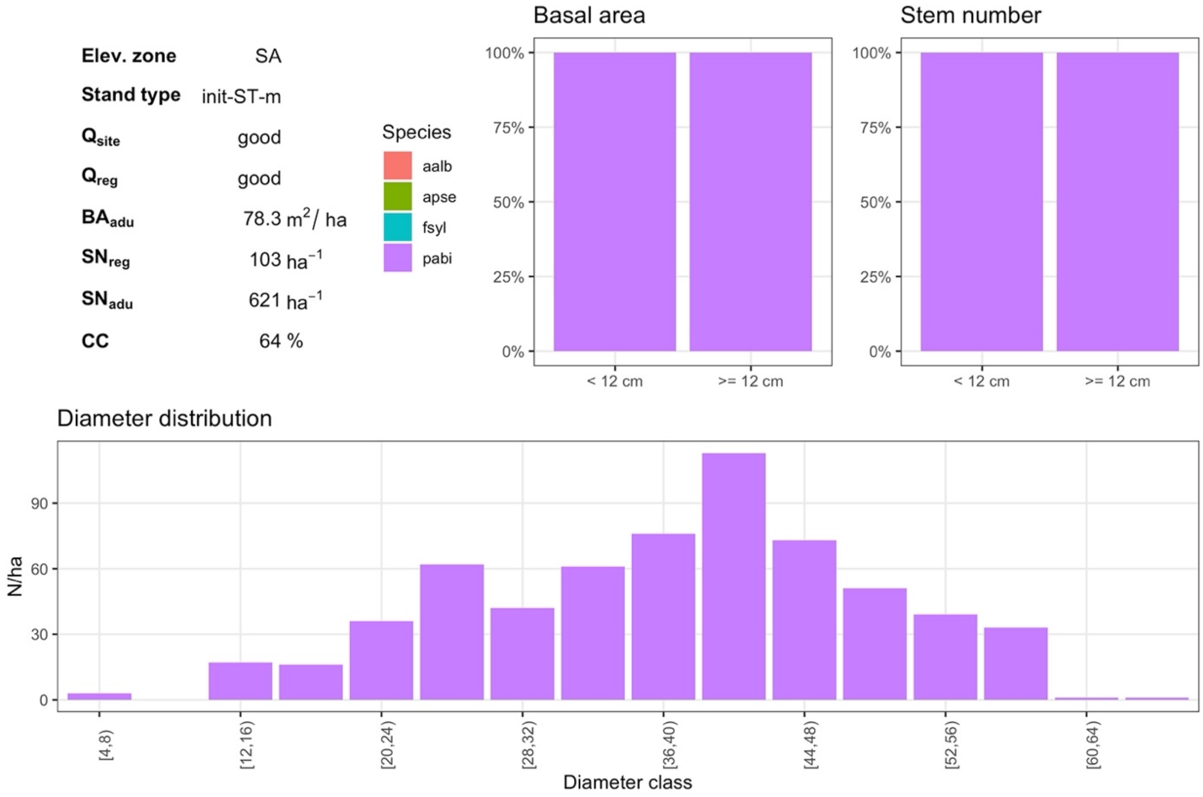


***Fig. S3.41*** *SA init-ST-m with good site quality (5) and good regeneration quality (5)*

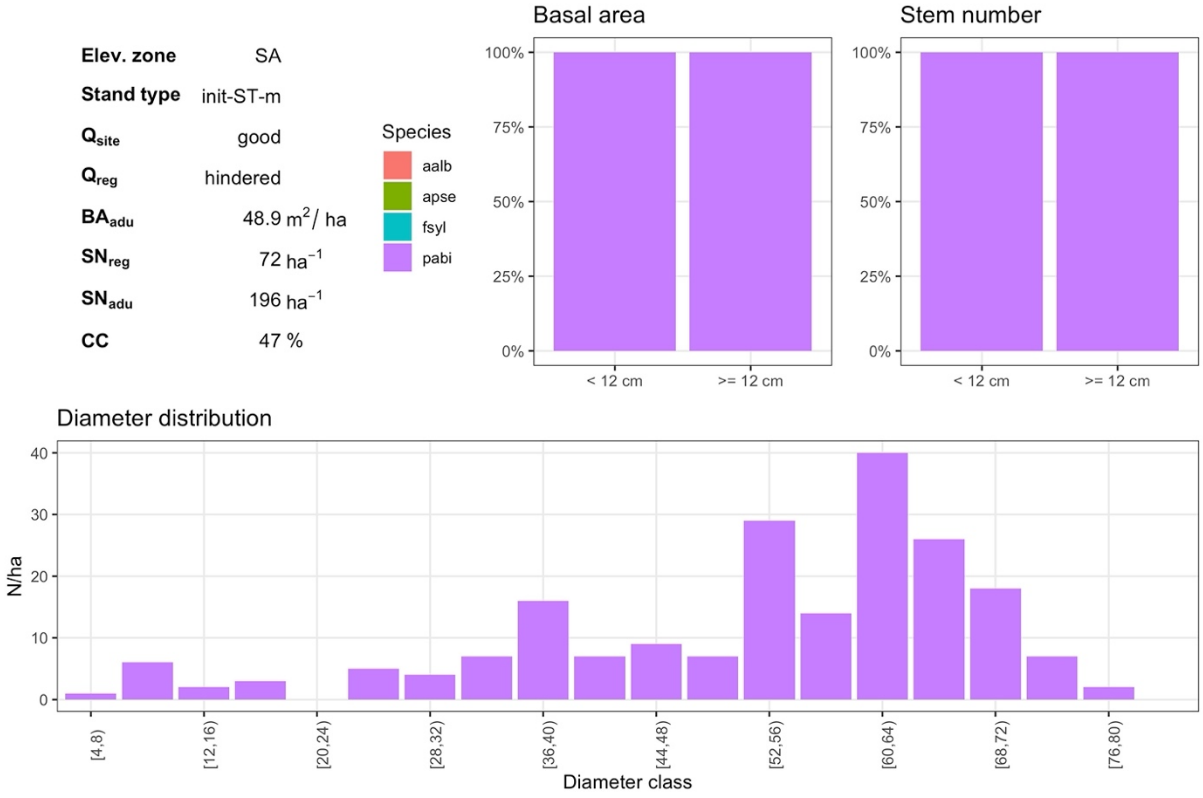


***Fig. S3.42*** *SA init-ST-m with good site quality (5) and hindered regeneration quality (1)*

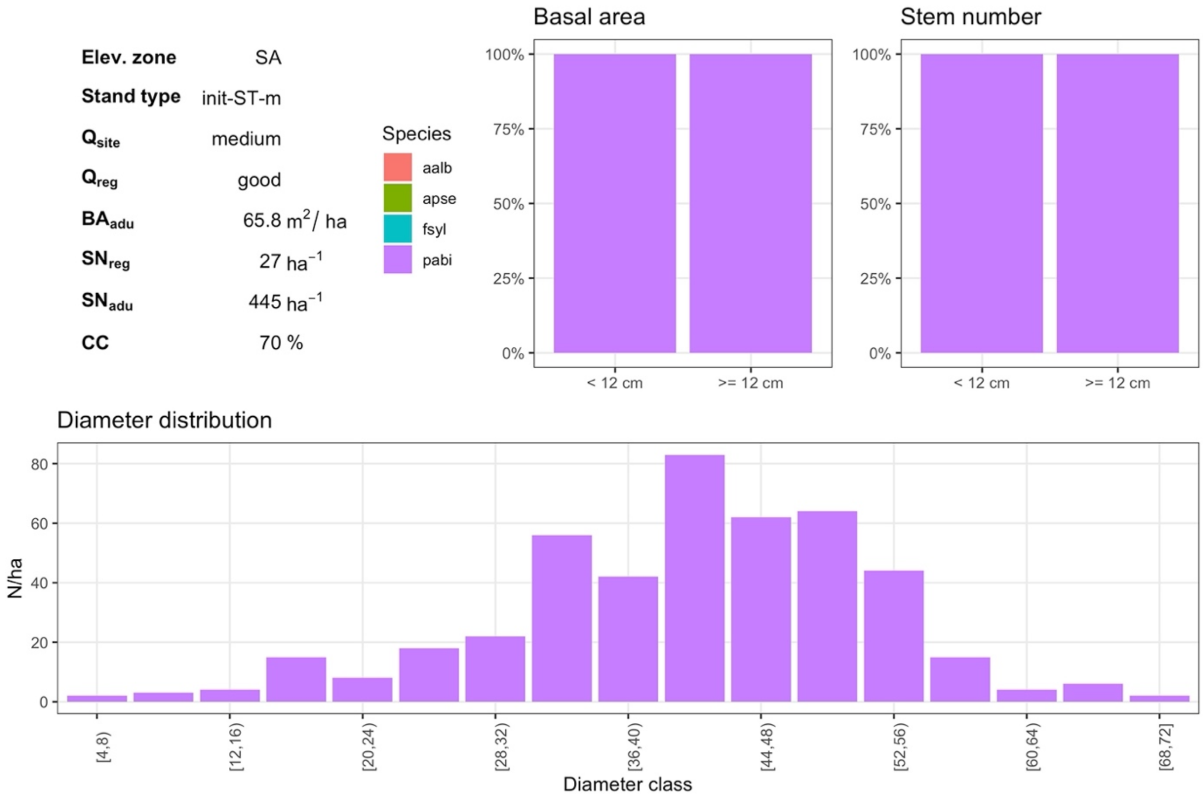


***Fig. S3.43*** *SA init-ST-m with medium site quality (3) and good regeneration quality (3)*

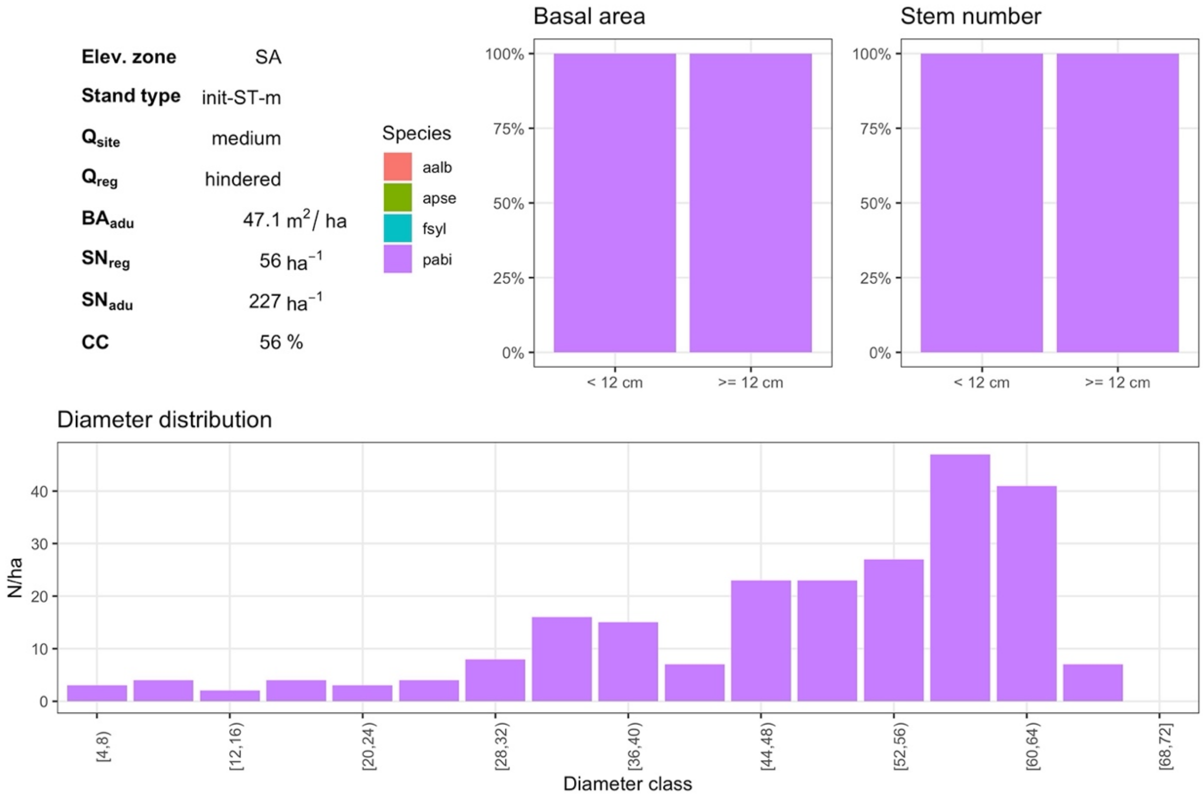


***Fig. S3.44*** *SA init-ST-m with medium site quality (3) and hindered regeneration quality (1)*

#### 3.4. Long-term simulation (init-LT)

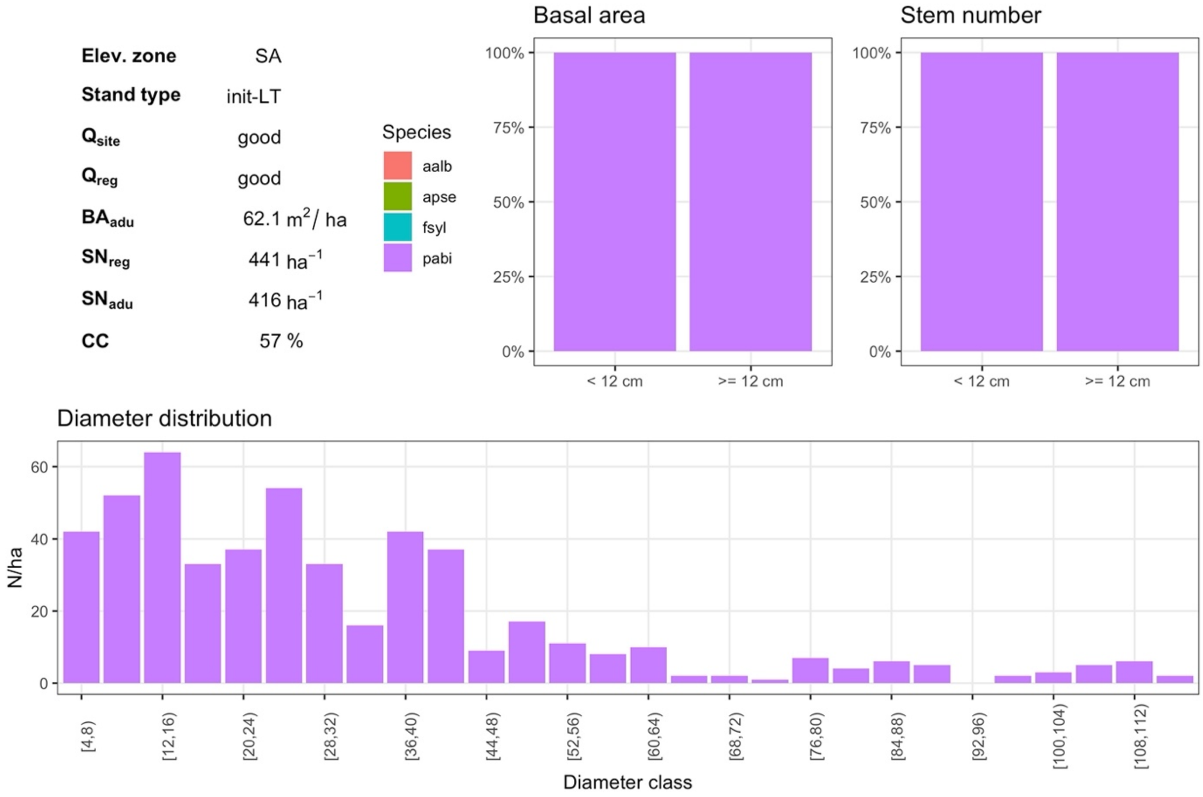


***Fig. S3.45** SA init-LT with good site quality (5) and good regeneration quality (5)*

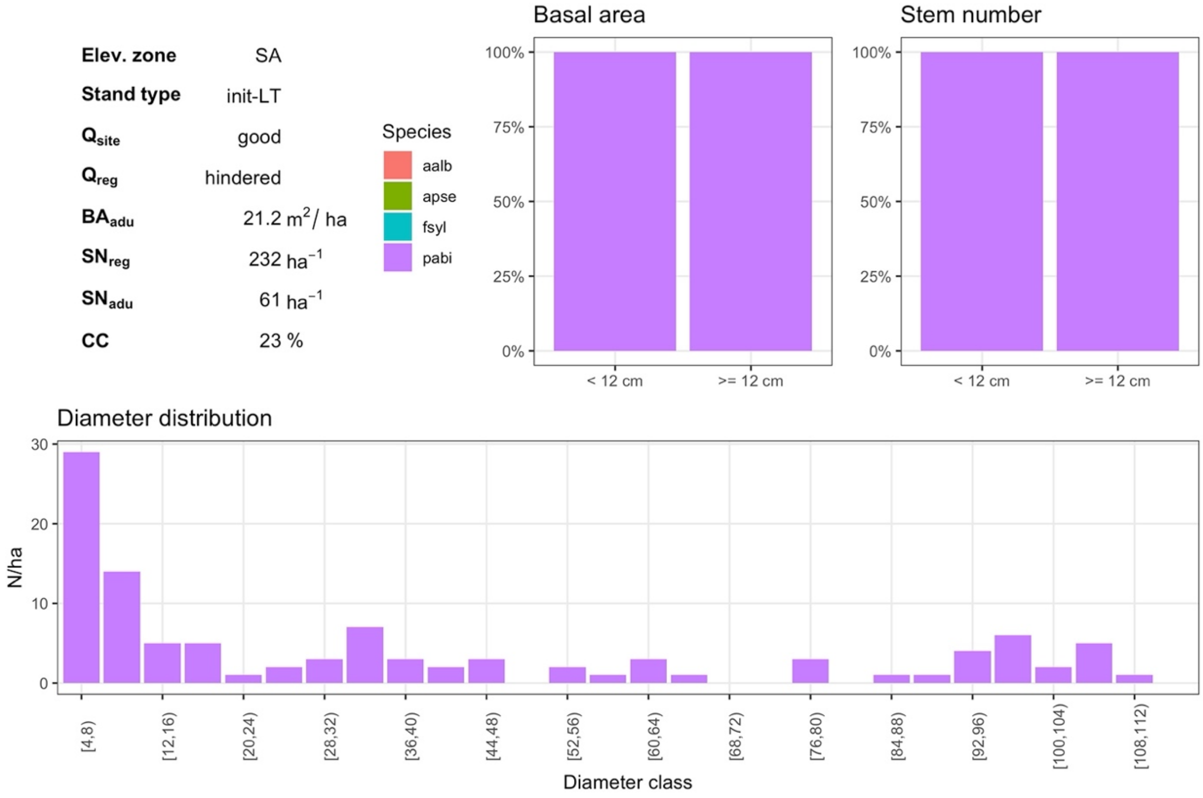


***Fig. S3.46** SA init-LT with good site quality (5) and hindered regeneration quality (1)*

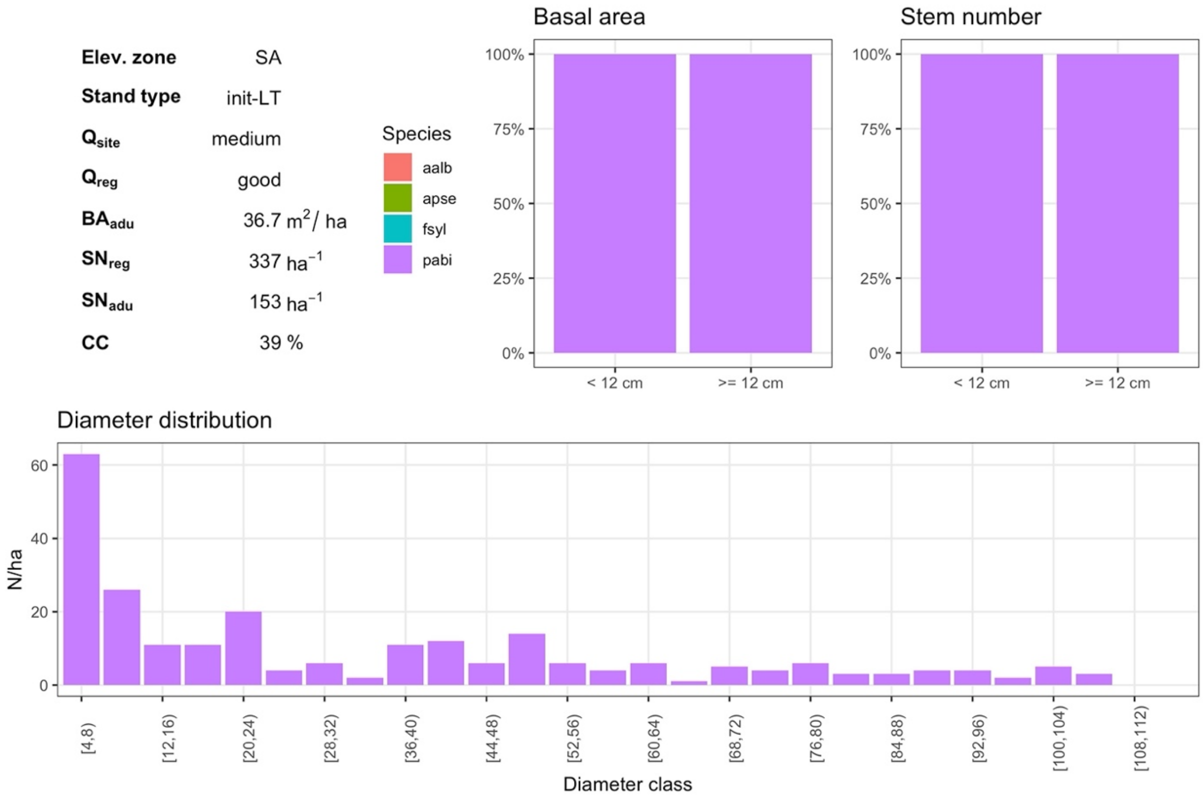


***Fig. S3.47*** *SA init-LT with medium site quality (3) and good regeneration quality (3)*

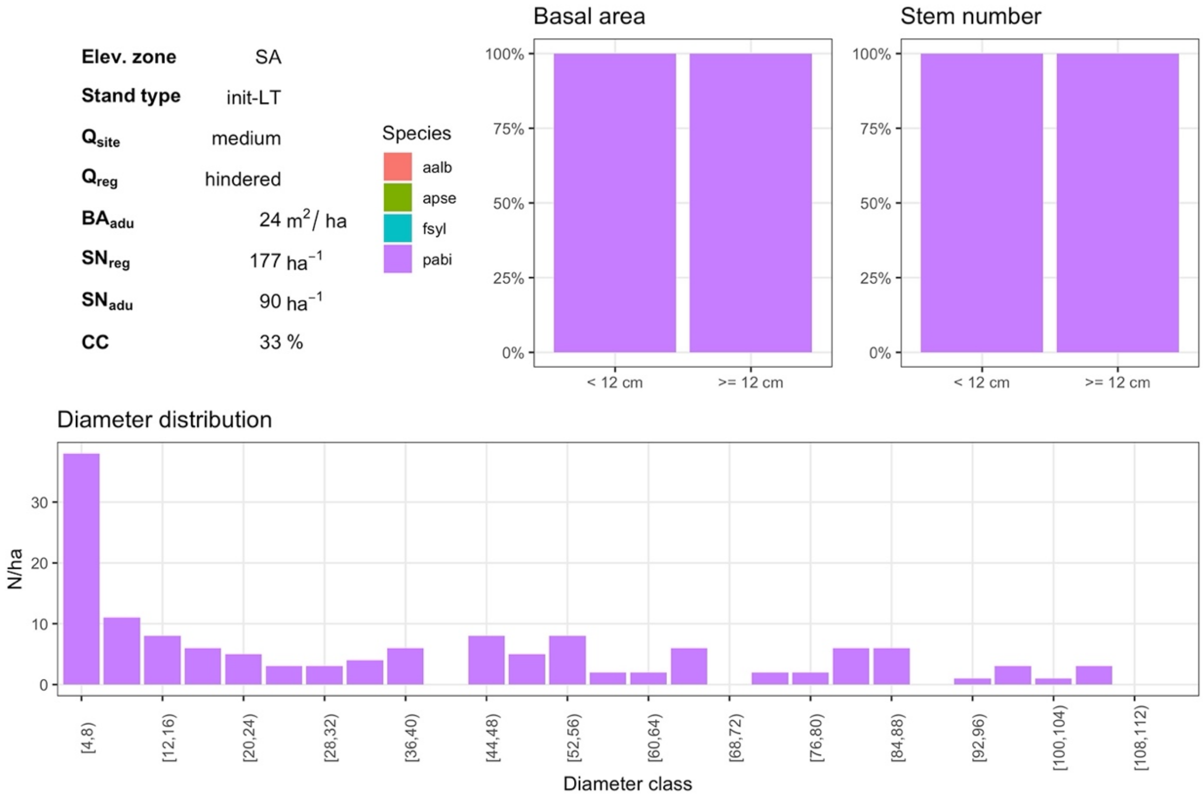


***Fig. S3.48*** *SA init-LT with medium site quality (3) and hindered regeneration quality (1)*
