## Supplementary material S4 for "Management recommendations for Alpine protection forests: the importance of regeneration quality and initial stand composition"

### Supplementary material S4: Management settings

S4 describes the management settings characterizing the management regimes. For a detailed description of the management algorithms and the parameters, cf. Supplementary material S1.

#### 1. Shared management parameters

The settings for the six management types (STS, GRS1, GRS2, CAB1, CAB2, and SC) shared certain parameters (Table S4.1).

***Table S4.1*** *Management settings applied to all six management types.*

| Parameter | Explanation | Value |
|---|---|---|
| $interv_{first}$ | Time step of the first intervention | 1 |
| $interv_{interval}$ | Interval between intervention in years | 10, 20, 30, 40 |
| $interv_{BAred}$ | Share of total basal area reduction per intervention | 0.1, 0.2, 0.3, 0.4 |
| $interv_{Dmin}$ | Minimal DBH in cm of trees eligible for harvest | 12 |
| $BAmin_i$ | Share of stand basal area per species at the time of management, below which they are not eligible for harvest | NaiS profile thresholds for ideal profile (differs between elevational zones) |
| $mgmunit_{buff}$ | Initial buffer width in number of cells to be left unmanaged around interventions | 1 |

#### 2. Management regime STS

The management regime STS used management type STS (single tree selection). The applied weighting factors are listed in Table S4.2.

***Table S4.2*** *Weighting factors used in management regime STS.*

| Parameter | Explanation | Value |
|---|---|---|
| $mgmw_{spc}$ | Weight given to tree species | 10 |
| $mgmw_{BA}$ | Weight given to basal area | 100 |
| $mgmw_{CR}$ | Weight given to crown ratio | 40 |

#### 3. Management regimes GRS1 and GRS2

The management regimes GRS1 and GRS2 used management type GRS (group selection). The only difference between the two regimes was the group size ($mgmunit_{agg}$) which was two cells in GRS1 and four cells in GRS2. The applied weighting factors are listed in Table S4.3.

***Table S4.3*** *Weighting factors used in management regimes GRS1 and GRS2.*

| Parameter | Explanation | Value |
|---|---|---|
| $mgmw_{spc}$ | Weight given to tree species | 10 |
| $mgmw_{BA}$ | Weight given to basal area | 100 |
| $mgmw_{CR}$ | Weight given to crown ratio | 20 |
| $mgmw_{RegUnit}$ | Weight given to regeneration within harvest unit | 40 |
| $mgmw_{RegNeighb}$ | Weight given to regeneration in cells neighboring harvest unit | 20 |

#### 4. Management regimes CAB1 and CAB2

The management regimes CAB1 and CAB2 used management type CAB (cable yarding). The applied weighting factors are listed in Table S4.4.

***Table S4.4*** *Weighting factors used in management regimes CAB1 and CAB2.*

| Parameter | Explanation | Value |
|---|---|---|
| $mgmw_{BA}$ | Weight given to basal area | 100 |
| $mgmw_{RegUnit}$ | Weight given to regeneration within harvest unit | 40 |
| $mgmw_{RegNeighb}$ | Weight given to regeneration in cells neighboring harvest unit | 20 |

The cable layout in CAB1 contained three cable lines with a distance of approximately 30 m in between (28 to 33 m, depending on the elevational zone). The parameter values for the spatial layout are listed in Table S4.5. The simulation grid dimensions were 10x10 cells in UM, 14x14 cells in HM, and 16x16 cells in SA. The resulting slit lengths were between 9 and 13 m and the slit widths between 13 and 22 m. In UM and SA, one column of cells was left unmanaged, in HM, one column was harvested from two lines.

***Table S4.5*** *Spatial layout parameters used in management regime CAB1.*

| | | **Values** | | |
|---|---|---|---|---|
| **Parameter** | **Explanation** | **UM** | **HM** | **SA** |
| $mgmunit_{CL1}$ | Grid column with cable line 1 | 2 | 3 | 3 |
| $mgmunit_{CL2}$ | Grid column with cable line 2 | 5 | 8 | 8 |
| $mgmunit_{CL3}$ | Grid column with cable line 3 | 9 | 13 | 14 |
| $mgmunit_{SL}$ | Slit length in cells | 1 | 2 | 2 |
| $mgmunit_{SW}$ | Slit width in cells | 2 | 2 | 2 |

The cable layout in CAB2 contained two cable lines with a distance of 47 m in between. The parameter values for the spatial layout are listed in Table S4.6. The resulting slit lengths were between 17 and 20 m and the slit widths between 13 and 22 m. In SA, two columns of cells were left unmanaged.

***Table S4.6*** *Spatial layout parameters used in management regime CAB2.*

| | | **Values** | | |
|---|---|---|---|---|
| **Parameter** | **Explanation** | **UM** | **HM** | **SA** |
| $mgmunit_{CL1}$ | Grid column with cable line 1 | 3 | 3 | 4 |
| $mgmunit_{CL2}$ | Grid column with cable line 2 | 8 | 11 | 12 |
| $mgmunit_{SL}$ | Slit length in cells | 2 | 3 | 3 |
| $mgmunit_{SW}$ | Slit width in cells | 2 | 2 | 2 |

### 5. Management regimes SC

The management regimes SC used management type CSC (slit cuts). The applied weighting factors are listed in Table S4.7.

***Table S4.7*** *Weighting factors used in management regimes SC.*

| Parameter | Explanation | Value |
|---|---|---|
| $mgmw_{spc}$ | Weight given to tree species | 0 |
| $mgmw_{BA}$ | Weight given to basal area | 100 |
| $mgmw_{CR}$ | Weight given to crown ratio | 0 |
| $mgmw_{RegUnit}$ | Weight given to regeneration within harvest unit | 40 |
| $mgmw_{RegNeighb}$ | Weight given to regeneration in cells neighboring harvest unit | 20 |

The size of the slits was chosen to be approximately 1.5 tree lengths long and 0.5 tree lengths wide. One tree length, i.e., height, was assumed to be 35 m in UM and HM, and 27 m in SA. The resulting parameter values for the spatial layout are listed in Table S4.8.

***Table S4.8*** *Spatial layout parameters used in management regime CAB2.*

| | | Values | | |
|---|---|---|---|---|
| Parameter | Explanation | UM | HM | SA |
| $mgmunit_{SL}$ | Slit length in cells | 5 | 7 | 6 |
| $mgmunit_{SW}$ | Slit width in cells | 2 | 2 | 2 |
