## Supplementary material S5 for "Management recommendations for Alpine protection forests: the importance of regeneration quality and initial stand composition"

### Supplementary material S5: Model outputs

S5 consists of a series of folders for each simulation with a management regime applied. The results of the corresponding simulation without management (NOM) are included in the graphs. The data can be downloaded as three zip-archives (one per elevational zone) from zenodo: 10.5281/zenodo.10938936 (25.7 GB total)

The **folder names** relate to the individual simulations as follows:

Example: SA[1]_5[2]1[3]_ST[4]_init1[5]_STS[6]_t30[7]_i40[8]

1. elevational zone (either UM, HM, or SA)
2. site quality (either 3 or 5)
3. regeneration quality (either 1, 3, or 5)
4. simulation type (either LT for long-term simulations or ST for short-term simulations)
5. initialization stand type (initLT = long-term simulation; init1 = young stand; init2 = well-structured stand; init3 = mature stand)
6. management type (either CAB1/CAB2 (cable yarding), GRS1/GRS2 (group selection), SC (slit cuts), or STS (single tree selection)
7. intervention interval (either 10, 20, 30, or 40 [years])
8. intervention intensity (either 10, 20, 30, or 40 [% basal area removal])

Each folder contains the following **files**:

- F2_*simulation name*_A.png: NaiS-indices arranged analogous to the NaiS-form 2 for the natural hazard snow avalanches (A)
- F2_*simulation name*_A.png: NaiS-indices arranged analogous to the NaiS-form 2 for the natural hazard landslides, erosion, debris flow (LED)
- Gaps_*simulation name*.jpg: Gaps according to the assessment of the protective function at six time steps during the assessed simulation period.
- Sim_*simulation name*.jpg: Stand characteristics over time for the assessed simulation period
    - Basal area (total and by species)
    - Stem number (total and by species)
    - Canopy cover (total)
    - Average crown ratio of the 100 trees/ha with the largest DBH (per species)
    - DHB-distribution (total) at six time steps
- NaiS_subindices_A / NaiS_subindices_LED: subfolder with detailed subindices and the underlying stand characteristics including threshold values from the NaiS-profiles used for the assessment of the protective function against avalanches (A) and landslides, erosion, debris flow (LED)
    - 0_F2.jpg: NaiS-indices arranged analogous to the NaiS-form 2
    - 1a_mix_stand.jpg: Stand characteristics for the calculation of the index “species mixture”
    - 1b_mix_ind.jpg: Subindices underlying the index “species mixture”
    - 2a_vert_dclass.jpg: Stand characteristics for the calculation of the subindex “dclass” (diameter distribution)
    - 2b_vert_ind.jpg: Subindices underlying the index “vertical structure”
    - 3a_horiz_*A/LED*_stand.jpg: Stand characteristics for the calculation of the index “horizontal structure”
    - 3b_horiz_ind.jpg: Subindices underlying the index “horizontal structure”
    - 4_supptr.jpg: Stand characteristics and subindices underlying the index “stability of support trees”
    - 5_seedl.jpg: Stand characteristics and subindices underlying the index “seedlings”
    - 6a_sapthi_stand.jpg: Stand characteristics for the calculation of the index “saplings and thicket”
    - 6b_sapthi_ind.jpg: Subindices underlying the index “saplings and thicket”
