## Supplementary material S6 for "Management recommendations for Alpine protection forests: the importance of regeneration quality and initial stand composition"

### Supplementary material S6: Protective quality of simulated stands

In the following graphs, the protective quality for simulations with normal *regeneration quality* are graphically displayed (corresponds to the simulations used for the “reduced models”). Figures are divided into two columns for good and medium *site quality* and two rows for the minimal (displayed response variable $sMP_{abs}$) and the ideal profile ($sIP_{abs}$). In each quadrant, the six panels refer to the management types. The color scales are chosen so that the protective quality of the NOM simulation (bottom left point) determines the midpoint (gray), while improvements relative to NOM are displayed in blue and reductions in red. Each figure contains the results of a certain elevational zone, initialization stand / simulation type and natural hazard.

### 1. Upper montane zone

#### 1.1. Short-term simulation – young stand

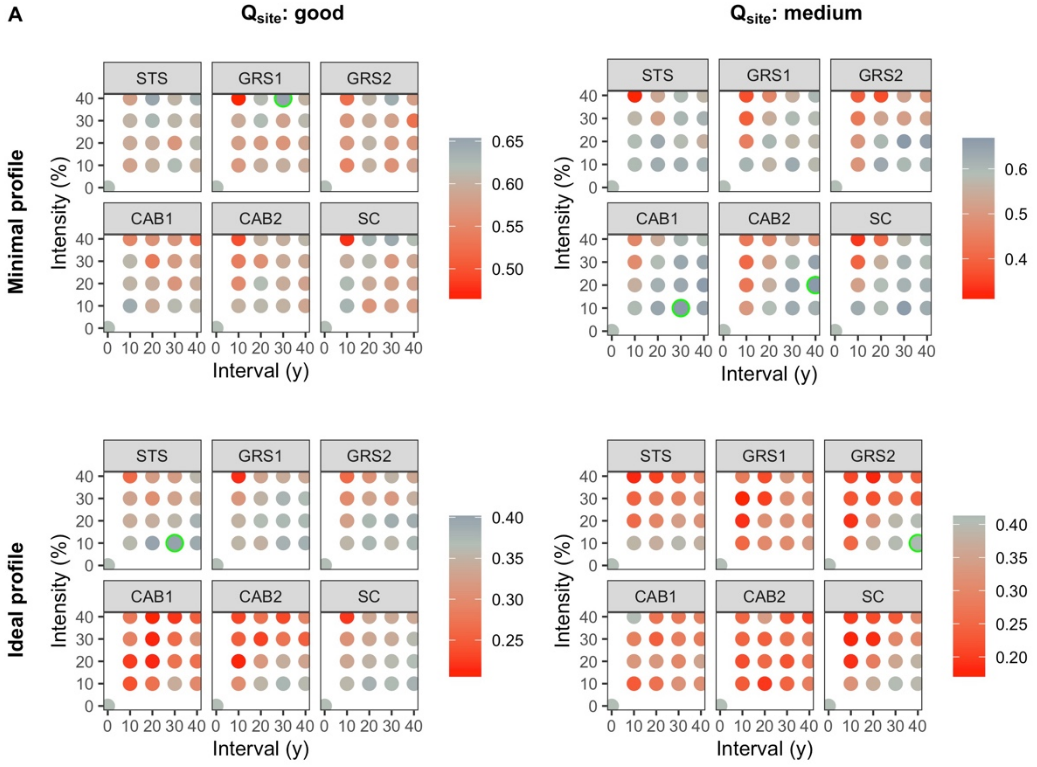


***Fig. S6.1*** *Protective qualities of short-term UM simulations initialized with a young stand and a normal regeneration quality using the NaiS-profile "avalanches".*

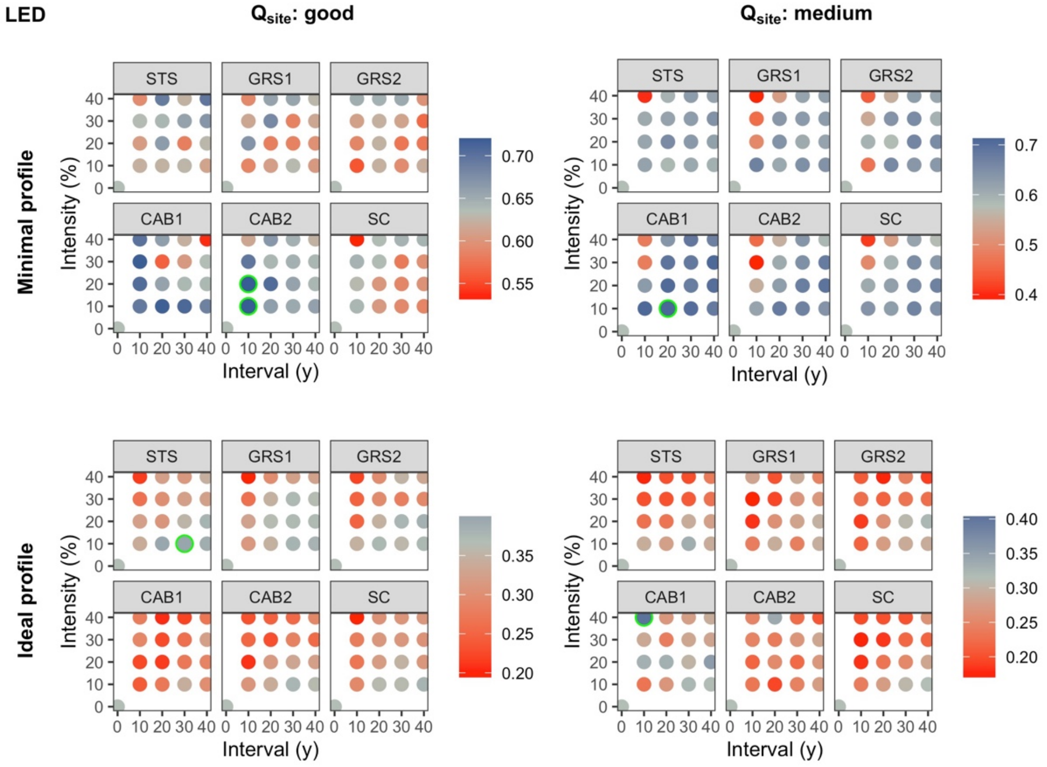


***Fig. S6.2*** *Protective qualities of short-term UM simulations initialized with a young stand and a normal regeneration quality using the NaiS-profile "landslides, erosion, debris flow".*

#### 1.2. Short-term simulation – well-structured stand

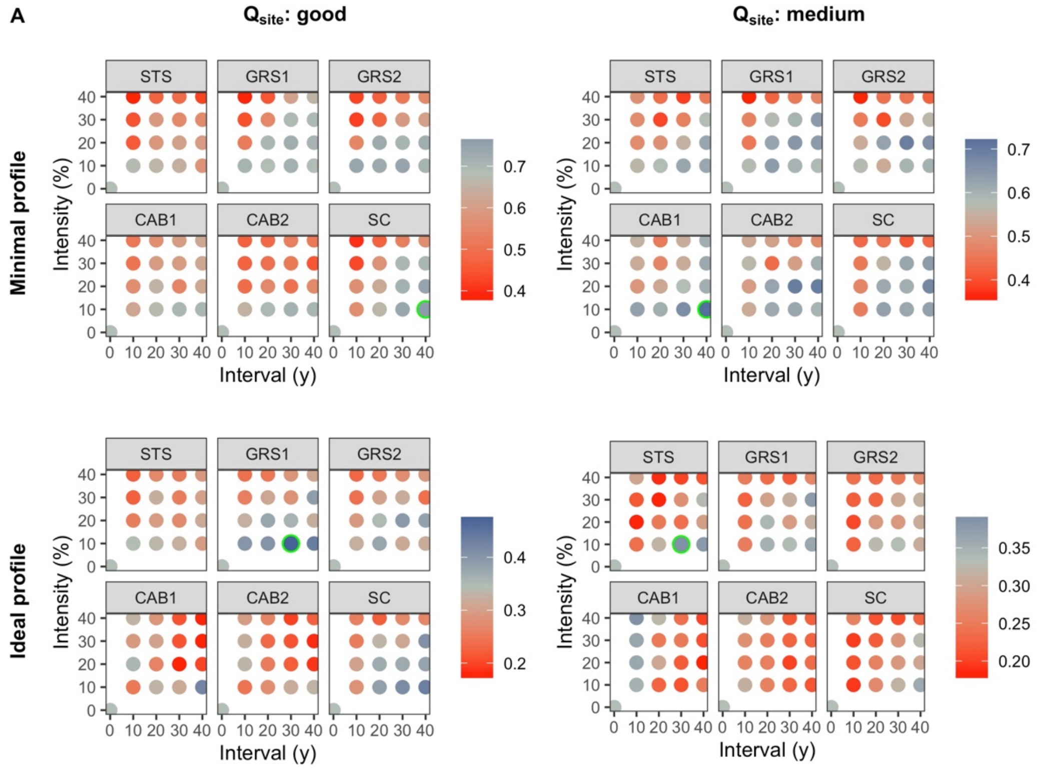


***Fig. S6.3*** *Protective qualities of short-term UM simulations initialized with a well-structured stand and a normal regeneration quality using the NaiS-profile "avalanches".*

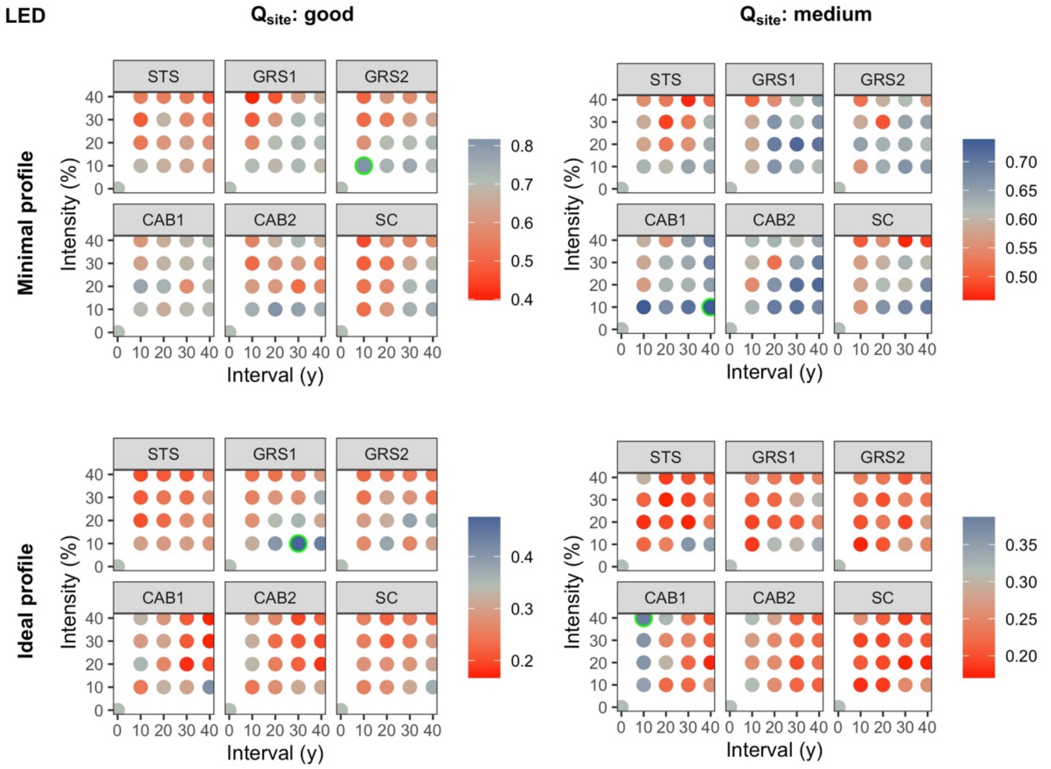


***Fig. S6.4*** *Protective qualities of short-term UM simulations initialized with a well-structured stand and a normal regeneration quality using the NaiS-profile "landslides, erosion, debris flow".*

#### 1.3. Short-term simulation – mature stand

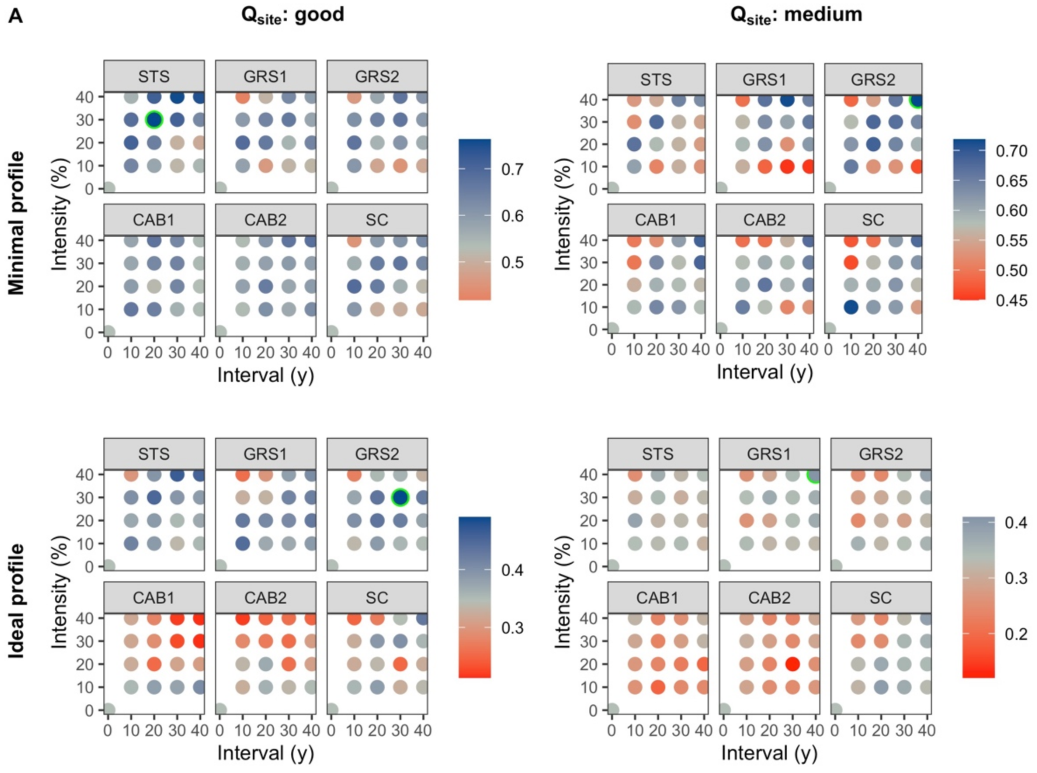


***Fig. S6.5** Protective qualities of short-term UM simulations initialized with a mature stand and a normal regeneration quality using the NaiS-profile "avalanches".*

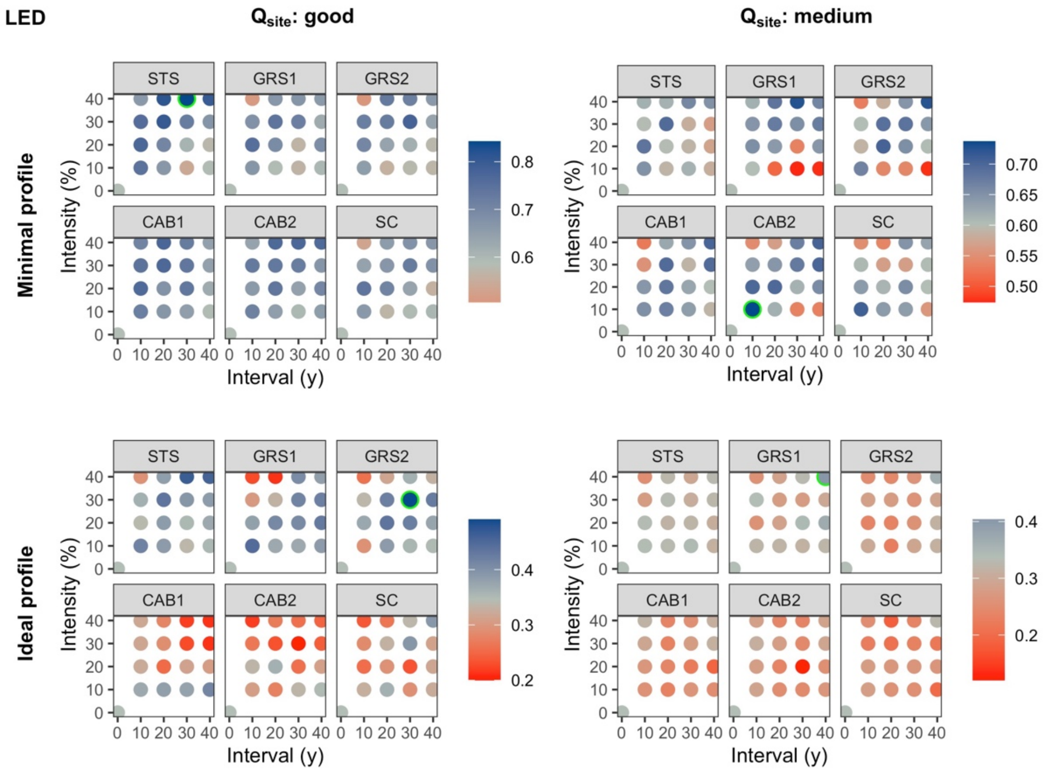


***Fig. S6.6** Protective qualities of short-term UM simulations initialized with a mature stand and a normal regeneration quality using the NaiS-profile "landslides, erosion, debris flow".*

#### 1.4. Long-term simulation

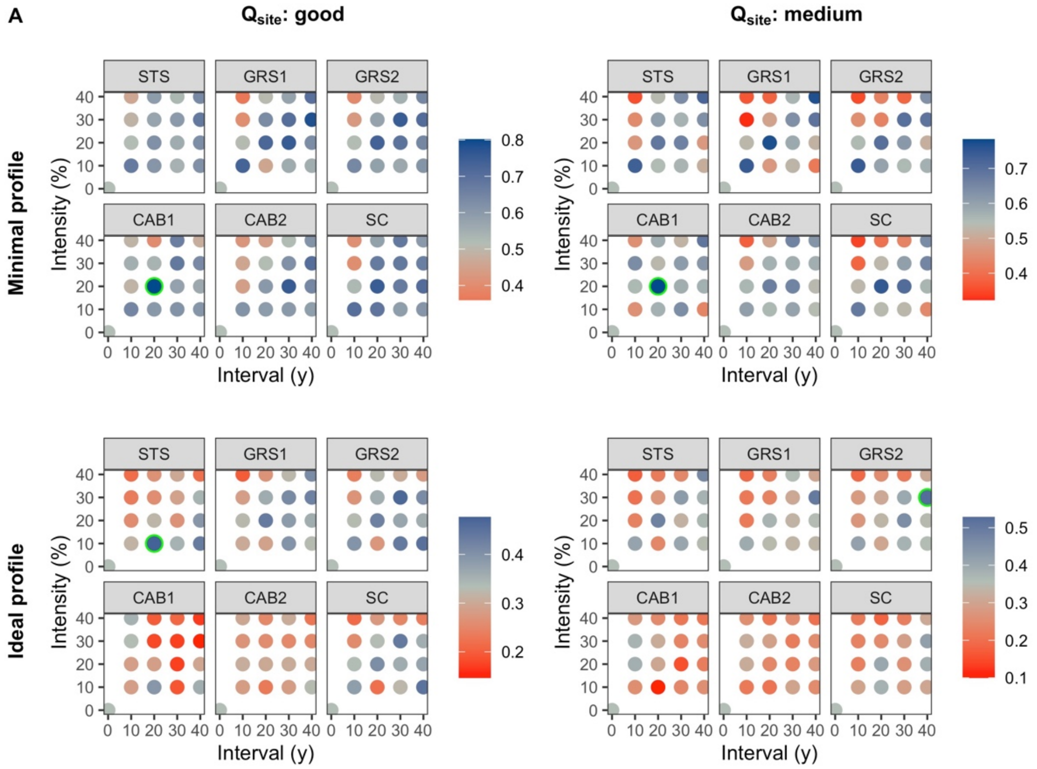


***Fig. S6.7** Protective qualities of long-term UM simulations and a normal regeneration quality using the NaiS-profile "avalanches".*

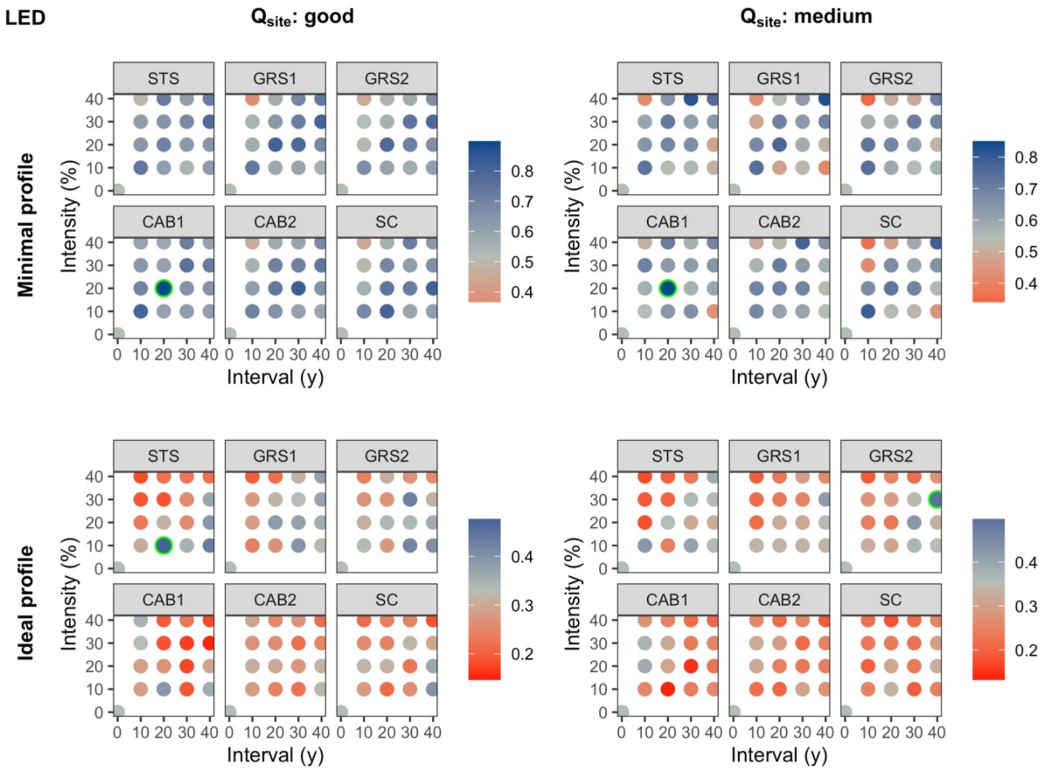


***Fig. S6.8** Protective qualities of long-term UM simulations and a normal regeneration quality using the NaiS-profile "landslides, erosion, debris flow".*

### 2. High montane zone

#### 2.1. Short-term simulation – young stand

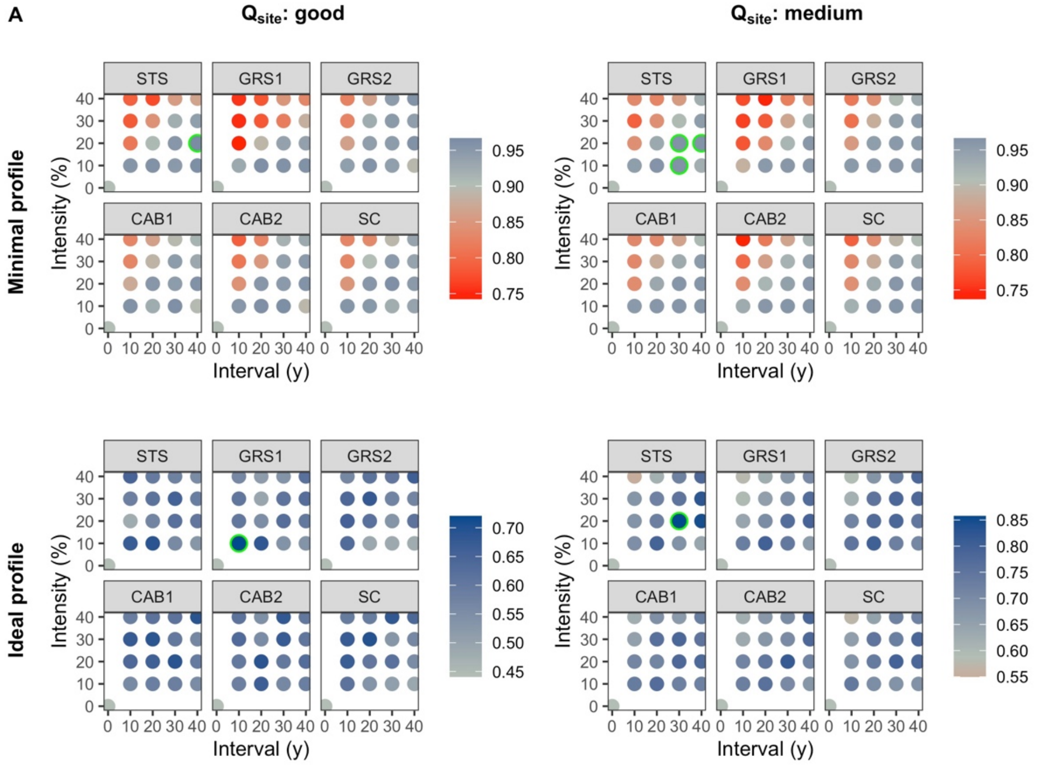


***Fig. S6.9*** *Protective qualities of short-term HM simulations initialized with a young stand and a normal regeneration quality using the NaiS-profile "avalanches".*

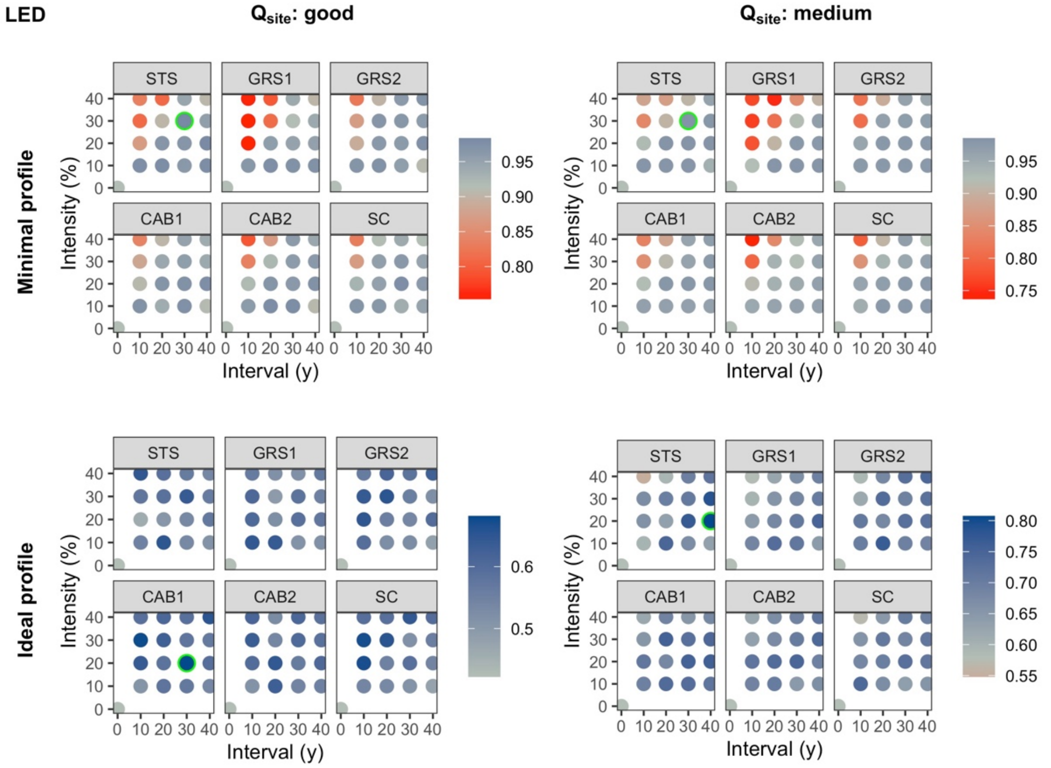


***Fig. S6.10*** *Protective qualities of short-term HM simulations initialized with a young stand and a normal regeneration quality using the NaiS-profile "landslides, erosion, debris flow".*

#### 2.2. Short-term simulation – well-structured stand

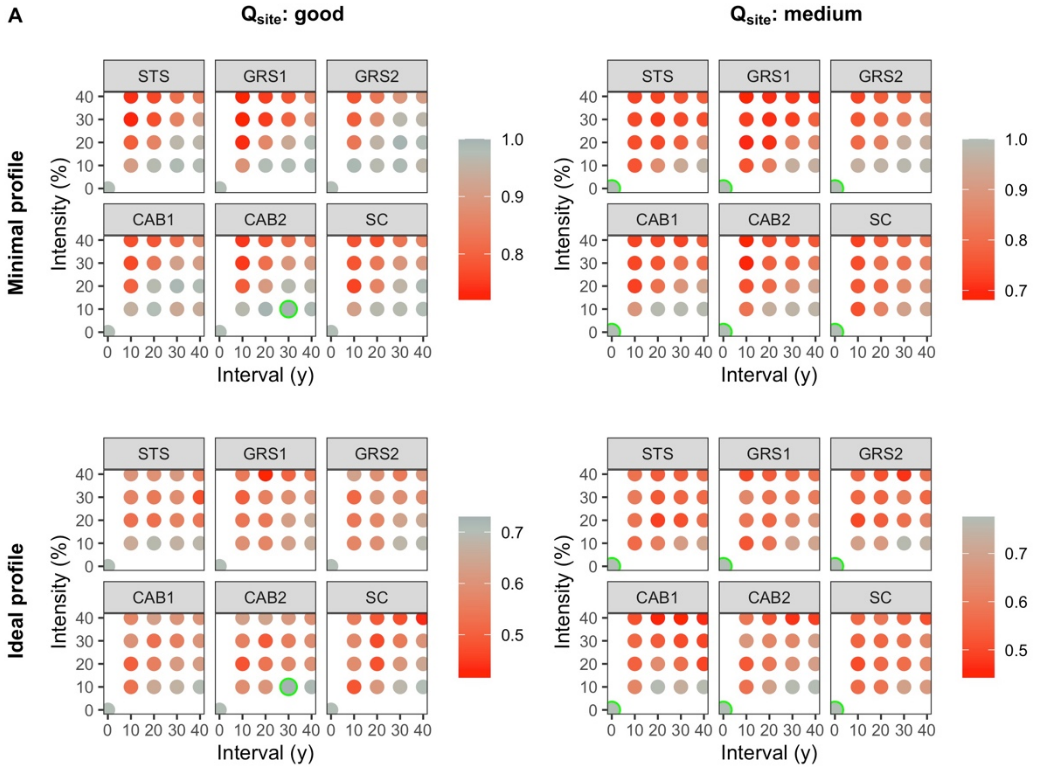


***Fig. S6.11*** *Protective qualities of short-term HM simulations initialized with a well-structured stand and a normal regeneration quality using the NaiS-profile "avalanches".*

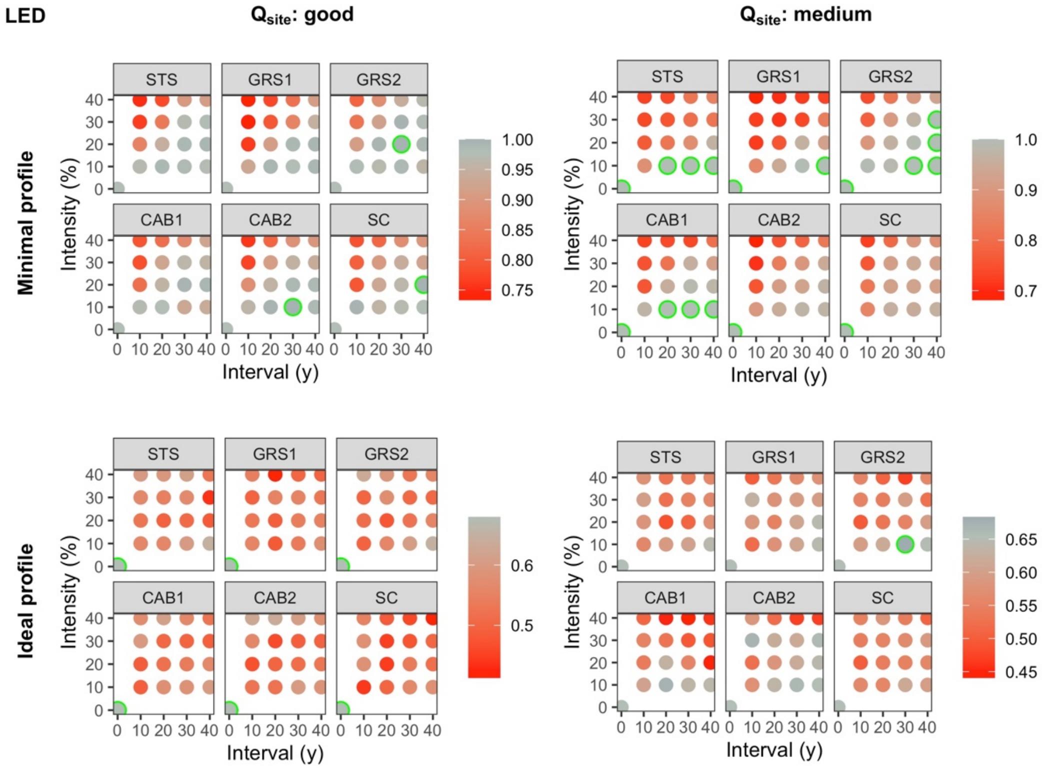


***Fig. S6.12*** *Protective qualities of short-term HM simulations initialized with a well-structured stand and a normal regeneration quality using the NaiS-profile "landslides, erosion, debris flow".*

#### 2.3. Short-term simulation – mature stand

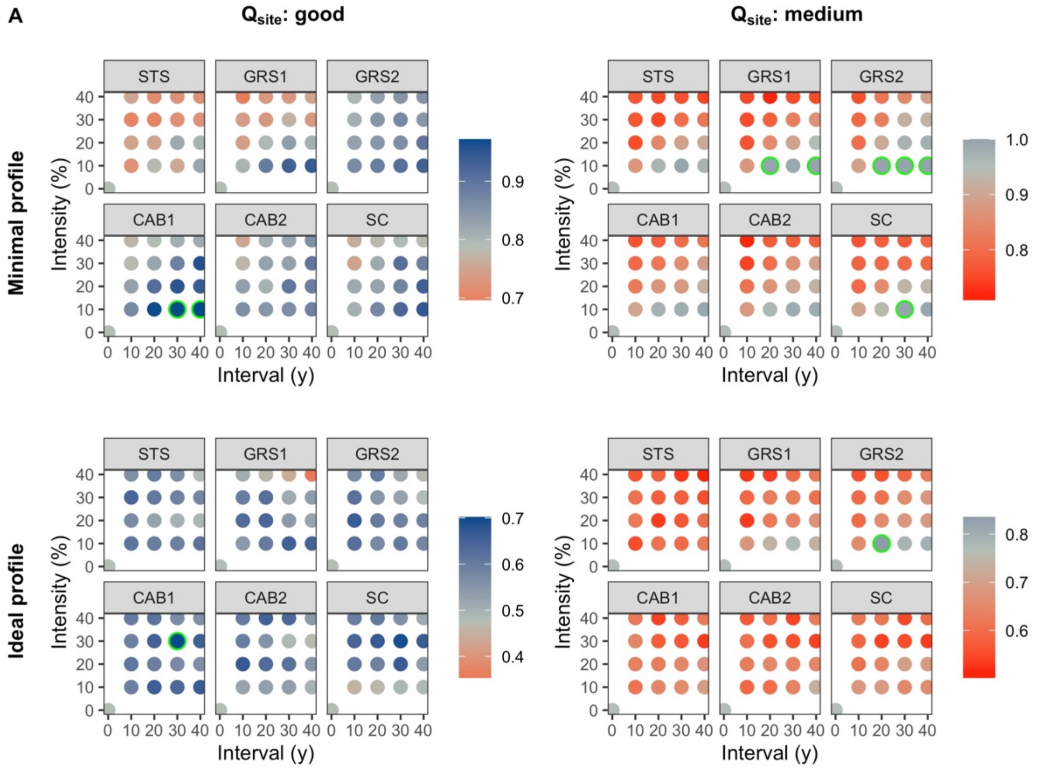


***Fig. S6.13*** *Protective qualities of short-term HM simulations initialized with a mature stand and a normal regeneration quality using the NaiS-profile "avalanches".*

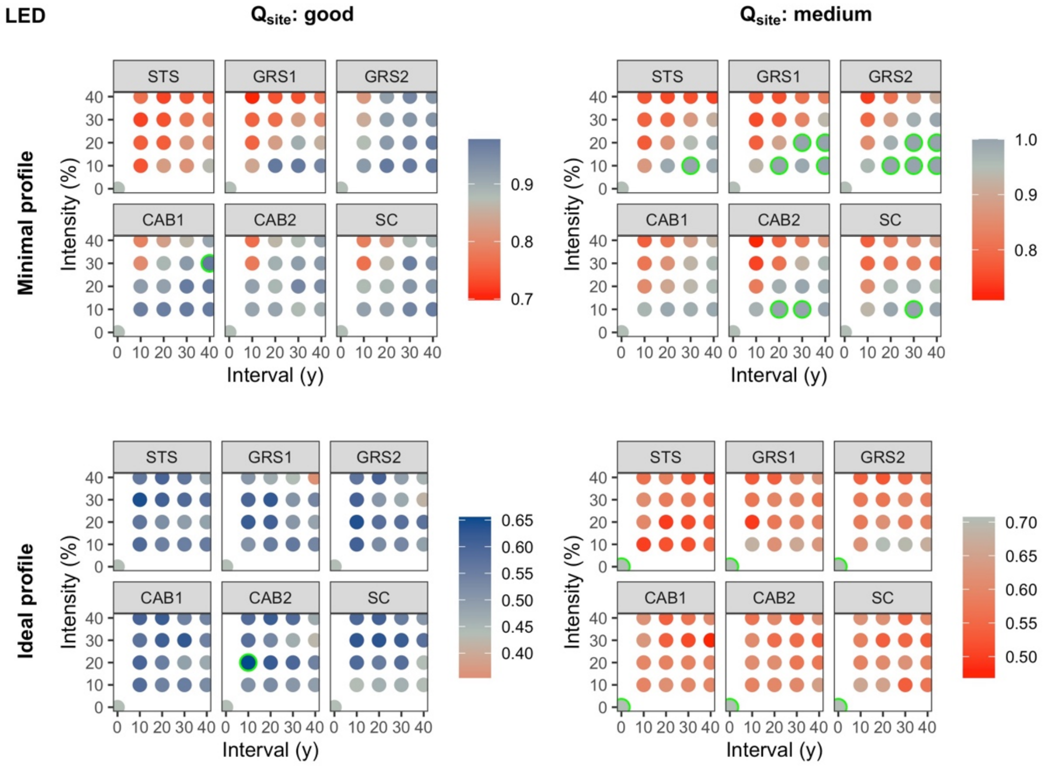


***Fig. S6.14*** *Protective qualities of short-term HM simulations initialized with a mature stand and a normal regeneration quality using the NaiS-profile "landslides, erosion, debris flow".*

#### 2.4. Long-term simulation

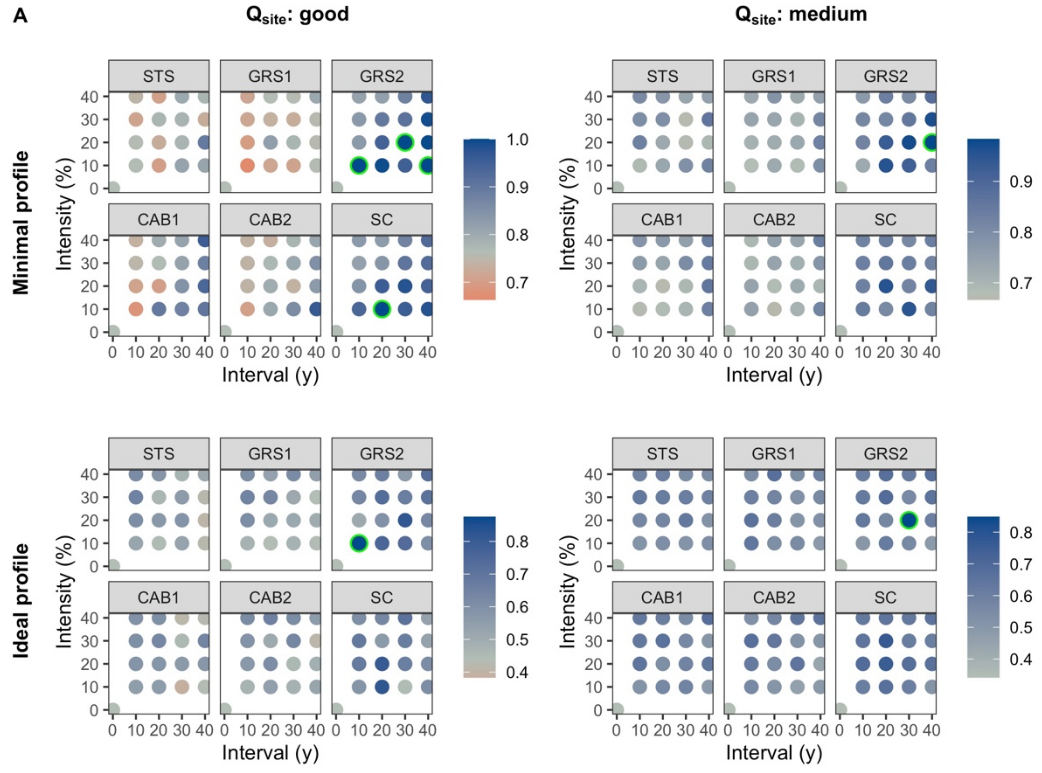


***Fig. S6.15*** *Protective qualities of long-term HM simulations and a normal regeneration quality using the NaiS-profile “avalanches”.*

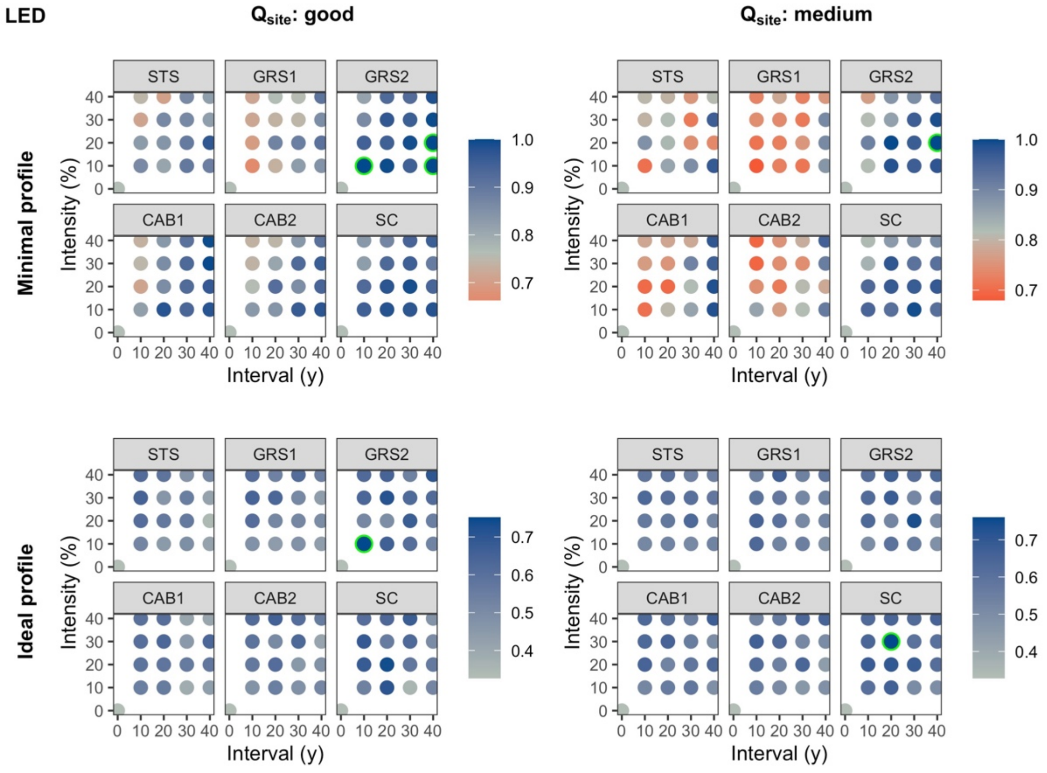


***Fig. S6.16*** *Protective qualities of long-term HM simulations and a normal regeneration quality using the NaiS-profile “landslides, erosion, debris flow”.*

### 3. Subalpine zone

#### 3.1. Short-term simulation – young stand

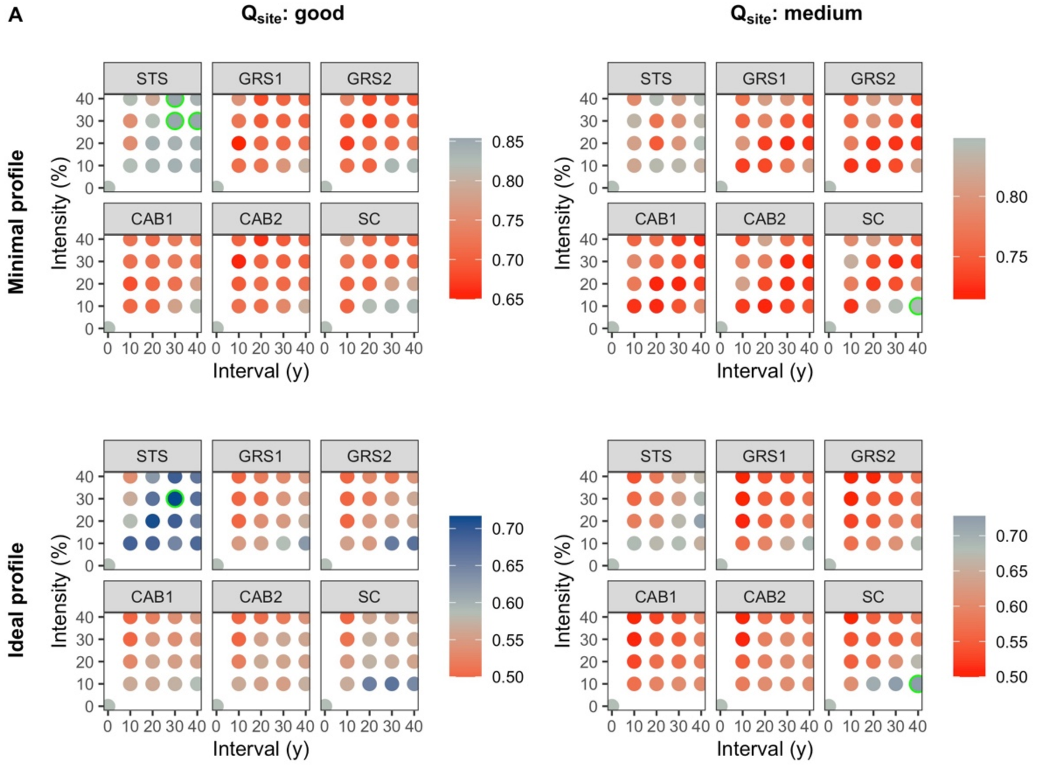


***Fig. S6.17*** *Protective qualities of short-term SA simulations initialized with a young stand and a normal regeneration quality using the NaiS-profile “avalanches”.*

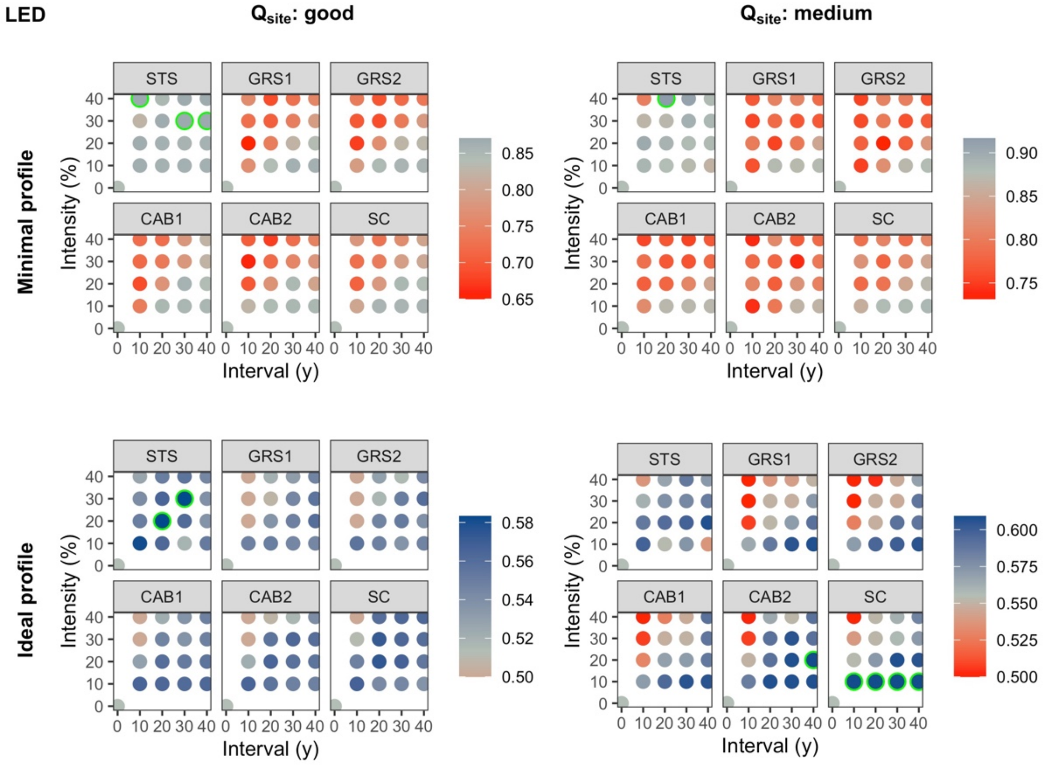


***Fig. S6.18*** *Protective qualities of short-term SA simulations initialized with a young stand and a normal regeneration quality using the NaiS-profile “landslides, erosion, debris flow”.*

#### 3.2. Short-term simulation – well-structured stand

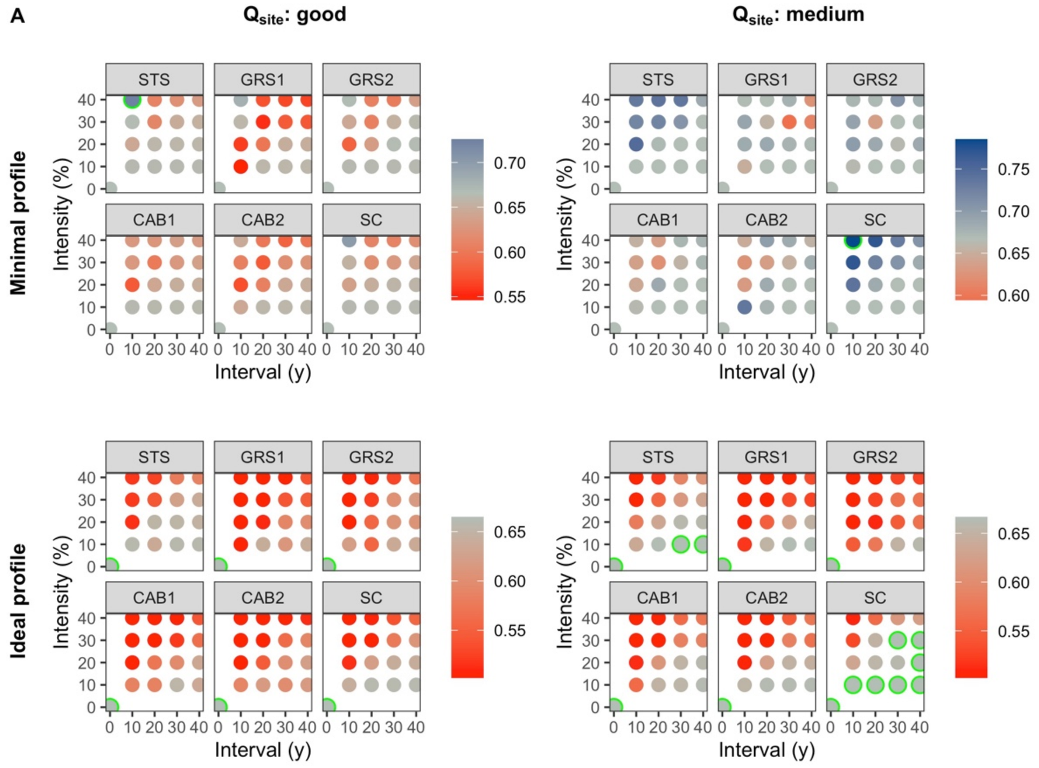


***Fig. S6.19*** *Protective qualities of short-term SA simulations initialized with a well-structured stand and a normal regeneration quality using the NaiS-profile “avalanches”.*

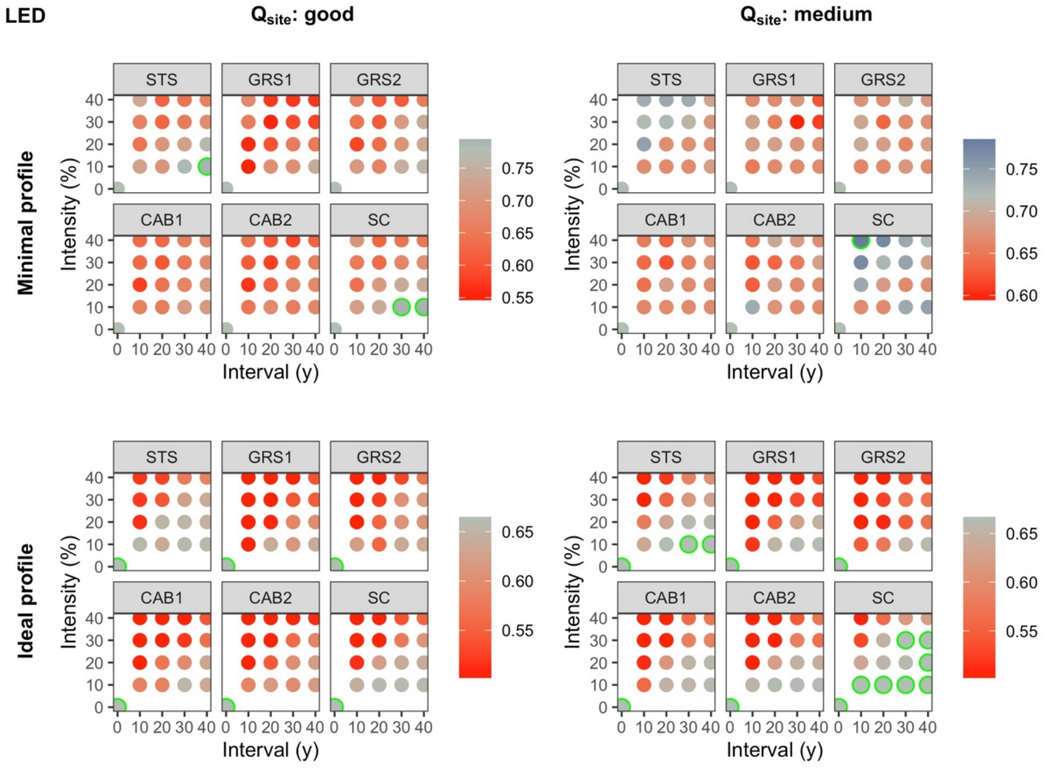


***Fig. S6.20*** *Protective qualities of short-term SA simulations initialized with a well-structured stand and a normal regeneration quality using the NaiS-profile “landslides, erosion, debris flow”.*

#### 3.3. Short-term simulation – mature stand

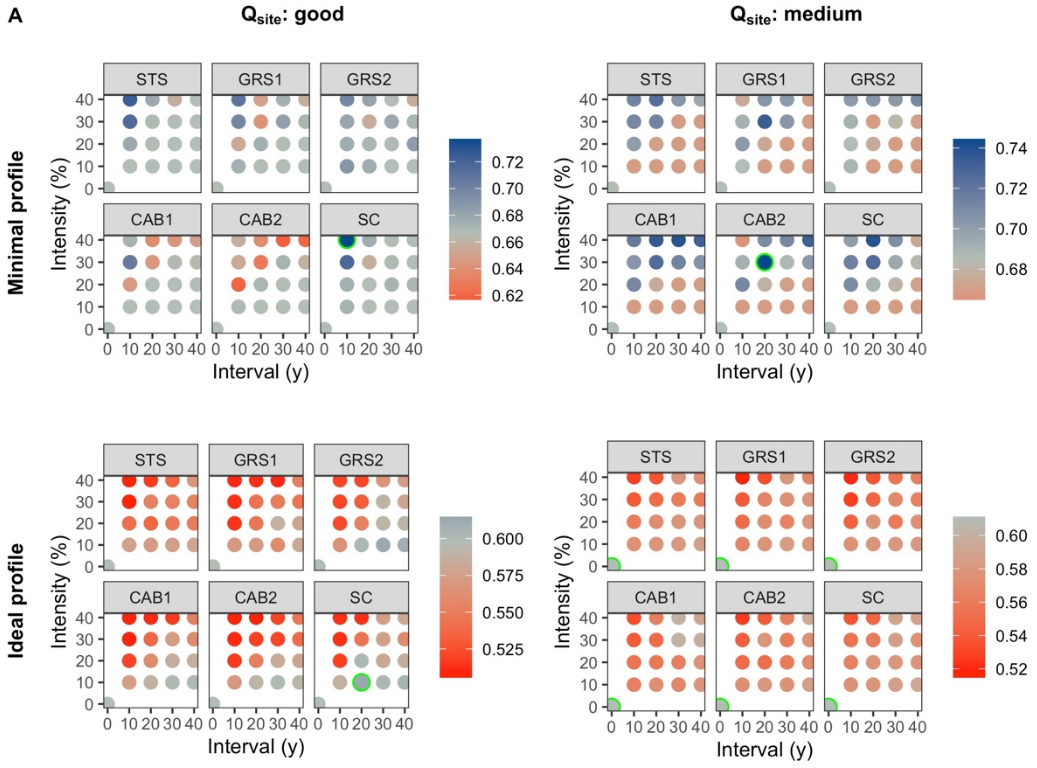


***Fig. S6.21*** *Protective qualities of short-term SA simulations initialized with a mature stand and a normal regeneration quality using the NaiS-profile "avalanches".*

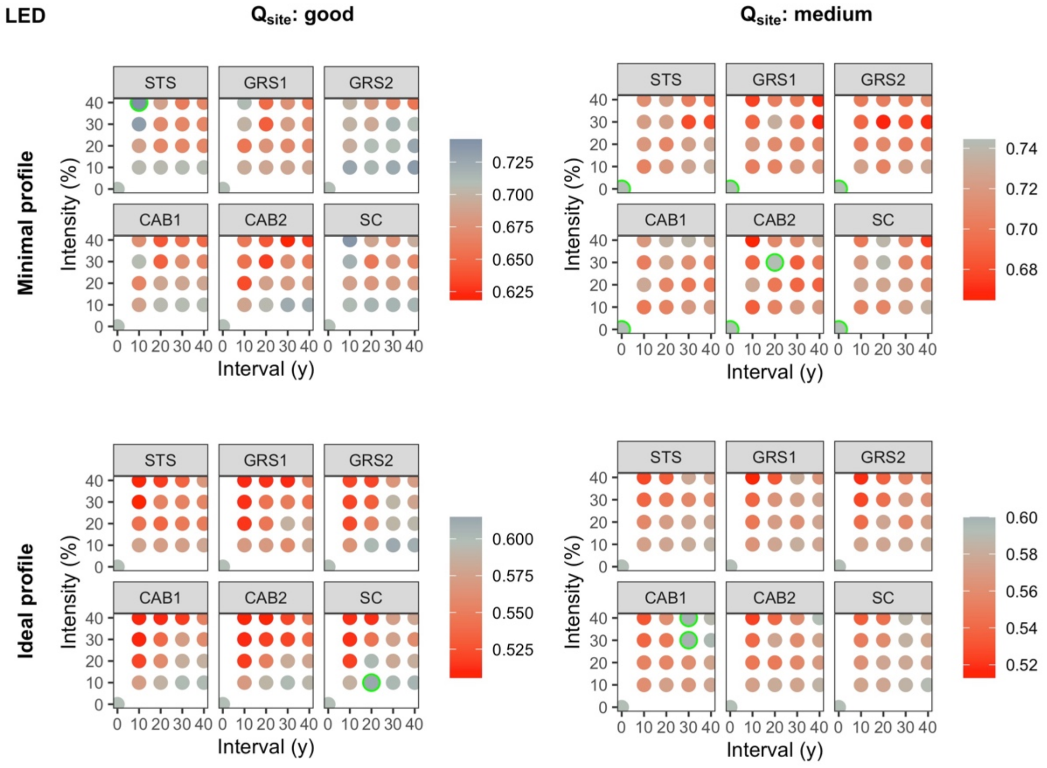


***Fig. S6.22*** *Protective qualities of short-term SA simulations initialized with a mature stand and a normal regeneration quality using the NaiS-profile "landslides, erosion, debris flow".*

##### 3.4. Long-term simulation

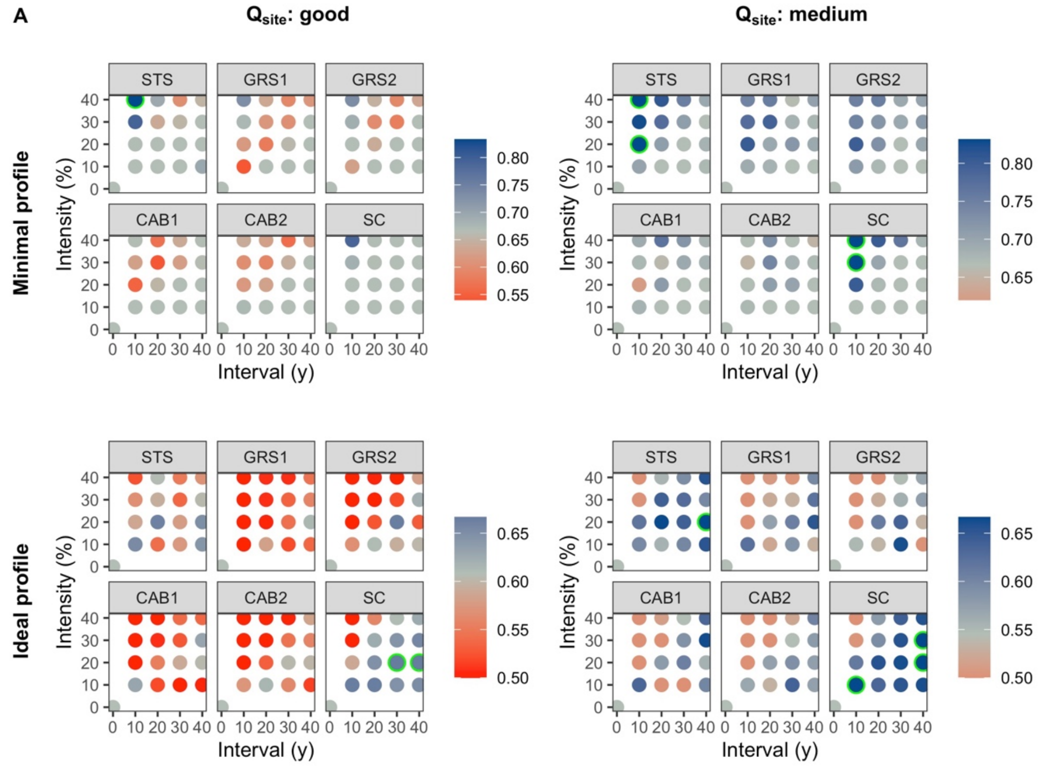


***Fig. S6.23** Protective qualities of long-term SA simulations and a normal regeneration quality using the NaiS-profile "avalanches".*

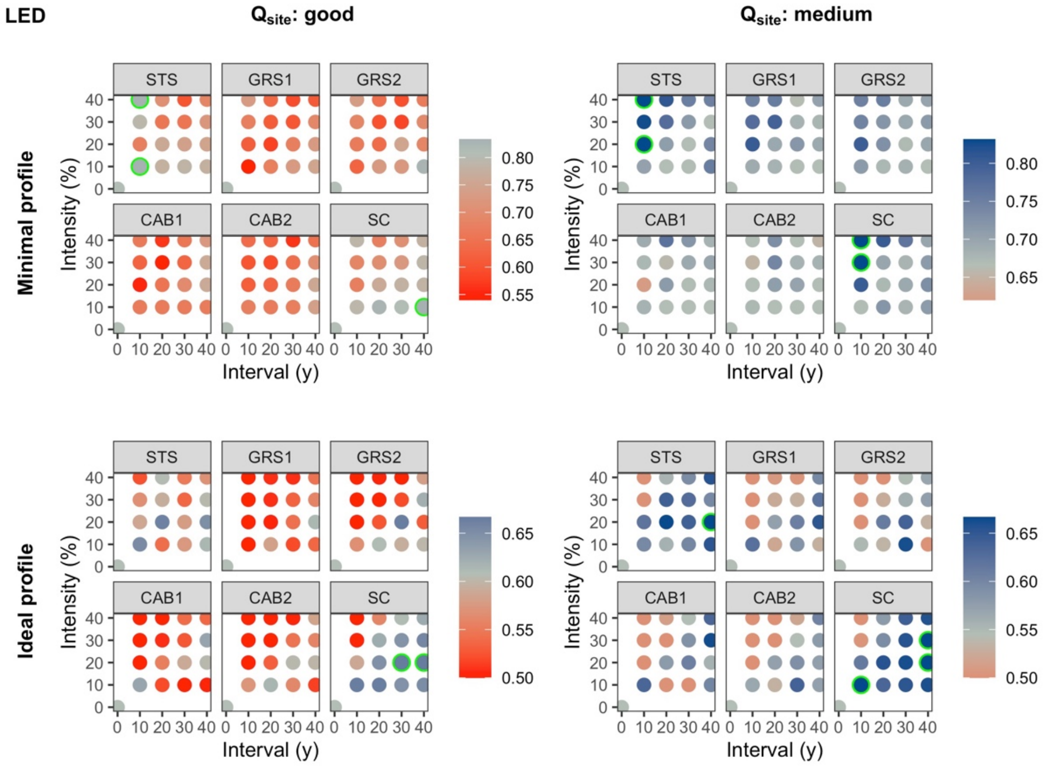


***Fig. S6.24** Protective qualities of long-term SA simulations and a normal regeneration quality using the NaiS-profile "landslides, erosion, debris flow".*
