## Supplementary material S7 for "Management recommendations for Alpine protection forests: the importance of regeneration quality and initial stand composition"

### Supplementary material S7: Further results of statistical models

#### 1. Full models – BRT – Model statistics

***Table S7.1*** *Model details (number of trees and learning rate) and cross-validation statistics (deviance and correlation with mean and standard error, respectively) of the BRT models using all simulations and explanatory variables (full models) by response variable (abs = absolute protective function; diff = difference in protective function compared to NOM), simulation type (LT = long-term; ST = short-term), NaiS-profile type, and natural hazards (A = snow avalanches; LED = landslide/erosion/debris flow).*

| Response variable | Simulation type | NaiS-Profile | Natural hazard | # Trees | Learning rate | Deviance mean | Deviance SE | Correlation mean | Correlation SE |
|---|---|---|---|---|---|---|---|---|---|
| abs | LT | MP | A | 1550 | 0.05 | 2.6E-03 | 1.5E-04 | 0.97 | 1.7E-03 |
| | | | LED | 1950 | 0.04 | 2.7E-03 | 1.1E-04 | 0.97 | 1.3E-03 |
| | | IP | A | 1200 | 0.03 | 2.3E-03 | 1.5E-04 | 0.96 | 3.1E-03 |
| | | | LED | 1200 | 0.03 | 1.9E-03 | 1.5E-04 | 0.97 | 2.7E-03 |
| | ST | MP | A | 1800 | 0.10 | 1.1E-03 | 2.4E-05 | 0.99 | 3.5E-04 |
| | | | LED | 1750 | 0.10 | 1.2E-03 | 3.3E-05 | 0.99 | 5.8E-04 |
| | | IP | A | 1600 | 0.10 | 9.5E-04 | 2.9E-05 | 0.99 | 4.6E-04 |
| | | | LED | 1450 | 0.10 | 7.5E-04 | 1.9E-05 | 0.99 | 2.9E-04 |
| diff | LT | MP | A | 1250 | 0.06 | 2.5E-03 | 1.6E-04 | 0.92 | 4.0E-03 |
| | | | LED | 1900 | 0.04 | 2.7E-03 | 1.1E-04 | 0.93 | 2.4E-03 |
| | | IP | A | 1600 | 0.02 | 2.2E-03 | 1.4E-04 | 0.91 | 7.9E-03 |
| | | | LED | 1550 | 0.02 | 1.9E-03 | 1.4E-04 | 0.93 | 5.7E-03 |
| | ST | MP | A | 3550 | 0.10 | 1.0E-03 | 2.3E-05 | 0.94 | 1.5E-03 |
| | | | LED | 3200 | 0.10 | 1.2E-03 | 3.5E-05 | 0.94 | 1.8E-03 |
| | | IP | A | 3000 | 0.10 | 9.5E-04 | 3.6E-05 | 0.93 | 2.7E-03 |
| | | | LED | 2250 | 0.10 | 7.5E-04 | 2.1E-05 | 0.93 | 1.9E-03 |

#### 2. Full models – Beta regression – Marginal effects

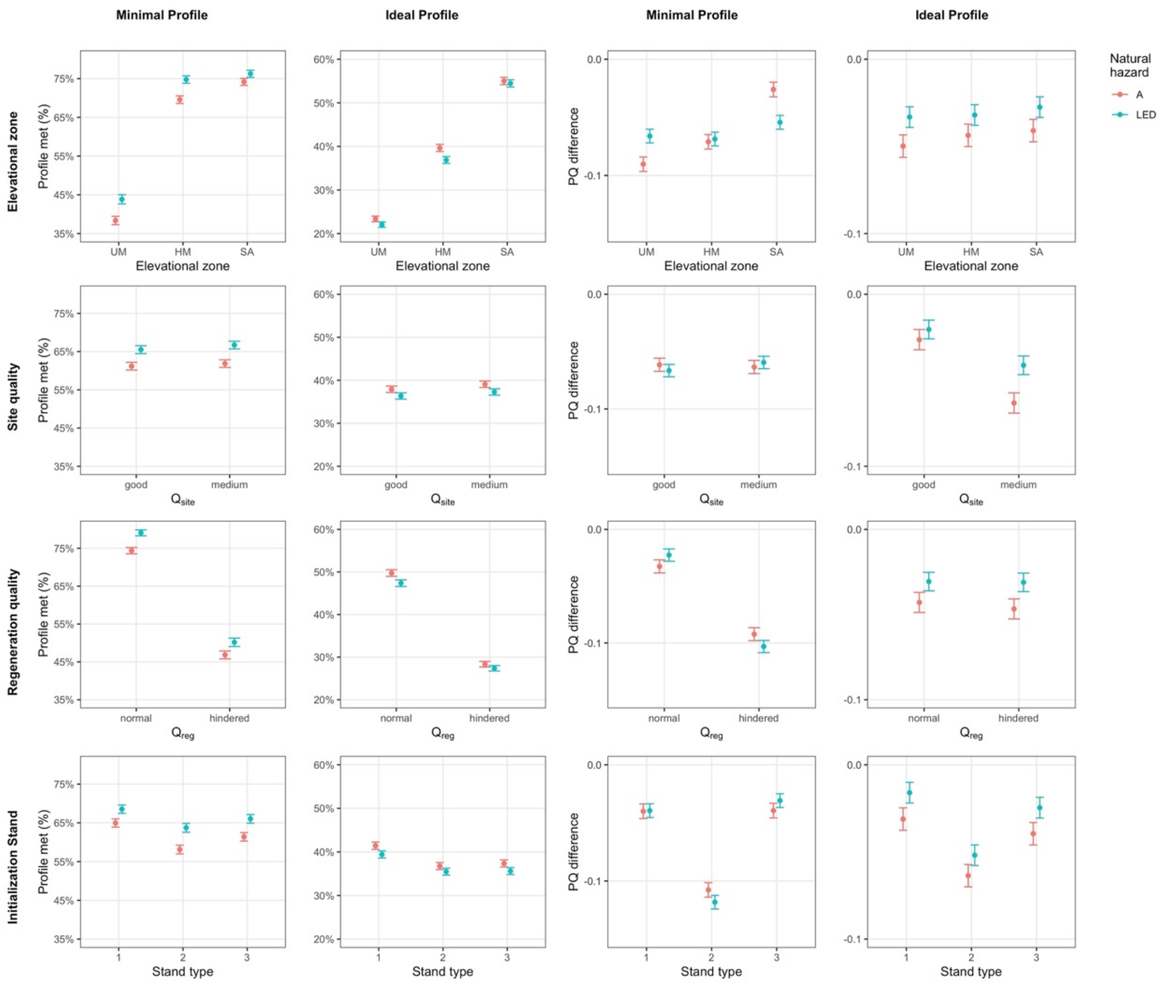


***Fig. S7.1*** *Marginal effects (mean and 95% confidence interval) of the* ***site variables*** *(variables that cannot be influenced by forest management) in beta regression models on* ***short-term*** *simulation results explaining the absolute protective function (left) and effect of management on the protective function (difference in protective function compared to NOM, right) using all simulations and explanatory variables (full models). Bar colors differentiate the natural hazards snow avalanches (A) and landslide/erosion/debris flow (LED).*

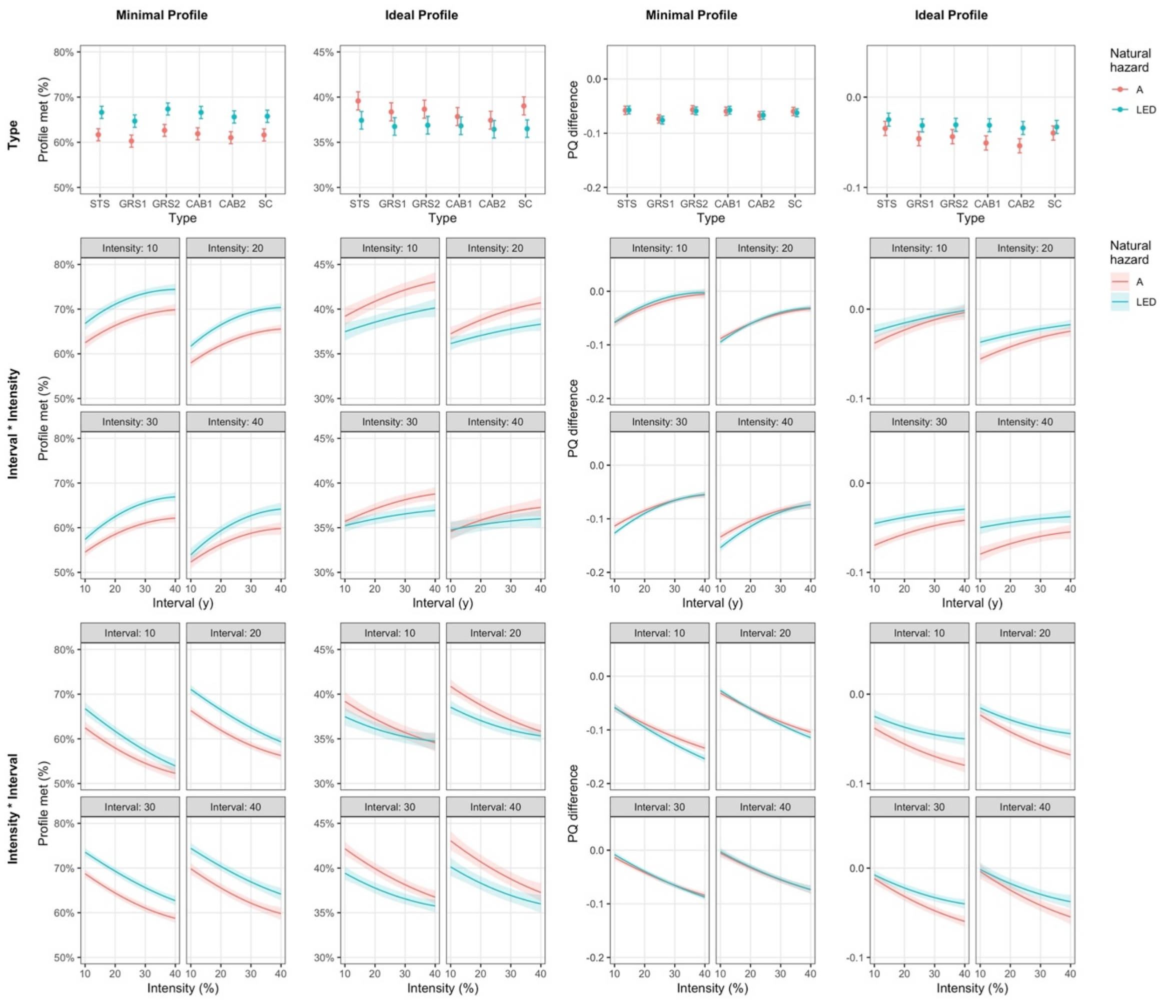


***Fig. S7.2*** *Marginal effects (mean and 95% confidence interval) of the* ***management variables*** *in beta regression models on* ***short-term*** *simulation results explaining the absolute protective function (left) and effect of management on the protective function (difference in protective function compared to NOM, right) using all simulations and explanatory variables (full models). Colors differentiate the natural hazards snow avalanches (A) and landslide/erosion/debris flow (LED).*

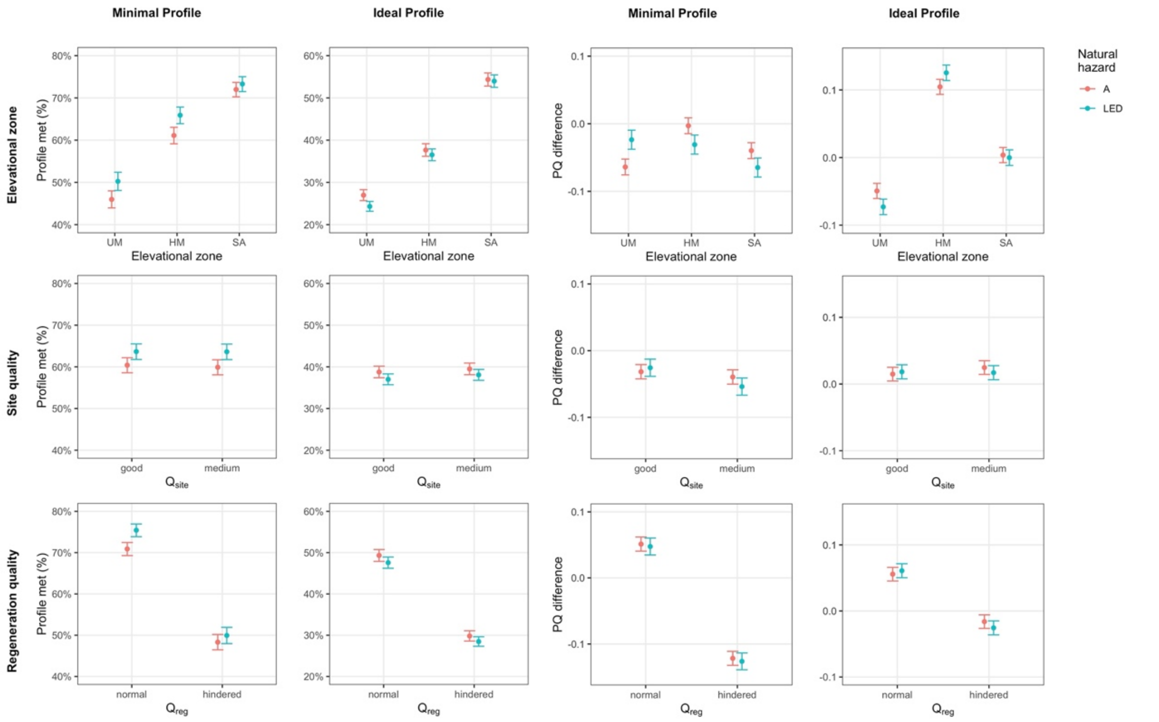


***Fig. S7.3*** *Marginal effects (mean and 95% confidence interval) of the* ***site variables*** *(variables that cannot be influenced by forest management) in beta regression models on* ***long-term*** *simulation results explaining the absolute protective function (left) and effect of management on the protective function (difference in protective function compared to NOM, right) using all simulations and explanatory variables (full models). Bar colors differentiate the natural hazards snow avalanches (A) and landslide/erosion/debris flow (LED).*

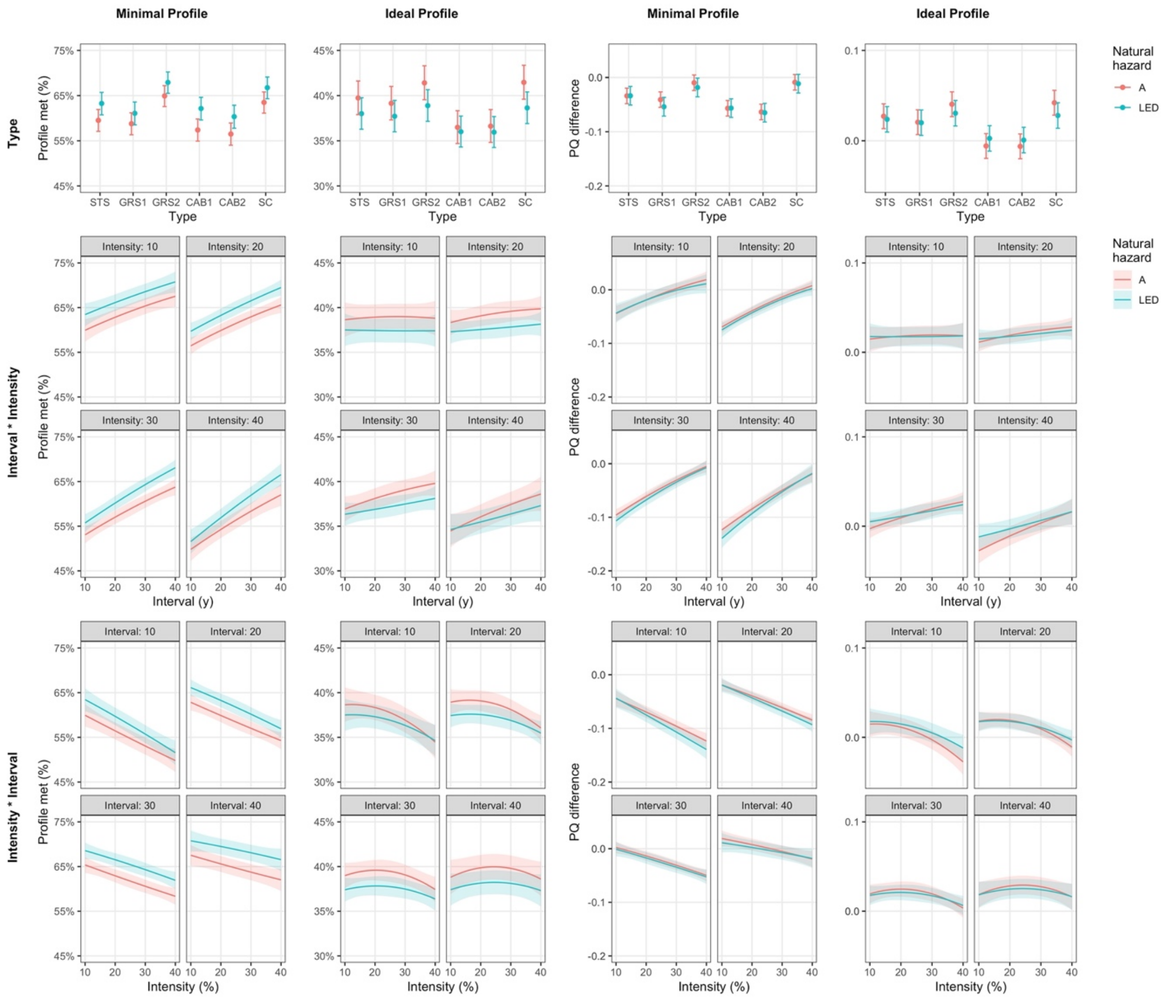


***Fig. S7.4*** *Marginal effects (mean and 95% confidence interval) of the* ***management variables*** *in beta regression models on* ***long-term*** *simulation results explaining the absolute protective function (left) and effect of management on the protective function (difference in protective function compared to NOM, right) using all simulations and explanatory variables (full models). Colors differentiate the natural hazards snow avalanches (A) and landslide/erosion/debris flow (LED).*

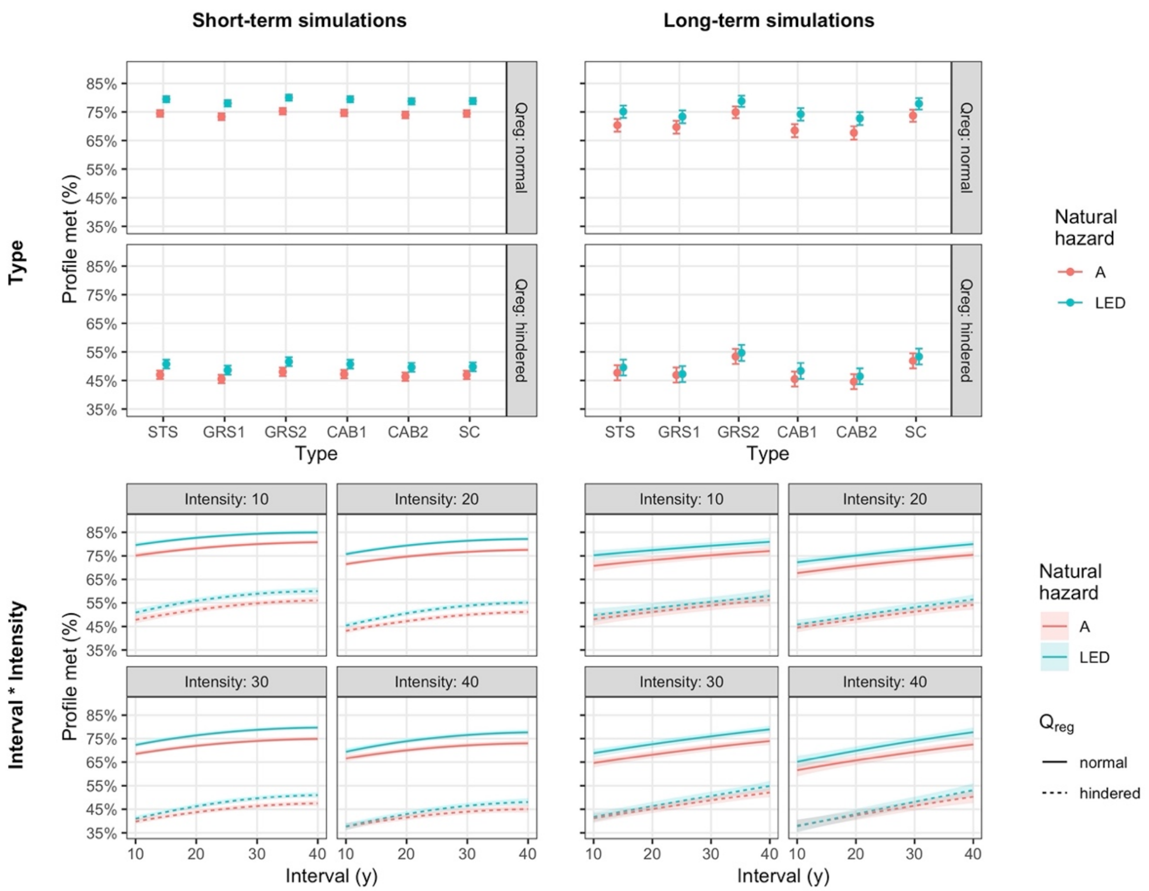


***Fig. S7.5*** *Marginal effects (mean and 95% confidence interval) of **management type** (upper row) and the interaction of **interval and intensity** (lower row) in beta regression models explaining the **absolute protective function** assessed by the **minimal profile** using all explanatory variables (full models). The natural hazards snow avalanches (A) and landslide/erosion/debris flow (LED) are differentiated by color, the two levels of regeneration quality ($Q_{reg}$) by panels in the upper row and by line type in the lower row.*

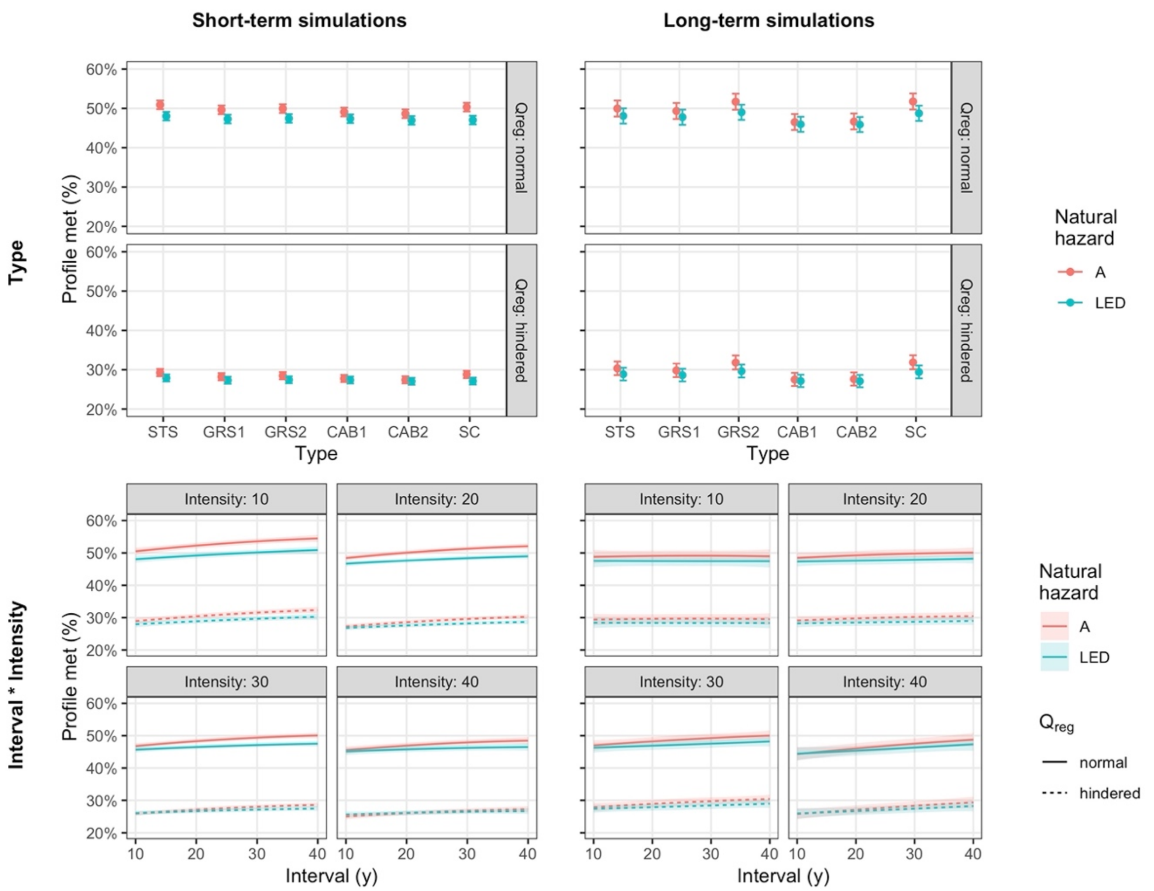


***Fig. S7.6*** *Marginal effects (mean and 95% confidence interval) of* ***management type*** *(upper row) and the interaction of* ***interval and intensity*** *(lower row) in beta regression models explaining the* ***absolute protective function*** *assessed by the* ***ideal profile*** *using all explanatory variables (full models). The natural hazards snow avalanches (A) and landslide/erosion/debris flow (LED) are differentiated by color, the two levels of regeneration quality ($Q_{reg}$) by panels in the upper row and by line type in the lower row.*

### 3. Full models – Beta regression – Goodness of fit

***Table S7.2*** *Pseudo-$R^2$ of the beta regression models using all simulations and explanatory variables (full models) by response variable (abs = absolute protective function; diff = difference in protective function compared to NOM), simulation type (LT = long-term; ST = short-term), NaiS-profile type, and natural hazards (A = snow avalanches; LED = landslide/erosion/debris flow). Colors visualize the pseudo-$R^2$ values with dark green corresponding to the highest and dark red to the lowest values.*

| | | Pseudo $R^2$ | | | |
|---|---|---|---|---|---|
| Response | Simulation | Minimal profile | | Ideal profile | |
| variable | type | A | LED | A | LED |
| abs | LT | 0.48 | 0.48 | 0.67 | 0.71 |
| | ST | 0.64 | 0.58 | 0.76 | 0.76 |
| diff | LT | 0.60 | 0.52 | 0.51 | 0.58 |
| | ST | 0.41 | 0.49 | 0.16 | 0.10 |

### 4. Reduced models – BRT – Variable importance

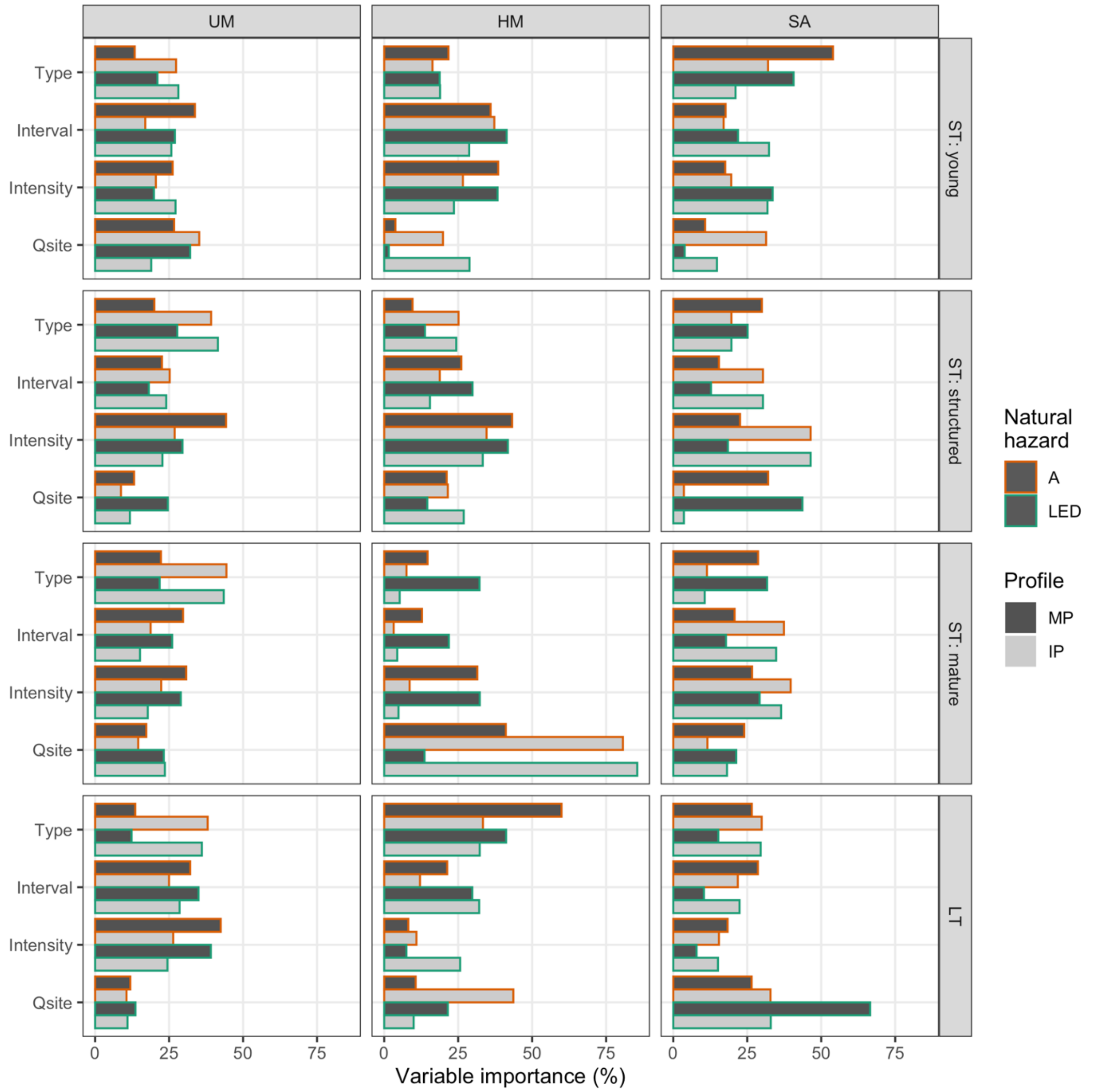


***Fig S7.7*** *Variable importance in BRT explaining the effect of management on the protective function (difference in protective function compared to NOM) using the reduced set of simulations and explanatory variables (reduced models) by elevational zone (panel columns) and simulation type and initialization stand type (ST = short-term; LT = long-term; panel rows). Bar colors differentiate minimal profile (MP) and ideal profile (IP), outline colors the natural hazards snow avalanches (A) and landslide/erosion/debris flow (LED). Abbreviations of explanatory variables: Type = management type; Qsite = site quality.*

### 5. Reduced models – BRT – Model statistics

***Table S7.3*** *Model details (number of trees and learning rate) and cross-validation statistics (deviance and correlation with mean and standard error, respectively) of the BRT models the reduced set of simulations and explanatory variables (reduced models) by elevational zone, simulation type and initialization stand type (ST = short-term; LT = long-term), NaiS-profile type, and natural hazards (A = snow avalanches; LED = landslide/erosion/debris flow).*

| Elevational zone | Simulation type / Init. stand type | NaiS-Profile | Natural hazard | # Trees | Learning rate | Deviance mean | Deviance SE | Correlation mean | Correlation SE |
|---|---|---|---|---|---|---|---|---|---|
| UM | ST: young | MP | A | 1100 | 0.010 | 1.6E-03 | 1.9E-04 | 0.78 | 3.7E-02 |
| | | | LED | 2350 | 0.005 | 1.9E-03 | 2.8E-04 | 0.73 | 3.8E-02 |
| | | IP | A | 1400 | 0.007 | 1.6E-03 | 3.0E-04 | 0.82 | 3.2E-02 |
| | | | LED | 1250 | 0.010 | 1.2E-03 | 2.3E-04 | 0.71 | 6.2E-02 |
| | ST: structured | MP | A | 2050 | 0.010 | 2.7E-03 | 2.8E-04 | 0.86 | 1.3E-02 |
| | | | LED | 1150 | 0.010 | 3.1E-03 | 1.9E-04 | 0.75 | 4.8E-02 |
| | | IP | A | 1850 | 0.007 | 1.7E-03 | 2.3E-04 | 0.74 | 3.5E-02 |
| | | | LED | 1800 | 0.010 | 1.4E-03 | 2.7E-04 | 0.73 | 3.4E-02 |
| | ST: mature | MP | A | 1550 | 0.010 | 2.8E-03 | 2.6E-04 | 0.71 | 2.5E-02 |
| | | | LED | 1750 | 0.010 | 2.3E-03 | 1.9E-04 | 0.76 | 2.2E-02 |
| | | IP | A | 1300 | 0.100 | 1.7E-03 | 1.5E-04 | 0.75 | 2.7E-02 |
| | | | LED | 2250 | 0.007 | 1.9E-03 | 1.2E-04 | 0.69 | 4.0E-02 |
| | LT | MP | A | 1350 | 0.010 | 5.9E-03 | 4.9E-04 | 0.70 | 4.3E-02 |
| | | | LED | 1400 | 0.008 | 5.7E-03 | 5.0E-04 | 0.68 | 4.8E-02 |
| | | IP | A | 1900 | 0.003 | 4.1E-03 | 4.9E-04 | 0.63 | 4.2E-02 |
| | | | LED | 1000 | 0.009 | 3.4E-03 | 3.4E-04 | 0.61 | 3.6E-02 |
| HM | ST: young | MP | A | 2150 | 0.100 | 4.5E-04 | 5.1E-05 | 0.95 | 5.5E-03 |
| | | | LED | 1450 | 0.010 | 8.4E-04 | 2.7E-04 | 0.91 | 1.5E-02 |
| | | IP | A | 1350 | 0.010 | 2.1E-03 | 1.9E-04 | 0.63 | 5.8E-02 |
| | | | LED | 1250 | 0.008 | 1.8E-03 | 1.5E-04 | 0.60 | 6.0E-02 |
| | ST: structured | MP | A | 1350 | 0.007 | 1.1E-03 | 1.7E-04 | 0.95 | 9.3E-03 |
| | | | LED | 1000 | 0.010 | 1.3E-03 | 1.5E-04 | 0.93 | 9.1E-03 |
| | | IP | A | 2650 | 0.030 | 2.2E-03 | 2.0E-04 | 0.81 | 1.9E-02 |
| | | | LED | 1550 | 0.010 | 1.7E-03 | 2.4E-04 | 0.73 | 3.6E-02 |
| | ST: mature | MP | A | 1600 | 0.010 | 8.5E-04 | 7.0E-05 | 0.96 | 5.9E-03 |
| | | | LED | 3450 | 0.100 | 1.1E-03 | 1.0E-04 | 0.93 | 9.1E-03 |
| | | IP | A | 1350 | 0.010 | 2.5E-03 | 3.1E-04 | 0.93 | 1.1E-02 |
| | | | LED | 1150 | 0.010 | 2.0E-03 | 2.1E-04 | 0.93 | 9.3E-03 |
| | LT | MP | A | 1300 | 0.005 | 2.3E-03 | 1.8E-04 | 0.83 | 2.3E-02 |
| | | | LED | 1400 | 0.007 | 2.7E-03 | 3.1E-04 | 0.87 | 1.8E-02 |
| | | IP | A | 1250 | 0.003 | 4.8E-03 | 6.4E-04 | 0.75 | 2.0E-02 |
| | | | LED | 1050 | 0.003 | 4.2E-03 | 5.1E-04 | 0.59 | 4.1E-02 |
| SA | ST: young | MP | A | 1800 | 0.007 | 8.5E-04 | 7.9E-05 | 0.78 | 3.8E-02 |
| | | | LED | 1000 | 0.005 | 8.1E-04 | 7.6E-05 | 0.87 | 2.0E-02 |
| | | IP | A | 2450 | 0.010 | 5.0E-04 | 6.3E-05 | 0.95 | 7.7E-03 |
| | | | LED | 1900 | 0.020 | 1.9E-04 | 1.8E-05 | 0.88 | 1.8E-02 |
| | ST: structured | MP | A | 2300 | 0.010 | 6.6E-04 | 7.6E-05 | 0.73 | 4.8E-02 |
| | | | LED | 1700 | 0.010 | 1.1E-03 | 1.3E-04 | 0.84 | 3.1E-02 |
| | | IP | A | 1150 | 0.010 | 6.8E-04 | 6.7E-05 | 0.92 | 7.8E-03 |
| | | | LED | 1150 | 0.010 | 6.8E-04 | 6.7E-05 | 0.92 | 7.8E-03 |
| | ST: mature | MP | A | 1700 | 0.010 | 2.7E-04 | 3.7E-05 | 0.63 | 4.2E-02 |
| | | | LED | 1050 | 0.010 | 3.5E-04 | 4.3E-05 | 0.60 | 4.8E-02 |
| | | IP | A | 1300 | 0.010 | 1.6E-04 | 1.3E-05 | 0.87 | 1.4E-02 |
| | | | LED | 1400 | 0.009 | 1.6E-04 | 1.4E-05 | 0.88 | 1.3E-02 |
| | LT | MP | A | 1450 | 0.010 | 1.2E-03 | 1.4E-04 | 0.76 | 3.2E-02 |
| | | | LED | 1400 | 0.009 | 1.9E-03 | 1.7E-04 | 0.90 | 9.3E-03 |
| | | IP | A | 1150 | 0.010 | 1.5E-03 | 1.3E-04 | 0.82 | 2.2E-02 |
| | | | LED | 1150 | 0.010 | 1.5E-03 | 1.3E-04 | 0.83 | 2.1E-02 |

### 6. Reduced models – Beta regression – Goodness of fit

***Table S7.4*** *Pseudo-$R^2$ of the beta regression models explaining the effect of management on the protective function (difference in protective function compared to NOM) using the reduced set of simulations and explanatory variables (reduced models) by elevational zone, simulation type and initialization stand type (ST = short-term; LT = long-term), NaiS-profile type, and natural hazards (A = snow avalanches; LED = landslide/erosion/debris flow). Colors visualize the pseudo-$R^2$ values with dark green corresponding to the highest and dark red to the lowest values.*

| | | Pseudo $R^2$ | | | |
|---|---|---|---|---|---|
| | | Minimal profile | | Ideal profile | |
| Elevational zone | Simulation type / Init. stand type | A | LED | A | LED |
| UM | ST: young | 0.46 | 0.39 | 0.63 | 0.38 |
| | ST: structured | 0.67 | 0.60 | 0.33 | 0.28 |
| | ST: mature | 0.47 | 0.55 | 0.50 | 0.49 |
| | LT | 0.55 | 0.53 | 0.39 | 0.34 |
| HM | ST: young | 0.81 | 0.77 | 0.32 | 0.33 |
| | ST: structured | 0.87 | 0.83 | 0.62 | 0.45 |
| | ST: mature | 0.87 | 0.78 | 0.83 | 0.85 |
| | LT | 0.66 | 0.74 | 0.57 | 0.41 |
| SA | ST: young | 0.58 | 0.70 | 0.86 | 0.67 |
| | ST: structured | 0.49 | 0.65 | 0.79 | 0.79 |
| | ST: mature | 0.24 | 0.35 | 0.69 | 0.72 |
| | LT | 0.48 | 0.76 | 0.73 | 0.73 |
